# A new class of lipid transfer proteins is required for the recycling of lipids from the *P. falciparum* digestive vacuole

**DOI:** 10.64898/2026.09.11.750981

**Authors:** Andrés Guillén-Samander, Yara Ahmed, Ana Ribeiro-Holbein, José Cubillán-Marín, Katharina Höhn, Cristian Rocha-Roa, Jana Dröge, Lea S. Panitzsch, Paul-Christian Burda, Tim-Wolf Gilberger, Stefano Vanni, Tobias Spielmann

**Affiliations:** Bernhard Nocht Institute for Tropical Medicine, Hamburg, Germany; Department of Biology, University of Fribourg, Fribourg, Switzerland; Centre for Structural Systems Biology, Hamburg, Germany; University of Hamburg, Hamburg, Germany

**Author notes:** Institute of Veterinary Pathology, Justus Liebig University Giessen, Giessen, Germany (Paul-Christian Burda).

## Abstract

Malaria parasites endocytose large quantities of hemoglobin from the host erythrocyte, a process critical for parasite survival, leading to extensive membrane internalization. While hemoglobin degradation in the digestive vacuole (DV) is well studied, how the parasite deals with the membranes arriving within the DV is unknown. Here we identified PfTUPA, a previously uncharacterized lipid transfer protein in the DV membrane that is needed for this function. PfTUPA contains a soluble TULIP-like lipid transport domain exposed to the DV lumen and a transmembrane lipid transfer domain of bacterial origin (PqiA) in the DV membrane. Structural comparisons revealed proteins with various PqiA and TULIP-like domain combinations across distant eukaryotic clades, indicating this is a frequent functional partnership. Hence, PfTUPA belongs to a new class of eukaryotic lipid transfer proteins that in malaria parasites is needed for a key function of its biology.

## Introduction

Recent years have revealed the crucial role of non-vesicular lipid transport for cellular homeostasis as a mechanism to ensure proper lipid distribution in organellar membranes^1–3^. Most studies of these mechanisms come from gram negative bacteria, where lipid transport occurs between the inner and outer membranes^4,5^, and from opisthokont model organisms. In the latter, non-vesicular lipid transport mechanisms have been implicated in membrane homeostasis^6,7^, extension^8–10^, repair^11,12^, and recycling^13^, occurring through the function of lipid transfer proteins (LTPs) which can be soluble, in the cytosol or extracellular space, or enriched at sites of interaction between two organellar membranes, so called membrane contact sites (MCSs)^2,3,14–16^. Although studies have covered examples of these mechanisms in other organisms like plants^17,18^, the field remains underexplored, especially in protists.

Given their medical relevance, apicomplexans parasites are amongst the most studied protists of this phylum, with the malaria parasite *Plasmodium falciparum* causing hundreds of thousands of deaths every year^19^. *P. falciparum* parasites develop and differentiate in mosquitos, the human liver and in red blood cells (RBCs), going through a complex cell biological program to propagate in these different environments ^20^. In the RBC stage, the parasite invades and multiplies within a cell otherwise devoid of any internal membranous compartments, orchestrating a multiplication program involving massive membrane generation and remodeling to build up to 32 copies of itself^21,22^. This includes the generation of copies of all cellular organelles, biogenesis of new internal organelles required for invasion, extension of the vacuole hosting the parasite (parasitophorous vacuole) and biogenesis of additional compartments in the RBC cytosol, all within a 48 h replication cycle (Fig. 1A)^23,24^. This poses a strong pressure on the parasite for precise mechanisms of lipid distribution on a considerable scale. A limited number of studies already discovered key roles for certain parasite encoded LTPs in the RBC stage. These include PfSTART1 and PfNCR1, essential proteins with small lipid transfer modules facing the parasitophorous vacuole lumen^25–28^, and PfVPS13L1, a bridge-like LTP recently shown to be involved in inner membrane complex (IMC) extension^29^. However, the involvement of other LTPs in cellular processes remains unknown.

**Figure 1.**
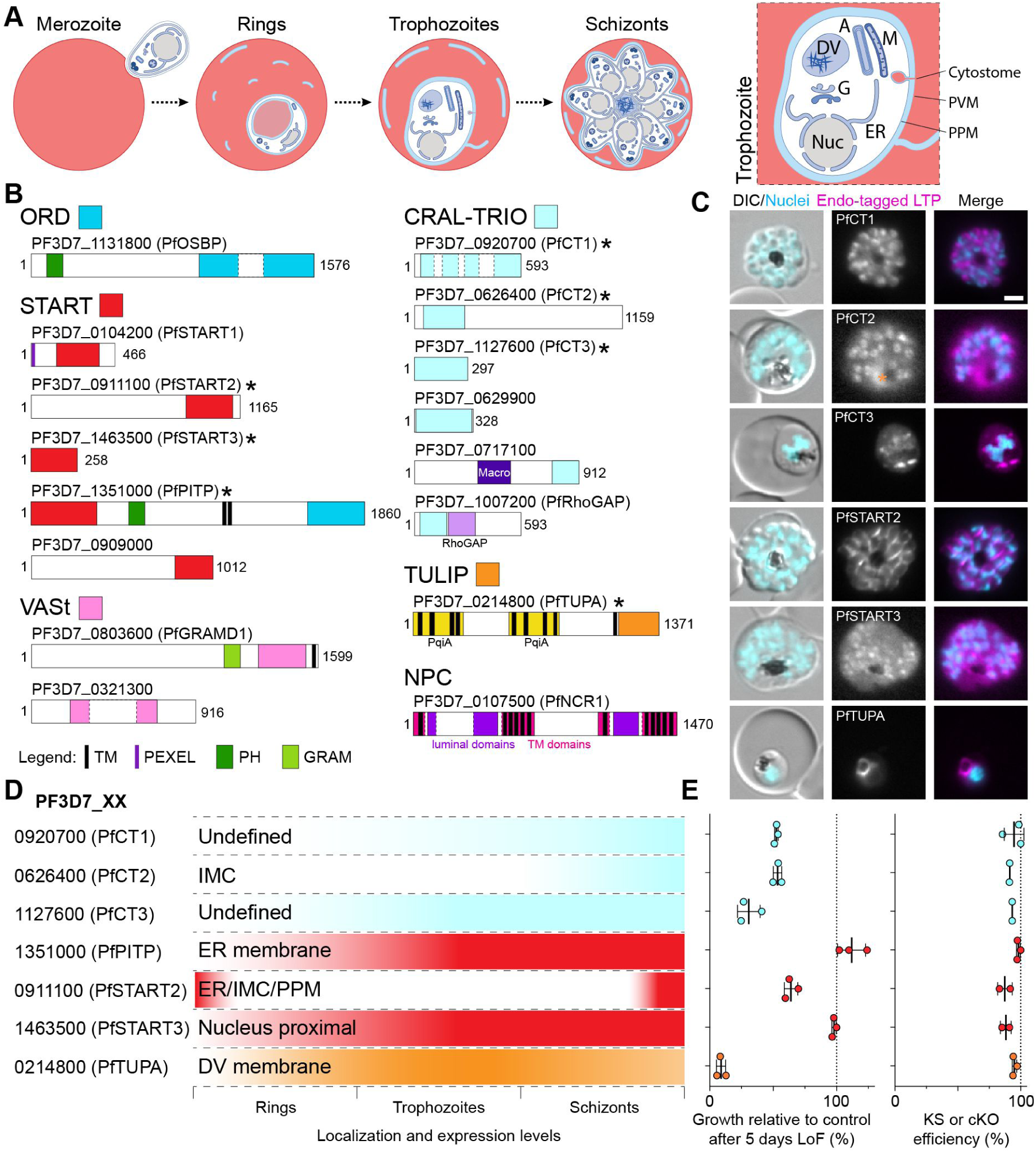
Proteins with small lipid transfer domains in *P. falciparum*. **A**, Schematic of *P. falciparum* growth in the RBC cycle (∼46 h). Right, typical compartments in a trophozoite. Nuc, nucleus; ER, Endoplasmic Reticulum; G, Golgi; M, mitochondria; A, apicoplast; DV, digestive vacuole; PVM, parasitophorous vacuole membrane; PPM, parasite plasma membrane. **B**, Domain cartoon of all proteins found via bioinformatics searches to contain domains characteristic of shuttle-like lipid transfer function in *P. falciparum*. Localization and essentiality for three of these proteins was previously studied^25,28,29^, whereas only localization had been determined for PfPITP. New data for seven proteins (marked with an asterisk) is included in this study. Numbers indicate amino acid number. **C**, Representative fluorescence microscopy images of the indicated endogenously GFP-tagged LTP. Images of different RBC stages in Fig. 3A for PfTUPA and Fig. S2 for the remaining proteins. Orange asterisk indicates DV luminal autofluorescence. **D**, Graphical representation of expression timing and localization of the tagged LTPs during the asexual blood-stage cycle. **E**, Growth relative to control after 5 days of loss of function (LoF) and quantification of the LoF efficiency. LoF was tested through conditional KO (cKO) for proteins with transmembrane regions and through knock-sideways (KS) for the others. Efficiency of cKO was assessed by percentage of cells with expression of the protein of interest after 1 excision cycle and of KS by percentage of cells with full or partial mislocalization (partial mislocalization was given half the weight) after 3h of KS induction, with the exception of PF3D7_0911100 where KS efficiency was assessed after 20h due to slower mislocalization kinetics. Growth graph shows mean with SD from n = 3 independent experiments and inactivation efficiency (KS or cKO) graph shows mean with SD (PF3D7_0920700, n = 3, 154 cells; PF3D7_0626400, n = 2, 80 cells; PF3D7_1127600, n = 2, 60 cells; PF3D7_1351000, n = 3, 124 cells; PF3D7_0911100, n = 2, 60 cells; PF3D7_1463500, n = 2, 89 cells; PF3D7_0214800, n = 2, 174 cells). DIC, differential interference contrast; Nuclei, Hoechst 33342; scale bars, 2 µm.

An essential process for parasite growth in RBCs is the endocytosis of host cell cytosol that consists primarily of hemoglobin, which after internalization is metabolized in the parasite’s digestive vacuole (DV)^30,31^. Up to 80% of the host cell cytosol is consumed in this manner, generating room for parasite growth and producing amino acids and heme^32,33^. The amino acids are to some extent used as building blocks^34–36^ and the heme is detoxified^31,37^. The importance of the DV function is demonstrated by the fact that these processes are the targets of many antimalarial drugs^38–40^ and drug resistance^41,42^. While host cell cytosol endocytosis is not fully understood^30^, double membrane vesicles derived from the cytostome, a membrane invagination of the parasite plasma membrane (PPM) and the parasitophorous vacuolar membrane (PVM) surrounding the parasite, transport hemoglobin to the DV^43,44^, where the outer membrane is thought to fuse with the DV membrane and the inner hemoglobin-containing vesicle is delivered to the DV lumen^45–47^. Only when perturbed, this process becomes evident through a build-up of hemoglobin-filled vesicles in the cytoplasm, or vesicles or undigested hemoglobin in the DV lumen, and illustrates the speed and magnitude of the process^43–45,48^. However, what happens to the membranes arriving in the DV lumen and how these lipids are metabolized or recycled is unknown.

Here, through a systematic analysis of LTPs with small lipid transfer domains in malaria RBC stage parasites, we identified a protein containing a tubular lipid-binding (TULIP)-like lipid transfer domain and an intramembrane domain homologous to the bacterial lipid transporter PqiA, that we named PfTUPA (TUlip-and-PqiA fusion protein). We detected this domain combination in diverse eukaryotic clades with high representation in protists, suggesting PfTUPA is a member of a new class of LTPs we here refer to as TUPAL (TUPA-like) proteins. Functional studies show that PfTUPA is an essential protein of the parasite’s DV membrane where it is needed to dispose of the large amounts of internalized membranes, revealing a so far elusive key function of malaria blood stage parasites. The diversity of domain arrangements in TUPALs suggest this combination has arisen more than once in eukaryotic evolution and indicates that the PqiA lipid transporter is a versatile module co-opted for LTP functions through fusion with TULIP-like domains.

## Results

### Characterization of small LTPs in *P. falciparum*

To complete a previously started survey of LTPs in malaria parasites^29^, we carried out sequence-and structure-based searches with small shuttle-like lipid transfer domains in *P. falciparum* parasites. This led to the identification of proteins containing the oxysterol-binding protein-related domain (ORD), the Steroidogenic Acute Regulatory lipid transfer (START) domain and the related VAD1 Analog of START (VASt), the CRAL-TRIO and the TULIP domain (Fig 1B). All of these proteins had also previously been identified^29,49–52^ with the exception of the TULIP-domain containing protein PF3D7_0214800 which was identified here using Foldseek structural similarity searches^53^. However, apart from PfSTART1 and PfNCR1^25–28^, little is known about these proteins. In a previous study of endoplasmic reticulum (ER)-related LTPs we reported the localization of PfPITP, PfGRAMD1 and PfOSBP to the ER and showed that PfOSBP was at ER-contact sites and was dispensable for RBC stages^29^.

In order to complete our survey, we attempted to study the localization and function of the remaining proteins by generating parasite lines where the genes were endogenously modified using selection-linked integration (Fig. S1A)^54^. Parasite lines for six not previously studied proteins were obtained (Fig. S1B) and the localization and expression levels during asexual RBC stages were determined (Fig. 1C-D and Fig. S2): PF3D7_0920700 (PfCT1) localized to nucleus-proximal hotspots in trophozoites and in segmenter (late schizont) stage parasites was enriched at a filamentous structure. PF3D7_0626400 (PfCT2) was expressed in schizont stages with a localization pattern indicative of IMC. PF3D7_1127600 (PfCT3) showed a punctate localization in proximity to the nucleus. PF3D7_0911100 (PfSTART2) localized to the merozoite periphery in late schizonts, which could be the IMC or PPM. PF3D7_1463500 (PfSTART3) showed a punctate pattern surrounding the nucleus. PF3D7_0214800 (PfTUPA) was at the DV membrane (see figures below for full cycle localization and expression of PfTUPA). The localization of PfPITP was known to be at the ER^29^, however, upon colocalization with an ER marker we noticed that this protein was enriched at hotspots along the ER, as expected if at a contact site with another organelle (Fig S2 and S3A-B). Besides the lipid transfer modules, PfPITP has two transmembrane (TM) domains expected to anchor the protein at the ER and a PH domain which could mediate interaction with another membrane (Fig. 1B). Accordingly, a tandem of this PH domain had a punctate pattern (Fig. S3C), suggestive of binding to a compartment, and confirming that our list of LTPs with small lipid transfer domains includes proteins behaving as expected for MCS proteins.

Next, we determined the essentiality of each of the selected LTPs for RBC stages using conditional gene knock-out (cKO; for membrane proteins) or knock-sideways (KS; for soluble proteins). PfSTART3 and PfPITP were dispensable, whereas the KS of PfCT1, PfCT2, PfCT3 and PfSTART2 caused a partial growth defect of 35 to 70%. The loss of PfTUPA by cKO resulted in the most severe growth defect (Fig. 1E). We conclude that malaria blood stages contain a series of proteins with small LTP domains at various cellular sites with important functions for efficient blood stage growth. The location of PfTUPA at the DV membrane, suggesting a potential role in the endosomal system, and the severity of its loss of function, prompted us to further study this protein.

### Structural analysis of PfTUPA reveals a new class of LTPs present in diverse eukaryotic clades

Structurally, PfTUPA contains two domains, an N-terminal region composed of 8 TM helices (split in two units of 4 TM helices each), and the TULIP-like lipid transfer domain in its C-terminal region (Fig. 2A-B, and Fig. S4A). Using Foldseek with the TULIP-like domain to search the RCSB-PDB database identified the closest similarity with the dust mite allergen DerP7 and the human bactericidal permeability increase (BPI), both belonging to the TULIP superfamily^55,56^. Although structural alignments using US-align^57^ suggested only a remote similarity (Fig. 2C and Fig. S4B), the hydrophobic cavity essential for the lipid transfer function was conserved in the domain of PfTUPA (Fig. 2D). The two blocks of 4 TM helices in PfTUPA showed structural similarity with the bacterial protein PqiA. The N-terminal region (PqiA-N) showed a highly similar fold to half of the PqiA protein of *E. coli* (Fig. 2E), whereas the more C-terminal four-TM region (PqiA-C) showed lower alignment scores to bacterial PqiA (Fig. 2F). This likely is due to the low confidence of *P. falciparum* structures, since PfTUPA homologs in other apicomplexans did show high similarities of their PqiA-C domains with bacterial PqiA (Fig. S4C-D).

**Figure 2.**
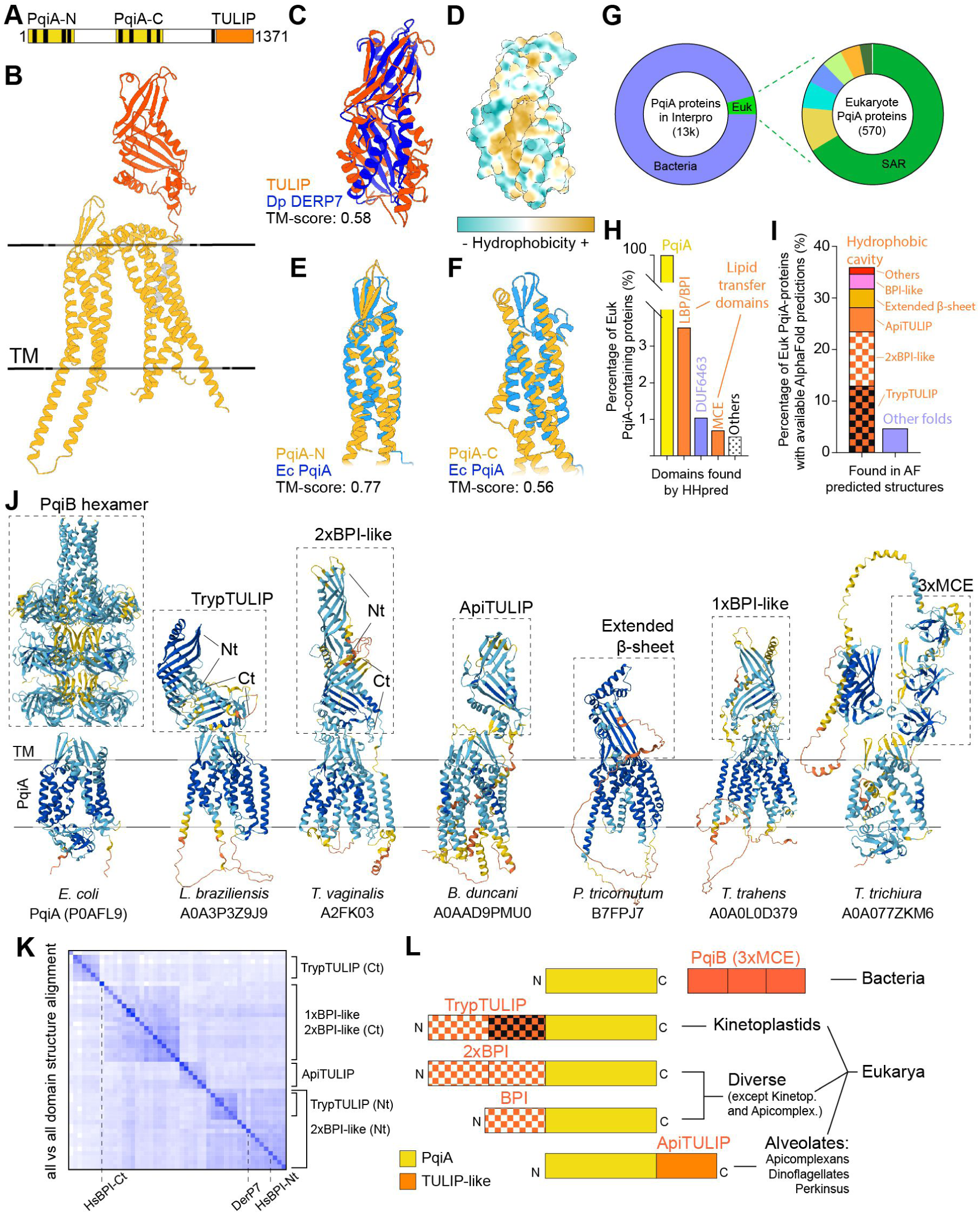
The intramembrane lipid transporter PqiA associates with exposed lipid transfer domains across eukaryotes. **A**, Domain cartoon of PfTUPA. Black columns represent TM regions; other colors as in B. **B**, Ribbon representation of AlphaFold3-predicted structure of PfTUPA colored by domain and with the disordered regions removed. Complete structure colored by confidence in Fig. S4A. Black lines represent the edges of the membrane bilayer. **C**, Structural alignment between the predicted structure of the ApiTULIP domain of PfTUPA and the DerP7 allergen (PDB 3H4Z). **D**, Surface representation of the predicted ApiTULIP domain structure colored by hydrophobicity. **E-F**, Structural alignments between the predicted structures of the two halves of the PqiA domain (PqiA-N, **E**; PqiA-C, **F**) formed by four TM regions each and the C-terminal portion of the *E. coli* PqiA protein. **G**, PqiA-containing proteins found in the Interpro database. In eukaryotes proteins were found in different clades (ordered by percentage): SAR, Discoba, Haptista, Amoebozoa, Viridiplantae, Opisthokonta, Metamonada, Apusozoa. **H-I**, Percentages of eukaryotic PqiA-containing proteins containing soluble lipid transfer domains identified by HHpred (**H**) or domains with hydrophobic cavities, likely involved in lipid transfer, observed in the AlphaFold predicted structures (**I**). **J**, AlphaFold predicted structures of PqiA-containing proteins. In bacteria, PqiA (inserted in the membrane) interacts with the periplasmic MCE-containing lipid transfer protein PqiB. In eukaryotes, the PqiA domains are largely found fused to a soluble lipid transfer domain, usually of the TULIP superfamily. Colors indicate confidence of AlphaFold prediction. **K**, All-vs-all structural alignment of a sample of the TULIP BPI-like domains found in eukaryotic PqiA-containing proteins. TrypTULIP and 2xBPI domains were separated in their N-terminal (Nt) and C-terminal (Ct) halves as indicated in (**J**). Structures of domains of the human BPI protein and the DerP7 allergen are included. **L**, Domain cartoons (not to scale) summarizing the association of PqiA proteins with soluble lipid transfer domains in bacteria and eukaryotes.

Interestingly, bacterial PqiA is part of the Paraquat inducible operon, comprising: PqiA, a TM protein in the bacterial inner membrane, PqiB, a protein composed of 3 bacterial Mammalian Cell Entry (MCE) lipid transfer domains in the periplasmic space, and PqiC, an adaptor to the outer membrane^4^. Together these proteins form one of the several tunnel-like lipid transfer systems between the inner and outer membrane of gram-negative bacteria. Recently, a close bacterial homolog of PqiA, LetA, was determined to work as a lipid transporter^58^, mobilizing lipids between the membrane and the exposed lipid transfer domain, which in the case of bacteria is PqiB or LetB^59^, and in the case of *P. falciparum* would be the TULIP-like domain.

The identification of an eukaryotic PqiA-containing protein fused to a lipid transfer domain prompted us to investigate the conservation of this domain combination in eukaryotes, which was also reported to exist in other protists^56,58^. The majority of PqiA-containing proteins in the Interpro database are found in bacterial genomes, but a small percentage is found in eukaryotes (excluding other distantly related eukaryotic proteins, such as the tetraspanin family of proteins^58^) (Fig. 2G). HHpred analysis of these proteins revealed that the BPI-like domain belonging to the TULIP superfamily of proteins was the most prevalent additional domain in these proteins (Fig. 2H). Since the TULIP-like domain of PfTUPA could only be identified via structural searches, we examined all the available predicted AlphaFold^60^ structures of eukaryotic PqiA-containing proteins in the Interpro database. We found that close to 35% of these proteins have the PqiA-domains fused to a domain with a hydrophobic cavity (Fig. 2I-J), always in the same topologic orientation. Interestingly, these hydrophobic cavities differed across eukaryotes. Most of these domains had similarities to the BPI domains of the TULIP superfamily, with some proteins having two copies of these N-terminally fused to the PqiA-domain, or one copy fused either N-or C-terminally. However, some proteins found in Stramenopiles had an extension of the β-sheet of the PqiA-domain which formed a hydrophobic surface (Fig. 2I-J), similar to what has been observed for the bacterial lipid transport complex formed by TamA and TamB^61^.

To classify the BPI-like domains fused to PqiA-type proteins, we performed an *all-vs-all* structural alignment (Fig. 2K) of a sample of 48 of them using the DALI server^62^. In proteins with two BPI-like domains (found across distinct eukaryotic evolutionary lineages), the N-terminal half had strong similarities and also clustered with the N-terminal domain of the human BPI protein (Fig. 2K-L and Fig. S4E-G). However, differences were found between the C-terminal parts of the BPI-like doublet. In kinetoplastids, the C-terminal half of the domain does not have similarities with other eukaryotic domains, suggesting this extension is a unique development of this phylum, hence we refer to this tandem domain as TrypTULIP (Fig. 2K-L and Fig. S4E&G). In the non-kinetoplastid proteins with two BPI-like domains the C-terminal half also had similarities with other eukaryotic BPI-like domains, and aligned well with its N-terminal counterpart (example from Viridiplantae in Fig. S4F-G). These domains were also similar to the ones found in proteins with a single BPI-like domain in their N-terminus (Fig. 2K). Interestingly, the single BPI-like domains found in proteins of alveolates, such as the one in PfTUPA, did not cluster with the other domains and were the only domains found on the C-terminal end of PqiA (Fig. 2K-L). We refer to this BPI-like domain as ApiTULIP, as it was exclusively found in apicomplexans, whereas some other alveolates also have proteins containing 2xBPI-like domains in their N-terminus (such as the dinoflagellate species *P. glaciallis*, Fig. S4I). Accordingly, using sequence-based cluster analysis (CLANS) the ApiTULIP and TrypTULIP domains were shown to cluster independently of each other and of other known domains of the TULIP superfamily extracted from the Pfam database (Fig. S4H). Given the identified widespread presence of proteins with lipid transport modules in similar arrangements to PfTUPA across several eukaryotic clades, we refer to this class of proteins of the TULIP superfamily as TUPA-Like proteins (TUPALs).

Taken together, these results argue for a strong functional partnership between PqiA-domains and lipid transfer domains across evolution, ranging from the bacterial PqiA protein interaction with the MCE-containing protein PqiB, to TUPALs, a class of proteins across diverse eukaryotic lineages with differently arranged combinations of PqiA and BPI-like domains (Fig. S4I-J). The diversity in structures of BPI-like domains in TUPALs suggests that this combination has occurred multiple times in evolution which is also supported by the finding that the BPI-like domain can be found fused to the N-or C-terminus of PqiA (Fig. 2L). To gain insight into this type of protein and its function in malaria parasites, we carried out a detailed analysis of PfTUPA.

### PfTUPA is at the DV membrane with its ApiTULIP domain facing the lumen

For better visualization of PfTUPA in the parasite, we generated parasite lines where it was endogenously tagged with Halo on its N-terminus (Halo-PfTUPA^endo^) (Fig. S1B). As observed with the GFP-tagged version, the protein was expressed in all asexual blood stages and localized to the DV membrane, with some additional lower signal at the ER, the PPM and smaller cytosolic compartments (Fig. 3A). These additional signals are likely from the biosynthetic and trafficking route of the protein, as pulse-chase experiments using different Halo-dyes determined that the ER and PPM signal is more prominent with protein translated in the last 3 hours, whereas most of the older pool of protein was found at the DV membrane (Fig. S5). ER-PPM-DV trafficking has also been proposed for the DV luminal protease Plasmepsin II (PMII), indicating this is a shared route by some DV proteins^63^.

**Figure 3.**
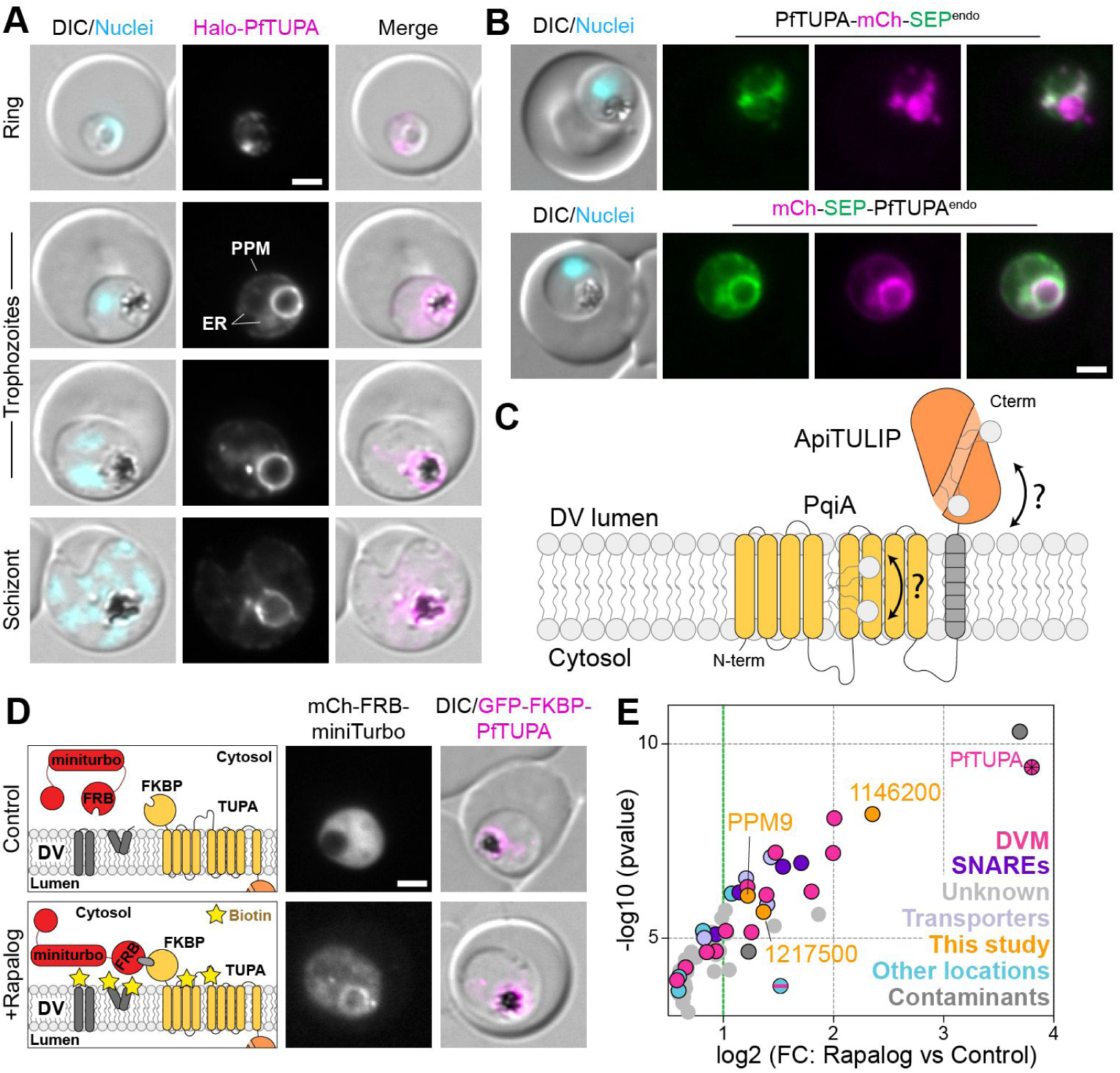
PfTUPA exposes its ApiTULIP lipid transfer domain to the DV lumen. **A**, Fluorescence microscopy images of endogenously Halo-tagged PfTUPA in asexual blood stages. **B**, Fluorescence microscopy images of endogenously tagged PfTUPA with mCh-SEP either C-or N-terminally (top and bottom, respectively). **C**, Schematic of the proposed molecular function of PfTUPA. Lipids are likely transferred between the DV lumen via the exposed ApiTULIP domain (orange) and the DV membrane via the TM PqiA domain (yellow). **D**, Schematic (left) and fluorescence microscopy images (right) showing rapalog-mediated recruitment of the episomally expressed miniTurbo biotinilyzer to the cytosolic side of PfTUPA. **E**, Plot of the PfTUPA dimerization-induced BioID (DiQ-BioID)^42^ outlined in (**D**) showing enrichment of proteins in plus rapalog over control. Hits are considered proteins with an absolute fold change (FC) of 2 or more and a false discovery rate (FDR) of 0.05 or less (27 proteins, to the right of the green dotted line); the remaining proteins in this plot are considered enriched, absolute FC>1.5, FDR <0.2 (35 proteins). Averages from *n* = 3 independent replicates analysed side by side in the same run (data in Table S1). Moderated two-tailed t-test was applied as implemented in the limma package. Hits are further annotated in Figure S6A-B and Table S1. Three of the proteins with previously unknown localization identified as hits were Halo-tagged for this study and are indicated in orange. DIC, differential interference contrast; Nuclei, Hoechst 33342; scale bars, 2 µm.

To determine the topology of PfTUPA in the DV membrane, we generated lines where we endogenously tagged the protein with a tandem of mCherry and a pH-sensitive phluorin-like protein (superecliptic phluorin, SEP^64^) in the N-or C-terminus of PfTUPA (Fig. S1B). When these proteins reach the DV membrane, the mCherry protein should fluoresce independently of the topology, whereas SEP fluorescence – due to its sensitivity to low pH - should only be observed if the protein was exposed to the cytosol, which occurred only when SEP was at the N-terminus (Fig. 3B). This topology indicates that PfTUPA might transport lipids between the DV intraluminal ApiTULIP lipid transfer domain and the DV membrane embedded PqiA region (Fig. 3C).

Confirming the results from the topology experiments, episomal expression of a cytosolic miniTurbo biotinylizer construct (miniTurbo fused to mCherry and an FRB domain) in the GFP-2xFKBP-PfTUPA^endo^ line (Fig. S1B) allowed for rapalog-inducible recruitment of this construct to, and hence biotinylation at, the cytosolic side of the DV membrane (Fig. 3D). Biotinylated proteins in the rapalog condition enriched over control identified many proteins known to act at the DV membrane, such as the transporters MDR1, MDR2, VIT, DMT, AAT1 and CRT, confirming the accuracy of the labelling assay (Fig. 3E, Fig. S6A-B and Table S1). Most of the remaining hits had an unknown localization but their inferred function based on homology was in many cases compatible with a location at the DV membrane (such as SNAREs or transporters). We studied some of the highest-ranked hits with unknown localization and function to assess a potential functional interaction with PfTUPA. While the top hit (PF3D7_1146200) localized to the DV membrane (Fig. S6C), cKO showed it is dispensable (Fig. S6D), and two further tested hits were only partially (PF3D7_1217500) or not (PF3D7_0520100) at the DV membrane (Fig. S6E-F). Hence, while this experiment re-enforced the topology of PfTUPA in the DV membrane and revealed a plausible DV surface proteome useful as a resource, it did not provide evidence for functionally relevant interaction partners at its cytosolic face, prompting a deeper direct functional analysis focusing on PfTUPA.

### PfTUPA is needed for recycling of membranes from the DV

To understand the function of PfTUPA in the biology of the parasite we better characterized the growth phenotype of the *Pftupa* cKO. We induced the excision of *Pftupa* in ring stage parasites and monitored the growth of the parasites from the following cycle, when the protein had been lost (Fig. 4A). The cKO parasites were delayed in development when compared with controls, and there was a 4-fold decrease in the number of progeny rings in the next cycle (Fig. 4A). Upon closer examination, we noticed a severely enlarged DV in the cKO parasites, which seemed to worsen with further progress of parasite development (Fig. 4B). A percentage of the parasites, although much delayed (Fig. 4C), was able to go through rounds of nuclear division and produce merozoites despite the substantially enlarged DV (Fig. 4A). However, much of this progeny presented with the same phenotype in the following cycle, which now halted its development, leading to a 60-fold difference in parasitemia compared with the control after 2 cycles without PfTUPA (Fig. 4A).

**Figure 4.**
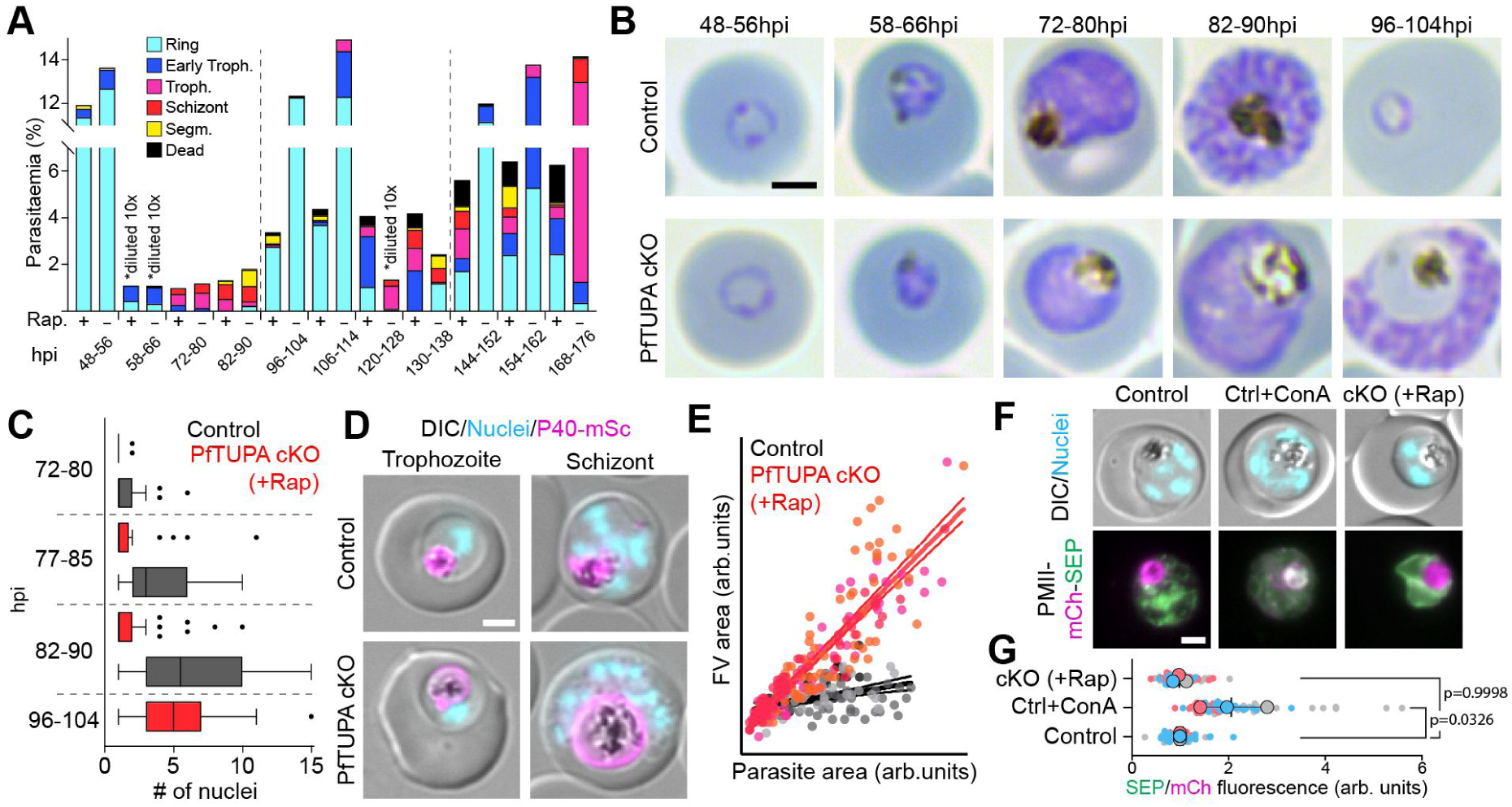
PfTUPA is essential for DV function independent of acidification. **A**, Parasite growth and stage progression assay, along two and a half cycles, in synchronous parasites (8 h stage window) of control and Pf*tupa* cKO (GFP-2xFKBP-PfTUPA^endo^ parasites with or without one cycle of rapalog-activated diCre excision), based on Giemsa smears. Samples were taken after one cycle (48hpi) of synchronization and induction. Parasites were diluted 10x when parasitemia surpassed 10% to prevent overgrowth. **B**, Representative Giemsa smears of control and Pf*tupa* cKO parasites along the first cycle after PfTUPA excision. **C**, Graph of number of nuclei in parasites at the indicated hpi. Parasites along the first cycle after excision from n=2 independent experiments; a total of 82, for control, and 144, for cKO, individual parasites were quantified. **D-E**, Representative fluorescence microscopy images (**D**) and graph of DV and parasite area measurements (**E**) of control and Pf*tupa* cKO trophozoites and schizonts expressing the PI3P-binding protein and DV membrane marker P40-mSc, used to measure DV area. Parasites along the first cycle after PfTUPA excision from n=3 independent experiments; a total of 109, for control, and 187, for cKO, individual parasites were quantified in (**E**). Different shades of color represent different experiments, lines represent simple linear regressions with 95% confidence intervals, slopes are significantly different with a p-value<0.0001 using ANCOVA. **F-G**, Representative fluorescence microscopy images (**F**) and graph of SEP/mCh fluorescence intensity ratio in the DV lumen (**G**) of control (with or without ConA-mediated inhibition of the DV V-ATPase) and Pf*tupa* cKO parasites expressing a tandem construct of mCh and the pH-sensitive SEP protein fused to PMII for targeting to the DV lumen. Values are normalized to the control and presented as a superplot^67^, from n=3 independent experiments with a total of 60, 59 and 69 cells quantified for the control, conA-treated and Pf*tupa* cKO parasites, respectively; colors indicate independent experiments (small dots, individual parasites; large dots, average of each experiment; black lines, mean and SD; p-value, Dunnett’s multiple comparisons test of the means). DIC, differential interference contrast; Nuclei, Hoechst 33342; scale bars, 2 µm.

As the DV size increases with development, and the cKO presented a significant growth delay, we compared DV area to parasite area (Fig. 4D-E). This showed that the phenotype was specific and not due to a stage delay (Fig. 4E) and in schizonts, the DV area of the cKO was ∼3x that of control cells (Fig. 4D-E). Previously published swelled DV phenotypes can be due to inhibition of luminal cysteine proteases that degrade the endocytosed hemoglobin^48^ or impaired DV acidification^45,47^. However, although seemingly similar upon Giemsa staining or DIC microscopy, these swelling phenotypes are not identical: disrupted acidification leads to accumulations of intact vesicles in the DV lumen, whereas protease inhibition leads to the accumulation of undigested hemoglobin evenly distributed in the DV. We thus explored the Pf*tupa* cKO phenotype further to understand which materials were accumulating in the DV lumen.

First, we tested whether a failure in DV lumen acidification could be behind the observed swelling. For this, we episomally expressed a PMII construct fused to a tandem of mCherry and SEP proteins (PMII-mCh-SEP) in the Halo-PfTUPA^endo^, and induced *Pftupa* gene excision. An increase in fluorescence of SEP in relation to mCherry would indicate a defect in acidification, but this was not observed for the Pf*tupa* cKO (Fig. 4F-G) while treatment with concanamycin A, an inhibitor of the V-ATPase previously used in *P. falciparum* parasites ^45,65^, increased fluorescence (Fig. 4G), validating the probe used for this assay. These findings differentiated the Pf*tupa* cKO phenotype from the one observed upon loss of the V-ATPase.

Next, we assessed the nature of the DV phenotype by ultrastructure expansion microscopy (U-ExM) in different stages after PfTUPA loss. Staining with NHS-ester in U-ExM allows for general protein or membrane detection, depending on the fixative and the presence of BSA during sample staining^66^. We took advantage of this to determine the composition of the accumulated content in the swollen Pf*tupa* cKO DVs. In cKO single nucleus trophozoites, the DV lumen was full of membranous accumulations, while amounts of protein in the DV varied between different cells (Fig. 5A-D and S7A). Quantification of this phenotype by scoring the imaged cells based on percentage of DV area with accumulated protein, indicated that while the DVs were all filled with membrane material, there was only partial accumulation of protein (Fig. 5D). This clearly separated the phenotype also from that of DV protease inhibition. In cKO schizonts similar membranous accumulations were observed, although with an increase in the fraction of cells with protein-filled DVs (Fig. 5A-D and S7B) that showed some similarity to the accumulations observed upon loss of the V-ATPase function^45^. Together, these findings indicate that the membranes arriving in the DV are not properly cleared if PfTUPA is absent and with increased progression through the cycle this also leads to a defect in release or digestion of hemoglobin.

**Figure 5.**
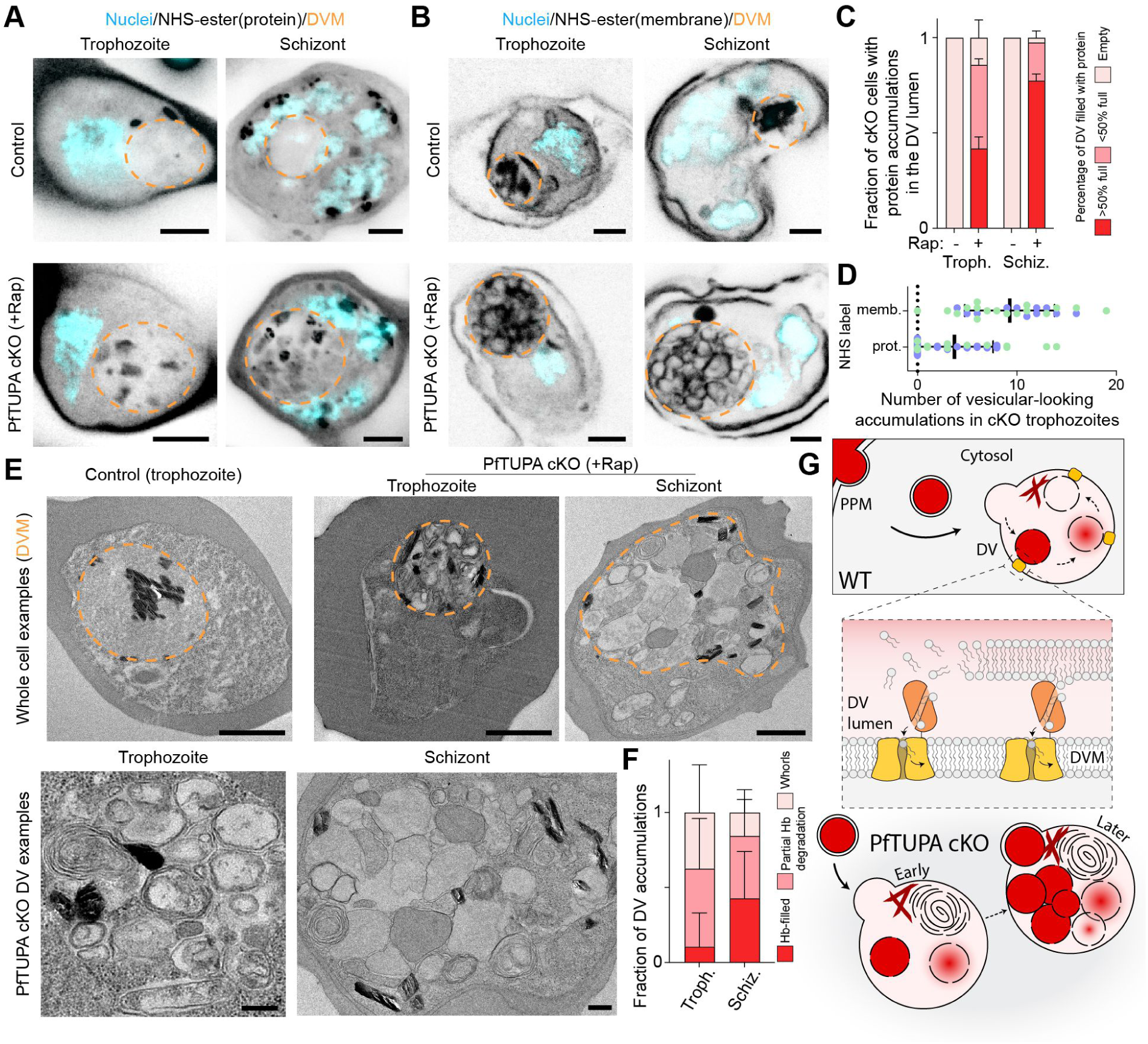
Depletion of PfTUPA leads to an accumulation of membranes in the DV lumen. **A-B**, Ultrastructure Expansion Microscopy (U-ExM) images of control and Pf*tupa* cKO parasites in trophozoite or schizont stages stained with NHS-ester in the absence (**A**) or presence (**B**) of BSA, used to stain proteins or membranes, respectively. The inferred DV periphery is represented with an orange dashed line. **C**, Graph of percentage of cells showing protein accumulations in the DV lumen in the stage and conditions indicated in U-ExM experiments (n=2 experiments for each; 24&13 trophozoites and 20&10 schizonts counted). **D**, Graph of number of vesicular looking accumulations in Pf*tupa* cKO trophozoites stained for protein or membranes (n=2 experiments for each, 24&13 and 11&17 trophozoites counted for protein and membrane staining, respectively). **E**, Transmission electron microscopy (TEM) of Pf*tupa* cKO parasites in trophozoite and schizont stages. Examples of whole-cell images and insets of DV content are presented. The inferred DV periphery is represented with an orange dashed line in the whole-cell examples. **F**, Graph of fraction of different types of accumulations in the DV lumen of Pf*tupa* cKO parasites as assessed by TEM (n=2 for trophozoites, 48 and 10 cells counted; n=1 for schizonts, 22 cells counted). Accumulations were classified according to their apparent content, examples of classifications in Fig. S7C. **G**, Schematic of proposed function of PfTUPA. In WT cells (upper two panels), PfTUPA transports glycerolipids or their degradation products from the DV lumen to the membrane, via its ApiTULIP and PqiA domains. Loss of this protein leads to accumulation of membranous material in the lumen (bottom panel). In the cell schematics, dark red vesicular content represents hemoglobin; dark red crystals, hemozoin; black lines, membranes; black arrows, vesicular trafficking; dashed-black arrows, time progression. Nuclei, Hoechst 33342; scale bars, 5 µm in **A,** 1 µm in **E** (whole cell) and 0.25 µm in **E** (insets).

To confirm these observations, we carried out electron microscopy (EM) with both stages of Pf*tupa* cKO and control parasites. Similarly to what was observed in U-ExM, trophozoites showed predominantly membranous accumulations (i.e. membrane whorls or empty vesicular structures) (Fig. 5E(top)-F and S7C) and protein accumulations became obvious only in late stages of the phenotype (Fig. 5E(bottom)-F and S7C). This suggests that the swollen DVs produced by the Pf*tupa* cKO differs from previously reported DV defects. Considering the topology and inferred lipid transporter capability of PfTUPA, it is likely that it is involved in recycling of membranes that reach the DV lumen after endocytosis (Fig. 5G, early). As the parasite grows and endocytosis continues, these membrane accumulations likely lead to further defects in DV function, causing luminal accumulation of intact hemoglobin-filled vesicles (Fig. 5G, later).

### ApiTULIP lipid transfer domain is important for the function of PfTUPA

Having established the essentiality of PfTUPA for proper *P. falciparum* growth and clearance of membranes from the DV, we aimed to elucidate the underlying molecular mechanism of PfTUPA function. As expression of *P. falciparum* proteins can be challenging and we had not so far succeeded in producing recombinant PfTUPA, we resorted to *in silico* approaches. We first investigated the lipid-binding ability of the ApiTULIP domain by performing coarse-grained molecular dynamics (CG-MD) simulations following an established protocol (Fig. S8A)^68^, using the Alphafold3 prediction of the ApiTULIP domain (residues 1145-1371, pLDDT=85.85) as starting structure (Fig S8B). In these simulations, POPC spontaneously and repeatedly bound to the ApiTULIP domain (Fig. 6A-B). Analysis of the entry pathway for various replicates showed that the lipid enters the cavity directly through the open conformation of the domain, where L1158, I1170 and V1320 are located (Fig. 6B). Further, we identified residues L1232, L1234, V1273, V1275, I1291, V1328, and L1352 as those that interacted the most (> 75% of simulation time) with POPC during the simulations (Fig. 6C). Based on our analyses and previous studies^68,69^, we next attempted to design a mutant with impaired lipid-binding ability: to do so, we mutated two small hydrophobic residues (one close to the entry of the cavity and one inside the cavity) into bulky tryptophane residues (L1232W/V1320W). Our simulations confirmed that the mutations succeeded at blocking the entry of POPC into the ApiTULIP domain (Fig. 6D, Fig. S8C). Finally, we investigated whether the cavity can accommodate multiple lipids. To this end, we iteratively added POPC molecules to ApiTULIP^68^, and we observed that the domain was able to bind multiple lipids (Fig. S8D-F).

**Figure 6.**
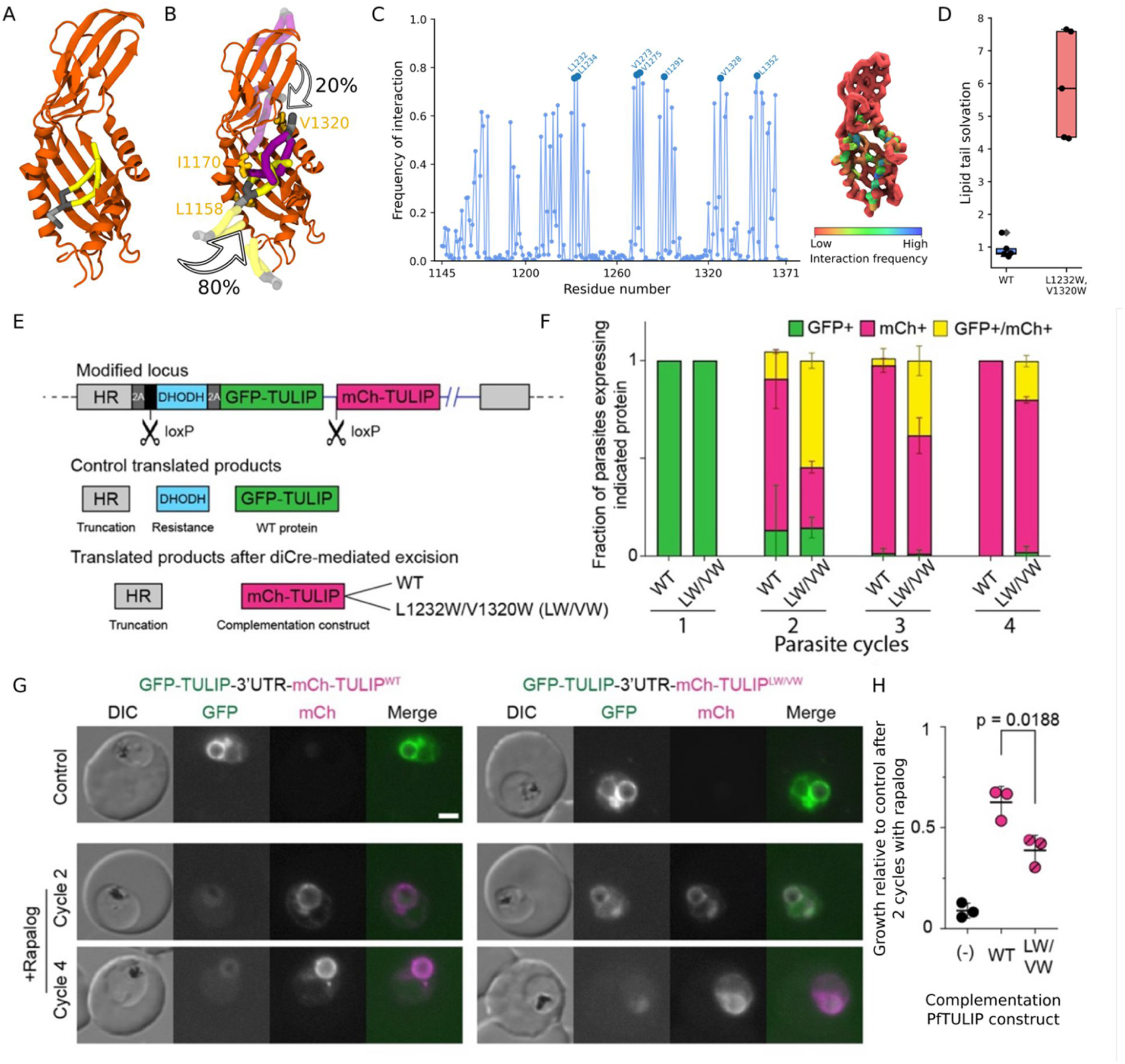
Lipid transfer capability of the exposed ApiTULIP domain is confirmed by CG-MD simulations and is important for PfTUPA function. **A**, Representative final state from CG-MD simulations showing one POPC molecule (tails in yellow; headgroup in grey) bound inside the hydrophobic cavity of the ApiTULIP domain (orange). **B**, Entry pathways of POPC into the ApiTULIP cavity. The major pathway, 80%, is represented by the POPC molecule in yellow while the secondary route, 20%, is represented by the POPC molecule in purple. The residues at the entry of both pathways are shown in orange licorice. **C**, Average frequency of interaction with POPC of each residue from all replicas. On the right, the ApiTULIP domain colored according to the interaction frequency. **D**, Lipid tail solvation during the last 100 ns of CG-MD trajectories indicating lipid binding within the cavity of wildtype (WT) ApiTULIP and lipid remaining in bulk water in presence of the L1232W/V1320W double mutant. **E**, Schematic of the genomic modification at the PfTUPA endogenous locus to allow for the diCre-mediated switching between expression of GFP-and mCh-tagged PfTUPA, the latter in its WT or mutated form to test for complementation of the WT function. GFP-and mCh-tagged PfTUPA are separated by a 3’UTR sequence. **F-G**, Quantification of the efficiency of switching (**F**) and representative fluorescence microscopy images (**G**) of the GFP-2xFKBP-PfTUPA-3’UTR-mCh-PfTUPA (WT or LW/VW)^endo^ parasites (trophozoites) before and after rapalog-activated diCre excision of the GFP-tagged protein. Graph shows mean with SD from n=3 independent experiments (cycle 2: 5, 15, 27 and 7, 32, 20; cycle 3: 23, 21, 14 and 30, 29, 20; cycle 4: 18, 12, 30 and 31, 36, 40 parasites assessed for “WT” and “LW/VW”, respectively, in each experiment). **H**, Growth relative to control 2 cycles after induction of diCre excision and, hence, expression of the mCh-tagged complementation construct (WT or mutated). Growth graph shows mean with SD from n=3 independent experiments. Data of “None” complementation is the same as the cKO presented in Fig. 1E. DIC, differential interference contrast; scale bars, 2 µm.

Based on the CG-MD analysis, we next used the L1232W/V1320W double mutant to test *in vivo* the essentiality of the lipid transfer domain. Unfortunately, episomal expression of PfTUPA failed, as the transfected parasites rapidly lost expression in repeated transfection attempts. To circumvent this, we used a complementation approach based on genetic editing. We modified the endogenous locus of *Pftupa* to encode two copies of the protein separated by a 3’UTR, and with the first copy flanked by loxP sites (Fig. 6E). In this way, in control conditions only the first copy of PfTUPA, GFP-tagged and always WT, is expressed. However, upon diCre-mediated excision, this copy is “switched” for a second copy, mCherry-tagged, to be expressed under the same native promoter (Fig. 6E). We generated two parasite lines, one where the second copy was a WT sequence or one where the second copy had the L1232W/V1320W mutations tested for abolished lipid binding to the ApiTULIP cavity (Fig. S1B). After a cycle of diCre-mediated excision, the mCherry-tagged copy was expressed as expected (Fig. 6F-G). Although the PfTUPA WT constructs did not fully complement the function of the original GFP-tagged WT copy, there was a significantly larger growth defect in parasites where the protein was replaced by the mutant copy (Fig. 6H), highlighting an important role for the ApiTULIP domain and supporting a function of this protein in lipid transfer. Interestingly, whereas the switching was highly efficient in the case of the line with the second WT copy, there was a delay in loss of cells additionally still expressing the endogenous copy when the second copy was the mutated version. It is possible that the impaired complementing activity of the mutated copy contributed to the slower establishment of parasites lacking expression of the original GFP-tagged WT copy.

### MD simulations provide a molecular basis for PfTUPA function

Results from the *in silico* lipid binding assay of the lipid-binding incompetent ApiTULIP double mutant (L1232W/V1320W) highlighted two regions on the periphery of the domain (Fig S8C) where the lipid preferentially localized, suggesting that these regions may serve as membrane association sites. Interestingly, prediction of the domain’s orientation relative to the membrane using OPM^70^ identified one of these regions, the furthermost end opposite the TM domains of the protein, as the membrane-associated surface (Fig. S9A). Following these observations, we performed CG-MD simulations to identify the possible membrane-associated regions of the ApiTULIP domain. Two different sets of simulations were performed: (i) one where the domain was associated with the membrane’s surface as predicted by OPM, and (ii) one where the domain was placed away from the membrane and thus allowed to diffuse and spontaneously bind to the membrane in an unbiased manner (Fig. S9B, insets of initial states). In both cases, the ApiTULIP domain showed preferential membrane association through residues T1201, F1202, L1249-T1260 and S1307-V1313, corresponding to the distal end opposite to the TM domains of PfTUPA (Fig. 7A, Fig. S9B). To determine whether the presence of lipids within the cavity influences membrane binding by the ApiTULIP domain, we repeated the CG-MD simulations described above with six POPC molecules initially placed inside the cavity. Results from the lipid-filled domain are comparable to the empty one, further confirming the membrane binding region (Fig. S9B). Notably, no lipid exchange events were observed in these simulations.

**Figure 7.**
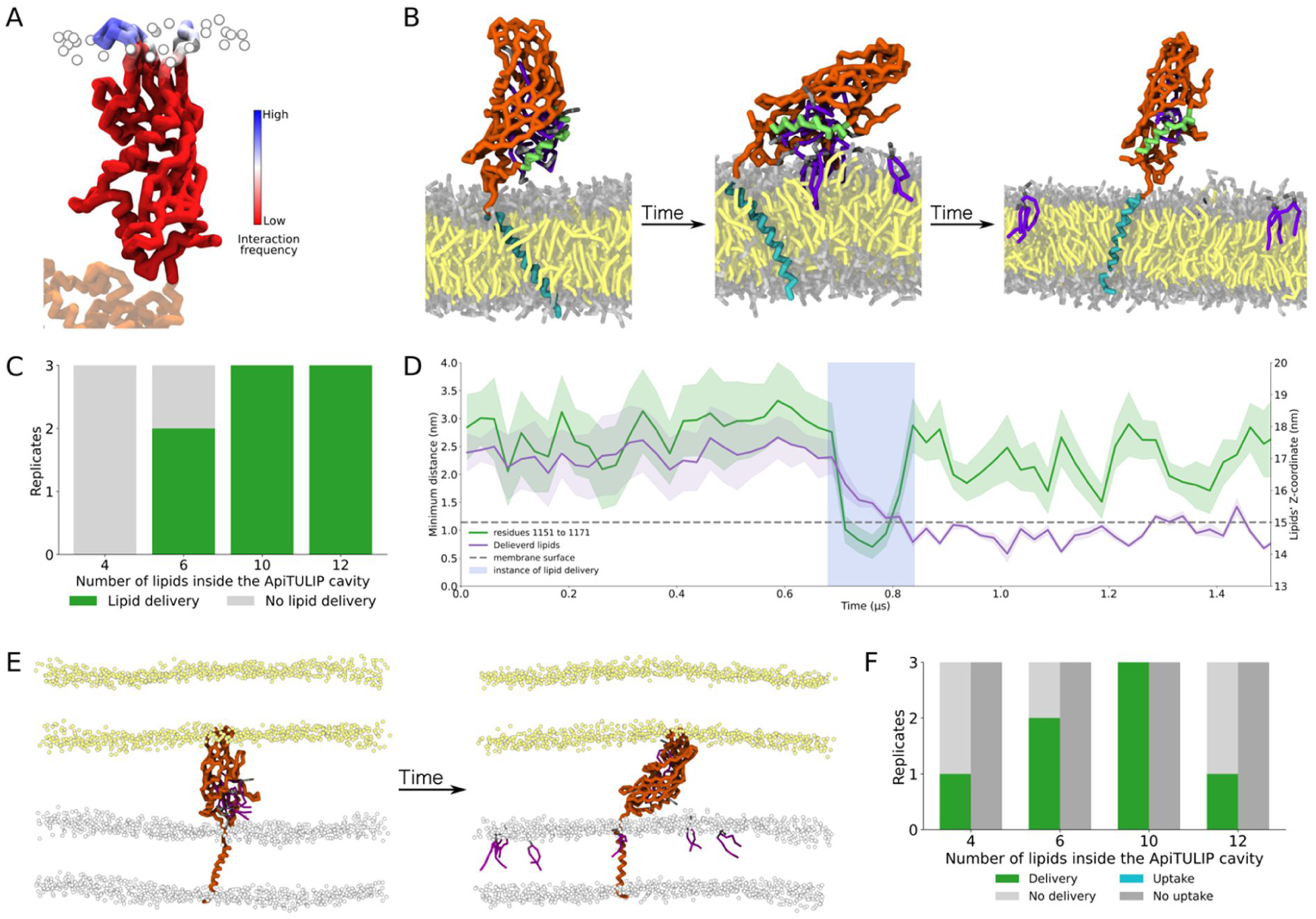
CG-MD simulations of the ApiTULIP domain determine its membrane binding and lipid delivery mechanism. **A**, Representative snapshot showing the membrane binding region (opposite to the TM domain) of the ApiTULIP domain in unbiased CG-MD simulations. The ApiTULIP domain is colored according to the residue’s interaction frequency with the membrane and the rest of the protein (not included in the simulations but added for clarification of the orientation relative to the membrane), is in faint orange. Membrane lipids’ phosphate groups are represented by white beads. **B**, Representative snapshots showing the ApiTULIP domain with 10 POPC molecules in the cavity and the connected TM helix embedded in a membrane showing lipid delivery to the proximal membrane. The initial, intermediate and final states show the ApiTULIP orientation relative to the membrane at different instances during the lipid delivery process. Orange, ApiTULIP domain; green, amphipathic helix; cyan, TM helix; lipids in the membrane: faint yellow (tails) and grey (headgroups); lipids initially in the cavity: purple (tails) and grey (headgroups). **C**, Occurrence of lipid delivery events across the three distinct replicates simulated for systems with different initial lipid occupancies in the ApiTULIP cavity. **D**, Mechanism of lipid delivery (light blue highlight, from one replicate with 10 POPC molecules initially in the cavity) represented by the average z-coordinate of the delivered lipids (purple line, dark traces, right axis) occurring immediately after the amphipathic helix orients closer to the membrane, indicated by the average minimum distance between the helix (residues 1198-1204) and the membrane’s phosphate groups (green line, dark traces, left axis). The light traces indicate the standard deviation across residues/lipids; the membrane surface is indicated by a dashed grey line. **E**, Representative snapshots of the initial and final states of a dual membrane system with 6 POPC molecules in the ApiTULIP cavity, showing lipid delivery towards the DV membrane (with the anchored TM helix) and no lipid uptake from (or delivery to) the membrane on the opposite side. The delivered lipids are shown in purple (tails) and grey (headgroups); the membranes’ surfaces are represented by yellow beads (top) and white beads (DV, bottom). **F**, Occurrence of lipid delivery or lipid uptake events across the three distinct replicates of dual membrane simulations, showing lipid delivery for all initial number of lipids in the ApiTULIP cavity and no lipid uptake events.

The ApiTULIP domain (residues 1145-1371) is directly connected to a TM region, which consists of the PqiA-like domain (residues 471-839) plus one additional TM helix (residues 1113-1144) that is directly connected to the lipid transfer domain. Since the arrangement of the ApiTULIP domain plus the connected TM helix shows structural resemblance to the ER-anchored TULIP domain-containing protein Mmm1 in yeast (Fig. S9C), which was reported to be capable of functioning as a sole lipid transporter at ER-mitochondrial contact sites^71^, we simulated the corresponding truncation of PfTUPA (residues 1113-1371) to investigate the interactions with the proximal membrane and test whether the lipid transport module could deliver lipids to it (Fig. 7B). To explore several saturation conditions, we performed simulations of the ApiTULIP domain filled with 4, 6, 10 and 12 lipids (Fig. 7B-C). Except for the case with 4 lipids, in all other conditions (6, 10 and 12 lipids) lipid release from the peripheral domain into the membrane was observed (Fig. 7C). Mechanistically, lipid release into the membrane followed the interaction of an amphipathic helix from the ApiTULIP domain (residues 1151-1171) with the surface of the membrane (Fig. 7B,D). Once the helix approached the membrane surface, the cavity was positioned in contact with the membrane, allowing the release of the lipids into the bilayer (Fig. 7D, Fig. S9D-E)

Taken together, our MD simulations suggest that (i) the TULIP domain can release lipids into the proximal DV membrane (Fig. 7B-C), and (ii) that it can potentially bind lipids and membranes in its distal end (Fig. 7A). To test whether the ApiTULIP domain could receive its lipids via a membrane associated to its distal end, we constructed an MCS model of the ApiTULIP domain plus the TM helix truncation, in which the TM helix was anchored to one membrane (mimicking the DV) while the opposite end of the protein was positioned in contact with a second membrane (Fig. 7E). Filling the ApiTULIP domain with 4, 6, 10 and 12 lipids as before, lipid release into the helix-anchored membrane was now observed with all lipid content (Fig. 7F). In contrast, while the interaction with the opposing membrane was maintained (Fig. S9F), no lipid uptake from that membrane was observed (Fig. 7E-F).

Next, to characterize the role of the PqiA-like domain in the lipid transport mechanism by PfTUPA, we simulated the full-length PfTUPA protein (Fig. S9G). Microsecond-long all-atom (AA)-MD simulations of full-length PfTUPA showed that, similar to what was observed in LetA MD simulations^58^, lipids were displaced from their bilayer-like orientation and adopted a tilted conformation inside the PqiA-like TM domain (Fig. 8A). Concomitantly water molecules also penetrated deep into the membrane close to the protein structure (Fig. 8A, Fig. S9H). We repeated these simulations with the ApiTULIP domain filled with lipids. However, due to the intrinsic time-scale limitations of AA-MD simulations, lipid delivery was not observed during the simulation duration (Fig. S9I), as the amphipathic helix identified by CG-MD simulations did not sample conformations sufficiently close to the membrane’s surface, (Fig. S9I).

**Figure 8.**
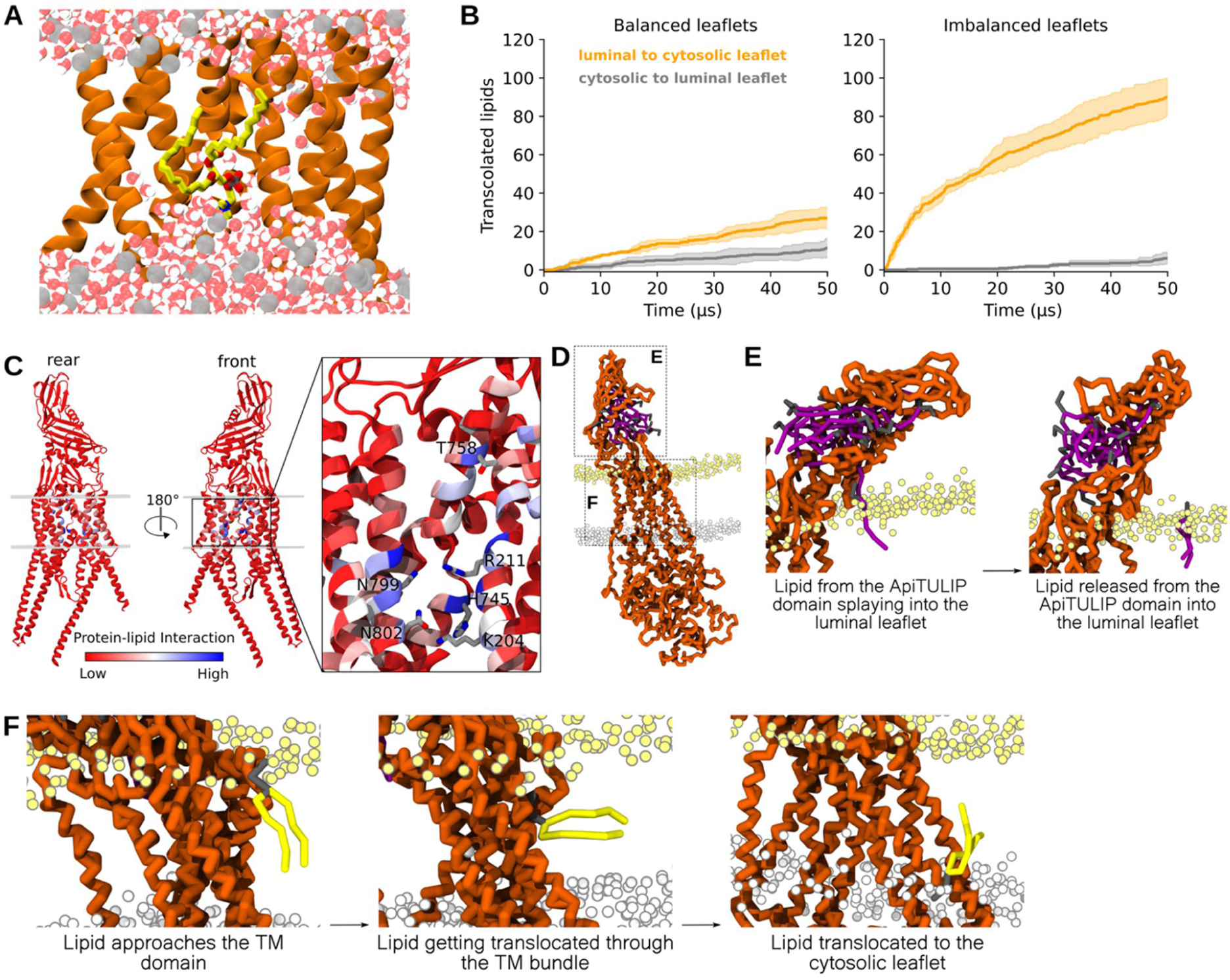
MD simulations suggest that PfTUPA deliver lipids from the DV luminal space to the cytosolic leaflet of the DV membrane. **A**, Representative close-up of an AA-MD snapshot showing lipids in a tilted orientation within the bilayer allowing water molecules to penetrate the membrane near the TM region of PfTUPA. **B**, Lipid translocations mediated by PfTUPA in balanced versus imbalanced membranes shown by the average number of lipids translocated from the luminal-to-cytosolic leaflet (dark yellow traces) and cytosolic-to-luminal leaflet (dark grey traces). The light traces indicate the standard deviation across four independent replicates. **C**, Top, PfTUPA from CG-MD simulations shown from two viewpoints, rear and front, and colored according to lipid interaction to indicate the region of lipid translocation (blue). Bottom, representative close-up of the TM region highlighting the residues that interacted the most with the translocated lipids. **D**, Full-length PfTUPA embedded in a POPC membrane with the ApiTULIP domain filled with 12 POPC molecules. **E**, Representative close-up views of the ApiTULIP domain showing the steps of the lipid delivery process during the simulation. Lipid delivery begins by one lipid from ApiTULIP splaying into the luminal leaflet of the membrane followed shortly by its release into the leaflet. **F**, Initial, intermediate, and final snapshots of a close-up view on the TM domain of PfTUPA showing the steps of lipid translocation from the luminal leaflet to the cytosolic leaflet. Lipid translocation occurs when a lipid approaches the TM region of translocation where its headgroup interacts with polar residues that facilitate its movement across the hydrophobic bilayer via maintaining the polar interactions, eventually translocating the lipid to the cytosolic leaflet. In panels D, E, and F, the protein is shown in orange; the translocated lipid is shown in yellow (tails) and grey (headgroups) and lipids initially in the cavity of the ApiTULIP domain are shown in purple (tails) and grey (headgroups). The luminal leaflet is represented by yellow beads, and the cytosolic leaflet is represented by grey beads.

We next asked whether the PqiA-like TM domain can redistribute the lipid imbalance created by lipid delivery from the ApiTULIP domain (Fig. 7B,E). To address this mechanism, we performed CG-MD simulations under two conditions: (i) PfTUPA embedded in an almost symmetric membrane, and (ii), PfTUPA embedded in a membrane with ∼10% excess lipids in the luminal leaflet. Our CG-MD simulations revealed that the PfTUPA TM domain preferentially translocates lipids from the luminal to the cytosolic leaflet (Fig. 8B, left panel) and that this activity is largely enhanced when the luminal leaflet has excess lipids (Fig. 8B, right panel). These translocations occur preferentially at a specific region of the protein (Fig. 8C), which corresponds to the same region previously reported for bacterial LetA to mediate lipid translocation^58^. At this interface, several polar amino acids appear to be strongly involved in lipid translocations, including K204, R211, H745, T758, N799, and N802 (Fig. 8C).

Finally, to concomitantly investigate the two molecular functions of PfTUPA, lipid delivery by the ApiTULIP domain, and transbilayer lipid translocation by the PqiA-like TM domain, we performed CG-MD simulations of the full-length protein in which the ApiTULIP domain was filled with lipids. These simulations revealed both lipid delivery into the luminal leaflet of the membrane and lipid translocation events (Fig. 8D-F), albeit not for the same lipids. Taken together, our simulations propose a mechanism where the various domains of PfTUPA work in tandem to deliver lipids from the DV luminal space to both leaflets of its surrounding membrane.

## Discussion

Despite an increasing realization of the importance of protein-mediated lipid transfer processes in eukaryotic cell biology and evidence that non-vesicular lipid transport is the dominant mechanism maintaining the homeostasis of the cellular lipid distribution^1^, these processes remain poorly studied in protists, including organisms of high medical importance such as *P. falciparum* parasites. Establishment of the RBC infection by malaria parasites involves extensive membrane rearrangements^24^ and endocytosis of large amounts of host cell cytosol^30^, consuming it almost to completion by the end of the cycle. The ingested hemoglobin is transported in double membrane vesicles^43,44,72^, an inner membrane originating from the PVM and an outer membrane from the PPM, leading to internalization of large quantities of membrane material (Fig. 5G). How the parasite deals with these membranes is not known and due to efficient removal under physiological conditions, the large amounts of membrane arriving at the DV have never been apparent up to date.

In a screen of LTPs we here identified an essential DV membrane protein we named PfTUPA. PfTUPA turned out to be a member of a class of proteins named TUPALs that contain a region homologous to PqiA, an intramembrane domain of bacterial origin recently shown to have lipid transfer function^58^, fused to a TULIP-like lipid transfer module. We detected TUPALs in different clades ranging from plants to opisthokonts and to diverse protist groups. Strikingly, the arrangement of the two domains is different between evolutionary clades and involves diverse small lipid transfer domains, suggesting this combination might have arisen multiple times during evolution. It is possible that the PqiA domain of bacterial origin provides a versatile building block co-opted for lipid-transfer functionalities in eukaryotes that are similar to that of the bacterial PqiABC operon, which includes a second lipid transfer protein exposed to the periplasm (the periplasmic exposed PqiB) that interacts with PqiA to enable lipid exchange between the inner and outer bacterial membranes^4,58,59^. In this scenario, eukaryotic TUPALs may serve analogous functions and the diversity of the associated TULIP-like domains and their arrangement may reflect the different tasks and target membranes in their organism.

Our data indicates that PfTUPA, the Plasmodial TUPAL, is needed to dispose membranes arriving in the DV lumen due to endocytosis by transferring lipids between the DV lumen and its membrane (Fig. 5G). Importantly the large build-up of membranes in the *PfTUPA* cKO highlights that considerable amounts of lipids can be mobilized by this type of LTP. According to our model, the TULIP-like domain facing the lumen picks up lipids, either as free products of an unknown DV lipase or directly from the intraluminal membranes via a MCSs in a bridge-like manner, and the PqiA domain distributes them between the two leaflets of the DV membrane. Our study is the first investigated protein example where a PqiA-like domain is directly fused with an exposed lipid transferring TULIP-like domain and the first evidence for the conservation of such a function for PqiA-containing proteins in eukaryotes, opening the possibility that other proteins containing PqiA-like domains and its remote homologs (such as the human tetraspanin proteins^58^) could cooperate with TULIP-like domains in other eukaryotic clades to fulfill similar functions.

Depletion of PfTUPA leads to a swollen DV, initially consisting primarily of membranous structures, concordant with a build-up of lipids due to their failed transfer when PfTUPA is missing (Fig. 5G). It is possible that the accumulation of membranes leads to a more general disruption of the DV function which is reflected in the later accumulation of hemoglobin-filled vesicles, resembling previously observed phenotypes of defective DV function^45,47^. However, we cannot fully exclude a more direct function of PfTUPA in hemoglobin release or digestion in the DV.

While the DV of malaria parasites processes a particularly high amount of intraluminal membranes, other eukaryotes also experience related challenges. In opisthokonts, processing of autophagy-derived vesicles or the endocytosis of low density lipoproteins also lead to the delivery of lipids to the vacuole/lysosome, where they are catabolized by lipases^73,74^ and the products effluxed^75,76^. Cholesterol esters, in particular, are metabolized and the free cholesterol molecules are transported from the lumen to the membrane of the vacuole or lysosome by the function of the membrane-anchored lipid transfer protein NPC1^77^, homolog of the *P. falciparum* protein PfNCR1^28^. The function of opisthokont NPC1 could be similar to the one reported here for PfTUPA, with the difference that TULIP-like domains are known to transfer phospholipids. Further studies should clarify whether PfTUPA has preference for a certain lipid species and how these lipids reach the lipid transfer domain.

In conjunction with previous studies^25,28,29^, our work provides a nearly complete overview of all at present identifiable LTPs in the RBC stages of the parasite. As several of these genes are important for growth and found at MCSs, it establishes a basis to unravel these critical mechanisms in apicomplexan cell biology, with potential applications in drug development, as has been already shown for two LTPs^26,27^. PfTUPA, in particular, functions in the DV which is a target of drugs and drug resistance^38^ and indeed mutations in the *P. chabaudi* homolog were identified as conferring chloroquine (CQ) resistance^78^. Although this mutation did not affect CQ IC_50_ in *P. falciparum* (Fig. S10), it nevertheless highlights potential relevance for drug development and resistance.

Overall the study of malaria proteins with small LTP domains revealed PfTUPA, the first functionally studied member of TUPALs, a diverse new class of LTPs consisting of an intramembrane lipid transporter module of bacterial origin fused to a TULIP-like domain with a wide evolutionary distribution.

## Methods

### Parasite culture, synchronization and transfection

*P. falciparum* 3D7 parasites were cultured with 5% hematocrit of human O+ erythrocytes in RPMI1640 with 0.5% Albumax II (Gibco #11021045), 12 mM sodium bicarbonate, 10 mM Glucose and 20 μg/mL Gentamycin (Ratiopharm). Cells were kept in a controlled gas environment of 5% O2, 5% CO2, 90% N2 at 37 °C, following standard culture protocol. Transfusion human O+ blood was commercially purchased from Universitätsklinikum Hamburg-Eppendorf (Approval number 10569a/96-1). Age, sex and identity of blood donors was unknown.

Synchronization was performed by incubation on 5% sorbitol for 10 minutes, for ring stage parasites, or by separation with 60% Percoll and centrifugation, for schizont stage parasites. Transfections were performed on Percoll-enriched late schizonts (originating from 400 μl of erythrocytes at 5% parasitaemia per transfection). Briefly, parasites were resuspended in 90 μl of transfection buffer (90 mM NaPO4, 5 mM KCl, 0.15 mM CaCl2 and 50 mM HEPES pH 7.6) and electroporated with 50 μg of plasmid in 10μl of TE buffer, using the Amaxa system (Lonza Nucleofector II AAD-1001N, program U-033) as previously described^79^. A drug resistance gene was encoded in the plasmid for selection of positively transfected parasites. Drugs used were 4 nM WR99210 (Jacobus Pharmaceuticals), 0.9 μM DSM1 (Merck #5.33304.0001), 400 μg/mL G418 (Merck #A1720), or 2 μg/μl blasticidin S (Invitrogen #R21001) for plasmids encoding the hDHFR, yDHODH, Neo-R, or BSD resistance genes, respectively.

### Plasmids

Plasmids generated for this study were cloned by Gibson ligation reaction (a mixture of T5 exonuclease (Epicenter #T5E4111K), Phusion DNA polymerase (NEB #M0530S), Taq DNA ligase (NEB #M0208L), dNTPs (Roth #K039.1), NAD (Sigma-Aldrich #n6522), DTT, MgCl_2_ and Tris-HCl pH 7.5), the backbones were linearized with restriction enzymes and the inserts were amplified by PCR with primers containing 20nt of overlap with the backbone. Oligonucleotides were ordered from Sigma-Aldrich, restriction enzymes from NEB and in cases where a large (above 120aa) recodonized sequence was required, synthesized DNA fragments were ordered from BioCat GmbH (Heidelberg, DE). A summary of all the plasmids (and the respective cloning reactions) generated in this study is found in Table S2, and the expected full plasmid sequences were deposited in a public database^80^.

Other plasmids used for transfections in this work, mSc-TmSec61β diCre, mCh-FRB-miniTurbo, mSc-TmSec61β NLS-FRB, IMC-mCh NLS-FRB, or used as templates or backbones for further cloning reactions, pSLI-N-GFP-2xFKBP-loxP (Addgene #85792), pSLI-C-GFP-Sandwich (also pSLI-sandwich, Addgene #85790), pSLI-N-Halo, pSLI-C-Halo, pSLI-N-PfPITP-mNG, crt-P40-mSc NLS-FRB, pSLI-N-GFP-3’UTR-mCh, were previously generated in our laboratory^29,42,44,54^, SEP DNA sequence was a kind gift from J. Matz (BNITM, Hamburg, DE).

### Selection linked integration (SLI)

All genes analysed in this study were endogenously edited at their endogenous loci using the SLI approach, with slight differences depending on whether the modification was N-or C-terminal (as shown in Fig. S1A). SLI was carried out as described^54^. Parasites were transfected with a SLI plasmid and selected with WR99210. Successfully transfected and growing parasites were then treated with a different drug to which only parasites that had integrated the plasmid in their genome were resistant (G418, in the case of C-terminal gene modifications, or DSM1, in the case of N-terminal gene modifications. Once integrated parasites resurfaced and were growing at usual rates, integration was evaluated by a diagnostic PCR from genomic DNA (collected using Monarch #T3010L kit). Diagnostic PCR involved reactions testing the absence of the unmodified gene locus and spanning the two integration junctions, performed using Firepol enzyme (Solis biodyne #01-01-02000). Genomic sequences of the original and modified loci, including the primers and amplicon sizes for diagnostic PCR, are provided in Data S1.

### Labelling of parasites and live-cell microscopy

For direct staining, parasites were incubated with 50 ng/mL Hoechst 33342 (Biomol #ABD-17533), and/or 50 nM Halo-JF571, -JF585, -JFX650 or -JFX673 ligands^81^ (kind gift from L. Lavis, Janelia Farm, Ashburn VA or Promega #HT1040/#HT1070) for nuclear and/or Halo-tag visualization, respectively. Labelling was carried out for 20 minutes at 37 °C in RPMI-1640 medium after which the parasites were washed with medium, with an additional 10 minute incubation and wash for Halo labelling. For imaging experiments involving pharmacological inhibition of the V-ATPase, parasites were incubated with 50 nM of Concanamycin A (MedChem #HY-N1724) for 30 minutes. For imaging, parasites in RPMI medium were placed between a coverslip and a glass slide.

Most of the wide-field fluorescence microscopy was carried out in a Zeiss AxioImager microscope equipped with a Hamamatsu Orca C4742-95 camera and a plan apochromat objective (63x, 1.4 NA, Oil DIC), and using the AxioVision software (version 4.7). Wide-field fluorescence microscopy of line PfCT3-mNG was carried out in a Leica DM6 B equipped with a K8 CMOS camera and a plan apochromat objective (100x, 1.4 NA, Oil), and using the Leica Application Suite X (LAS X) software (version 3.9.1.28433, Leica Microsystems). Laser scanning confocal microscopy was carried out in an Olympus FluoView FV3000 system equipped with a universal plan apochromat objective (60×, 1.5 NA, oil) and a cell Vivo incubation system (at 37°C). All images were processed with FIJI^82^ to adjust for brightness and contrast and in the case of confocal images apply a gaussian filter.

### Ultra-expansion microscopy (U-ExM)

U-ExM was performed as previously described^22^. Briefly, parasites were allowed to attach for 20 minutes on 3 mm PDL-coated coverslips at 37°C. After a PBS wash, samples were fixed with 4% formaldehyde (Electron Microscopy Sciences), for NHS-ester membrane staining, or 4% formaldehyde/0.0075% glutaraldehyde (Electron Microscopy Sciences), for NHS-ester protein staining, in PBS for 20 minutes, and left overnight in post-fix solution [1.4% formaldehyde and 2% acrylamide (Merck #A4058)], all at 37 °C. Coverslips were washed and placed on 35 μl of monomer solution [19% sodium acrylate (Sigma-Aldrich #408220)}, 10% acrylamide and 0.1% N,N’-methylenebisacrylamide (Merck #M1533)] with freshly added 0.5% TEMED (EMD Millipore #1.10732) and 0.5% APS (Sigma-Aldrich #A3678). Gel polymerization was carried out at 37 °C for 1 hour and gels were then separated from the coverslip by shaking at RT on denaturation buffer (200 mM SDS, 200 mM NaCl, 50 mM Tris-HCl pH 9). Gel denaturation was carried out for 90 minutes at 95°C, after which they were either frozen in 50% PBS/glycerol or placed in water for three rounds of expansion of 30 minutes each. The gels were shrunk in PBS and a slice was cut for antibody staining. This slice was blocked with 3% BSA (Biomol #9048-46-8) in PBS for 30 minutes at RT, and left incubating with NHS-ester (1/250, Thermo Fischer #46403) and Hoechst 33342 in PBS (for NHS-ester protein staining) or 3%BSA-PBS (for NHS-ester membrane staining) overnight at RT. Gels were washed four times with 0.5% Tween in PBS and placed in water for three rounds of expansion. Gels were placed on a PDL-coated glass-bottom dish (Ibidi #80427) for imaging in an Olympus FluoView FV3000 laser scanning confocal microscope equipped with a universal plan super apochromat objective (60x, 1.3 NA, silicone). Images were processed using FIJI^82^ to adjust brightness and contrast, apply a gaussian blur filter of 0.5 pixel radius and obtain Z-stack max intensity projections of the number of Z-slices specified in each image.

### Transmission electron microscopy

Transmission electron microscopy was done as previously described^43^. Pf*tupa* cKO parasites, induced in ring stages in the previous cycle, were fixed with 2.5% glutaraldehyde in 50 mM cacodylate buffer (pH 7.4) for 1h at RT, at either trophozoites (29-37 hours post infection) or schizonts (45-49 hours post induction). For postfixation and staining of the cell membranes, samples were incubated with 2% osmium tetroxide (OsO4) in dH20 for 40 min on ice in the dark. After 3 washes with dH20, samples were incubated with uranyl acetate (Agar Scientific) for 30 min at room temperature, washed 2 times with dH20 and dehydrated in an ethanol series (50%, 70%, 90% (2×), 95% (3×), and 100%) for 5 min each. Samples were then incubated with an epon-ethanol mixture (1:1) overnight at room temperature shaking, followed by an incubation with 100% epon (Carl Roth GmbH & Co. KG) for 6 h. Epon was replaced and left to polymerize at 60°C for 1 to 3 days. Samples were cut into 60 nm sections with an Ultracut UC7 (Leica) and examined with a Tecnai Spirit transmission electron microscope (FEI), equipped with a LaB6 filament and operated at an acceleration voltage of 80 kV. The EM image.emi/.ser-files were converted to 8-bit TIFF files using the TIA Reader Plugin for ImageJ.

## DiQ-BioID

### Sample preparation

DiQ-BioID was carried out in an asynchronous culture of GFP-2xFKBP-PfTUPA^endo^ parasites episomally expressing miniTurbo proximity biotinylation enzyme, as previously done^29,42^. Briefly, 200 mL of culture in biotin-free RPMI-1640 medium (Biozol #USB-R9002-01) were split into two, and in one half the dimerization of miniturbo with PfTUPA was induced by addition of 250 nM rapalog. Immediately after, 50 μM of biotin (Sigma-Aldrich #B4639) was added to both halves for 30 minutes of labelling. Thereafter, RBCs were harvested and parasites were isolated by lysis with 0.03% saponin (in 1xPBS) on ice for 10 minutes, washed five times with 1xPBS and lysed in 2 mL of lysis buffer (50mM Tris-HCL pH 7.5, 500 mM NaCl, 1% Triton-X-100, 0.4% SDS) supplemented with 1 mM DTT, 1 mM PMSF and 1 x protease inhibitor cocktail (Roche #11836170001). The lysate was frozen at -80 °C until affinity purification was carried out.

After 3 freeze-thaw cycles, lysates were centrifuged at 16,000xg for 60 minutes to clear the lysate, at 4°C. The supernatant was diluted 3-fold in 50 mM Tris-HCl pH 7.5 and incubated with 50 μl of pre-treated (for protease-resistance^83^) streptavidin sepharose beads rotating overnight at 4°C. The beads were washed twice with lysis buffer, once with dH_2_0, twice with Tris-HCl pH 7.5 and three times in 100 mM TEAB (Sigma-Aldrich #T7408) pH 8.5, before adding 50 μl of elution buffer (2 M urea and 10 mM DTT in 100 mM Tris-HCl pH 7.5) and incubation for 20 minutes with shaking at 1400rpm at RT. For protein alkylation, 5 μl iodoacetamide to a final concentration of 50 mM was added to the beads and incubated for 10 minutes shaking in the dark, and on-bead digestion was carried out then by adding 500 ng of Trypsin (Sigma-Aldrich #T0303) and shaking for 2 hours, all at RT. The supernatant was collected and the beads were incubated with fresh 50 μl of elution buffer for 15 min, and the new supernatant was combined with the previous one. 200 ng of additional trypsin was added and left shaking at RT overnight. Tryptic peptides were frozen at -80°C and shipped to the EMBL proteomics core facility (Heidelberg, DE), for MS analysis.

### Offline fractionation

Peptides were dried and reconstituted in 10 µl of 100 mM Heps/NaOH, pH 8.5 and reacted for 60 min at RT with 80 µg of TMT6plex (Thermo Fischer #90066) dissolved in 4 µl of acetonitrile. Excess TMT reagent was quenched by the addition of 4 µl of an aqueous 5% hydroxylamine solution (Sigma-Aldrich #438227). Peptides were reconstituted in 0.1 % formic acid, mixed and purified by a reverse phase clean-up step (OASIS HLB 96-well µElution Plate, Waters #186001828BA). Offline high-pH reversed-phase fractionation^84^ was carried out using an Agilent 1200 Infinity high-performance liquid chromatography (HPLC) system, equipped with a Gemini C18 analytical column (3 μm particle size, 110 Å pore size, dimensions 100 x 1.0 mm, Phenomenex) and a Gemini C18 SecurityGuard pre-column cartridge (4 x 2.0 mm, Phenomenex). The mobile phases consisted of 20 mM ammonium formate adjusted to pH 10.0 (Buffer A) and 100% acetonitrile (Buffer B). The peptides were separated at a flow rate of 0.1 mL/min using the following linear gradient: 100% Buffer A for 2 minutes, ramping to 35% Buffer B over 59 minutes, increasing rapidly to 85% Buffer B within 1 minute, and holding at 85% Buffer B for an additional 15 minutes. Subsequently, the column was returned to 100% Buffer A and re-equilibrated for 13 minutes. During the LC separation, 48 fractions were collected. These were pooled into six fractions by combining every sixth fraction. The pooled fractions were then dried using vacuum centrifugation.

### LC-MS/MS analysis

An UltiMate 3000 RSLCnano LC system (Thermo Fisher Scientific) equipped with a trapping cartridge (µ-Precolumn C18 PepMap™ 100, 300 µm i.d. × 5 mm, 5 µm particle size, 100 Å pore size; Thermo Fisher Scientific) and an analytical column (nanoEase™ M/Z HSS T3, 75 µm i.d. × 250 mm, 1.8 µm particle size, 100 Å pore size; Waters) was used. Samples were trapped at a constant flow rate of 30 µL/min using 0.05% trifluoroacetic acid (TFA) in water for 6 minutes. After switching in-line with the analytical column, which was pre-equilibrated with solvent A (3% dimethyl sulfoxide [DMSO], 0.1% formic acid in water), the peptides were eluted at a constant flow rate of 0.3 µL/min using a gradient of increasing solvent B concentration (3% DMSO, 0.1% formic acid in acetonitrile). The gradient was as follows: 2% to 8% in 6 minutes (min), 8% to 28% in 72 min, 28% to 40% in 4 min, 40-80% in 0.1 min, washed at 80 % for 2.9 min and re-equilibrated to 2% B for 5 min.

Peptides were introduced into an Orbitrap Fusion™ Lumos™ Tribrid™ mass spectrometer (Thermo Fisher Scientific) via a Pico-Tip emitter (360 µm OD × 20 µm ID; 10 µm tip, CoAnn Technologies) using an applied spray voltage of 2.4 kV. The capillary temperature was maintained at 275 °C. Full MS (MS1) scans were acquired in profile mode over an m/z range of 375–1,500, with a resolution of 60,000 at m/z 200 in the Orbitrap. The maximum injection time was set to 50 ms, the AGC target limit was set to ’standard’. The instrument was operated in data-dependent acquisition (DDA) mode, with MS/MS scans acquired in the Orbitrap at a resolution of 15,000. The maximum injection time was set to 54 ms, with an AGC target limit of 200%. Fragmentation was performed using higher-energy collisional dissociation (HCD) with a normalized collision energy of 36%. MS2 spectra were acquired in profile mode. The quadrupole isolation window was set to 0.7 m/z, and dynamic exclusion was enabled with a duration of 60 seconds. Only precursor ions with charge states 2–7 were included for fragmentation.

### Database search and analysis

Raw files were converted to mzML format using MSConvert from ProteoWizard, using peak picking, 64-bit encoding and zlib compression, and filtering for the 1000 most intense peaks. Files were then searched using MSFragger in FragPipe^85^ (22.1-build02) against FASTA database Uniprot Plasmodium falciparum isolate 3D7 fasta databas e (UP000001450, ID 36329, 5372 entries, date: 06.03.2023, downloaded: 17.05.2023) containing common contaminants and reversed sequences. The following modifications were included into the search parameters: Carbamidomethylation (C, 57.0215), TMT (K, 229.1629) as fixed modifications; Oxidation (M, 15.9949), Acetylation (protein N-terminus, 42.0106), TMT (peptide N-terminus, 229.1629) as variable modifications. For the full scan (MS1) a mass error tolerance of 20 pp m and for MS/MS (MS2) spectra of 20 ppm was set. For protein digestion, ’trypsin’ was used as protease with an allowance of maximum 2 missed cleavages requiring a minimum peptide length of 7 amino acids. The false discovery rate on peptide and protein level was set to 0.01. The standard settings of the FragPipe workflow ’Default’ were used.

The FragPipe output was processed using R (ISBN 3-900051-07-0). Proteins quantified with >1 razor peptide were considered (contaminants and reverse proteins filtered out). ’removeBatchEffect’ of the limma package^86^ was used to clean Log2 transformed raw TMT reporter ion intensities from batch effects and ’normalizeVSN’ for further normalization the (VSN -variance stabilization normalization^87^). Testing for differential expression of proteins was done with a moderated t-test using ’lmFit’ and ’eBayes’ of the limma package. Replicate information was factored as an argument in the design matrix of the ’lmFit’ function of limma. Proteomics results have been deposited to the ProteomeXchange Consortium via the PRIDE partner repository with the dataset identifier PXD077762.

### Loss-of-function induction by knock sideways or conditional gene excision

Knock-sideways (KS) was used to study the loss-of-function of soluble proteins as done previously ^54^. This was induced by addition of 250 nM Rapalog to a cell line expressing the POI endogenously fused to 4 copies of the FKBP domain and an episomal FRB protein fused to a nuclear localization signal (NLS). Mislocalization to the nucleus was assessed by microscopy after 4 hours of induction or, in the case of PF3D7_0911100, after 20 hours. Mislocalization was considered partial when there was protein signal in its endogenous localization in addition to the nucleus. For the percentage presented in Fig. 1D, cells with partial mislocalization were given half the weight as a full mislocalization.

Gene excision (conditional KO, cKO) was induced by addition of 250 nM Rapalog to a cell line with two loxP sites at the endogenous locus of the POI (Fig. S1A) and an episomally expressed diCre protein^54^. For all Pf*tupa* cKO experiments, excision was induced at rings (either 0-4hpi or 0-10hpi, depending on synchronization method) and analysis were performed in the following cycle, were the protein expression was first lost. Efficiency in Fig. 1D was determined as percentage of cells with no protein expression, assessed by microscopy. Most Pf*tupa* cKO experiments were performed with the GFP-2xFKBP-PfTUPA^endo^ cell line, with the exception of the acidification experiment in Fig. 5F-G, which was done with the Halo-PfTUPA^endo^ cell line.

### Parasite growth assays

For protein essentiality assays, the growth of asynchronous (for KS) or double sorbitol-synchronized (by two 10-minute incubations with 5% sorbitol 10 hours apart, for diCre-mediated cKO) cultures (control vs plus rapalog) was measured every day over 7 (for diCre excision) or 5 (for KS) days. Considering excision happens during the first cycle in cKO, this would be time for 2 cycles with protein loss of function in both cases. For IC50 assays, an asynchronous culture with 0.8% of parasiteamia was exposed to a range of CQ (0 to 100 nM) concentrations for 48h, with replenishment of fresh medium and CQ at 24h. Growth was assessed to % of parasiteamia in untreated control.

For parasiteamia measurements, samples were stained with Hoechst 33342 and Dihydroethidium (Cayman #12013-5) for 20 minutes at room temperature and measured as the percentage of positively stained cells over 100000 events counted with an LSRII flow cytometer (BD Biosciences) with a FACSDiva software (BD Biosciences) as described^54,88^. Curves were plotted using Graph Pad Prism. For stage specific growth assays Giemsa (Merck) smears were taken at the indicated time points.

### Sequence and structural analysis

Protein sequences were retrieved from PlasmoDB (release 67)^89^. For conservation analysis across *Plasmodium*, apicomplexans and other related alveolate species (Fig. S4C), a maximum-likelihood phylogenetic tree was generated in MEGA11^90^ with 100 bootstrapping repeats from an alignment of PfTUPA homologs identified in reference genomes using BLAST. TULIP-like sequences found across different eukaryotes downloaded from Interpro were clustered using CLANS^91^. The sequences used for these analyses were deposited in a public database^80^.

LTPs were identified by sequence and structural homology. For sequence-based searches, alignments of the different small lipid transfer domains from opisthokonts were used to query the *P. falciparum* proteome using HHpred^92^, and for structure-based searches, solved structures of the different domains were downloaded from the PDB and used to query the *P. falciparum* proteome using Foldseek^53^. Structures of the identified LTPs were downloaded from the AlphaFold public database^60,93^ and analysed using PyMOL^94^ and UCSF ChimeraX^95^ (developed by the Resource for Biocomputing, Visualization, and Informatics at the University of California, San Francisco, with support from National Institutes of Health R01-GM129325 and the Office of Cyber Infrastructure and Computational Biology, National Institute of Allergy and Infectious Diseases). Structural alignments between two indicated proteins was done using US-align^57^. All vs all structural alignments of a sample of 50 domains found in eukaryotic PqiA-containing proteins downloaded from The Encyclopedia of Domains (TED^96^) was done using the DALI server^62^. The domain structures used for this alignment were deposited in a public database^80^.

### Molecular dynamics simulations

#### Structure prediction

All structures were predicted using the Alphafold3 server^97^. To predict the ApiTULIP domain alone, residues P1145-Y1371 from Uniprot accession number O96228 were used as sequence input. The average predicted local distance difference test (pLDDT) score of this domain was 85.85. To create the L1232W/V1320W double-mutant of the ApiTULIP domain, chimera was used to directly introduce the mutations to the predicted structure. To include the TM helix that is connected to the peripheral ApiTULIP domain, residues E1113-Y1371 were used, providing a prediction with an average pLDDT of 81.47. Finally, to predict the full-length structure, the complete sequence was used as input, resulting in a prediction with an average pLDDT of 51.4. To improve the prediction, we omitted the unstructured loops from the input sequence and modeled the three structured domains: the Nter PqiA domain (residues M1-E300), the Cter PqiA domain (residues D471-D839) and the ApiTULIP domain with the connected TM helix (residues E1113-Y1371) separately as a multimer complex. The resulting prediction had an average pLDDT of 67.7. The disordered loops were then re-integrated into the structure using SWISS-MODEL^98^.

### Systems setup

#### CG-MD lipid binding assay

The atomistic structures were converted to CG using the martinize script^99^. An elastic network with a force constant of 500 kJ mol^−1^ nm^−2^ was applied to retain the secondary structure of the ApiTULIP domain in CG, with upper and lower elastic bond cutoffs of 0.8 and 0 nm, respectively^68^. The ApiTULIP domain was centered in a cubic box, with a minimum distance of 2 nm between the domain and each box edge. One POPC molecule was randomly placed in the bulk solvent without applying any distance restraints, independently randomizing the position of the lipid in the box for each replicate. The system was then solvated using a van der Waals cutoff distance of 0.21 nm and ionized with 0.12 M NaCl. The same protocol was followed for the double-mutant domain. To test the domain’s propensity to bind multiple lipids, POPC addition was performed iteratively, with the protein structure containing n bound POPCs serving as the starting configuration for the addition of the (n + 1)th POPC. Following each lipid insertion, the system was re-solvated and re-ionized before proceeding to the next iteration. This process was repeated until a total of 12 lipids were present in the system. Five independent replicates were simulated for each iteration for 1 µs each.

### CG-MD membrane binding assay

To determine the membrane associating region of the ApiTULIP domain, four independent systems were simulated at CG resolution. The elastic network of the domain was kept as previously described. The insane Python script was used to generate the bilayers for CG-MD systems^100^. A POPC membrane was generated in a box of 16 nm x 16 nm x 20 nm in x, y and z, respectively. In the first system, the empty ApiTULIP domain was positioned at least 2.5 nm away from the membrane. In the second, the empty domain was replaced with the lipid-loaded structure obtained from the lipid binding assay simulations, containing six POPC molecules within its cavity, while maintaining the same initial position relative to the membrane. In the third, the empty ApiTULIP domain was placed in a membrane-bound orientation predicted by the OPM server^70^, which was used to determine the domain’s orientation with respect to the membrane. Finally, the fourth system consisted of the membrane-bound ApiTULIP domain loaded with six POPC molecules. The systems were solvated and ionized with 0.15M NaCl. All systems were then equilibrated using a five-step protocol that progressively increased the time step while gradually reducing the position restraints on the lipid headgroups, ultimately removing all restraints in the final step to allow the lipids to move freely. Restraints on the protein backbone were treated in a similar manner. Three independent replicates of each system were simulated for 8 µs each.

### CG-MD lipid delivery assay

The truncation of PfTUPA (residues 1113-1371) that includes the ApiTULIP domain and the TM helix that is directly connected to it were embedded in a POPC membrane in a simulation box with dimensions 20 nm x 20 nm x 35 nm in x, y and z, respectively. An elastic network with a force constant of 700 kJ mol^−1^ nm^−2^ was applied to retain the secondary structure. The ApiTULIP domain was kept empty and was filled with 4, 6, 10, and 12 POPC molecules, giving a total of 5 distinct systems. The systems were then solvated and ionized with 0.15M NaCl. Equilibrations were performed as previously described. Three independent replicates of each system were simulated for 8 µs each.

To create the double membrane systems of this PfTUPA truncation, a POPC membrane of the same dimensions was added to the opposite end of the simulation box, where it associated with the region of the ApiTULIP domain that is opposite to the TM helix (∼6 nm away from the first membrane). A pore with a 2 nm diameter was present in the newly-added membrane to allow uninterrupted water and ion exchange and ensure sufficient equilibration. The pore was eventually sealed after sequential equilibrations during which the lipid tails were restrained to stay away from the pore using a flat-bottom restraint potential^101^. The systems were then solvated and ionized with 0.15 M NaCl. Three independent replicates of each system were simulated for 10 µs each.

### CG-MD local lipid remodeling simulations

Full-length PfTUPA was simulated using an elastic network, excluding the unstructured regions, with a force constant of 500 kJ mol^−1^ nm^−2^. The membrane was composed of 100% POPC with dimensions of 30 nm x 30 nm x 36 nm in x, y, and z, respectively. PfTUPA was oriented in the membrane according to the OPM prediction. One set of systems included the empty ApiTULIP domain while another set included the lipid-filled (12 POPC molecules) ApiTULIP domain. The systems were then solvated and ionized with 0.15 M NaCl. Equilibrations were performed as previously described. Four independent replicates of each system were simulated. The systems with the empty ApiTULIP domain were simulated for 50 µs each while the systems with the filled ApiTULIP domain were simulated for 20 µs each. For the simulations with imbalanced leaflets, the luminal leaflet proximal to the ApiTULIP domain contained an excess of 10% (∼200 lipids) relative to the cytosolic leaflet.

### Simulation details CG-MD

All CG-MD simulations were performed with the GROMACS package^102^ using the Martini 3 force field^103^. All systems were initially minimized using a steepest descent algorithm. For all systems except the lipid binding assay simulations, an NPT equilibration was run for 250 ps with restraints on the protein backbone before production runs. Instead, for the lipid binding assay simulations of the ApiTULIP domain, an MD run of 125 ps was performed prior to the production run. For all systems, simulations were kept at 310K using a velocity-rescale thermostat^104^ that separately coupled the protein, membrane (where applicable), and solvent. The Parrinello–Rahman barostat^105^ was used to maintain the pressure constant at 1 bar using a semi-isotropic pressure coupling scheme for membrane-containing systems and an isotropic scheme for systems without membranes. To calculate the nonbonded interactions, a Verlet scheme with a buffer tolerance of 0.005 was used. Reaction-field electrostatics was used to compute the coulombic interactions, while the cutoff method was used for the van der Waals terms. Both former terms have a cutoff distance of 1.1 nm and follow the Verlet cutoff scheme for the potential shift. A time step of 20 fs was used with the md integrator.

### AA-MD

The CHARMM36 force field^106^ was used in combination with the GROMACS package^107^ for AA-MD simulations. The membrane builder tool on CHARMM-GUI^108^ was used to build the system of PfTUPA embedded in a membrane in AA. Systems were initially minimized for 5,000 steps. Next, two equilibrations in the NVT ensemble were run for 125 ps, followed by four equilibrations in the NPT ensemble. With each iteration of the NPT equilibrations, constraints on the protein backbone and lipid bilayer were gradually removed until the entire system was allowed to move freely^109^. For the production runs, a time step of 2 fs was used with the md integrator. Three independent replicas were simulated for 5 µs each. Temperature was kept at 310K using a v-rescale thermostat^104^, while pressure was maintained at 1 bar using a semi-isotropic C-rescale barostat^110^. A Verlet cutoff scheme with a cutoff value of 1.2 nm was used to calculate van der Waals and coulombic interactions. Beyond 1.2 ns, particle mesh Ewald was used to compute longrange interactions. Hydrogen bonds were constrained using the LINCS algorithm^111^.

### Simulation analysis

To identify the residues that interacted the most with the lipid/membrane, GROMACS’ *gmx mindist* was used.

To obtain the fractional occupancy of the residues throughout the simulation duration, *gmx select* was used with a distance cutoff of 0.7 nm for CG-MD simulations.

To determine the lipid entry pathways, the center of mass of the lipid along the aligned trajectories was computed and saved in a separate PDB file using *gmx traj* tool. The percentage of occurrence of each pathway was calculated by dividing the number of replicas in which the lipid entered the domain via the specific pathway over the total number of replicas in which the lipid entered the protein cavity.

An in-house tcl script was used to calculate the solvation number of the lipid, counting the number of water molecules within 5.0 Å of the lipid tail beads (headgroup and backbone were excluded) for each frame of the trajectory, for each replica^68^.

To determine the radius of gyration of the ApiTULIP domain at each lipid iteration, *gmx gyrate* was used to calculate the average expansion of the domain throughout the simulation.

To trace lipid delivery from the peripheral domain into the membrane, the z-coordinates of selected lipids were computed using *gmx traj*.

To compute the density of the system components, *gmx density* was used. HeliQuest was used to determine the amphipaticity of residues I1151-H1171^112^.

Lipid translocation events were computed by tracing the angle between each lipid and the z-axis of the membrane along the entire simulation^113^. To decrease the susceptibility to noise coming from incomplete lipid translocations, we defined a buffer region between 50° and 130°, where events were not counted. Additionally, block average with bin size of 50ns was applied. Angles were computed every 1 ns with the GROMACS tool *gmx gangle*. To identify the residues involved with the lipid translocation, we performed protein–lipid contact analysis, for which only the translocated lipids were used. These contacts were computed with the GROMACS tool *gmx select*. A contact was considered when either the headgroup or phosphate bead from the translocated lipid was at a distance of 0.45 nm or lower from any protein bead.

All graphical representations were plotted using Matplotlib^114^. All visual representations were created using VMD^115^.

## Acknowledgements

We thank members of the malaria cell biology laboratory (BNITM, Hamburg) for discussion, Hannah M. Behrens for proofreading the manuscript and J. Matz (BNITM, Hamburg) for discussion and sharing of reagents. We thank P. Haberkant, F. Stein, and the Proteomics Core Facility of the EMBL (Heidelberg, DE) for LC-MS/MS analysis and the BNITM flow cytometry core for experimental support. AGS was supported by an EMBO long-term postdoctoral fellowship (ALTF 166-2022). JD, LSP and TS acknowledge funding from the European Research Council (ERC) under the European Union’s Horizon 2020 research and innovation program (grant agreement no. 101021493). SV acknowledges support from the Swiss National Science Foundation (grant CR00I5-236020 to SV), and by the European Research Council under the European Union’s Horizon 2020 research and innovation program (grant agreement no. 803952). This work was supported by grants from the Swiss National Supercomputing Centre under projects ID s1269 and lp69.

## Author contributions

A.G.S. and T.S. conceptualized the project. A.G.S. designed and performed most experiments and analysed the data. A.R.H. contributed with the DV acidification experiments and the analysis and tagging of DiQ-BioID interacting candidates. K.H. performed the TEM experiments. J.D. developed the genetic complementation assay. L.S.P. provided experimental support. J.C.M., P.C.B, and T.G. contributed data related to PF3D7_1127600 (PfCT3). Y.A., C.R.R. and S.V. performed and analysed the MD simulation experiments. The original draft was written by A.G.S., S.V. and T.S., and reviewed by all the authors.

