## Supplemental Figures for "A new class of lipid transfer proteins is required for the recycling of lipids from the *P. falciparum* digestive vacuole"

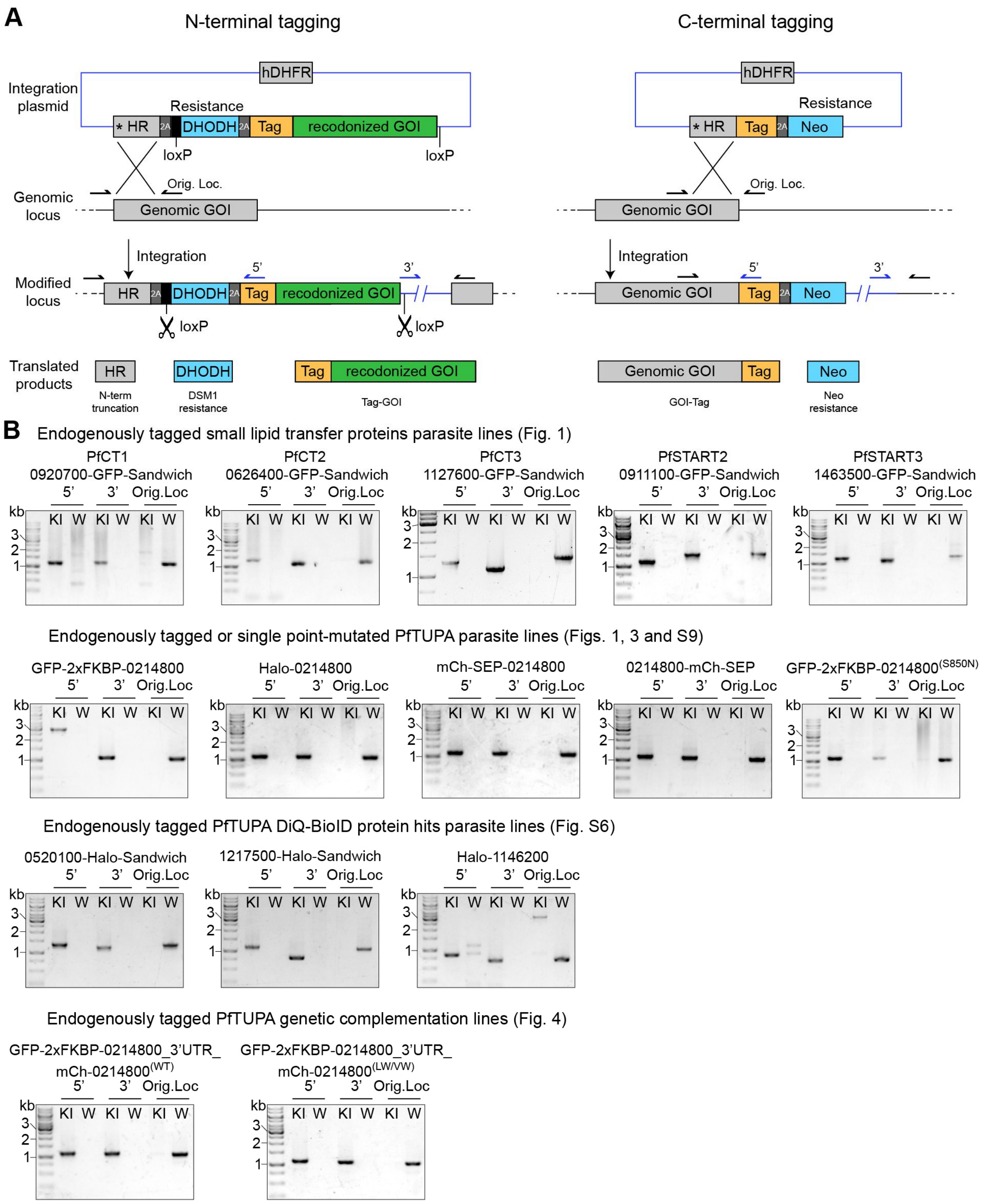


**Supplementary Figure 1. Genomic modifications of *P. falciparum* parasite lines. A**, Schematic illustrating the selection-linked integration (SLI) strategies used to edit the genomic loci of proteins included in this study. The genes for DSM1 (yDHODH) or Neomycin resistance were fused with a skip peptide (T2A) to the gene of interest (for N-terminal or C-terminal modifications, respectively) and were used to select edited parasites. The asterisk indicates a stop codon. Primers used for genomic PCR-based integration evaluation in (**b**) are indicated as arrows. “Tag” varies depending on the cell line, used in this study were 2xFKBP-GFP-2xFKBP (referred to as GFP-Sandwich or GFP-SW), mNeonGreen (mNG), Halo-SW and mCherry-Supereclyptic phluorin (mCh-SEP). **B**, Agarose gels with PCR products amplified from genomic DNA of the indicated parasite lines confirming genomic modifications: the original locus amplicon (Orig. Loc.) shows presence or absence of the unmodified loci whereas 5’ and 3’ are amplicons generated across the integration junctions only present when the parasites have been successfully modified, comparing parental (W) and knock-in (KI) lines. Genomic sequences and expected PCR amplicon sizes are included in Data S1.

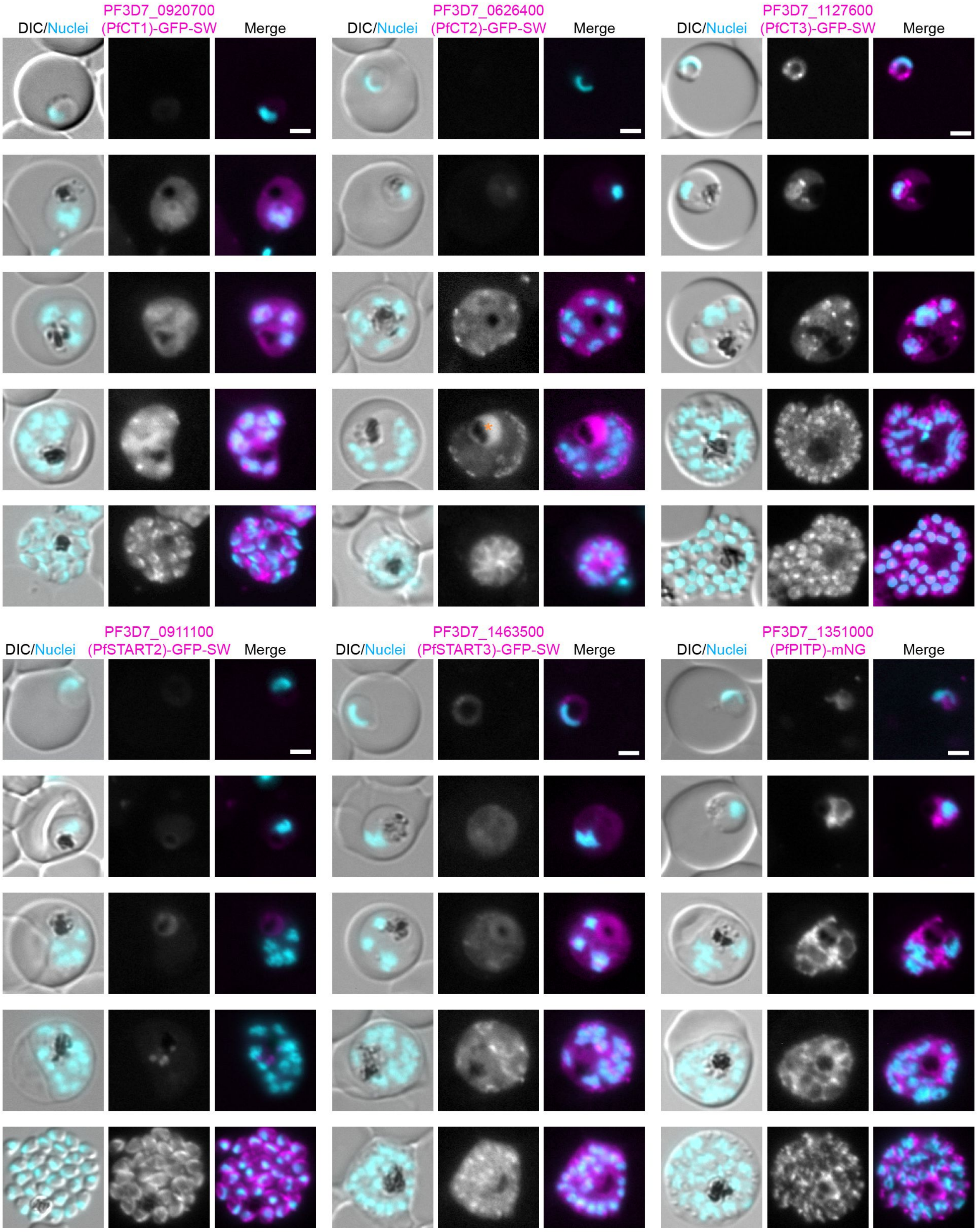


**Supplementary Figure 2. Localization and expression of selected lipid transfer proteins across all asexual RBC stages.** Representative fluorescence microscopy images of the LTPs endogenously tagged as indicated.
DIC, differential interference contrast; Nuclei, Hoechst 33342; scale bars, 2 µm.

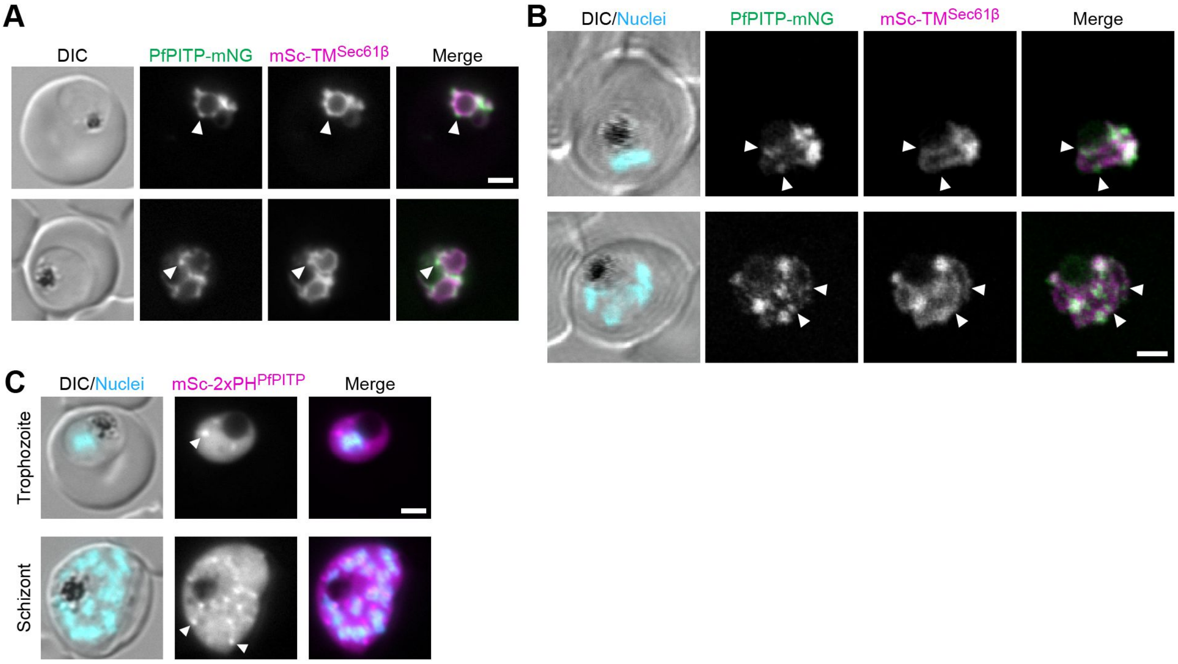


**Supplementary Figure 3: PfPITP is enriched at hospots at the ER. A-B**, Representative fluorescence microscopy images (**A**) and confocal images (**B**) of PfPITP-mNG$^{endo}$ parasites episomally expressing the TM region of Sec61β fused to mSc as an ER marker. **C**, Representative fluorescence microscopy images of two copies of the PH domain of PfPITP arranged in tandem and fused to mSc.

DIC, differential interference contrast; Nuclei, Hoechst 33342; scale bars, 2 µm.

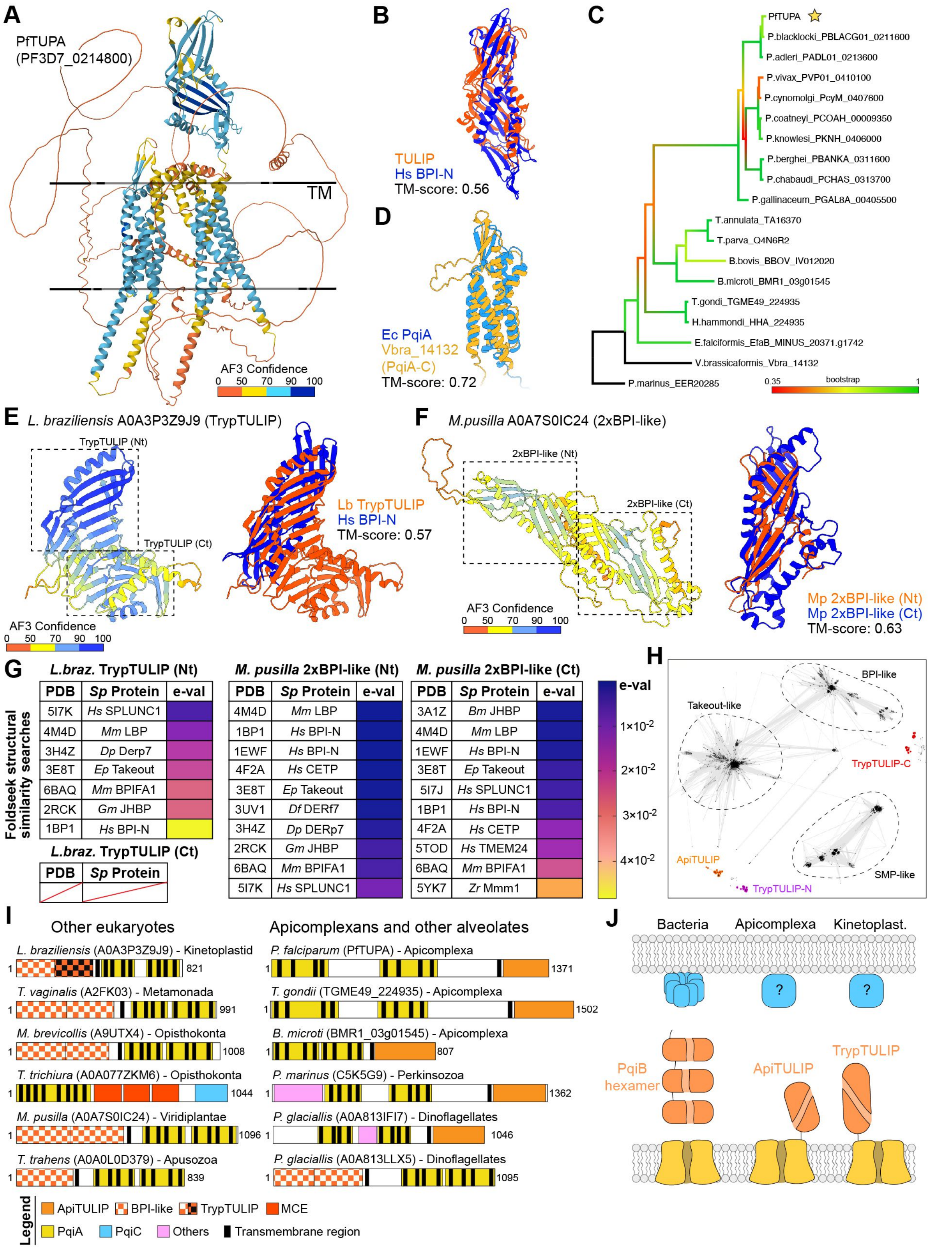


**Supplementary Figure 4. Diversity of TULIP-like lipid transfer domains associated with PqiA transmembrane domains. A**, Ribbon representation of AlphaFold3-predicted structure of

PfTUPA colored by prediction confidence. Black lines represent the edges of the membrane bilayer. **B**, Structural alignment between the predicted structure of the ApiTULIP domain of PfTUPA and the N-terminal BPI domain of the human BPI protein (PDB 1EWF). **C**, Phylogenetic tree of the PfTUPA homologues found in apicomplexan species using BLAST. The protein was not found in the cryptosporidium lineage. **D**, Structural alignment between the predicted structures of the PqiA-C region of the *V. brassicaformis* homologue of PfTUPA and the C-terminal portion of the *E. coli* PqiA protein. **E**, AlphaFold predicted structure of the TrypTULIP domain of the *L. braziliensis* PqiA-containing protein, colored by confidence (left panel) and aligned with the N-terminal BPI domain of the human BPI protein (PDB 1EWF, right panel). **F**, AlphaFold predicted structure of the two BPI-like domains (2xBPI-like) of the *M. pusilla* PqiA-containing protein colored by confidence (left panel) and a structural alignment of the two single domains. **G**, Results of Foldseek structural similarity searches to both lipid transfer domains (N- and C-term) of the TrypTULIP and 2xBPI-like domains of the PqiA-containing proteins found in *L. braziliensis* and *M. pusilla.* The top 10 hits with an e-value below 0.05 were considered. **H**, Cluster analysis of sequences (CLANS) of domains of the TULIP superfamily. Apicomplexan TULIP (ApiTULIP) and N- and C-terminal portions of Trypanosomatid TULIP (TrypTULIP) domains are highlighted in orange, purple and red, respectively. **I**, To scale domain cartoons of PqiA-containing proteins found in a variety of eukaryotic clades (left panel) and alveolates (right panel). The C-terminal ApiTULIP domain is exclusively found in alveolates. **J**, Schematics of PqiA proteins in bacteria, apicomplexans and kinetoplastids.

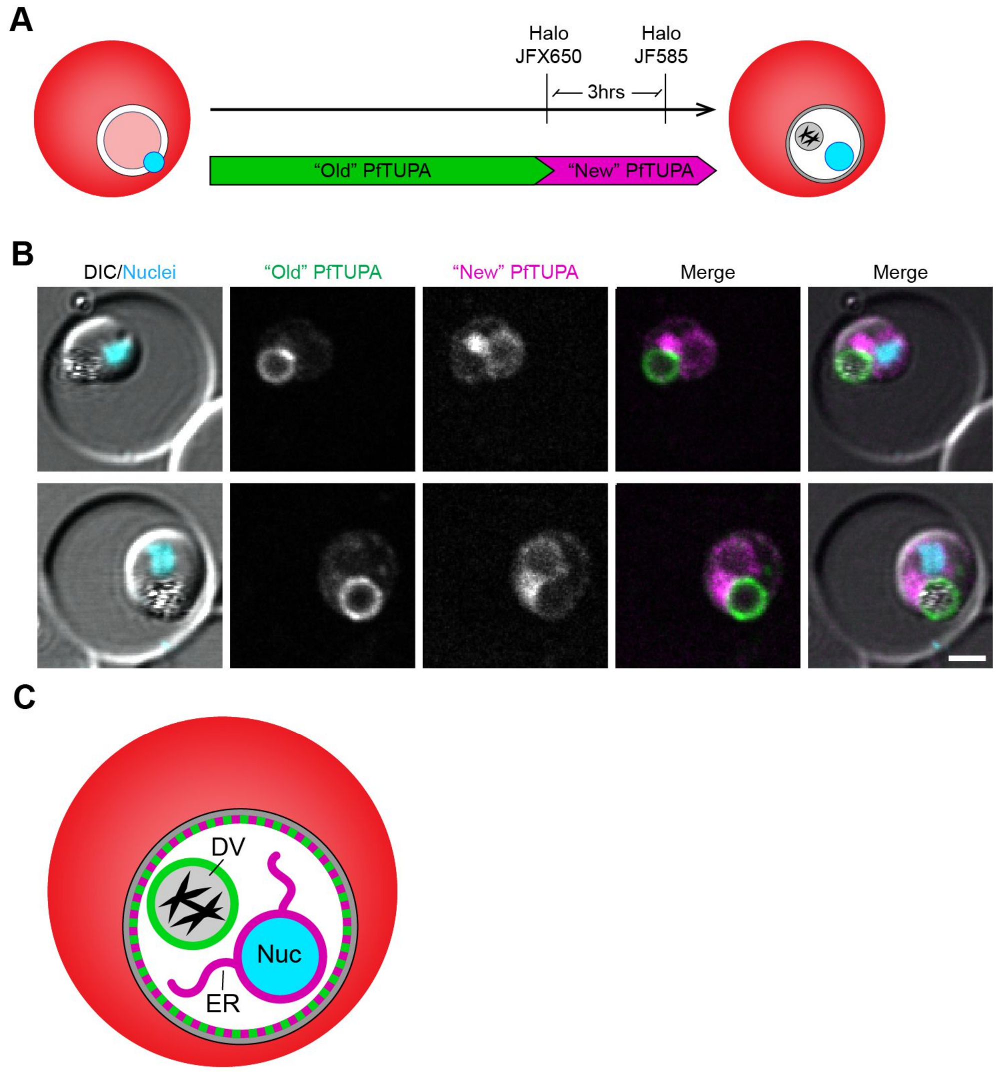


**Supplementary Figure 5. PfTUPA is synthesized in the ER and transported to the DV. A**, Schematic of labelling and microscopy experiment to differentiate pools of Halo-fused PfTUPA protein. "New" protein corresponds to the pool synthesized within the last 3 hours, whereas "Old" refers to the pool synthesized before that. **B**, Representative confocal microscopy images of "New" and "Old" PfTUPA pools of protein in trophozoites. **C**, Schematic of the localization of the "New" (magenta) and "Old" (green) pools of PfTUPA protein.
DIC, differential interference contrast; Nuclei, Hoechst 33342; scale bars, 2 µm.

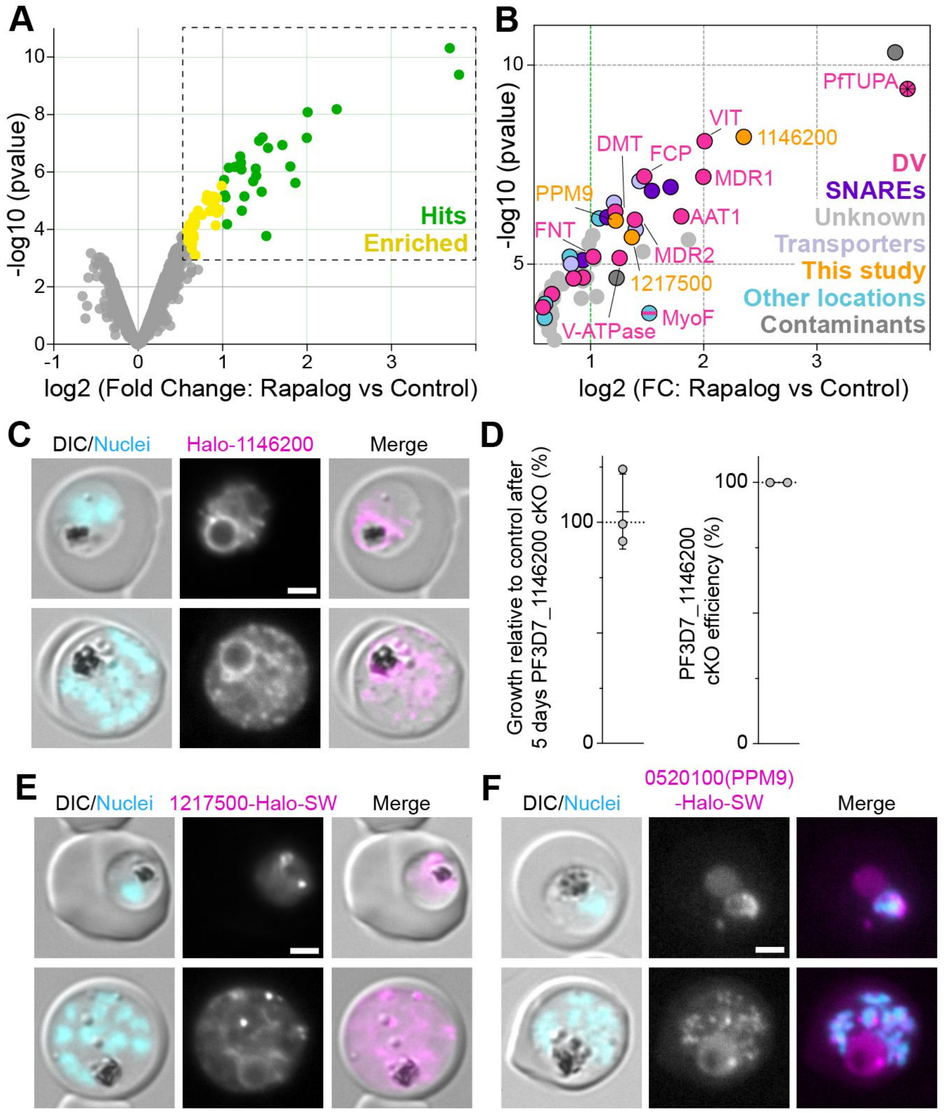


**Supplementary Figure 6. DiQ-BioID of the cytosolic face of PfTUPA identifies a novel DV membrane protein. A**, Volcano plot of the PfTUPA dimerization-induced BioID (DiQ-BioID) (Birnbaum et al., 2020) showing enrichment of proteins in plus rapalog over control. Hits, absolute fold change (FC) of 2 or more and a false discovery rate (FDR) of 0.05 or less (27 proteins); enriched, absolute FC>1.5, FDR <0.2 (35 proteins). Averages from *n* = 3 independent replicates analysed side by side in the same run (data in Table S1). Moderated two-tailed t-test was applied as implemented in the limma package. Dashed box is enlarged in (**B**). **B**, Proteins considered hits or enriched candidates in the PfTUPA DiQ-BioID (same classification as in Fig. 3E) with known DVM hits individually labelled. **C**, Fluorescence microscopy images of trophozoites and schizonts expressing endogenously N-terminal Halo-tagged versions of PF3D7_1146200, unknown protein identified in proximity of PfTUPA and detected at the DV membrane. **D**, Growth relative to control after 5 days of PF3D7_1146200 cKO (left) and quantification of the cKO efficiency (right). Efficiency of cKO was assessed by percentage of cells with expression of PF3D7_1146200 after 1 excision cycle. Growth graph shows mean with SD from n=3 independent

experiments and cKO efficiency graph shows mean with SD from n=2 independent experiments (34 and 42 cells assessed in each experiment). **E-F**, Fluorescence microscopy images of trophozoites and schizonts expressing endogenously C-terminal Halo-tagged versions of two unknown proteins identified in proximity to PfTUPA. PF3D7_1217500 was partially found at the DV membrane, with some foci in proximity to it in trophozoites, whereas PF3D7_0520100 (PPM9) was identified in foci in the nuclear periphery.
DIC, differential interference contrast; Nuclei, Hoechst 33342; scale bars, 2 µm.

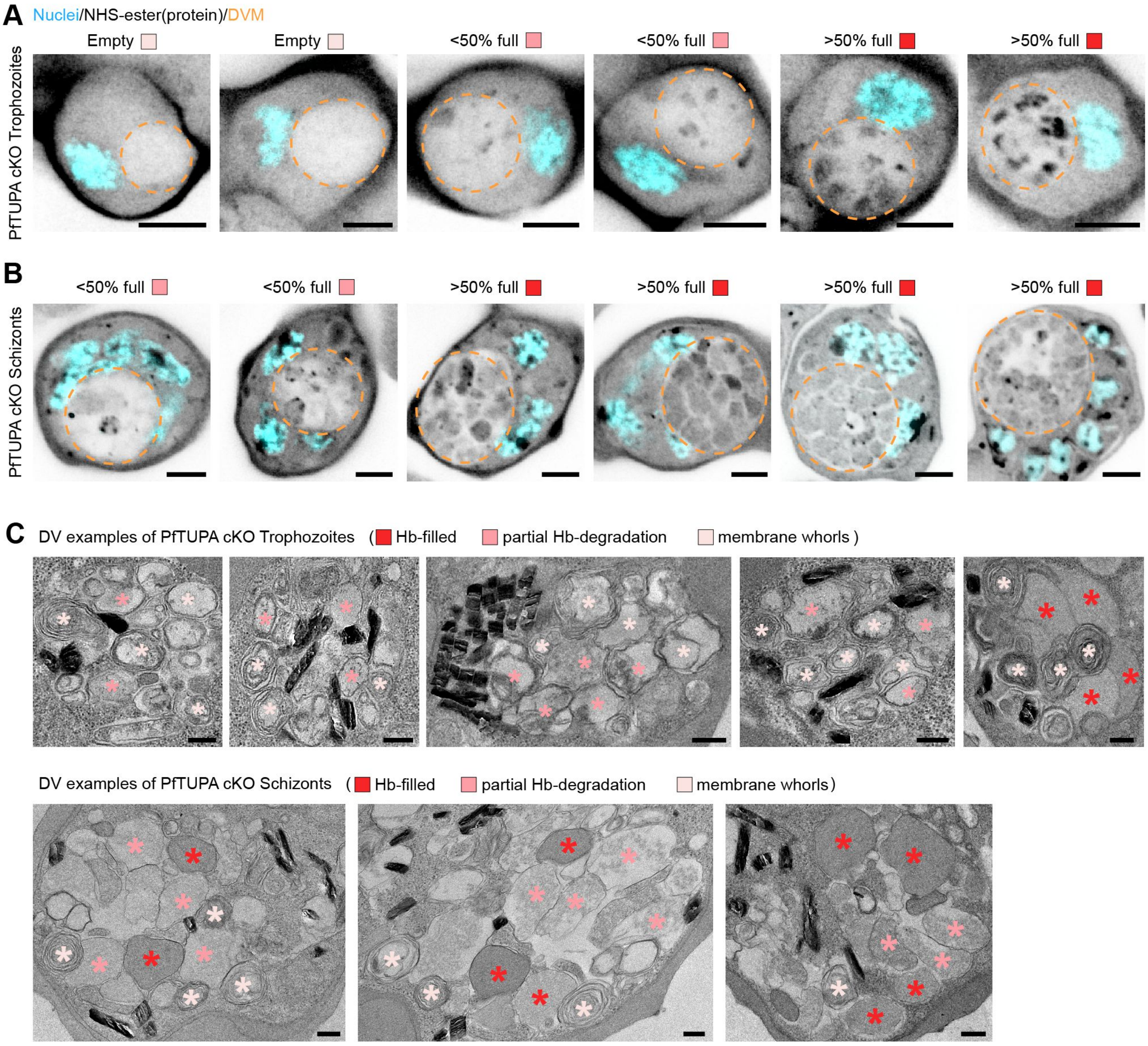


**Supplementary Figure 7. Examples of intraluminal accumulations in PfTUPA cKO**. **A-B**, Ultrastructure Expansion Microscopy (U-ExM) images of PfTUPA cKO parasites in trophozoite (**A**) or schizont (**B**) stages stained with NHS-ester (for labelling proteins). The inferred DV periphery is represented with an orange dashed line. Results were classified depending on the amount of protein content in the DV lumen (as indicated above each image) and a summary of this quantification is shown in Fig. 6C. **C**, Transmission electron microscopy (TEM) of DVs of PfTUPA cKO parasites in trophozoite and schizont stages. The apparent compartments observed in the DV lumen were classified depending on its content: Hb-filled, content of similar staining density to the host cell cytosol; partial Hb-degradation, content with lower staining density; membrane whorls, compartments surrounded by 2 or more membranes. The luminal content classification is summarized in Fig. 6F.

Nuclei, Hoechst 33342; scale bars, 5 µm in **A-B** and 0.25 µm in **C**.

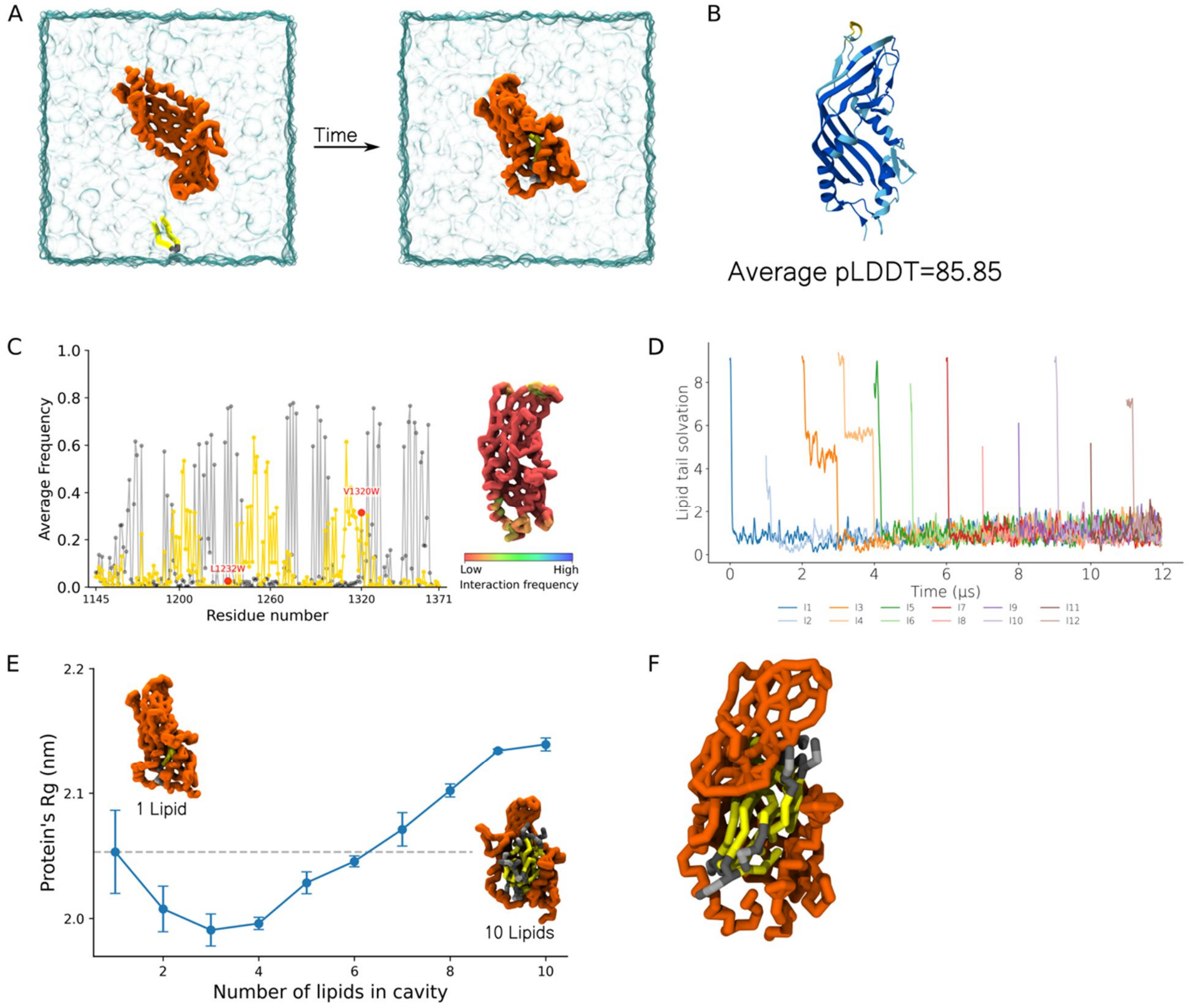


**Supplementary Figure 8. CG-MD simulations of the ApiTULIP domain**. **A**, Snapshots from the CG-MD simulations, showing the initial system setup and the final state where POPC is bound to the ApiTULIP domain. **B**, Alphafold3 prediction of the ApiTULIP alone. **C,** Average frequency of interaction of each residue from the wildtype (black) and the L1232W/V1320W double-mutant (yellow), showing a shift of the per-residue interaction profile from cavity residues to surface residues. **D**, Lipid tail solvation of 12 POPC molecules bound to the ApiTULIP domain of PfTUPA. **E**, Radius of gyration (Rg) of the ApiTULIP domain as a function of the number of POPC molecules inside the cavity. The insets show the domain bound to one POPC molecule (left), and the domain bound to 10 POPC molecules (right). **F**, Final state from CG-MD simulations showing 6 POPC molecules bound to the hydrophobic cavity of the ApiTULIP domain.

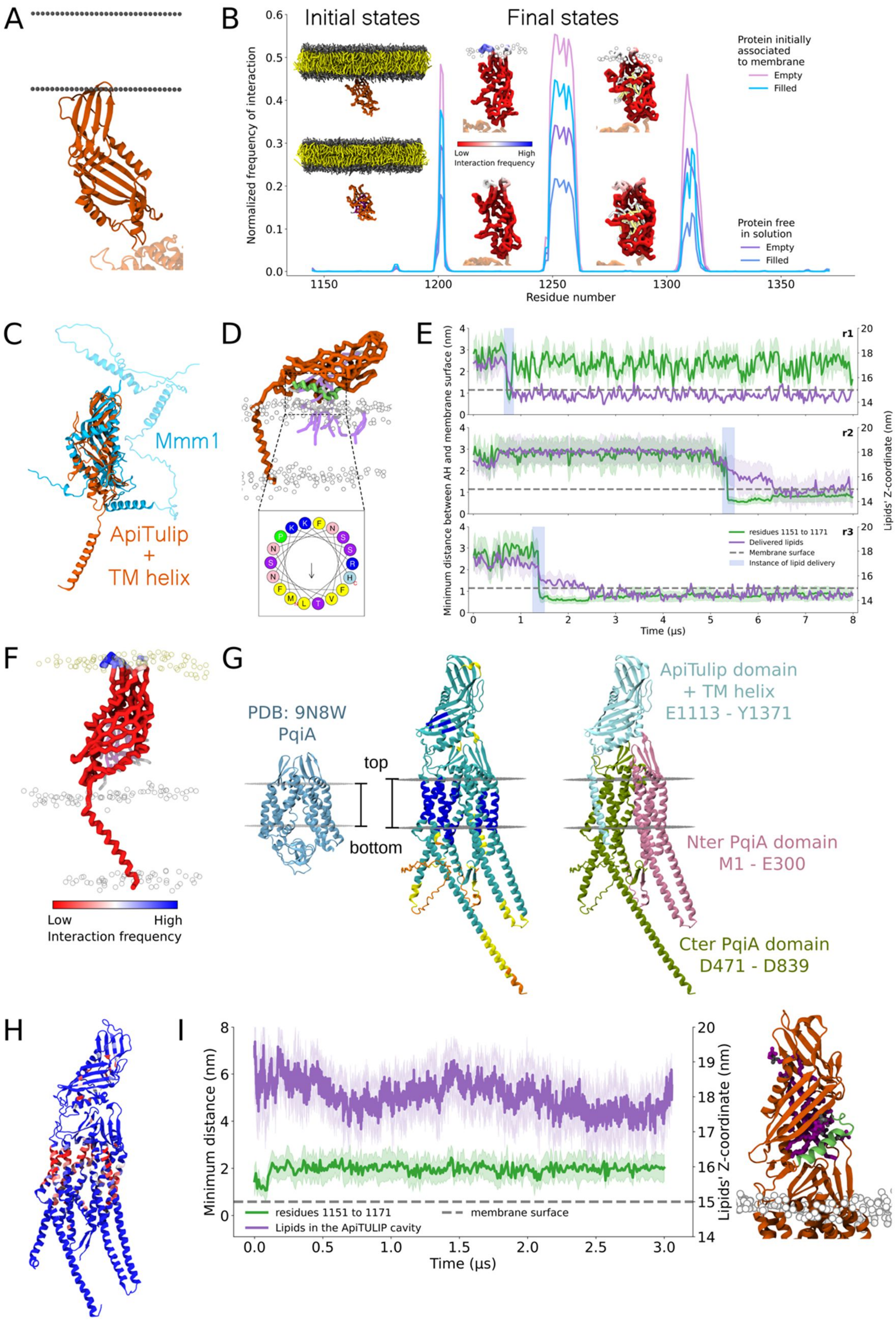


**Supplementary Figure 9. MD simulations of ApiTULIP and PfTUPA. A,** ApiTULIP orientation relative to the membrane as predicted by OPM. **B,** Frequency of interaction of the ApiTULIP domain with the membrane throughout the simulation duration for four different systems where the domain is either lipid-filled or empty and initially associated to the membrane or initially free in solution. The insets on the far left show two representative initial states of the membrane-bound and unbound setups of the ApiTULIP domain. The remaining insets show the binding region (opposite to the TM domains) of the ApiTULIP domain from the four systems. The ApiTULIP domain is colored according to the residue's interaction frequency with the membrane and the rest of the protein, which was not included in the simulations but added here for clarification of the orientation, is shown in faint orange. For the initially membrane-associated systems, phosphate groups are represented by white beads. **C,** Structural alignment of the truncation (residues 1113-1371) of the ApiTULIP domain (residues 1145-1371) with the connected TM helix (residues 1113-1144) (orange) to Mmm1 in yeast (cyan). **D,** Representative snapshot from CG-MD showing the amphipathic helix (green) from the ApiTULIP domain and its respective amphipaticity from HeliQuest. **E,** Time traces from three distinct replicates showing the instance of lipid delivery (light blue highlight) in each replicate. The average z-coordinate of the delivered lipids (purple line, dark traces, right axis) occurred immediately after the amphipathic helix oriented closer to the membrane, indicated by the average minimum distance between the helix (residues 1198-1204) and the membrane's phosphate groups (green line, dark traces, left axis). The light traces indicate the standard deviation across residues/lipids; the membrane surface is indicated by a grey dashed line. **F,** The PfTUPA truncation colored according to the frequency of interaction with the membrane opposite to the TM domains from CG-MD simulations. **G,** Improved structural prediction of PfTUPA (middle), which was obtained by first excluding the unstructured regions then integrating them afterwards using Swissmodel, in comparison to the electron microscopy (EM-) resolved structure of PqiA (left), showing more structural similarity. The illustration on the right indicates the input supplied to Alphafold3 to obtain the improved model. **H,** Full-length PfTUPA from AA-MD simulations colored according to the frequency of interaction with water showing regions interacting with water within the bilayer. **I,** Left, Averaged time traces of the minimum distance between the amphipathic helix and the membrane (green, dark trace, left axis) plotted with the average z-coordinates of the lipids in the cavity (purple, dark trace, right axis) from a representative AA-MD replicate showing no lipid delivery during the simulation time. Right, a representative frame showing the system's details relative to the plot.

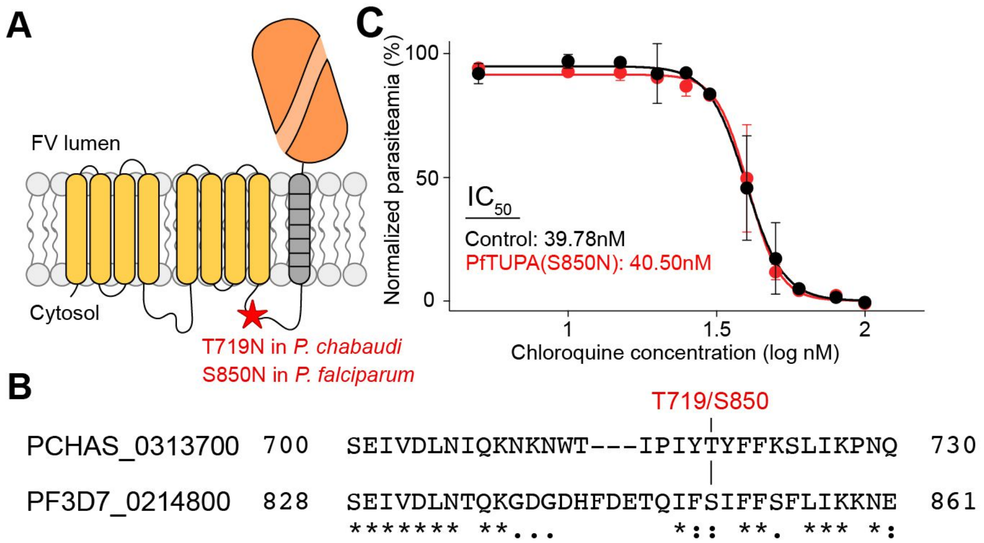


**Supplementary Figure 10. Evaluation of chloroquine resistance in PfTUPA S850N mutant. A-B**, PfTUPA schematic (**A**) and protein sequence alignment (**B**) highlighting the point mutation identified as a CQ resistance conferring mutation in *P. chabaudi* and its corresponding amino acid in *P. falciparum* tested for CQ resistance in (**C**). **C**, CQ $IC_{50}$ curves for PfTUPA(S850N) mutant and WT parental lines.
