## Supplementary material for "A new class of lipid transfer proteins is required for the recycling of lipids from the *P. falciparum* digestive vacuole": Data S1

**Original and modified genomic sequences**

Shown for each gene:

- The unedited genomic sequence.
- The modified genomic sequence after integration.
- The ATG is highlighted in green.
- Introns are in italics.
- The exons are shown in upper case.
- The homology region is shown in bold letters.
- The gene specific primers used for amplification of the original locus are highlighted in yellow.
- The common primers used for amplification of 5’ and 3’ junctions are highlighted in cyan.
- Other gene specific color labels are indicated for each sequence.
- Predicted size of PCR products are at the end of each genomic sequence.

**Index:**

- **PF3D7_0920700 (PfCT1)**	**02-05**
- **PF3D7_0626400 (PfCT2)**	**06-10**
- **PF3D7_1127600 (PfCT3)**	**11-15**
- **PF3D7_0911100 (PfSTART2)**	**16-21**
- **PF3D7_1463500 (PfSTART3)**	**22-25**
- **PF3D7_0214800 (PfTUPA)**	**26-53**
- **PF3D7_1146200**	**54-57**
- **PF3D7_1217500**	**58-61**
- **PF3D7_0520100** **(PPM9)	62-66**

### > PF3D7_0920700 (PfCT1)

ctatgatttaaaaaacaaatagaccatatgatgatccatataatatatatgtcatttcatccatatagtttttatttaaactttgtcaaattataatatatatatatatatatatatatatatatattaatattaatattaatattcttaatgaaaatgtataaaaaattgaaaaattaattctgttcatataagtattattatatatttaatatatatatgatatattatttattttttctcatgttttgtaaaataaatttttctttttttatattttttaaatggaaacttttatat

tatcctaaatattctataaatagtaatatatgaaagatgttttttttttttttttttttttttatttgttttgttttgttttgttttatttatttattattatttttttttttatttatttttttttttttttttatgttcacatgtgtgtgtgtaaatttataaatattatggaataaaaaaaaaaaaaaaaaaaaaaaaaaaaacaaatttaactgctatattatatatattatgtattttaatttatatccatttattccaaaattgagatgttaaaaaaaaagtaaaaataaaaataaataaatatatatataatataatatagcacatatatatacatatatatattaatatgtatatatttatttttaccatttatttaatttattttttcatatttaacttttttttttttttttttttttctttctttttataaaatcatttaacttctatacgggctgaagtagaggcagaacaaatttggtaaaaaaagaagaaaataaaataaaatgaagtcaaatgaaataaaataaaaaaggaaatatttataatataaataaataaataaacatccaaaaatattatgtactaaatttggatgtattaaaaaattttttgtttcaacataataaatcctatatagtacatatatgcaagtatgtaattataatgctatatatatgtaacgaaatatatatatatatatatatatatatatataattatttatcattcatattatgtatatgttatcaaatatatatttatgattacatatttatgtgtaatgaaatgtattcattattttaatataatcattaaaaattatgtatatatatatatatatatgtatatattcaaattaataaacatttaaaaaacattttataaatt

tgcttcatttttcataaaaaataaataggaatatatatatatatatatttaataattatattatattataataatttaattatcagataaaattatacaagattcaaattttataatttcataatattatccttctttgtgtcttgtttttcccttcatcagtatccaaaaaaaaaaaaaaaaaaaaaaATGAGTGATAATGAAAGCTACATTTCTGCAGAAAATAACAAGAGTGATGATGGGAAAAAATCCAATGTAAACCTTGATAATTTTTTAAATTATGATATAGATTATTATATG

AATGAAAAAAATAAATCAAATTATGTAGGAGTAGATATATTACAACGTATTGATTATTATAATATAAAGAGTGCTAATAATGAGTATATAATGAATGGAAATTTAAGTAATAATAATAATAATAGTCATTATCATCATAATATATTATTATATAATAGAAAAAAAGAACAAGTTGATATATTTTATAAATTAAAAAAAGAATTATATAATATAGCATATACAAATGAAGATAATAAAATAAATAATGTCGTAACAACTAATAATAATATAACGGTTCAAAAAAATTCAAAGAAAACATCA

AGTTTTAATACAAAAATCCTTAAAAAAAATTTTAATTTATTTCATTCGAAAAATAAAGATAAAACTAATAATAAAAAATCATATGATTTGTTAAATAATGATTTATGTATAACCTTAGGAAATAAAATAGAAGATACTTATAAAGATATTAAAAATAGTGTTAATTGGCTAGGAAACTTCTTTTCAAACAATGATAATCAAGAAGAAAAAAAATATGTTAATGTTTATTATGCGGATTTATATTTTTTCAACGATATAACCTTAATACGTTTTCTAGATACATACAATTATAACATGTTGAAAACTTTGAATAAAATAATTAAGTTTATAGTTTGGAGACAAATGCATATACAAGATTTTAATAATAAATG**GGATACCTTGAAATTATCTGAGCAATTAACAAAAGAAAATGATAACAATAATAATGATAATAATGATAACAATAATAATGATAATAATAATGATAATAATAACAATGATGATGATAAAGATAAAGATAATAACTGTAATATTTTTCTTAACCCTTCTAATATTAAAGATACTCTTATAAAGTCAAACATCTTCAGATGTGGATACGATATGTATTCCAGACCCATACTATATGTTAAGATACAAGAAAAGCTAGACATGAGTGAAGACGATCTTTTTCTCATGTTAGTTTATCACGTGGATATGTGTATTAATAGTGTAGACTACAAAAAATGTTATCCAGATGCATTTACATCGGAGAAGGACAACATCAATAATGATAATGATAATAATGATAATAACAACAATAATAATAACAACAATAATAATAACAACAATAATGACAAAAACAATAATGACAAAAACAATAATGAAGACGCAGATAAAAGCAAAAGTAATTATAACAGCAGCAGTGATAGCAACAATAAGAAT**

**TATTGTTATAAACCTAATACAGAAAATATTAAAATAGATGAAACGACCTTACAACTTGTTATTGTTGTGGATTGCCTTAAGTTTGAATTAAATAATATGATTAGCGTGGAATGTATAAAAAAAATGATAAATGTATTTAACGAGTTTTATACGGATGTTTTATTCAGAATATATGTTATTAATGTACCATCCTTTTTTAAGAAGGTTTGGTGTCTTTTCAATATGTTTATTGATAATCATACATTAAATAAAATTATTTTTATTAATAAAAAAAACATTAATTTAATGTATGAACATATT**

**CCTATTCATATTATGGAACACTTAGATAAAAATGCAACAAAATCAGAGCAACAAAAATCAGTCTTTTTTCCTTCTAGTTCTTTATATTACAAGTATGATGAAATGTATTATAAAAAGCTTATGCATTATGTCAATATTTGGGTAGACAAAATGATAAAGACTAATCAC**TAGttattaaatgggatcaataagcaaataaacaaaacaaaataaaataaaataatataatatgaaataatatatgtagaaaagtatagtgggagatatgtgaaaaatgtgtaatgccaaattaaaaatttt

ttttttttttttttttttttttgtgtaaagatgtttcatatttaccagcatgagtaacatatataatatatatatatatatataatatatttatttttatttttttgctacataataccacttggttacctttcatttatgttatgtaaaaatatttattttattttgtgtttatattgctttgttttggttatattaaaaaaattccatatgtgcaataatattttaggagtatttattaatttttattgtttttttttttttttttttttttttttttctcttatttttttttaatat

ttttaatagttaacaattaaatgaatcgtttttattgtgatttattaaatataattatgatttagaataatataaaataaaccctgcattcaatgttgataaatttaatatgtagtgaccatgttggatgttactatatttttcttatatattataaagaaataaaaaaaaaaaaaaaaaattaaaagaaaataa

**Original Locus (OL) amplicon size: 1156bp**

### > PF3D7_0920700-GFP-Sandwich GFP Neo-R

ctatgatttaaaaaacaaatagaccatatgatgatccatataatatatatgtcatttcatccatatagtttttatttaaactttgtcaaattataatatatatatatatatatatatatatatatattaatattaatattaatattcttaatgaaaatgtataaaaaattgaaaaattaattctgttcatataagtattattatatatttaatatatatatgatatattatttattttttctcatgttttgtaaaataaatttttctttttttatattttttaaatggaaacttttatat

tatcctaaatattctataaatagtaatatatgaaagatgttttttttttttttttttttttttatttgttttgttttgttttgttttatttatttattattatttttttttttatttatttttttttttttttttatgttcacatgtgtgtgtgtaaatttataaatattatggaataaaaaaaaaaaaaaaaaaaaaaaaaaaaacaaatttaactgctatattatatatattatgtattttaatttatatccatttattccaaaattgagatgttaaaaaaaaagtaaaaataaaaataaataaatatatatataatataatatagcacatatatatacatatatatattaatatgtatatatttatttttaccatttatttaatttattttttcatatttaacttttttttttttttttttttttctttctttttataaaatcatttaacttctatacgggctgaagtagaggcagaacaaatttggtaaaaaaagaagaaaataaaataaaatgaagtcaaatgaaataaaataaaaaaggaaatatttataatataaataaataaataaacatccaaaaatattatgtactaaatttggatgtattaaaaaattttttgtttcaacataataaatcctatatagtacatatatgcaagtatgtaattataatgctatatatatgtaacgaaatatatatatatatatatatatatatatataattatttatcattcatattatgtatatgttatcaaatatatatttatgattacatatttatgtgtaatgaaatgtattcattattttaatataatcattaaaaattatgtatatatatatatatatatgtatatattcaaattaataaacatttaaaaaacattttataaatt

tgcttcatttttcataaaaaataaataggaatatatatatatatatatttaataattatattatattataataatttaattatcagataaaattatacaagattcaaattttataatttcataatattatccttctttgtgtcttgtttttcccttcatcagtatccaaaaaaaaaaaaaaaaaaaaaaATGAGTGATAATGAAAGCTACATTTCTGCAGAAAATAACAAGAGTGATGATGGGAAAAAATCCAATGTAAACCTTGATAATTTTTTAAATTATGATATAGATTATTATATG

AATGAAAAAAATAAATCAAATTATGTAGGAGTAGATATATTACAACGTATTGATTATTATAATATAAAGAGTGCTAATAATGAGTATATAATGAATGGAAATTTAAGTAATAATAATAATAATAGTCATTATCATCATAATATATTATTATATAATAGAAAAAAAGAACAAGTTGATATATTTTATAAATTAAAAAAAGAATTATATAATATAGCATATACAAATGAAGATAATAAAATAAATAATGTCGTAACAACTAATAATAATATAACGGTTCAAAAAAATTCAAAGAAAACATCA

AGTTTTAATACAAAAATCCTTAAAAAAAATTTTAATTTATTTCATTCGAAAAATAAAGATAAAACTAATAATAAAAAATCATATGATTTGTTAAATAATGATTTATGTATAACCTTAGGAAATAAAATAGAAGATACTTATAAAGATATTAAAAATAGTGTTAATTGGCTAGGAAACTTCTTTTCAAACAATGATAATCAAGAAGAAAAAAAATATGTTAATGTTTATTATGCGGATTTATATTTTTTCAACGATATAACCTTAATACGTTTTCTAGATACATACAATTATAACATGTTGAAAACTTTGAATAAAATAATTAAGTTTATAGTTTGGAGACAAATGCATATACAAGATTTTAATAATAAATG**GGATACCTTGAAATTATCTGAGCAATTAACAAAAGAAAATGATAACAATAATAATGATAATAATGATAACAATAATAATGATAATAATAATGATAATAATAACAATGATGATGATAAAGATAAAGATAATAACTGTAATATTTTTCTTAACCCTTCTAATATTAAAGATACTCTTATAAAGTCAAACATCTTCAGATGTGGATACGATATGTATTCCAGACCCATACTATATGTTAAGATACAAGAAAAGCTAGACATGAGTGAAGACGATCTTTTTCTCATGTTAGTTTATCACGTGGATATGTGTATTAATAGTGTAGACTACAAAAAATGTTATCCAGATGCATTTACATCGGAGAAGGACAACATCAATAATGATAATGATAATAATGATAATAACAACAATAATAATAACAACAATAATAATAACAACAATAATGACAAAAACAATAATGACAAAAACAATAATGAAGACGCAGATAAAAGCAAAAGTAATTATAACAGCAGCAGTGATAGCAACAATAAGAATTATTGTTATAAACCTAATACAGAAAATATTAAAATAGATGAAACGACCTTACAACTTGTTATTGTTGTGGATTGCCTTAAGTTTGAATTAAATAATATGATTAGCGTGGAATGTATAAAAAAAATGATAAATGTATTTAACGAGTTTTATACGGATGTTTTATTCAGAATATATGTTATTAATGTACCATCCTTTTTTAAGAAGGTTTGGTGTCTTTTCAATATGTTTATTGATAATCATACATTAAATAAAATTATTTTTATTAATAAAAAAAACATTAATTTAATGTATGAACATATTCCTATTCATATTATGGAACACTTAGATAAAAATGCAACAAAATCAGAGCAACAAAAATCAGTCTTTTTTCCTTCTAGTTCTTTATATTACAAGTATGATGAAATGTATTATAAAAAGCTTATGCATTATGTCAATATTTGGGTAGACAAAATGATAAAGACTAATCAC**CCTAGGTCAGGATTGAGATCAAGATCTGCTGCTGCTGGTGCTGGTGGTGCTGCTAGAGCTGCTCTGCAGAGAGGAGTACAAGTTGAAACAATATCACCAGGAGATGGTCGTACATTTCCAAAAAGAGGTCAAACTTGTGTTGTACATTATACTGGAATGCTTGAAGATGGAAAGAAATTTGATTCATCTCGTGATAGAAATAAACCATTTAAATTTATGCTAGGTAAACAAGAAGTAATACGAGGTTGGGAAGAAGGAGTTGCTCAAATGAGTGTAGGTCAAAGAGCAAAACTTACTATATCTCCAGATTATGCTTATGGTGCAACTGGACATCCAGGTATAATTCCACCTCATGCAACTCTTGTATTTGATGTGGAGCTTCTAAAACTAGAAACTAGAGGTGTTCAGGTTGAAACAATTTCACCTGGAGATGGCAGAACCTTTCCTAAAAGAGGACAGACTTGCGTAGTTCATTATACAGGCATGCTAGAGGATGGTAAGAAATTTGATTCTAGTCGAGATAGAAATAAGCCATTCAAGTTTATGCTAGGTAAACAGGAAGTAATAAGAGGTTGGGAAGAGGGTGTAGCACAGATGTCAGTTGGACAAAGAGCAAAGTTAACAATATCACCAGATTATGCATACGGTGCAACAGGCCATCCTGGCATCATCCCTCCACATGCAACTTTAGTATTCGACGTTGAATTGTTAAAGTTAGAGACAACGCGTGCTAGAGGTGCTGCTGCTGGTGCTGGAGGTGCAGGTAGACGTACGATGAGTAAAGGAGAAGAACTTTTCACTGGAGTTGTCCCAATTCTTGTTGAATTAGATGGTGATGTTAATGGGCACAAATTTTCTGTCAGTGGAGAGGGTGAAGGTGATGCAACATACGGAAAACTTACCCTTAAATTTATTTGCACTACTGGAAAACTACCTGTTCCATGGCCAACACTTGTCACTACTTTCGCGTATGGTCTTCAATGCTTTGCGAGATACCCAGATCATATGAAACAGCATGACTTTTTCAAGAGTGCCATGCCCGAAGGTTATGTACAGGAAAGAACTATATTTTTCAAAGATGACGGGAACTACAAGACACGTGCTGAAGTCAAGTTTGAAGGTGATACCCTTGTTAATAGAATCGAGTTAAAAGGTATTGATTTTAAAGAAGATGGAAACATTCTTGGACACAAATTGGAATACAACTATAACTCACACAATGTATACATCATGGCAGACAAACAAAAGAATGGAATCAAAGTTAACTTCAAAATTAGACACAACATTGAAGATGGAAGCGTTCAACTAGCAGACCATTATCAACAAAATACTCCAATTGGCGATGGCCCTGTCCTTTTACCAGACAACCATTACCTGTCCACACAATCTGCCCTTTCGAAAGATCCCAACGAAAAGAGAGACCACATGGTCCTTCTTGAGTTTGTAACAGCTGCTGGGATTACACATGGCATGGATGAGCTCTACAAAGTCGACGCCAGGGGAGCAGCCGCAGGAGCAGGGGGGGCAGGAAGGCGTGGTGTTCAGGTCGAGACTATTAGCCCTGGAGATGGACGCACGTTTCCTAAGCGTGGACAGACATGCGTAGTTCACTACACAGGTATGTTGGAGGACGGTAAAAAGTTCGACAGCTCACGCGACCGCAATAAACCTTTCAAGTTTATGCTTGGCAAGCAGGAGGTTATTCGTGGATGGGAGGAGGGTGTAGCACAGATGTCTGTTGGACAGCGTGCTAAGTTGACAATTTCACCTGACTATGCTTATGGCGCTACGGGCCATCCCGGGATCATTCCGCCACATGCGACTCTGGTATTCGACGTTGAATTATTAAAGTTAGAGACAGCTAGAGGGGCCGCTGCAGGTGCTGGTGGAGCTGGAAGACGTGGAGTACAAGTAGAGACTATCTCTCCAGGTGACGGTCGCACTTTCCCAAAGCGTGGCCAAACCTGTGTTGTACATTACACTGGTATGCTGGAGGATGGGAAAAAGTTCGATTCCAGTCGCGACCGTAACAAACCGTTCAAATTCATGTTGGGAAAGCAGGAAGTGATCCGCGGGTGGGAGGAAGGCGTGGCGCAAATGAGCGTCGGTCAGCGGGCTAAATTGACCATTTCCCCTGACTACGCGTATGGGGCTACTGGGCACCCAGGGATTATTCCGCCTCACGCTACACTTGTGTTTGATGTCGAACTTTTGAAACTGGAAACTGTCGACGGAGAAGGAAGAGGAAGTTTATTAACATGTGGAGATGTAGAAGAAAATCCAGGACCAATGATTGAACAAGATGGATTGCACGCAGGTTCTCCGGCCGCTTGGGTGGAGAGGCTATTCGGCTATGACTGGGCACAACAGACAATCGGCTGCTCTGATGCCGCCGTGTTCCGGCTGTCAGCGCAGGGGCGCCCGGTTCTTTTTGTCAAGACCGACCTGTCCGGTGCCCTGAATGAACTGCAGGACGAGGCAGCGCGGCTATCGTGGCTGGCCACGACGGGCGTTCCTTGCGCAGCTGTGCTCGACGTTGTCACTGAAGCGGGAAGGGACTGGCTGCTATTGGGCGAAGTGCCGGGGCAGGATCTCCTGTCATCTCACCTTGCTCCTGCCGAGAAAGTATCCATCATGGCTGATGCAATGCGGCGGCTGCATACGCTTGATCCGGCTACCTGCCCATTCGACCACCAAGCGAAACATCGCATCGAGCGAGCACGTACTCGGATGGAAGCCGGTCTTGTCGATCAGGATGATCTGGACGAAGAGCATCAGGGGCTCGCGCCAGCCGAACTGTTCGCCAGGCTCAAGGCGCGCATGCCCGACGGCGAGGATCTCGTCGTGACCCATGGCGATGCCTGCTTGCCGAATATCATGGTGGAAAATGGCCGCTTTTCTGGATTCATCGACTGTGGCCGGCTGGGTGTGGCGGACCGCTATCAGGACATAGCGTTGGCTACCCGTGATATTGCTGAAGAGCTTGGCGGCGAATGGGCTGACCGCTTCCTCGTGCTTTACGGTATCGCCGCTCCCGATTCGCAGCGCATCGCCTTCTATCGCCTTCTTGACGAGTTCTTCTAActcgagggatatggcagcttaatgttcgtttttcttatttatatatttataccaattgattgtatttataactgtaaaaatgtgtatgttgtgtgcatatttttttttgtgcatgcacatgcatgtaaatagctaaaattatgaacattttattttttgttcagaaaaaaaaaactttacacacataaaatggctagtatgaatagccatattttatataaattaaatcctatgaatttatgaccatattaaaaatttagatatttatggaacataatatgtttgaaacaataagacaaaattattattattattattatttttactgttataattatgtgtctccttcaatgattcataaatagttggacttgatttttaaaatgtttataatatgattagcatagttaaataaaaaaagttgaaaaattaaaaaaaaacatataaacacaaatgatggtttttccttcaatttcgatatcaatttatagaaacaaaatatatacttgtataattttatttttttatataaatcattacatatataattatacaatattttttctaagagataattatatattaatatatataaaaaaaggtgttttttttttttttttttatttttatttttattttatggtaatattttattttccttattttataaattatattagtttatatgtgattaattttatatattatcaatttatatatttttaaatgcttacttaattatctttttttttttttttttttttttttcccctctttttatattaatttatttttgaaaaaattgatatatatatatatatataatatatatatatacatgtagtagtattaaacaatgtataatatatataaataatatatttatatatttcatttcaattttaattttttttggttttttttttttttctttttgtcatatttaaaaaaaattatattcatataagttatgcattttttataaacattattcaatatatgtataatataatatatatatatatattaatgtattattccaatgtgcatgataaaagaaaaaaataatatttataaaaaaaaagaaaaataaaacaaaaaaagaaaaaaaaaaaaaaaaaaaaaaaaatacaaaaataaataatataatttataattatatattcttgtcacaataaaaatatatatatatatatatatatttataatatgtatattttaaactagaaaaggaataactaatattttatttattatcattcaagatttatattttataataataaatacctaatagaaatatatcaggatccatgcatggttcgctaaactgcatcgtcgctgtgtcccagaacatgggcatcggcaagaacggggactacccctggccaccgctcaggaacgaatttagatatttccagagaatgaccacaacctcttcagtagaaggtaaacagaatctggtgattatgggtaagaagacctggttctccattcctgagaagaatcgacctttaaagggtagaattaatttagttctcagcagagaactcaaggaacctccacaaggagctcattttctttccagaagtctagatgatgccttaaaacttactgaacaaccagaattagcaaataaagtagacatggtctggatagttggtggcagttctgtttataaggaagccatgaatcacccaggccatcttaaactatttgtgacaaggatcatgcaagactttgaaagtgacacgttttttccagaaattgatttggagaaatataaacttctgccagaatacccaggtgttctctctgatgtccaggaggagaaaggcattaagtacaaatttgaagtatatgagaagaatgattaagcttatttaataatagattaaaaatattataaaaataaaaacataaacacagaaattacaaaaaaaatacatatgaattttttttttgtaatcttccttataaatatagaataatgaatcatataaaacatatcattattcatttatttacatttaaaattattgtttcagtatctttaatttattatgtatatataaaaataacttacaattttattaataaacaatatatgtttattaattcatgttttgtaatttatgggatagcgattttttttactgtctgtatttttcttttttaattatgttttaattgtattttatttttattattgttctttttatagtattattttaaaacaaaatgtattttctaagaacttataataataataaatataaattttaataaaaattatatttatcttttacaatatgaacataaagtacaacattaatatatagcttttaatatttttattcctaatcatgtaaatcttaaatttttctttttaaacatatgttaaatatttatttctcattatatataagaacatatttattaaatctagaattctatagtgagtcgtattacaattcactggccgtcgttttacaacgtcgtgactgggaaaaccctggcgttacccaacttaatcgccttgcagcacatccccctttcgccagctggcgtaatagcgaagaggcccgcaccgatcgcccttcccaacagttgcgcagcctgaatggcgaatggcgcctgatgcggtattttctccttacgcatctgtgcggtatttcacaccgcatatggtgcactctcagtacaatctgctctgatgccgcatagttaagccagccccgacacccgccaacacccgctgacgcgccctgacgggcttgtctgctcccggcatccgcttacagacaagctgtgaccgtctccgggagctgcatgtgtcagaggttttcaccgtcatcaccgaaacgcgcgagacgaaagggcctcgtgatacgcctatttttataggttaatgtcatgataataatggtttcttagacgtcaggtggcacttttcggggaaatgtgcgcggaacccctatttgtttatttttctaaatacattcaaatatgtatccgctcatgagacaataaccctgataaatgcttcaataatattgaaaaaggaagagtatgagtattcaacatttccgtgtcgcccttattcccttttttgcggcattttgccttcctgtttttgctcacccagaaacgctggtgaaagtaaaagatgctgaagatcagttgggtgcacgagtgggttacatcgaactggatctcaacagcggtaagatccttgagagttttcgccccgaagaacgttttccaatgatgagcacttttaaagttctgctatgtggcgcggtattatcccgtattgacgccgggcaagagcaactcggtcgccgcatacactattctcagaatgacttggttgagtactcaccagtcacagaaaagcatcttacggatggcatgacagtaagagaattatgcagtgctgccataaccatgagtgataacactgcggccaacttacttctgacaacgatcggaggaccgaaggagctaaccgcttttttgcacaacatgggggatcatgtaactcgccttgatcgttgggaaccggagctgaatgaagccataccaaacgacgagcgtgacaccacgatgcctgtagcaatgccaacaacgttgcgcaaactattaactggcgaactacttactctagcttcccggcaacaattaatagactggatggaggcggataaagttgcaggaccacttctgcgctcggcccttccggctggctggtttattgctgataaatctggagccggtgagcgtgggtctcgcggtatcattgcagcactggggccagatggtaagccctcccgtatcgtagttatctacacgacggggagtcaggcaactatggatgaacgaaatagacagatcgctgagataggtgcctcactgattaagcattggtaactgtcagaccaagtttactcatatatactttagattgatttaaaacttcatttttaatttaaaaggatctaggtgaagatcctttttgataatctcatgaccaaaatcccttaacgtgagttttcgttccactgagcgtcagaccccgtagaaaagatcaaaggatcttcttgagatcctttttttctgcgcgtaatctgctgcttgcaaacaaaaaaaccaccgctaccagcggtggtttgtttgccggatcaagagctaccaactctttttccgaaggtaactggcttcagcagagcgcagataccaaatactgtccttctagtgtagccgtagttaggccaccacttcaagaactctgtagcaccgcctacatacctcgctctgctaatcctgttaccagtggctgctgccagtggcgataagtcgtgtcttaccgggttggactcaagacgatagttaccggataaggcgcagcggtcgggctgaacggggggttcgtgcacacagcccagcttggagcgaacgacctacaccgaactgagatacctacagcgtgagctatgagaaagcgccacgcttcccgaagggagaaaggcggacaggtatccggtaagcggcagggtcggaacaggagagcgcacgagggagcttccagggggaaacgcctggtatctttatagtcctgtcgggtttcgccacctctgacttgagcgtcgatttttgtgatgctcgtcaggggggcggagcctatcgaaaaacgccagcaacgcggcctttttacggttcctggccttttgctggccttttgctcacatgttctttcctgcgttatcccctgattctgtggataaccgtattaccgcctttgagtgagctgataccgctcgccgcagccgaacgaccgagcgcagcgagtcagtgagcgaggaagcggaagagcgcccaatacgcaaaccgcctctccccgcgcgttggccgattcattaatgcagctggcacgacaggtttcccgactggaaagcgggcagtgagcgcaacgcaattaatgtgagttagctcactcattaggcaccccaggctttacactttatgcttccggctcgtatgttgtgtggaattgtgagcggataacaatttcacacaggaaacagctatgaccatgattacgccaagctatttaggtgacactatagaatactcgcggccgcTAGGATACCTTGAAATTATCTGAGCAATTAACAAAAGAAAATGATAACAATAATAATGATAATAATGATAACAATAATAATGATAATAATAATGATAATAATAACAATGATGATGATAAAGATAAAGATAATAACTGTAATATTTTTCTTAACCCTTCTAATATTAAAGATACTCTTATAAAGTCAAACATCTTCAGATGTGGATACGATATGTATTCCAGACCCATACTATATGTTAAGATACAAGAAAAGCTAGACATGAGTGAAGACGATCTTTTTCTCATGTTAGTTTATCACGTGGATATGTGTATTAATAGTGTAGACTACAAAAAATGTTATCCAGATGCATTTACATCGGAGAAGGACAACATCAATAATGATAATGATAATAATGATAATAACAACAATAATAATAACAACAATAATAATAACAACAATAATGACAAAAACAATAATGACAAAAACAATAATGAAGACGCAGATAAAAGCAAAAGTAATTATAACAGCAGCAGTGATAGCAACAATAAGAATTATTGTTATAAACCTAATACAGAAAATATTAAAATAGATGAAACGACCTTACAACTTGTTATTGTTGTGGATTGCCTTAAGTTTGAATTAAATAATATGATTAGCGTGGAATGTATAAAAAAAATGATAAATGTATTTAACGAGTTTTATACGGATGTTTTATTCAGAATATATGTTATTAATGTACCATCCTTTTTTAAGAAGGTTTGGTGTCTTTTCAATATGTTTATTGATAATCATACATTAAATAAAATTATTTTTATTAATAAAAAAAACATTAATTTAATGTATGAACATATTCCTATTCATATTATGGAACACTTAGATAAAAATGCAACAAAATCAGAGCAACAAAAATCAGTCTTTTTTCCTTCTAGTTCTTTATATTACAAGTATGATGAAATGTATTATAAAAAGCTTATGCATTATGTCAATATTTGGGTAGACAAAATGATAAAGACTAATCACTAGttattaaatgggatcaataagcaaataaacaaaacaaaataaaataaaataatataatatgaaataatatatgtagaaaagtatagtgggagatatgtgaaaaatgtgtaatgccaaattaaaaattttttttttttttttttttttttttgtgtaaagatgtttcatatttaccagcatgagtaacatatataatatatatatatatatataatatatttatttttatttttttgctacataataccacttggttacctttcatttatgttatgtaaaaatatttattttattttgtgtttatattgctttgttttggttatattaaaaaaattccatatgtgcaataatattttaggagtatttattaatttttattgtttttttttttttttttttttttttttttctcttatttttttttaatatttttaatagttaacaattaaatgaatcgtttttattgtgatttattaaatataattatgatttagaataatataaaataaaccctgcattcaatgttgataaatttaatatgtagtgaccatgttggatgttactatatttttcttatatattataaagaaataaaaaaaaaaaaaaaaaattaaaagaaaataa

**5’ amplicon size: 1177bps**

**3’ amplicon size: 1205bps**

### > PF3D7_0626400 (PfCT2)

tacatatatacacattcattgtattttttttttttttttttttttttttttcacttccatttatattttaaacattatttttttattcttttattaatatacaaaagaaaaaaacatttgaaggacatgtttaaaaaataaatatatacacatatttatatataattatatatgatgtggaatttttaatgttcagtgtaacattttataagaagaataggagtaatatgaaaataataataacgttcaaatagaggaatataataaaataaaaacaataaaaaagaaaataatgaaaatccatacatatatttgaaaaagaaacatacatatgtatatatatatatatatatataaatgtacattgtgtgttgtaagaattataaacattatttatatttacacttttatttaattatttatttatatttatatttacttatttatttacatttacatttataattttaattttttttttttttttttttttgtgtacaaatgtatgaatttataaaaaaattaaaaaaaaaaaaaaaaataaggttgcatactgataatatgaaatatatttatcatttggtatgaatattatattaatagaatatatatatatatatatatatatatatatatgtatatattaaaggggcatatatatgtacatattatatattttgattatataagaaatatatataagctacatatataaaataatagctatccatttatacttgtatggacaaaataagaaaggaagaaaaaaaagaaaagaaaagaaaagtATGGAACTTTCAAAAGGAAAAGTTATTATTGAAAAAGATATAAATGAACACACCATAGATGACGAAGTGTTTATGTTTGAACCTAGCATTGATGATGTGTATGATAAGAATACGAATTTACGGTTTATATTTCATAATACATTTATAACATCAGAAGAAGAAATAGCTATAAGTGAATTTAGAAAATATTGTAAGAGTAGATGCTTAAAAATTAATAAAATATATTTCGAAAATGAATGCTTACGTTATTTATATTCAGCACAATTTGATTTTTCTAAGGCCATGGAATTAATAAAAAGTAATTATGAATTTCGGTTATCATCTATTTTACCAATAAAAGAAAAAGATGTGATATTTTATATAAATAAAGGAGTTATGTATTGGCATGGGAGAGATAAAAAATGTAGACCAATATTAATTATTAATTTATTAAAAGTTGAATTATTAAGTATTGATGATTTATCTAATTTATTTTTTTTCTGTTTTGAATTTTTTTTAAAATATTTATGTATCCCAGGAAAAATAGAAAACTATATATCAATAATCGATTGTTCTGGTATATCTATATCAAAATTCCCTATGACAACATTTATGAAATTGTTAGAAATTATGAATTCTAAATATAGATGTAGATTATTTAGAATGTATATTCTTGAGGCACCTAAAATTTTAAAAACATTTGGAAAATCATTTCTAAATTTTGCACCTACCTATATGACCAAAAAATTAAAAATATTAGATAACAATTATGCAGATTATTTAAGAGAAGAAATATTATCAACTCAATTAGAAAAAAAATATGGAGGAATTCAGGAAGATAAAATTAATAATTTTTATCCTTTCCACTTTTATCCAAGTTGTTATATATCACAACAACAAAAAAGGAAATCTCAAGATAATAAAGCAGTTATCGAAAAAAAAACAAATATTTTTAATAATGATCATATTTATAATATATTCCTATCTGGTTATTCTATGCATGTAATATTAGTCCAGAAGGATCAAAGGATAATTGATAATCATCGGTATAATGATATTCTATTAAATGATTCAAATTCTGCAAATTTGGAAAGGACTGCATGTATAGAAGCAGATACTAGTGATAACCATAAGGAAGATGCAGATGTTATCAAGAAAAATGTACACAAGGATATGGTAAAAAGTATGAGTATTGACAGCTTAAAACAAATTAGTAGTGATAACTTAAAATATATTAGTAGTGATAACTTAAAATATATTAGTAGTGATAACTTAAAATATATTAGTAGTGATAACCTAAAACATATTGGAAGCGATAACCTAAAACATATTGGAAGCGATAACTTCAAACATATTGGAAGCGATAACTTCAAACATATTGGAAGCGATAACTTCAAACATATTGGAAGCGATAACTTCAAACATATTGGAAGCGATAACTTCAAACATATTGGAAGCGATAACTTCAAACATATTGGAAGCGATAACTTCAAACATATTGGAAGCGATAACCTAAAACATATTGGAAGCGATAACTTCAAACATATTAGTGGTGATCACCATAATAACTATTATCATAAGAGGAAAAAGAAAAAGAAAAAAAAACATATCATTTTAAATGAAGAAGAATTATTATTACAACAGCAATTTGTGAATTATATGAAAAATGAAGAATATGATATTGAAAATGTTCATATTTTGAAAAATAATAAATTTATAAATTTTAAAAATAAATATGTTGTACATATTGATAGTATACATAAATGGATTTTTAAAATAAAAAATTTATTTTTATCAAATATAACAATTAATTATATTACGAAACGATTCCCATTTTTAAAAAATGTTATACTTGTCAAATCAAATTTTAAATGTATAAATGAGTATATAAAATATTTGAAAGAAAATGTACCACCTGAAATTAGTACTTCTCATCAGATTGTTTCTACAAATAATGAACATGATGAAGAGGATAATCACAAAAACAAAAGTAATATAATAATAAAAAATAATATGAATGAAAAGGAAGAAGAAATCAAAAAGGAAAGGTTATATAATAATCCTTCTACAGATAAAATAATAAATAAATTTGAAAACATTCAACGCTTAAGTAGTTGTAAATATGTAGATGATATGAAGGAAGAATATAATAATAATATAAAAATAGGGCATCATAAACATGATCCAAATATCATGTTAAAAACAGAAAAGGGTGTAAAAATAAAAAAAAATAAAAAAAAAATTGAAATAATGGAAAATAAAAATAAAAATAAGAGCAAAAATAAAATATCACATGATAATACTTCAAAAGAATATAATTATCCAGAGGTCAATATATATAATAACTGTGAATCTATTTTTGATGAATATATAAAAACTTCATCGATGGACAATAATAAAAAAAAAGGTTTTAAAGAGAAAAATAATGACAATAATAATAATAATAATAATAATAACACCAATATTGATAATAATGATAATAATATCGATAATAATAT**CGATAATAATATTGAAAACAATGATAATAATAATAATAATATGATAATGATGATGAAGAAAAAGAACGGTGAACCTGAGAATCCTTGTGGTGATAAAGTAAATTTAAAAAAAAAAGGAATAGAAAATGTTAAACAAAAACATAAAGAGGTGGAAATTAGTGAAGAACAAGGGAAATTAAAACAAAGCAGTGAAACAAAAGGACGTTTAAAACAACCACATACAATTAAGGTAACAGATAGATTATTTAAAGAAAGAGAAGATATGGAAAAAAATGAGAAAAGAGAATATATAGAAAAAAAAATTTTAAGTAAAGTAACTATGAATTTCTCCCCATGTGATAAGGAGACACAGAGTAAAAAGAATAAGAAAAAGAAAAAAAAAGGAATAGATGATGTAAAATTTATTTCTATGAATAAAGATGAATTGAGCAATATGTTATCCAAAGAATATGTGGAAAAAGGTAATATTAATTGTGATAATATGACATGTGAAAGTAATACAATTTGTGATAATTCAAAAAGTGAAAATTATATACCTTTAGATAATAAGACAAATAATAATAATAATAATAATAATAAAAAAAGTGCTATTAGTATAAAATCTATATTTCCAAAAAGACTTGATCATGCGCTTTCAAAATATAAAAAATATTCTACAACATCTATAAGTACTAGATTATCTAGGAGTTCATATAATAAATTCGTTTCAGAAGATAAAAATGAAGAAAACCATTTTGATAATATTGTCATAAATGATTCATATCAAAATAAGGTCAGTTTCTTTAGTCAAGTTTCTTTATCAACAGGCAATATAACTAATACTATTGATTTGGATACAAATAAAAATGAAAAAAAAAACGATGTGAATTTCAAGTCTACAATTGTCGATAATTCAAATAAACCTGTAAATCCAGAGGTGCCTGAAAAAAAAAGTAGAAAAAAGTTGAGTAAAATTAAAATTATCGGTCTTCAATTTTTTAATAAACAATCGTCAAGAAGA**TAA*ATTGTAAATTGGGCACTTATAAAAATGAACAAATATATATATATATATATATATATATATATTTATTTATTTATTGATTTATTTATTAAATTATATTCAATGTGTGTGTTCACATTCTTTTTTTTTTTTTTTTTTTTTTTTTTTTTGTTTTTGTATAATTTAAAGAATTAAGATTTTTTTTTTTTTTTTTTTTTTCTTCTTTACATATTATCATATGGACAATATATATACATATATATTTATATTTGTTTTTATTTGTATATTTATGTGTTTGAACCAGATTTAGAAGTACTAAATAGTACATACATATATGTTTTTATATATTTCTTTTAAATTATGAATAATTATATATATATATATATATGTATATAATTCTAATTAATATTCATTTTCGTTATTTTATTTTTTTTAATAATTATTTGAATAAAAATAAATAAATAATAAGTAATATGTTTAAGAGTATATTTTTTTTCATACATATATAAATATTTTGTATATTCTGTCTATAAATATG*

**Original Locus (OL) amplicon size: 1241bp**

### > PF3D7_0626400-GFP-Sandwich GFP Neo-R

tacatatatacacattcattgtattttttttttttttttttttttttttttcacttccatttatattttaaacattatttttttattcttttattaatatacaaaagaaaaaaacatttgaaggacatgtttaaaaaataaatatatacacatatttatatataattatatatgatgtggaatttttaatgttcagtgtaacattttataagaagaataggagtaatatgaaaataataataacgttcaaatagaggaatataataaaataaaaacaataaaaaagaaaataatgaaaatccatacatatatttgaaaaagaaacatacatatgtatatatatatatatatatataaatgtacattgtgtgttgtaagaattataaacattatttatatttacacttttatttaattatttatttatatttatatttacttatttatttacatttacatttataattttaattttttttttttttttttttttgtgtacaaatgtatgaatttataaaaaaattaaaaaaaaaaaaaaaaataaggttgcatactgataatatgaaatatatttatcatttggtatgaatattatattaatagaatatatatatatatatatatatatatatatatgtatatattaaaggggcatatatatgtacatattatatattttgattatataagaaatatatataagctacatatataaaataatagctatccatttatacttgtatggacaaaataagaaaggaagaaaaaaaagaaaagaaaagaaaagtATGGAACTTTCAAAAGGAAAAGTTATTATTGAAAAAGATATAAATGAACACACCATAGATGACGAAGTGTTTATGTTTGAACCTAGCATTGATGATGTGTATGATAAGAATACGAATTTACGGTTTATATTTCATAATACATTTATAACATCAGAAGAAGAAATAGCTATAAGTGAATTTAGAAAATATTGTAAGAGTAGATGCTTAAAAATTAATAAAATATATTTCGAAAATGAATGCTTACGTTATTTATATTCAGCACAATTTGATTTTTCTAAGGCCATGGAATTAATAAAAAGTAATTATGAATTTCGGTTATCATCTATTTTACCAATAAAAGAAAAAGATGTGATATTTTATATAAATAAAGGAGTTATGTATTGGCATGGGAGAGATAAAAAATGTAGACCAATATTAATTATTAATTTATTAAAAGTTGAATTATTAAGTATTGATGATTTATCTAATTTATTTTTTTTCTGTTTTGAATTTTTTTTAAAATATTTATGTATCCCAGGAAAAATAGAAAACTATATATCAATAATCGATTGTTCTGGTATATCTATATCAAAATTCCCTATGACAACATTTATGAAATTGTTAGAAATTATGAATTCTAAATATAGATGTAGATTATTTAGAATGTATATTCTTGAGGCACCTAAAATTTTAAAAACATTTGGAAAATCATTTCTAAATTTTGCACCTACCTATATGACCAAAAAATTAAAAATATTAGATAACAATTATGCAGATTATTTAAGAGAAGAAATATTATCAACTCAATTAGAAAAAAAATATGGAGGAATTCAGGAAGATAAAATTAATAATTTTTATCCTTTCCACTTTTATCCAAGTTGTTATATATCACAACAACAAAAAAGGAAATCTCAAGATAATAAAGCAGTTATCGAAAAAAAAACAAATATTTTTAATAATGATCATATTTATAATATATTCCTATCTGGTTATTCTATGCATGTAATATTAGTCCAGAAGGATCAAAGGATAATTGATAATCATCGGTATAATGATATTCTATTAAATGATTCAAATTCTGCAAATTTGGAAAGGACTGCATGTATAGAAGCAGATACTAGTGATAACCATAAGGAAGATGCAGATGTTATCAAGAAAAATGTACACAAGGATATGGTAAAAAGTATGAGTATTGACAGCTTAAAACAAATTAGTAGTGATAACTTAAAATATATTAGTAGTGATAACTTAAAATATATTAGTAGTGATAACTTAAAATATATTAGTAGTGATAACCTAAAACATATTGGAAGCGATAACCTAAAACATATTGGAAGCGATAACTTCAAACATATTGGAAGCGATAACTTCAAACATATTGGAAGCGATAACTTCAAACATATTGGAAGCGATAACTTCAAACATATTGGAAGCGATAACTTCAAACATATTGGAAGCGATAACTTCAAACATATTGGAAGCGATAACTTCAAACATATTGGAAGCGATAACCTAAAACATATTGGAAGCGATAACTTCAAACATATTAGTGGTGATCACCATAATAACTATTATCATAAGAGGAAAAAGAAAAAGAAAAAAAAACATATCATTTTAAATGAAGAAGAATTATTATTACAACAGCAATTTGTGAATTATATGAAAAATGAAGAATATGATATTGAAAATGTTCATATTTTGAAAAATAATAAATTTATAAATTTTAAAAATAAATATGTTGTACATATTGATAGTATACATAAATGGATTTTTAAAATAAAAAATTTATTTTTATCAAATATAACAATTAATTATATTACGAAACGATTCCCATTTTTAAAAAATGTTATACTTGTCAAATCAAATTTTAAATGTATAAATGAGTATATAAAATATTTGAAAGAAAATGTACCACCTGAAATTAGTACTTCTCATCAGATTGTTTCTACAAATAATGAACATGATGAAGAGGATAATCACAAAAACAAAAGTAATATAATAATAAAAAATAATATGAATGAAAAGGAAGAAGAAATCAAAAAGGAAAGGTTATATAATAATCCTTCTACAGATAAAATAATAAATAAATTTGAAAACATTCAACGCTTAAGTAGTTGTAAATATGTAGATGATATGAAGGAAGAATATAATAATAATATAAAAATAGGGCATCATAAACATGATCCAAATATCATGTTAAAAACAGAAAAGGGTGTAAAAATAAAAAAAAATAAAAAAAAAATTGAAATAATGGAAAATAAAAATAAAAATAAGAGCAAAAATAAAATATCACATGATAATACTTCAAAAGAATATAATTATCCAGAGGTCAATATATATAATAACTGTGAATCTATTTTTGATGAATATATAAAAACTTCATCGATGGACAATAATAAAAAAAAAGGTTTTAAAGAGAAAAATAATGACAATAATAATAATAATAATAATAATAACACCAATATTGATAATAATGATAATAATATCGATAATAATAT**CGATAATAATATTGAAAACAATGATAATAATAATAATAATATGATAATGATGATGAAGAAAAAGAACGGTGAACCTGAGAATCCTTGTGGTGATAAAGTAAATTTAAAAAAAAAAGGAATAGAAAATGTTAAACAAAAACATAAAGAGGTGGAAATTAGTGAAGAACAAGGGAAATTAAAACAAAGCAGTGAAACAAAAGGACGTTTAAAACAACCACATACAATTAAGGTAACAGATAGATTATTTAAAGAAAGAGAAGATATGGAAAAAAATGAGAAAAGAGAATATATAGAAAAAAAAATTTTAAGTAAAGTAACTATGAATTTCTCCCCATGTGATAAGGAGACACAGAGTAAAAAGAATAAGAAAAAGAAAAAAAAAGGAATAGATGATGTAAAATTTATTTCTATGAATAAAGATGAATTGAGCAATATGTTATCCAAAGAATATGTGGAAAAAGGTAATATTAATTGTGATAATATGACATGTGAAAGTAATACAATTTGTGATAATTCAAAAAGTGAAAATTATATACCTTTAGATAATAAGACAAATAATAATAATAATAATAATAATAAAAAAAGTGCTATTAGTATAAAATCTATATTTCCAAAAAGACTTGATCATGCGCTTTCAAAATATAAAAAATATTCTACAACATCTATAAGTACTAGATTATCTAGGAGTTCATATAATAAATTCGTTTCAGAAGATAAAAATGAAGAAAACCATTTTGATAATATTGTCATAAATGATTCATATCAAAATAAGGTCAGTTTCTTTAGTCAAGTTTCTTTATCAACAGGCAATATAACTAATACTATTGATTTGGATACAAATAAAAATGAAAAAAAAAACGATGTGAATTTCAAGTCTACAATTGTCGATAATTCAAATAAACCTGTAAATCCAGAGGTGCCTGAAAAAAAAAGTAGAAAAAAGTTGAGTAAAATTAAAATTATCGGTCTTCAATTTTTTAATAAACAATCGTCAAGAAGA**CCTAGGTCAGGATTGAGATCAAGATCTGCTGCTGCTGGTGCTGGTGGTGCTGCTAGAGCTGCTCTGCAGAGAGGAGTACAAGTTGAAACAATATCACCAGGAGATGGTCGTACATTTCCAAAAAGAGGTCAAACTTGTGTTGTACATTATACTGGAATGCTTGAAGATGGAAAGAAATTTGATTCATCTCGTGATAGAAATAAACCATTTAAATTTATGCTAGGTAAACAAGAAGTAATACGAGGTTGGGAAGAAGGAGTTGCTCAAATGAGTGTAGGTCAAAGAGCAAAACTTACTATATCTCCAGATTATGCTTATGGTGCAACTGGACATCCAGGTATAATTCCACCTCATGCAACTCTTGTATTTGATGTGGAGCTTCTAAAACTAGAAACTAGAGGTGTTCAGGTTGAAACAATTTCACCTGGAGATGGCAGAACCTTTCCTAAAAGAGGACAGACTTGCGTAGTTCATTATACAGGCATGCTAGAGGATGGTAAGAAATTTGATTCTAGTCGAGATAGAAATAAGCCATTCAAGTTTATGCTAGGTAAACAGGAAGTAATAAGAGGTTGGGAAGAGGGTGTAGCACAGATGTCAGTTGGACAAAGAGCAAAGTTAACAATATCACCAGATTATGCATACGGTGCAACAGGCCATCCTGGCATCATCCCTCCACATGCAACTTTAGTATTCGACGTTGAATTGTTAAAGTTAGAGACAACGCGTGCTAGAGGTGCTGCTGCTGGTGCTGGAGGTGCAGGTAGACGTACGATGAGTAAAGGAGAAGAACTTTTCACTGGAGTTGTCCCAATTCTTGTTGAATTAGATGGTGATGTTAATGGGCACAAATTTTCTGTCAGTGGAGAGGGTGAAGGTGATGCAACATACGGAAAACTTACCCTTAAATTTATTTGCACTACTGGAAAACTACCTGTTCCATGGCCAACACTTGTCACTACTTTCGCGTATGGTCTTCAATGCTTTGCGAGATACCCAGATCATATGAAACAGCATGACTTTTTCAAGAGTGCCATGCCCGAAGGTTATGTACAGGAAAGAACTATATTTTTCAAAGATGACGGGAACTACAAGACACGTGCTGAAGTCAAGTTTGAAGGTGATACCCTTGTTAATAGAATCGAGTTAAAAGGTATTGATTTTAAAGAAGATGGAAACATTCTTGGACACAAATTGGAATACAACTATAACTCACACAATGTATACATCATGGCAGACAAACAAAAGAATGGAATCAAAGTTAACTTCAAAATTAGACACAACATTGAAGATGGAAGCGTTCAACTAGCAGACCATTATCAACAAAATACTCCAATTGGCGATGGCCCTGTCCTTTTACCAGACAACCATTACCTGTCCACACAATCTGCCCTTTCGAAAGATCCCAACGAAAAGAGAGACCACATGGTCCTTCTTGAGTTTGTAACAGCTGCTGGGATTACACATGGCATGGATGAGCTCTACAAAGTCGACGCCAGGGGAGCAGCCGCAGGAGCAGGGGGGGCAGGAAGGCGTGGTGTTCAGGTCGAGACTATTAGCCCTGGAGATGGACGCACGTTTCCTAAGCGTGGACAGACATGCGTAGTTCACTACACAGGTATGTTGGAGGACGGTAAAAAGTTCGACAGCTCACGCGACCGCAATAAACCTTTCAAGTTTATGCTTGGCAAGCAGGAGGTTATTCGTGGATGGGAGGAGGGTGTAGCACAGATGTCTGTTGGACAGCGTGCTAAGTTGACAATTTCACCTGACTATGCTTATGGCGCTACGGGCCATCCCGGGATCATTCCGCCACATGCGACTCTGGTATTCGACGTTGAATTATTAAAGTTAGAGACAGCTAGAGGGGCCGCTGCAGGTGCTGGTGGAGCTGGAAGACGTGGAGTACAAGTAGAGACTATCTCTCCAGGTGACGGTCGCACTTTCCCAAAGCGTGGCCAAACCTGTGTTGTACATTACACTGGTATGCTGGAGGATGGGAAAAAGTTCGATTCCAGTCGCGACCGTAACAAACCGTTCAAATTCATGTTGGGAAAGCAGGAAGTGATCCGCGGGTGGGAGGAAGGCGTGGCGCAAATGAGCGTCGGTCAGCGGGCTAAATTGACCATTTCCCCTGACTACGCGTATGGGGCTACTGGGCACCCAGGGATTATTCCGCCTCACGCTACACTTGTGTTTGATGTCGAACTTTTGAAACTGGAAACTGTCGACGGAGAAGGAAGAGGAAGTTTATTAACATGTGGAGATGTAGAAGAAAATCCAGGACCAATGATTGAACAAGATGGATTGCACGCAGGTTCTCCGGCCGCTTGGGTGGAGAGGCTATTCGGCTATGACTGGGCACAACAGACAATCGGCTGCTCTGATGCCGCCGTGTTCCGGCTGTCAGCGCAGGGGCGCCCGGTTCTTTTTGTCAAGACCGACCTGTCCGGTGCCCTGAATGAACTGCAGGACGAGGCAGCGCGGCTATCGTGGCTGGCCACGACGGGCGTTCCTTGCGCAGCTGTGCTCGACGTTGTCACTGAAGCGGGAAGGGACTGGCTGCTATTGGGCGAAGTGCCGGGGCAGGATCTCCTGTCATCTCACCTTGCTCCTGCCGAGAAAGTATCCATCATGGCTGATGCAATGCGGCGGCTGCATACGCTTGATCCGGCTACCTGCCCATTCGACCACCAAGCGAAACATCGCATCGAGCGAGCACGTACTCGGATGGAAGCCGGTCTTGTCGATCAGGATGATCTGGACGAAGAGCATCAGGGGCTCGCGCCAGCCGAACTGTTCGCCAGGCTCAAGGCGCGCATGCCCGACGGCGAGGATCTCGTCGTGACCCATGGCGATGCCTGCTTGCCGAATATCATGGTGGAAAATGGCCGCTTTTCTGGATTCATCGACTGTGGCCGGCTGGGTGTGGCGGACCGCTATCAGGACATAGCGTTGGCTACCCGTGATATTGCTGAAGAGCTTGGCGGCGAATGGGCTGACCGCTTCCTCGTGCTTTACGGTATCGCCGCTCCCGATTCGCAGCGCATCGCCTTCTATCGCCTTCTTGACGAGTTCTTCTAActcgagggatatggcagcttaatgttcgtttttcttatttatatatttataccaattgattgtatttataactgtaaaaatgtgtatgttgtgtgcatatttttttttgtgcatgcacatgcatgtaaatagctaaaattatgaacattttattttttgttcagaaaaaaaaaactttacacacataaaatggctagtatgaatagccatattttatataaattaaatcctatgaatttatgaccatattaaaaatttagatatttatggaacataatatgtttgaaacaataagacaaaattattattattattattatttttactgttataattatgtgtctccttcaatgattcataaatagttggacttgatttttaaaatgtttataatatgattagcatagttaaataaaaaaagttgaaaaattaaaaaaaaacatataaacacaaatgatggtttttccttcaatttcgatatcaatttatagaaacaaaatatatacttgtataattttatttttttatataaatcattacatatataattatacaatattttttctaagagataattatatattaatatatataaaaaaaggtgttttttttttttttttttatttttatttttattttatggtaatattttattttccttattttataaattatattagtttatatgtgattaattttatatattatcaatttatatatttttaaatgcttacttaattatctttttttttttttttttttttttttcccctctttttatattaatttatttttgaaaaaattgatatatatatatatatataatatatatatatacatgtagtagtattaaacaatgtataatatatataaataatatatttatatatttcatttcaattttaattttttttggttttttttttttttctttttgtcatatttaaaaaaaattatattcatataagttatgcattttttataaacattattcaatatatgtataatataatatatatatatatattaatgtattattccaatgtgcatgataaaagaaaaaaataatatttataaaaaaaaagaaaaataaaacaaaaaaagaaaaaaaaaaaaaaaaaaaaaaaaatacaaaaataaataatataatttataattatatattcttgtcacaataaaaatatatatatatatatatatatttataatatgtatattttaaactagaaaaggaataactaatattttatttattatcattcaagatttatattttataataataaatacctaatagaaatatatcaggatccatgcatggttcgctaaactgcatcgtcgctgtgtcccagaacatgggcatcggcaagaacggggactacccctggccaccgctcaggaacgaatttagatatttccagagaatgaccacaacctcttcagtagaaggtaaacagaatctggtgattatgggtaagaagacctggttctccattcctgagaagaatcgacctttaaagggtagaattaatttagttctcagcagagaactcaaggaacctccacaaggagctcattttctttccagaagtctagatgatgccttaaaacttactgaacaaccagaattagcaaataaagtagacatggtctggatagttggtggcagttctgtttataaggaagccatgaatcacccaggccatcttaaactatttgtgacaaggatcatgcaagactttgaaagtgacacgttttttccagaaattgatttggagaaatataaacttctgccagaatacccaggtgttctctctgatgtccaggaggagaaaggcattaagtacaaatttgaagtatatgagaagaatgattaagcttatttaataatagattaaaaatattataaaaataaaaacataaacacagaaattacaaaaaaaatacatatgaattttttttttgtaatcttccttataaatatagaataatgaatcatataaaacatatcattattcatttatttacatttaaaattattgtttcagtatctttaatttattatgtatatataaaaataacttacaattttattaataaacaatatatgtttattaattcatgttttgtaatttatgggatagcgattttttttactgtctgtatttttcttttttaattatgttttaattgtattttatttttattattgttctttttatagtattattttaaaacaaaatgtattttctaagaacttataataataataaatataaattttaataaaaattatatttatcttttacaatatgaacataaagtacaacattaatatatagcttttaatatttttattcctaatcatgtaaatcttaaatttttctttttaaacatatgttaaatatttatttctcattatatataagaacatatttattaaatctagaattctatagtgagtcgtattacaattcactggccgtcgttttacaacgtcgtgactgggaaaaccctggcgttacccaacttaatcgccttgcagcacatccccctttcgccagctggcgtaatagcgaagaggcccgcaccgatcgcccttcccaacagttgcgcagcctgaatggcgaatggcgcctgatgcggtattttctccttacgcatctgtgcggtatttcacaccgcatatggtgcactctcagtacaatctgctctgatgccgcatagttaagccagccccgacacccgccaacacccgctgacgcgccctgacgggcttgtctgctcccggcatccgcttacagacaagctgtgaccgtctccgggagctgcatgtgtcagaggttttcaccgtcatcaccgaaacgcgcgagacgaaagggcctcgtgatacgcctatttttataggttaatgtcatgataataatggtttcttagacgtcaggtggcacttttcggggaaatgtgcgcggaacccctatttgtttatttttctaaatacattcaaatatgtatccgctcatgagacaataaccctgataaatgcttcaataatattgaaaaaggaagagtatgagtattcaacatttccgtgtcgcccttattcccttttttgcggcattttgccttcctgtttttgctcacccagaaacgctggtgaaagtaaaagatgctgaagatcagttgggtgcacgagtgggttacatcgaactggatctcaacagcggtaagatccttgagagttttcgccccgaagaacgttttccaatgatgagcacttttaaagttctgctatgtggcgcggtattatcccgtattgacgccgggcaagagcaactcggtcgccgcatacactattctcagaatgacttggttgagtactcaccagtcacagaaaagcatcttacggatggcatgacagtaagagaattatgcagtgctgccataaccatgagtgataacactgcggccaacttacttctgacaacgatcggaggaccgaaggagctaaccgcttttttgcacaacatgggggatcatgtaactcgccttgatcgttgggaaccggagctgaatgaagccataccaaacgacgagcgtgacaccacgatgcctgtagcaatgccaacaacgttgcgcaaactattaactggcgaactacttactctagcttcccggcaacaattaatagactggatggaggcggataaagttgcaggaccacttctgcgctcggcccttccggctggctggtttattgctgataaatctggagccggtgagcgtgggtctcgcggtatcattgcagcactggggccagatggtaagccctcccgtatcgtagttatctacacgacggggagtcaggcaactatggatgaacgaaatagacagatcgctgagataggtgcctcactgattaagcattggtaactgtcagaccaagtttactcatatatactttagattgatttaaaacttcatttttaatttaaaaggatctaggtgaagatcctttttgataatctcatgaccaaaatcccttaacgtgagttttcgttccactgagcgtcagaccccgtagaaaagatcaaaggatcttcttgagatcctttttttctgcgcgtaatctgctgcttgcaaacaaaaaaaccaccgctaccagcggtggtttgtttgccggatcaagagctaccaactctttttccgaaggtaactggcttcagcagagcgcagataccaaatactgtccttctagtgtagccgtagttaggccaccacttcaagaactctgtagcaccgcctacatacctcgctctgctaatcctgttaccagtggctgctgccagtggcgataagtcgtgtcttaccgggttggactcaagacgatagttaccggataaggcgcagcggtcgggctgaacggggggttcgtgcacacagcccagcttggagcgaacgacctacaccgaactgagatacctacagcgtgagctatgagaaagcgccacgcttcccgaagggagaaaggcggacaggtatccggtaagcggcagggtcggaacaggagagcgcacgagggagcttccagggggaaacgcctggtatctttatagtcctgtcgggtttcgccacctctgacttgagcgtcgatttttgtgatgctcgtcaggggggcggagcctatcgaaaaacgccagcaacgcggcctttttacggttcctggccttttgctggccttttgctcacatgttctttcctgcgttatcccctgattctgtggataaccgtattaccgcctttgagtgagctgataccgctcgccgcagccgaacgaccgagcgcagcgagtcagtgagcgaggaagcggaagagcgcccaatacgcaaaccgcctctccccgcgcgttggccgattcattaatgcagctggcacgacaggtttcccgactggaaagcgggcagtgagcgcaacgcaattaatgtgagttagctcactcattaggcaccccaggctttacactttatgcttccggctcgtatgttgtgtggaattgtgagcggataacaatttcacacaggaaacagctatgaccatgattacgccaagctatttaggtgacactatagaatactcgcggccgctagcgCGATAATAATATTGAAAACAATGATAATAATAATAATAATATGATAATGATGATGAAGAAAAAGAACGGTGAACCTGAGAATCCTTGTGGTGATAAAGTAAATTTAAAAAAAAAAGGAATAGAAAATGTTAAACAAAAACATAAAGAGGTGGAAATTAGTGAAGAACAAGGGAAATTAAAACAAAGCAGTGAAACAAAAGGACGTTTAAAACAACCACATACAATTAAGGTAACAGATAGATTATTTAAAGAAAGAGAAGATATGGAAAAAAATGAGAAAAGAGAATATATAGAAAAAAAAATTTTAAGTAAAGTAACTATGAATTTCTCCCCATGTGATAAGGAGACACAGAGTAAAAAGAATAAGAAAAAGAAAAAAAAAGGAATAGATGATGTAAAATTTATTTCTATGAATAAAGATGAATTGAGCAATATGTTATCCAAAGAATATGTGGAAAAAGGTAATATTAATTGTGATAATATGACATGTGAAAGTAATACAATTTGTGATAATTCAAAAAGTGAAAATTATATACCTTTAGATAATAAGACAAATAATAATAATAATAATAATAATAAAAAAAGTGCTATTAGTATAAAATCTATATTTCCAAAAAGACTTGATCATGCGCTTTCAAAATATAAAAAATATTCTACAACATCTATAAGTACTAGATTATCTAGGAGTTCATATAATAAATTCGTTTCAGAAGATAAAAATGAAGAAAACCATTTTGATAATATTGTCATAAATGATTCATATCAAAATAAGGTCAGTTTCTTTAGTCAAGTTTCTTTATCAACAGGCAATATAACTAATACTATTGATTTGGATACAAATAAAAATGAAAAAAAAAACGATGTGAATTTCAAGTCTACAATTGTCGATAATTCAAATAAACCTGTAAATCCAGAGGTGCCTGAAAAAAAAAGTAGAAAAAAGTTGAGTAAAATTAAAATTATCGGTCTTCAATTTTTTAATAAACAATCGTCAAGAAGATAA*ATTGTAAATTGGGCACTTATAAAAATGAACAAATATATATATATATATATATATATATATATTTATTTATTTATTGATTTATTTATTAAATTATATTCAATGTGTGTGTTCACATTCTTTTTTTTTTTTTTTTTTTTTTTTTTTTTGTTTTTGTATAATTTAAAGAATTAAGATTTTTTTTTTTTTTTTTTTTTTCTTCTTTACATATTATCATATGGACAATATATATACATATATATTTATATTTGTTTTTATTTGTATATTTATGTGTTTGAACCAGATTTAGAAGTACTAAATAG*

**5’ amplicon size: 1340bps**

**3’ amplicon size: 1133bps**

### > PF3D7_1127600 (PfCT3)

gattattatatatatataatatattagaactaggaaaatttatacgtatattattatatatatatatatatatatatatatataaatttttatgtgttgtttttttttttttttttttttttttcttaattatataaatatattttttttaatatttttaaaaattctattttatatattatatatgtatgtgcaattttttttttttcttgttttccatatttgaaaatttaaaaaatagaaaaaaaaaaaaaaaaaaattacacaaaattgtatagtaacacatatatatatttttaattttttaatttttttttattactattaattatatgtatatatatatatatatatatataaatatatatataatataatataataacgtaatatattattctttgttttaaatatatgtaaagatatatatatatatatatatataaatatataattatatatattacataaatctttgtcaattgaatttttttttttttttttttttttttaaatatataagtataaataataATGAACGACAATCCTGTTTCAAGTTATTTGTCGAAGGATGTTTTAGATTATGAACCGACTAAGGAGGAAATTAAATTT*gtaaaaataaaaaaaataataaaaaaaaatgagaaaaatagaattcacgctatgtaaattatgtagataaatatattatatgtgtgtatatcattactacgtacatgtatatatattgtattatacatataaataaatatatgtatatatatatatatatatatgtatatattcacatttatatatatttttttacatttgtag*TATCATAAAAATAACCTAAAAAGTTTAAGATATATATTTTGTGGAAAAGAGCTTGATGACTTTGAGAAACAGAAAATAAGAGAATTAAAAGAATTTGTTAATAAATTAAAATTAAAGGAAAAAGATAAAGAGAAAGATAAAGAAAAAGAAGTTGAAACATACCAAACTATTTTTAAAAATACCCTTTTTGATGATGATAATTATGTTCTAAGATTTCTACAAGG*gttaatatataaataataaaaaaaatattaaaataataatatgttatgtatcatattttaaatatataattataaacatatttaaaatgtaataaaacatttatatttctttttgtag*AAATGAATTTGTTTTTGAAAGATGTTATAACGATATGTTAAGACACTTAACGTGGAGAAAAGAAAATCTACCAATACCCCTGAGTGACGTTCAAATTTTTTTA*gtaacttacaaaaaaaaaaaaaaaaaaaaaggaaaaataataatagctcatatatatatatatatatatatatatttatttatttatatatttatttgtgtattttattttgtag*GACAAGGGATATTGTTATATTCATGGAAGGGATAAACAAATGCACCCAATCATAATTATTAATTGCAAAAATATTATATCAGCTAATACT*gtaatgatttaaattataaaaattatataggagaacttattcactttattattttattttattttattttaatttttttttttttttttttttgtttttacatatttcag*AAAGATGTTTTAAAAGTTCTTTATTATTGGCTAG*gtaacacacaaaaaaaaaaaaaaaaaaaaaaaaaaaaaaaaaattgtgtacttaatttatttttttccatttgtatatctaatggatatatttttttaatgttcattagtccttatgttaaatataaatatatttattttattttgtag*AATTTTGTATATCCAAATTA**TTAATAGAAGGAAAAGTAGAACAATGGAGAGTTATAATTGATTTAACATCATGTAGTGTTTTAAATATACCTATAGCAACTTTAAAGGATATCTCAAAAAATTTAAGT*gtaagaaaagaaaaatatatgaacatatatatatatatatatatatatatatttatttatttatttatatttatttatttatatttcccatcag*TGTAATTATCGATCAAGGCTGTGTAAAATGCTTGTGTTAAGTGCTCCTTTAGTTGTTACAGGAATATGGCATATGTTAAAATCCATCATACCA*gtaattgaataaattatcatatatatattatttatgtatttatattttctaatttattcataaatatattttatatatgatactttaaaaatatgtag*GTTGTAACACAACAAAAAATTACAATAACATCATCTGAAAAGGAG*gtaaataaggaataaaaataaaataaaaaggaataatatatagtatatatttatatatataatagtaaaaatatatacctttatatatttttattttttattttattatattattttatttttttttttttctttctctctttatag*AAGAAGTTGCTAGATCAAGTTCAGGCAAATCAATTAGAAA*gtaagcatatattattattacatatgccaatatatttgtcttttaaacctgtacacacatatatatatatatatatatatatatatatttatttatttatttacacatgtttattttatgttttttttaacag*AAAAATTCGGTGGAACCTGTGAGAATGCAACAGATTTTACAGAACCAATATTGCCA**TAAtttgttatatattcttttatgttaataattcgaataaaatatgttttatatgtataatttatatatatatatatattttttaatttttttatttgttttatattatgatcatataattattatatattcttttgttattatatatgcttcatttgattattgttatgttgctgttacataattccaagtttatttttctttatatcgcttgttgctgtatttgtatgcttatgcatacatacatatatatatatatatatatatatatattgtgtatagttttatatattttgtatacctcactatcttgtatgttataagtaaatatatatatttatatatatatgtttatttatttatttatttattaattttaatttaattttttttgtttacttttttttttttttttttttttttttttttttcaagttaaaaagccaaacaaaagaggtttatattttaaaacttctttatatatatatttatttattcataactttcatattttttaaaggaaaaagtataaattgtcaatttatttttcatattgtagaattagattcaaatatataatatatatatgtatggtatcataaatattttttttatatgtattatttaagatttataaacgtttaaaagttcgaaattatatatatatatttatatatatatttcaaaataatatacaaaagattgaaaatgttattattgagttaaaatatatatgtataattaaaatgtttcataagttatgaaatatataaattttataaaatggacatgtatcattcacttaaatatatattttataaatcttggtcataaaatatgtgatagacaattcttttcaagaaaattataacaaatccaaaaaagatatacgataaaaa

**Original Locus (OL) amplicon size: 1429bp**

### > PF3D7_1127600-GFP-Sandwich GFP Neo-R

gattattatatatatataatatattagaactaggaaaatttatacgtatattattatatatatatatatatatatatatatataaatttttatgtgttgtttttttttttttttttttttttttcttaattatataaatatattttttttaatatttttaaaaattctattttatatattatatatgtatgtgcaattttttttttttcttgttttccatatttgaaaatttaaaaaatagaaaaaaaaaaaaaaaaaaattacacaaaattgtatagtaacacatatatatatttttaattttttaatttttttttattactattaattatatgtatatatatatatatatatatataaatatatatataatataatataataacgtaatatattattctttgttttaaatatatgtaaagatatatatatatatatatatataaatatataattatatatattacataaatctttgtcaattgaatttttttttttttttttttttttttaaatatataagtataaataataATGAACGACAATCCTGTTTCAAGTTATTTGTCGAAGGATGTTTTAGATTATGAACCGACTAAGGAGGAAATTAAATTT*gtaaaaataaaaaaaataataaaaaaaaatgagaaaaatagaattcacgctatgtaaattatgtagataaatatattatatgtgtgtatatcattactacgtacatgtatatatattgtattatacatataaataaatatatgtatatatatatatatatatatgtatatattcacatttatatatatttttttacatttgtag*TATCATAAAAATAACCTAAAAAGTTTAAGATATATATTTTGTGGAAAAGAGCTTGATGACTTTGAGAAACAGAAAATAAGAGAATTAAAAGAATTTGTTAATAAATTAAAATTAAAGGAAAAAGATAAAGAGAAAGATAAAGAAAAAGAAGTTGAAACATACCAAACTATTTTTAAAAATACCCTTTTTGATGATGATAATTATGTTCTAAGATTTCTACAAGG*gttaatatataaataataaaaaaaatattaaaataataatatgttatgtatcatattttaaatatataattataaacatatttaaaatgtaataaaacatttatatttctttttgtag*AAATGAATTTGTTTTTGAAAGATGTTATAACGATATGTTAAGACACTTAACGTGGAGAAAAGAAAATCTACCAATACCCCTGAGTGACGTTCAAATTTTTTTA*gtaacttacaaaaaaaaaaaaaaaaaaaaaggaaaaataataatagctcatatatatatatatatatatatatatttatttatttatatatttatttgtgtattttattttgtag*GACAAGGGATATTGTTATATTCATGGAAGGGATAAACAAATGCACCCAATCATAATTATTAATTGCAAAAATATTATATCAGCTAATACT*gtaatgatttaaattataaaaattatataggagaacttattcactttattattttattttattttattttaatttttttttttttttttttttgtttttacatatttcag*AAAGATGTTTTAAAAGTTCTTTATTATTGGCTAG*gtaacacacaaaaaaaaaaaaaaaaaaaaaaaaaaaaaaaaaattgtgtacttaatttatttttttccatttgtatatctaatggatatatttttttaatgttcattagtccttatgttaaatataaatatatttattttattttgtag*AATTTTGTATATCCAAATTA**TTAATAGAAGGAAAAGTAGAACAATGGAGAGTTATAATTGATTTAACATCATGTAGTGTTTTAAATATACCTATAGCAACTTTAAAGGATATCTCAAAAAATTTAAGT*gtaagaaaagaaaaatatatgaacatatatatatatatatatatatatatatttatttatttatttatatttatttatttatatttcccatcag*TGTAATTATCGATCAAGGCTGTGTAAAATGCTTGTGTTAAGTGCTCCTTTAGTTGTTACAGGAATATGGCATATGTTAAAATCCATCATACCA*gtaattgaataaattatcatatatatattatttatgtatttatattttctaatttattcataaatatattttatatatgatactttaaaaatatgtag*GTTGTAACACAACAAAAAATTACAATAACATCATCTGAAAAGGAG*gtaaataaggaataaaaataaaataaaaaggaataatatatagtatatatttatatatataatagtaaaaatatatacctttatatatttttattttttattttattatattattttatttttttttttttctttctctctttatag*AAGAAGTTGCTAGATCAAGTTCAGGCAAATCAATTAGAAA*gtaagcatatattattattacatatgccaatatatttgtcttttaaacctgtacacacatatatatatatatatatatatatatatatttatttatttatttacacatgtttattttatgttttttttaacag*AAAAATTCGGTGGAACCTGTGAGAATGCAACAGATTTTACAGAACCAATATTGCCA**CCTAGGTCAGGATTGAGATCAAGATCTGCTGCTGCTGGTGCTGGTGGTGCTGCTAGAGCTGCTCTGCAGAGAGGAGTACAAGTTGAAACAATATCACCAGGAGATGGTCGTACATTTCCAAAAAGAGGTCAAACTTGTGTTGTACATTATACTGGAATGCTTGAAGATGGAAAGAAATTTGATTCATCTCGTGATAGAAATAAACCATTTAAATTTATGCTAGGTAAACAAGAAGTAATACGAGGTTGGGAAGAAGGAGTTGCTCAAATGAGTGTAGGTCAAAGAGCAAAACTTACTATATCTCCAGATTATGCTTATGGTGCAACTGGACATCCAGGTATAATTCCACCTCATGCAACTCTTGTATTTGATGTGGAGCTTCTAAAACTAGAAACTAGAGGTGTTCAGGTTGAAACAATTTCACCTGGAGATGGCAGAACCTTTCCTAAAAGAGGACAGACTTGCGTAGTTCATTATACAGGCATGCTAGAGGATGGTAAGAAATTTGATTCTAGTCGAGATAGAAATAAGCCATTCAAGTTTATGCTAGGTAAACAGGAAGTAATAAGAGGTTGGGAAGAGGGTGTAGCACAGATGTCAGTTGGACAAAGAGCAAAGTTAACAATATCACCAGATTATGCATACGGTGCAACAGGCCATCCTGGCATCATCCCTCCACATGCAACTTTAGTATTCGACGTTGAATTGTTAAAGTTAGAGACAACGCGTGCTAGAGGTGCTGCTGCTGGTGCTGGAGGTGCAGGTAGACGTACGATGAGTAAAGGAGAAGAACTTTTCACTGGAGTTGTCCCAATTCTTGTTGAATTAGATGGTGATGTTAATGGGCACAAATTTTCTGTCAGTGGAGAGGGTGAAGGTGATGCAACATACGGAAAACTTACCCTTAAATTTATTTGCACTACTGGAAAACTACCTGTTCCATGGCCAACACTTGTCACTACTTTCGCGTATGGTCTTCAATGCTTTGCGAGATACCCAGATCATATGAAACAGCATGACTTTTTCAAGAGTGCCATGCCCGAAGGTTATGTACAGGAAAGAACTATATTTTTCAAAGATGACGGGAACTACAAGACACGTGCTGAAGTCAAGTTTGAAGGTGATACCCTTGTTAATAGAATCGAGTTAAAAGGTATTGATTTTAAAGAAGATGGAAACATTCTTGGACACAAATTGGAATACAACTATAACTCACACAATGTATACATCATGGCAGACAAACAAAAGAATGGAATCAAAGTTAACTTCAAAATTAGACACAACATTGAAGATGGAAGCGTTCAACTAGCAGACCATTATCAACAAAATACTCCAATTGGCGATGGCCCTGTCCTTTTACCAGACAACCATTACCTGTCCACACAATCTGCCCTTTCGAAAGATCCCAACGAAAAGAGAGACCACATGGTCCTTCTTGAGTTTGTAACAGCTGCTGGGATTACACATGGCATGGATGAGCTCTACAAAGTCGACGCCAGGGGAGCAGCCGCAGGAGCAGGGGGGGCAGGAAGGCGTGGTGTTCAGGTCGAGACTATTAGCCCTGGAGATGGACGCACGTTTCCTAAGCGTGGACAGACATGCGTAGTTCACTACACAGGTATGTTGGAGGACGGTAAAAAGTTCGACAGCTCACGCGACCGCAATAAACCTTTCAAGTTTATGCTTGGCAAGCAGGAGGTTATTCGTGGATGGGAGGAGGGTGTAGCACAGATGTCTGTTGGACAGCGTGCTAAGTTGACAATTTCACCTGACTATGCTTATGGCGCTACGGGCCATCCCGGGATCATTCCGCCACATGCGACTCTGGTATTCGACGTTGAATTATTAAAGTTAGAGACAGCTAGAGGGGCCGCTGCAGGTGCTGGTGGAGCTGGAAGACGTGGAGTACAAGTAGAGACTATCTCTCCAGGTGACGGTCGCACTTTCCCAAAGCGTGGCCAAACCTGTGTTGTACATTACACTGGTATGCTGGAGGATGGGAAAAAGTTCGATTCCAGTCGCGACCGTAACAAACCGTTCAAATTCATGTTGGGAAAGCAGGAAGTGATCCGCGGGTGGGAGGAAGGCGTGGCGCAAATGAGCGTCGGTCAGCGGGCTAAATTGACCATTTCCCCTGACTACGCGTATGGGGCTACTGGGCACCCAGGGATTATTCCGCCTCACGCTACACTTGTGTTTGATGTCGAACTTTTGAAACTGGAAACTGTCGACGGAGAAGGAAGAGGAAGTTTATTAACATGTGGAGATGTAGAAGAAAATCCAGGACCAATGATTGAACAAGATGGATTGCACGCAGGTTCTCCGGCCGCTTGGGTGGAGAGGCTATTCGGCTATGACTGGGCACAACAGACAATCGGCTGCTCTGATGCCGCCGTGTTCCGGCTGTCAGCGCAGGGGCGCCCGGTTCTTTTTGTCAAGACCGACCTGTCCGGTGCCCTGAATGAACTGCAGGACGAGGCAGCGCGGCTATCGTGGCTGGCCACGACGGGCGTTCCTTGCGCAGCTGTGCTCGACGTTGTCACTGAAGCGGGAAGGGACTGGCTGCTATTGGGCGAAGTGCCGGGGCAGGATCTCCTGTCATCTCACCTTGCTCCTGCCGAGAAAGTATCCATCATGGCTGATGCAATGCGGCGGCTGCATACGCTTGATCCGGCTACCTGCCCATTCGACCACCAAGCGAAACATCGCATCGAGCGAGCACGTACTCGGATGGAAGCCGGTCTTGTCGATCAGGATGATCTGGACGAAGAGCATCAGGGGCTCGCGCCAGCCGAACTGTTCGCCAGGCTCAAGGCGCGCATGCCCGACGGCGAGGATCTCGTCGTGACCCATGGCGATGCCTGCTTGCCGAATATCATGGTGGAAAATGGCCGCTTTTCTGGATTCATCGACTGTGGCCGGCTGGGTGTGGCGGACCGCTATCAGGACATAGCGTTGGCTACCCGTGATATTGCTGAAGAGCTTGGCGGCGAATGGGCTGACCGCTTCCTCGTGCTTTACGGTATCGCCGCTCCCGATTCGCAGCGCATCGCCTTCTATCGCCTTCTTGACGAGTTCTTCTAActcgagggatatggcagcttaatgttcgtttttcttatttatatatttataccaattgattgtatttataactgtaaaaatgtgtatgttgtgtgcatatttttttttgtgcatgcacatgcatgtaaatagctaaaattatgaacattttattttttgttcagaaaaaaaaaactttacacacataaaatggctagtatgaatagccatattttatataaattaaatcctatgaatttatgaccatattaaaaatttagatatttatggaacataatatgtttgaaacaataagacaaaattattattattattattatttttactgttataattatgtgtctccttcaatgattcataaatagttggacttgatttttaaaatgtttataatatgattagcatagttaaataaaaaaagttgaaaaattaaaaaaaaacatataaacacaaatgatggtttttccttcaatttcgatatcaatttatagaaacaaaatatatacttgtataattttatttttttatataaatcattacatatataattatacaatattttttctaagagataattatatattaatatatataaaaaaaggtgttttttttttttttttttatttttatttttattttatggtaatattttattttccttattttataaattatattagtttatatgtgattaattttatatattatcaatttatatatttttaaatgcttacttaattatctttttttttttttttttttttttttcccctctttttatattaatttatttttgaaaaaattgatatatatataGTTGAAATATAAATTTCAAAAAAAATGATCACAAAATATACACTTAAATATAGGTACAATAAAAAAAAAAAATAAAAATATAATTACAAGATAATATTTTTTCCTGCTATCAAATTTTTATATATTCTCCTCAAAGAAAAAATATAAATAAATGAAGTAAATTAAAAAAAAAATTTTCTTTTTCTTCTTCTTTTGTAATTCCTTATTTATACATATTTTACTATATTTTCATAAAAATAAATTGTCATATTATATAAATATATATACCAAACCATAATTATATAGCCCTCAACATATTTTTATGATGTTTTTTCTTTTTAAATGTGATACGTAATTAAATATAATAATATATATTAAATATTATATTTTGTAATATTTACTTTCATGAGGTTTAATAAATTATAAAGAGAGAATAAAAAAAAAAAAAAAAAAAATTTATCCATATACAAATTATTAATTTATTTTTATTTTTTATTTCCCTTTGTATATATTATAAAAAAATAATCTACATAATTTTATATGATGATAATTACAATATAATATTTATAATATATATTATTTGTTAAGAGAAAAAAAAAATAAATATATACCTTCTTTTTGAGAATTGAATAAATTGTTTAATATATATATATATATATATAAATATATTATACATTTATGGTGAAAAAAAATGTTGTATTTAATTAGATTTAATATATATATATAAAAAATAGCGTATTTAAAATAATATATATATATATATATTATTATTATTACAAAATGACGGATATTATAAAAGTATATATCTATATATATGTATATATATATAATATTATTTTACTATATATATATAATATATAAATGTAAATGCATATAGTATCTATGTATATTATATATATATATAATATTAATTACATATCTAAGTTTTCTTTTCTTTTCTTTTTTTTTTTTTTTTTTTTTATTTTTTATAGAGAGCCCGTTATATATTATATATTAAAATTTTTTATAATAGCATTTATATACAATATTTTATTAACTAAAAGAAAAAAAAAAAAAAAAAAAAAAAAAAAACGAGGAAATTTATATTTCTTTAACAACATTTTAATTAAAATCATGATAATACAATTTCAAATCATTTTGTAATTATATAAATAAATATATATATATATATATATATATATATGTACTTTTAAATTAGGAATATTCTCATTTATAAATATATCTTATTTTTTAAATTGGTATAAAAAAAAAAAAAAAAATAAGAAACCGTTGATTAAATAATACATATATAATATAAATATATTTTATAAATATATATTTATATATATATATATATATATTTATAACGTATATCATTTTAAAGATAAACTAGTatggcaccaaaaaaaaaaagaaaagttacgcgtgatccaacaagaagtgcaaatagtggagcaggagcaggagcaggagcaatattaagtagagctagcatggcttctagaatcctctggcatgagatGtggcatgaaggcctggaagaggcatctcgtttgtactttggggaaaggaacgtgaaaggCatgtttgaggtgctggagcccttgcatgctatgatggaacggggcccccagactctgaaGgaaacatcctttaatcaggcctatggtcgagatttaatggaggcccaagagtggtgcagGaagtacatgaaatcagggaatgtcaaggacctcctccaagcctgggacctctattatcaTgtgttccgacgaatctcaaagGGTGAGGGTCGTGGTTCACTTCTTACTTGCGGTGACGTTGAGGAGAACCCTGGTCCTGGTACCggatccatgcatggttcgctaaactgcatcgtcgctgtgtcccagaacatgggcatcggcaagaacggggactacccctggccaccgctcaggaacgaatttagatatttccagagaatgaccacaacctcttcagtagaaggtaaacagaatctggtgattatgggtaagaagacctggttctccattcctgagaagaatcgacctttaaagggtagaattaatttagttctcagcagagaactcaaggaacctccacaaggagctcattttctttccagaagtctagatgatgccttaaaacttactgaacaaccagaattagcaaataaagtagacatggtctggatagttggtggcagttctgtttataaggaagccatgaatcacccaggccatcttaaactatttgtgacaaggatcatgcaagactttgaaagtgacacgttttttccagaaattgatttggagaaatataaacttctgccagaatacccaggtgttctctctgatgtccaggaggagaaaggcattaagtacaaatttgaagtatatgagaagaatgattaagcttatttaataatagattaaaaatattataaaaataaaaacataaacacagaaattacaaaaaaaatacatatgaattttttttttgtaatcttccttataaatatagaataatgaatcatataaaacatatcattattcatttatttacatttaaaattattgtttcagtatctttaatttattatgtatatataaaaataacttacaattttattaataaacaatatatgtttattaattcatgttttgtaatttatgggatagcgattttttttactgtctgtatttttcttttttaattatgttttaattgtattttatttttattattgttctttttatagtattattttaaaacaaaatgtattttctaagaacttataataataataaatataaattttaataaaaattatatttatcttttacaatatgaacataaagtacaacattaatatatagcttttaatatttttattcctaatcatgtaaatcttaaatttttctttttaaacatatgttaaatatttatttctcattatatataagaacatatttattaaatctagaattctatagtgagtcgtattacaattcactggccgtcgttttacaacgtcgtgactgggaaaaccctggcgttacccaacttaatcgccttgcagcacatccccctttcgccagctggcgtaatagcgaagaggcccgcaccgatcgcccttcccaacagttgcgcagcctgaatggcgaatggcgcctgatgcggtattttctccttacgcatctgtgcggtatttcacaccgcatatggtgcactctcagtacaatctgctctgatgccgcatagttaagccagccccgacacccgccaacacccgctgacgcgccctgacgggcttgtctgctcccggcatccgcttacagacaagctgtgaccgtctccgggagctgcatgtgtcagaggttttcaccgtcatcaccgaaacgcgcgagacgaaagggcctcgtgatacgcctatttttataggttaatgtcatgataataatggtttcttagacgtcaggtggcacttttcggggaaatgtgcgcggaacccctatttgtttatttttctaaatacattcaaatatgtatccgctcatgagacaataaccctgataaatgcttcaataatattgaaaaaggaagagtatgagtattcaacatttccgtgtcgcccttattcccttttttgcggcattttgccttcctgtttttgctcacccagaaacgctggtgaaagtaaaagatgctgaagatcagttgggtgcacgagtgggttacatcgaactggatctcaacagcggtaagatccttgagagttttcgccccgaagaacgttttccaatgatgagcacttttaaagttctgctatgtggcgcggtattatcccgtattgacgccgggcaagagcaactcggtcgccgcatacactattctcagaatgacttggttgagtactcaccagtcacagaaaagcatcttacggatggcatgacagtaagagaattatgcagtgctgccataaccatgagtgataacactgcggccaacttacttctgacaacgatcggaggaccgaaggagctaaccgcttttttgcacaacatgggggatcatgtaactcgccttgatcgttgggaaccggagctgaatgaagccataccaaacgacgagcgtgacaccacgatgcctgtagcaatgccaacaacgttgcgcaaactattaactggcgaactacttactctagcttcccggcaacaattaatagactggatggaggcggataaagttgcaggaccacttctgcgctcggcccttccggctggctggtttattgctgataaatctggagccggtgagcgtgggtctcgcggtatcattgcagcactggggccagatggtaagccctcccgtatcgtagttatctacacgacggggagtcaggcaactatggatgaacgaaatagacagatcgctgagataggtgcctcactgattaagcattggtaactgtcagaccaagtttactcatatatactttagattgatttaaaacttcatttttaatttaaaaggatctaggtgaagatcctttttgataatctcatgaccaaaatcccttaacgtgagttttcgttccactgagcgtcagaccccgtagaaaagatcaaaggatcttcttgagatcctttttttctgcgcgtaatctgctgcttgcaaacaaaaaaaccaccgctaccagcggtggtttgtttgccggatcaagagctaccaactctttttccgaaggtaactggcttcagcagagcgcagataccaaatactgtccttctagtgtagccgtagttaggccaccacttcaagaactctgtagcaccgcctacatacctcgctctgctaatcctgttaccagtggctgctgccagtggcgataagtcgtgtcttaccgggttggactcaagacgatagttaccggataaggcgcagcggtcgggctgaacggggggttcgtgcacacagcccagcttggagcgaacgacctacaccgaactgagatacctacagcgtgagctatgagaaagcgccacgcttcccgaagggagaaaggcggacaggtatccggtaagcggcagggtcggaacaggagagcgcacgagggagcttccagggggaaacgcctggtatctttatagtcctgtcgggtttcgccacctctgacttgagcgtcgatttttgtgatgctcgtcaggggggcggagcctatcgaaaaacgccagcaacgcggcctttttacggttcctggccttttgctggccttttgctcacatgttctttcctgcgttatcccctgattctgtggataaccgtattaccgcctttgagtgagctgataccgctcgccgcagccgaacgaccgagcgcagcgagtcagtgagcgaggaagcggaagagcgcccaatacgcaaaccgcctctccccgcgcgttggccgattcattaatgcagctggcacgacaggtttcccgactggaaagcgggcagtgagcgcaacgcaattaatgtgagttagctcactcattaggcaccccaggctttacactttatgcttccggctcgtatgttgtgtggaattgtgagcggataacaatttcacacaggaaacagctatgaccatgattacgccaagctatttaggtgacactatagaatactcgcggccgctaaTTAATAGAAGGAAAAGTAGAACAATGGAGAGTTATAATTGATTTAACATCATGTAGTGTTTTAAATATACCTATAGCAACTTTAAAGGATATCTCAAAAAATTTAAGTgtaagaaaagaaaaatatatgaacatatatatatatatatatatatatatatttatttatttatttatatttatttatttatatttcccatcagTGTAATTATCGATCAAGGCTGTGTAAAATGCTTGTGTTAAGTGCTCCTTTAGTTGTTACAGGAATATGGCATATGTTAAAATCCATCATACCAgtaattgaataaattatcatatatatattatttatgtatttatattttctaatttattcataaatatattttatatatgatactttaaaaatatgtagGTTGTAACACAACAAAAAATTACAATAACATCATCTGAAAAGGAGgtaaataaggaataaaaataaaataaaaaggaataatatatagtatatatttatatatataatagtaaaaatatatacctttatatatttttattttttattttattatattattttatttttttttttttctttctctctttatagAAGAAGTTGCTAGATCAAGTTCAGGCAAATCAATTAGAAAgtaagcatatattattattacatatgccaatatatttgtcttttaaacctgtacacacatatatatatatatatatatatatatatatttatttatttatttacacatgtttattttatgttttttttaacagAAAAATTCGGTGGAACCTGTGAGAATGCAACAGATTTTACAGAACCAATATTGCCATAAtttgttatatattcttttatgttaataattcgaataaaatatgttttatatgtataatttatatatatatatatattttttaatttttttatttgttttatattatgatcatataattattatatattcttttgttattatatatgcttcatttgattattgttatgttgctgttacataattccaagtttatttttctttatatcgcttgttgctgtatttgtatgcttatgcatacatacatatatatatatatatatatatatatattgtgtatagttttatatattttgtatacctcactatcttgtatgttataagtaaatatatatatttatatatatatgtttatttatttatttatttattaattttaatttaattttttttgtttacttttttttttttttttttttttttttttttttcaagttaaaaagccaaacaaaagaggtttatattttaaaacttctttatatatatatttatttattcataactttcatattttttaaaggaaaaagtataaattgtcaatttatttttcatattgtagaattagattcaaatatataatatatatatgtatggtatcataaatattttttttatatgtattatttaagatttataaacgtttaaaagttcgaaattatatatatatatttatatatatatttcaaaataatatacaaaagattgaaaatgttattattgagttaaaatatatatgtataattaaaatgtttcataagttatgaaatatataaattttataaaatggacatgtatcattcacttaaatatatattttataaatcttggtcataaaatatgtgatagacaattcttttcaagaaaattataacaaatccaaaaaagatatacgataaaaa

**5’ amplicon size: 1328bps**

**3’ amplicon size: 1145bps**

### > PF3D7_0911100 (PfSTART2)

attatgacaacctgagaacgttatacagaaatatgattatattatcatgttatataaaaatatatatatatatataaacttatttattatatatatataatatatattttattttttgtgttttatgaatatacatataattatataaacaacatacttatataaattcatagtcttaatgactacacacctaactatgcatgcatcaatatagtggattattaaaggaaccataaaaatatatatatatatatatatatatatatatatataataaataaataaaataaaaaaagaaaaaaaaattaaaaataaaaataaaaataaataaaacgcaaaagcacaaaaaatacacacacaaaaaataaaaaaagagaaaagaaaggaaatcacaaaaactaaaatatatatatatatatatatatatatatatatatatattatatatctatattttttttttttttttttttttttttctgagtacgaaaaatagagatagagaaaaaaaaaaaaaaATGAGTTGCGTACTAAAAAAAATTACAGTGAGTGAAGAGAACATCTGTTGTTACACAGGTGAAGAGGAAATATTAGAAAAAGAAAAAAAAAAGAATGCAAGAGATTTGTTAGTAGAGTCGAAAAAAGAGAAAAATAAAAAAATAGAACAAAAAAAACAACTACCACTAAGAGAAAAATTACCAACCTACTTTTCACAAAATTCATCTAATAGTTTGAAAAATTCTAATTTTGATAAAAAACAATTTTGTGAAAGTGGAACAAAATATTTAATAAAATATAAAATCTATAATTTGATAAAAAATCAATTTTATCAAGTTGATGTACATCATGCATCTGTTTCTCCTAGTTTGGATAAAAAAAAATTAAATTCCACAAAAAATATCATAAGAGAAAAAAACAAACAAACAAACAAAAAAAAAAAAAAATTACCAAAAATACCTGAAAAAAAAAAAAAAGATTTTCTAATAAATATTGCTACCAATTTTAGAAAAAATAAACCAAAAAAAAATTCCGAAAATTCTAATAAAAAAAAAAAAAAAAAAAATTACACCAATCAGCAAGCCAAAAAAGATAAAAGTAATAATAATATTATTAATAACAGTAATAGTAATATTAATAATAATAGTAACAGTAATAGTAATAGTAACAGTAATAATAATAATAATTACTATTATTATTATTATAATAATAAAAATAAAAATAAAAATAGTAATAATAATAATAATAATAATAATAATTTGAAAAATAAGAATAAAAATAACAAAAGGAATAGAAAAGGAAAAAAGAATATTAAAAAAAAAATTTACATAGAGAAAAATAGTCTGACTCTTCCAAAAACAAATAGATATTGTAATTATAAATATGAAGAATATTACGATATATCTGATGAAAAAATGGAAGCAACAAACTATGATGAAACATATTGTTATGATGAAGAACTAAAAAATGATAGTAAAACAACCATATATGATGAAGGTGTTTCAAAAAAAAATATATACAATTGTGATGTGAATCATATATTCGATAATTTTTGTAATTATTTTAATTGTGTAGATAAAAAAGAAAGTATGAACGATGACACAGCTAATTCAGAAATAACATTAAATTCTAATAATATAAAAACACAAGACATTATAATAGATAATAATGAAAAATCCTTTTTATATGAAAAAAATAGACAACAAAATATTAATGTATATACACAAACTTCTAATAAAAAATCTAGATGGTTATGTTCTACTGCAGATGATAGTCATAATGAATATTTAACAGAATATGAAAAAAAAATTGATAGTTATGACAATATTGATAACAATGTTCATAAGGATTATTCTAATATTCATAAAAATGTAACTATAGCAAATAATAACAATATAACATTAAATTACGATACAAATGATTTTTCTTCATATACAACACAACCCATTCAATTATTCAATGATCAAGAAAATAATACATATGACATGGGAACAAATATTGTAAGGAAAGGAAAAAGTTTATATGAAGAAACAAGAAAAATAAGTAATGCCCTTTTTGAAATGTACTCTACAAATGTAAATAAGGATAATATGAATAATAATATGAATGGCAATAATAATAATAATAATAATAATAACGAAAATGTTGATAAATTATGTAATAAAAACCAATTAGAAGCTCCTACCCCATCCCTATTTACAACATATATAAATCAATCTCATAATTATCATTCTATTGTTGATAAAAATATCGTATTAGTTCAAATTAAGAATTTTTCTCAAAAAATTGATGAATATATTTCTGATGAAAAAATATTCAGAGCACAGAAATTAATTGATCATGTTAAAAAATATATAGAATTTTATATAAACTATTATAAAAAATATAATGATAAAGAAGTTGTGGAGAAGTTAGAATTATATCATGAAATGCATTGTATGCATAAAAATATCAAATATAATATTAATAATTTAAAAGTTAATATTATTATGCATTTTTTAAATTTTTTTCATTTAAATGATATGTATACTATATTGAATGATTATACATATTACTTCTCAGTAGATATGGTGAATAGCATAACATCAGAAAGCATTACTAATTCAAACGAAAATACAAATTCTATAAATCATTCGGATATTAATAATATGAAAACTAAAAATTCTATACATTCATATGCTCCTTCCGAGTTAGGTGGACTATTTAATTCTTCTATTAAATGTTATAGTATTGACGAAAGTCTGAATTCTTCAAAAATGTATTTAACACCAACATTAGGTAATAATAATAATAATAATATTAATGTATCAAATAATGAAGCAGAAAATATCGAAAATGTACATAAAAGTGATAATCTAAATAATAATAATAATAATAATGAGGATGATGGTGTCAATGTAGATAGCGGTATAAATAATAACAGCAAATATAATACAGACGATTCAGATATTAAATATGAAGATATATCCAGTGTGATAGTAGATAAAATTTATCATCAACAATATAGTAATTGTAATATGCTTAATGGAAATAAAACTTTAATAAATCAAAAAAGTATAAATCCAGATAATTCAGCAAATAGAAAAAATAGTAATCGTCATATGGTTAAAATTATACAAAAATATTCTAAGAGATTTAAAAAATTCAGAAAAAGTAAGCAAAATAGTGAAGGGTGGATTAAAGAAAATGATAAATATCTTGATTTATCACATAGAGTAGATAAAGATAATAATATTTC**TGTACATATAAGAGCAAAACTACCATATGAAGTTAATCGAATCTTATCTATATTAAATGAAACAGAATTAAGTGTTAATTGGGCTCCATTCTTAACATCAGCAAAAAAAATTAAAAACTTGTCAAGAGCAAGTGCTATAATAACTCAGCTATATGAATATCCAATTATAGGAAAGAAAGAATCACTCATGTATTGTTTAG*gtaacaaaaacggataatatttatgcataaataaataaatataaatatatataaatatatatatatatatatattaatttttatgtgaacactattttggttttttatttaaatacagacgtttcatcacccgttttaatatataccataattatgtatatttatttatttatttatctatctatttattatttttttttttttttttcatcctattttag*GAGCGAATTCTTTGGAAGAACTCGGATGCATAATTTTGTGCTGCAAAGCCCCACCAGAATTTAATAAGGACATTTTATTTTATGAAAACATGTGCGAAAAAATAAATATAAACAAATTTGGGGAAATTATAAAAGTCAAAGAAATACCAATAAAATTTAGAAAGACATACAAAGAAGTAACATTTTTCGATTATACTTTACCTGAACCAGTACCAAAATTAGACAGACAAAGAGCAGCTAATTTATGTTTCCTCTTATATCCAATGAATAATGGAAAATCAACAGTTCTTGAATTATTTTTACATTTCGAAAATGAATTTAAATATACACCTATTAAAATGGTTACCTTCTTCATAAAGAAAATCGTAAAAAACATGTATGAAAATATTATAAAATCCTGTAGAAATTATGATCTTTTATATTCAGAATTCTTAATGAATAATGCCGAATTCTACATTTGGCTAGATGATCAAATTAAAAGATATATGAAAGGAAAAAATGATTCTAAACTTTTATACTCAATAAGTCTCGAAAGTTATGATGAACCAGAACATAATGAGGAACTGGATAGTAAGACT**TAG*GCACAAAAAAAAAAAAAAAAAATAAATAATAATAAATATATATATATATATATATGTATATATTATAACTATTAGATATAAACATAAATTTCAAAACAAATATATAATTAAATATCTTTAAAAAATATAAGACAAAATGTTATCCATTTTATTAAAATATGTTATATCGATATATCTATATTCAATTTGTCATCTTTTTACTTAATTAATTATATACAGTGTTTTTATATTTTATTTTGTTTATTTTCTTGTTAATATTTTTATTATATTCATATAATGTTCATTCTTTTAATTAAATTTTGTGCTTATTTATACATTGTTATTTTTTTTATTTTTTGTTTTGTTTTGTTTTGTTTTCTATATATAATATTCTTAGAATTAATTTTATATACATTTATATATAATATTATAGATACACATAGCATTATAACACAAAAATAATGTGTATATATTGCATGTTTTTATGTATATATAATATATATATATATATATATATTATAATGGCATACAAAAAAGAAAAAGAAAAATTGATATAAAGTAACCTTTTTTTATGTATAAATTATTTAAAGAAAGTTTATTTTATTTTAATTTATTTTATATTTATGTATTTATTTCATTTCATTTTTGAGTTTTCTAAACATAGGCCCTTTATAAAATTTACAATACATCATTGTGATATTAAATTTTTTCTCAAAAAATTGTTTGAAATTATATATATATATTATATATATGTATGTATTTTTTTATATATTTCCTTTTATTTATTTTATCTCTATTATTTCTTTATATTTAAATATAAATATTAAACCATTTCTATTAATACAGGTTTTATTAAAATTACGATTTGAATGTTCTGTAAATACATATTTTTTAATAAGAAAAAAGTAAAATGAAATAAGAATTGAATTGAATTGAATTTAATGAAGAATATAAATATAAAAATTTCCTTTTTTTGGAGTCGCACATATGAATATAATACACACTTGTATTATTAATATATATATATATATATGTAAATTATTTCGACGAATTTTATTTTTATTACGCATATATTTTTTTTCTTTAAAAATTAATAATATTTTCTTCTTTTTTTGTAAATGAAGGGAAAAAAAAAAAAAAAAAAAAAAAAAAGGAAAGGATAGGAAAATTACTTAAATTTTTATTTATTTATTTATTTATTATTATTATTTTATTTCTTTTTTTTTTTATACATATTTCAAATTTTCTTATAAATAATTTTTTCTCATTTTGTTTGCTAATATAATCGATTAGCTATATTTAAAAATATTATGTAATATTGAGGCTTTTTTTTTTTTCTTTTTTTTTTTTTTTTAATTTATTTTAAAGAAATTAATATATTTTAATTTTATGTAACGTTATTCCATATATTGTTTCCCTTTTTATTTATGAATTATTATAAATTTCTTTTATTTTAATTTTAATATATATATATATATATATATGTATGTGTATAAATTAAGGACAAGAAAAAAAA*

**Original Locus (OL) amplicon size: 1532bp**

### > PF3D7_0911100-GFP-Sandwich GFP Neo-R

attatgacaacctgagaacgttatacagaaatatgattatattatcatgttatataaaaatatatatatatatataaacttatttattatatatatataatatatattttattttttgtgttttatgaatatacatataattatataaacaacatacttatataaattcatagtcttaatgactacacacctaactatgcatgcatcaatatagtggattattaaaggaaccataaaaatatatatatatatatatatatatatatatatataataaataaataaaataaaaaaagaaaaaaaaattaaaaataaaaataaaaataaataaaacgcaaaagcacaaaaaatacacacacaaaaaataaaaaaagagaaaagaaaggaaatcacaaaaactaaaatatatatatatatatatatatatatatatatatatattatatatctatattttttttttttttttttttttttttctgagtacgaaaaatagagatagagaaaaaaaaaaaaaaATGAGTTGCGTACTAAAAAAAATTACAGTGAGTGAAGAGAACATCTGTTGTTACACAGGTGAAGAGGAAATATTAGAAAAAGAAAAAAAAAAGAATGCAAGAGATTTGTTAGTAGAGTCGAAAAAAGAGAAAAATAAAAAAATAGAACAAAAAAAACAACTACCACTAAGAGAAAAATTACCAACCTACTTTTCACAAAATTCATCTAATAGTTTGAAAAATTCTAATTTTGATAAAAAACAATTTTGTGAAAGTGGAACAAAATATTTAATAAAATATAAAATCTATAATTTGATAAAAAATCAATTTTATCAAGTTGATGTACATCATGCATCTGTTTCTCCTAGTTTGGATAAAAAAAAATTAAATTCCACAAAAAATATCATAAGAGAAAAAAACAAACAAACAAACAAAAAAAAAAAAAAATTACCAAAAATACCTGAAAAAAAAAAAAAAGATTTTCTAATAAATATTGCTACCAATTTTAGAAAAAATAAACCAAAAAAAAATTCCGAAAATTCTAATAAAAAAAAAAAAAAAAAAAATTACACCAATCAGCAAGCCAAAAAAGATAAAAGTAATAATAATATTATTAATAACAGTAATAGTAATATTAATAATAATAGTAACAGTAATAGTAATAGTAACAGTAATAATAATAATAATTACTATTATTATTATTATAATAATAAAAATAAAAATAAAAATAGTAATAATAATAATAATAATAATAATAATTTGAAAAATAAGAATAAAAATAACAAAAGGAATAGAAAAGGAAAAAAGAATATTAAAAAAAAAATTTACATAGAGAAAAATAGTCTGACTCTTCCAAAAACAAATAGATATTGTAATTATAAATATGAAGAATATTACGATATATCTGATGAAAAAATGGAAGCAACAAACTATGATGAAACATATTGTTATGATGAAGAACTAAAAAATGATAGTAAAACAACCATATATGATGAAGGTGTTTCAAAAAAAAATATATACAATTGTGATGTGAATCATATATTCGATAATTTTTGTAATTATTTTAATTGTGTAGATAAAAAAGAAAGTATGAACGATGACACAGCTAATTCAGAAATAACATTAAATTCTAATAATATAAAAACACAAGACATTATAATAGATAATAATGAAAAATCCTTTTTATATGAAAAAAATAGACAACAAAATATTAATGTATATACACAAACTTCTAATAAAAAATCTAGATGGTTATGTTCTACTGCAGATGATAGTCATAATGAATATTTAACAGAATATGAAAAAAAAATTGATAGTTATGACAATATTGATAACAATGTTCATAAGGATTATTCTAATATTCATAAAAATGTAACTATAGCAAATAATAACAATATAACATTAAATTACGATACAAATGATTTTTCTTCATATACAACACAACCCATTCAATTATTCAATGATCAAGAAAATAATACATATGACATGGGAACAAATATTGTAAGGAAAGGAAAAAGTTTATATGAAGAAACAAGAAAAATAAGTAATGCCCTTTTTGAAATGTACTCTACAAATGTAAATAAGGATAATATGAATAATAATATGAATGGCAATAATAATAATAATAATAATAATAACGAAAATGTTGATAAATTATGTAATAAAAACCAATTAGAAGCTCCTACCCCATCCCTATTTACAACATATATAAATCAATCTCATAATTATCATTCTATTGTTGATAAAAATATCGTATTAGTTCAAATTAAGAATTTTTCTCAAAAAATTGATGAATATATTTCTGATGAAAAAATATTCAGAGCACAGAAATTAATTGATCATGTTAAAAAATATATAGAATTTTATATAAACTATTATAAAAAATATAATGATAAAGAAGTTGTGGAGAAGTTAGAATTATATCATGAAATGCATTGTATGCATAAAAATATCAAATATAATATTAATAATTTAAAAGTTAATATTATTATGCATTTTTTAAATTTTTTTCATTTAAATGATATGTATACTATATTGAATGATTATACATATTACTTCTCAGTAGATATGGTGAATAGCATAACATCAGAAAGCATTACTAATTCAAACGAAAATACAAATTCTATAAATCATTCGGATATTAATAATATGAAAACTAAAAATTCTATACATTCATATGCTCCTTCCGAGTTAGGTGGACTATTTAATTCTTCTATTAAATGTTATAGTATTGACGAAAGTCTGAATTCTTCAAAAATGTATTTAACACCAACATTAGGTAATAATAATAATAATAATATTAATGTATCAAATAATGAAGCAGAAAATATCGAAAATGTACATAAAAGTGATAATCTAAATAATAATAATAATAATAATGAGGATGATGGTGTCAATGTAGATAGCGGTATAAATAATAACAGCAAATATAATACAGACGATTCAGATATTAAATATGAAGATATATCCAGTGTGATAGTAGATAAAATTTATCATCAACAATATAGTAATTGTAATATGCTTAATGGAAATAAAACTTTAATAAATCAAAAAAGTATAAATCCAGATAATTCAGCAAATAGAAAAAATAGTAATCGTCATATGGTTAAAATTATACAAAAATATTCTAAGAGATTTAAAAAATTCAGAAAAAGTAAGCAAAATAGTGAAGGGTGGATTAAAGAAAATGATAAATATCTTGATTTATCACATAGAGTAGATAAAGATAATAATATTTC**TGTACATATAAGAGCAAAACTACCATATGAAGTTAATCGAATCTTATCTATATTAAATGAAACAGAATTAAGTGTTAATTGGGCTCCATTCTTAACATCAGCAAAAAAAATTAAAAACTTGTCAAGAGCAAGTGCTATAATAACTCAGCTATATGAATATCCAATTATAGGAAAGAAAGAATCACTCATGTATTGTTTAG*gtaacaaaaacggataatatttatgcataaataaataaatataaatatatataaatatatatatatatatatattaatttttatgtgaacactattttggttttttatttaaatacagacgtttcatcacccgttttaatatataccataattatgtatatttatttatttatttatctatctatttattatttttttttttttttttcatcctattttag*GAGCGAATTCTTTGGAAGAACTCGGATGCATAATTTTGTGCTGCAAAGCCCCACCAGAATTTAATAAGGACATTTTATTTTATGAAAACATGTGCGAAAAAATAAATATAAACAAATTTGGGGAAATTATAAAAGTCAAAGAAATACCAATAAAATTTAGAAAGACATACAAAGAAGTAACATTTTTCGATTATACTTTACCTGAACCAGTACCAAAATTAGACAGACAAAGAGCAGCTAATTTATGTTTCCTCTTATATCCAATGAATAATGGAAAATCAACAGTTCTTGAATTATTTTTACATTTCGAAAATGAATTTAAATATACACCTATTAAAATGGTTACCTTCTTCATAAAGAAAATCGTAAAAAACATGTATGAAAATATTATAAAATCCTGTAGAAATTATGATCTTTTATATTCAGAATTCTTAATGAATAATGCCGAATTCTACATTTGGCTAGATGATCAAATTAAAAGATATATGAAAGGAAAAAATGATTCTAAACTTTTATACTCAATAAGTCTCGAAAGTTATGATGAACCAGAACATAATGAGGAACTGGATAGTAAGACT**CCTAGGTCAGGATTGAGATCAAGATCTGCTGCTGCTGGTGCTGGTGGTGCTGCTAGAGCTGCTCTGCAGAGAGGAGTACAAGTTGAAACAATATCACCAGGAGATGGTCGTACATTTCCAAAAAGAGGTCAAACTTGTGTTGTACATTATACTGGAATGCTTGAAGATGGAAAGAAATTTGATTCATCTCGTGATAGAAATAAACCATTTAAATTTATGCTAGGTAAACAAGAAGTAATACGAGGTTGGGAAGAAGGAGTTGCTCAAATGAGTGTAGGTCAAAGAGCAAAACTTACTATATCTCCAGATTATGCTTATGGTGCAACTGGACATCCAGGTATAATTCCACCTCATGCAACTCTTGTATTTGATGTGGAGCTTCTAAAACTAGAAACTAGAGGTGTTCAGGTTGAAACAATTTCACCTGGAGATGGCAGAACCTTTCCTAAAAGAGGACAGACTTGCGTAGTTCATTATACAGGCATGCTAGAGGATGGTAAGAAATTTGATTCTAGTCGAGATAGAAATAAGCCATTCAAGTTTATGCTAGGTAAACAGGAAGTAATAAGAGGTTGGGAAGAGGGTGTAGCACAGATGTCAGTTGGACAAAGAGCAAAGTTAACAATATCACCAGATTATGCATACGGTGCAACAGGCCATCCTGGCATCATCCCTCCACATGCAACTTTAGTATTCGACGTTGAATTGTTAAAGTTAGAGACAACGCGTGCTAGAGGTGCTGCTGCTGGTGCTGGAGGTGCAGGTAGACGTACGATGAGTAAAGGAGAAGAACTTTTCACTGGAGTTGTCCCAATTCTTGTTGAATTAGATGGTGATGTTAATGGGCACAAATTTTCTGTCAGTGGAGAGGGTGAAGGTGATGCAACATACGGAAAACTTACCCTTAAATTTATTTGCACTACTGGAAAACTACCTGTTCCATGGCCAACACTTGTCACTACTTTCGCGTATGGTCTTCAATGCTTTGCGAGATACCCAGATCATATGAAACAGCATGACTTTTTCAAGAGTGCCATGCCCGAAGGTTATGTACAGGAAAGAACTATATTTTTCAAAGATGACGGGAACTACAAGACACGTGCTGAAGTCAAGTTTGAAGGTGATACCCTTGTTAATAGAATCGAGTTAAAAGGTATTGATTTTAAAGAAGATGGAAACATTCTTGGACACAAATTGGAATACAACTATAACTCACACAATGTATACATCATGGCAGACAAACAAAAGAATGGAATCAAAGTTAACTTCAAAATTAGACACAACATTGAAGATGGAAGCGTTCAACTAGCAGACCATTATCAACAAAATACTCCAATTGGCGATGGCCCTGTCCTTTTACCAGACAACCATTACCTGTCCACACAATCTGCCCTTTCGAAAGATCCCAACGAAAAGAGAGACCACATGGTCCTTCTTGAGTTTGTAACAGCTGCTGGGATTACACATGGCATGGATGAGCTCTACAAAGTCGACGCCAGGGGAGCAGCCGCAGGAGCAGGGGGGGCAGGAAGGCGTGGTGTTCAGGTCGAGACTATTAGCCCTGGAGATGGACGCACGTTTCCTAAGCGTGGACAGACATGCGTAGTTCACTACACAGGTATGTTGGAGGACGGTAAAAAGTTCGACAGCTCACGCGACCGCAATAAACCTTTCAAGTTTATGCTTGGCAAGCAGGAGGTTATTCGTGGATGGGAGGAGGGTGTAGCACAGATGTCTGTTGGACAGCGTGCTAAGTTGACAATTTCACCTGACTATGCTTATGGCGCTACGGGCCATCCCGGGATCATTCCGCCACATGCGACTCTGGTATTCGACGTTGAATTATTAAAGTTAGAGACAGCTAGAGGGGCCGCTGCAGGTGCTGGTGGAGCTGGAAGACGTGGAGTACAAGTAGAGACTATCTCTCCAGGTGACGGTCGCACTTTCCCAAAGCGTGGCCAAACCTGTGTTGTACATTACACTGGTATGCTGGAGGATGGGAAAAAGTTCGATTCCAGTCGCGACCGTAACAAACCGTTCAAATTCATGTTGGGAAAGCAGGAAGTGATCCGCGGGTGGGAGGAAGGCGTGGCGCAAATGAGCGTCGGTCAGCGGGCTAAATTGACCATTTCCCCTGACTACGCGTATGGGGCTACTGGGCACCCAGGGATTATTCCGCCTCACGCTACACTTGTGTTTGATGTCGAACTTTTGAAACTGGAAACTGTCGACGGAGAAGGAAGAGGAAGTTTATTAACATGTGGAGATGTAGAAGAAAATCCAGGACCAATGATTGAACAAGATGGATTGCACGCAGGTTCTCCGGCCGCTTGGGTGGAGAGGCTATTCGGCTATGACTGGGCACAACAGACAATCGGCTGCTCTGATGCCGCCGTGTTCCGGCTGTCAGCGCAGGGGCGCCCGGTTCTTTTTGTCAAGACCGACCTGTCCGGTGCCCTGAATGAACTGCAGGACGAGGCAGCGCGGCTATCGTGGCTGGCCACGACGGGCGTTCCTTGCGCAGCTGTGCTCGACGTTGTCACTGAAGCGGGAAGGGACTGGCTGCTATTGGGCGAAGTGCCGGGGCAGGATCTCCTGTCATCTCACCTTGCTCCTGCCGAGAAAGTATCCATCATGGCTGATGCAATGCGGCGGCTGCATACGCTTGATCCGGCTACCTGCCCATTCGACCACCAAGCGAAACATCGCATCGAGCGAGCACGTACTCGGATGGAAGCCGGTCTTGTCGATCAGGATGATCTGGACGAAGAGCATCAGGGGCTCGCGCCAGCCGAACTGTTCGCCAGGCTCAAGGCGCGCATGCCCGACGGCGAGGATCTCGTCGTGACCCATGGCGATGCCTGCTTGCCGAATATCATGGTGGAAAATGGCCGCTTTTCTGGATTCATCGACTGTGGCCGGCTGGGTGTGGCGGACCGCTATCAGGACATAGCGTTGGCTACCCGTGATATTGCTGAAGAGCTTGGCGGCGAATGGGCTGACCGCTTCCTCGTGCTTTACGGTATCGCCGCTCCCGATTCGCAGCGCATCGCCTTCTATCGCCTTCTTGACGAGTTCTTCTAActcgagggatatggcagcttaatgttcgtttttcttatttatatatttataccaattgattgtatttataactgtaaaaatgtgtatgttgtgtgcatatttttttttgtgcatgcacatgcatgtaaatagctaaaattatgaacattttattttttgttcagaaaaaaaaaactttacacacataaaatggctagtatgaatagccatattttatataaattaaatcctatgaatttatgaccatattaaaaatttagatatttatggaacataatatgtttgaaacaataagacaaaattattattattattattatttttactgttataattatgtgtctccttcaatgattcataaatagttggacttgatttttaaaatgtttataatatgattagcatagttaaataaaaaaagttgaaaaattaaaaaaaaacatataaacacaaatgatggtttttccttcaatttcgatatcaatttatagaaacaaaatatatacttgtataattttatttttttatataaatcattacatatataattatacaatattttttctaagagataattatatattaatatatataaaaaaaggtgttttttttttttttttttatttttatttttattttatggtaatattttattttccttattttataaattatattagtttatatgtgattaattttatatattatcaatttatatatttttaaatgcttacttaattatctttttttttttttttttttttttttcccctctttttatattaatttatttttgaaaaaattgatatatatatatatatataatatatatatatacatgtagtagtattaaacaatgtataatatatataaataatatatttatatatttcatttcaattttaattttttttggttttttttttttttctttttgtcatatttaaaaaaaattatattcatataagttatgcattttttataaacattattcaatatatgtataatataatatatatatatatattaatgtattattccaatgtgcatgataaaagaaaaaaataatatttataaaaaaaaagaaaaataaaacaaaaaaagaaaaaaaaaaaaaaaaaaaaaaaaatacaaaaataaataatataatttataattatatattcttgtcacaataaaaatatatatatatatatatatatttataatatgtatattttaaactagaaaaggaataactaatattttatttattatcattcaagatttatattttataataataaatacctaatagaaatatatcaggatccatgcatggttcgctaaactgcatcgtcgctgtgtcccagaacatgggcatcggcaagaacggggactacccctggccaccgctcaggaacgaatttagatatttccagagaatgaccacaacctcttcagtagaaggtaaacagaatctggtgattatgggtaagaagacctggttctccattcctgagaagaatcgacctttaaagggtagaattaatttagttctcagcagagaactcaaggaacctccacaaggagctcattttctttccagaagtctagatgatgccttaaaacttactgaacaaccagaattagcaaataaagtagacatggtctggatagttggtggcagttctgtttataaggaagccatgaatcacccaggccatcttaaactatttgtgacaaggatcatgcaagactttgaaagtgacacgttttttccagaaattgatttggagaaatataaacttctgccagaatacccaggtgttctctctgatgtccaggaggagaaaggcattaagtacaaatttgaagtatatgagaagaatgattaagcttatttaataatagattaaaaatattataaaaataaaaacataaacacagaaattacaaaaaaaatacatatgaattttttttttgtaatcttccttataaatatagaataatgaatcatataaaacatatcattattcatttatttacatttaaaattattgtttcagtatctttaatttattatgtatatataaaaataacttacaattttattaataaacaatatatgtttattaattcatgttttgtaatttatgggatagcgattttttttactgtctgtatttttcttttttaattatgttttaattgtattttatttttattattgttctttttatagtattattttaaaacaaaatgtattttctaagaacttataataataataaatataaattttaataaaaattatatttatcttttacaatatgaacataaagtacaacattaatatatagcttttaatatttttattcctaatcatgtaaatcttaaatttttctttttaaacatatgttaaatatttatttctcattatatataagaacatatttattaaatctagaattctatagtgagtcgtattacaattcactggccgtcgttttacaacgtcgtgactgggaaaaccctggcgttacccaacttaatcgccttgcagcacatccccctttcgccagctggcgtaatagcgaagaggcccgcaccgatcgcccttcccaacagttgcgcagcctgaatggcgaatggcgcctgatgcggtattttctccttacgcatctgtgcggtatttcacaccgcatatggtgcactctcagtacaatctgctctgatgccgcatagttaagccagccccgacacccgccaacacccgctgacgcgccctgacgggcttgtctgctcccggcatccgcttacagacaagctgtgaccgtctccgggagctgcatgtgtcagaggttttcaccgtcatcaccgaaacgcgcgagacgaaagggcctcgtgatacgcctatttttataggttaatgtcatgataataatggtttcttagacgtcaggtggcacttttcggggaaatgtgcgcggaacccctatttgtttatttttctaaatacattcaaatatgtatccgctcatgagacaataaccctgataaatgcttcaataatattgaaaaaggaagagtatgagtattcaacatttccgtgtcgcccttattcccttttttgcggcattttgccttcctgtttttgctcacccagaaacgctggtgaaagtaaaagatgctgaagatcagttgggtgcacgagtgggttacatcgaactggatctcaacagcggtaagatccttgagagttttcgccccgaagaacgttttccaatgatgagcacttttaaagttctgctatgtggcgcggtattatcccgtattgacgccgggcaagagcaactcggtcgccgcatacactattctcagaatgacttggttgagtactcaccagtcacagaaaagcatcttacggatggcatgacagtaagagaattatgcagtgctgccataaccatgagtgataacactgcggccaacttacttctgacaacgatcggaggaccgaaggagctaaccgcttttttgcacaacatgggggatcatgtaactcgccttgatcgttgggaaccggagctgaatgaagccataccaaacgacgagcgtgacaccacgatgcctgtagcaatgccaacaacgttgcgcaaactattaactggcgaactacttactctagcttcccggcaacaattaatagactggatggaggcggataaagttgcaggaccacttctgcgctcggcccttccggctggctggtttattgctgataaatctggagccggtgagcgtgggtctcgcggtatcattgcagcactggggccagatggtaagccctcccgtatcgtagttatctacacgacggggagtcaggcaactatggatgaacgaaatagacagatcgctgagataggtgcctcactgattaagcattggtaactgtcagaccaagtttactcatatatactttagattgatttaaaacttcatttttaatttaaaaggatctaggtgaagatcctttttgataatctcatgaccaaaatcccttaacgtgagttttcgttccactgagcgtcagaccccgtagaaaagatcaaaggatcttcttgagatcctttttttctgcgcgtaatctgctgcttgcaaacaaaaaaaccaccgctaccagcggtggtttgtttgccggatcaagagctaccaactctttttccgaaggtaactggcttcagcagagcgcagataccaaatactgtccttctagtgtagccgtagttaggccaccacttcaagaactctgtagcaccgcctacatacctcgctctgctaatcctgttaccagtggctgctgccagtggcgataagtcgtgtcttaccgggttggactcaagacgatagttaccggataaggcgcagcggtcgggctgaacggggggttcgtgcacacagcccagcttggagcgaacgacctacaccgaactgagatacctacagcgtgagctatgagaaagcgccacgcttcccgaagggagaaaggcggacaggtatccggtaagcggcagggtcggaacaggagagcgcacgagggagcttccagggggaaacgcctggtatctttatagtcctgtcgggtttcgccacctctgacttgagcgtcgatttttgtgatgctcgtcaggggggcggagcctatcgaaaaacgccagcaacgcggcctttttacggttcctggccttttgctggccttttgctcacatgttctttcctgcgttatcccctgattctgtggataaccgtattaccgcctttgagtgagctgataccgctcgccgcagccgaacgaccgagcgcagcgagtcagtgagcgaggaagcggaagagcgcccaatacgcaaaccgcctctccccgcgcgttggccgattcattaatgcagctggcacgacaggtttcccgactggaaagcgggcagtgagcgcaacgcaattaatgtgagttagctcactcattaggcaccccaggctttacactttatgcttccggctcgtatgttgtgtggaattgtgagcggataacaatttcacacaggaaacagctatgaccatgattacgccaagctatttaggtgacactatagaatactcgcggccgcTAGtgtacatataagagcaaaactaccatatgaagttaatcgaatcttatctatattaaatgaaacagaattaagtgttaattgggctccattcttaacatcagcaaaaaaaattaaaaacttgtcaagagcaagtgctataataactcagctatatgaatatccaattataggaaagaaagaatcactcatgtattgtttaggtaacaaaaacggataatatttatgcataaataaataaatataaatatatataaatatatatatatatatatattaatttttatgtgaacactattttggttttttatttaaatacagacgtttcatcacccgttttaatatataccataattatgtatatttatttatttatttatctatctatttattatttttttttttttttttcatcctattttaggagcgaattctttggaagaactcggatgcataattttgtgctgcaaagccccaccagaatttaataaggacattttattttatgaaaacatgtgcgaaaaaataaatataaacaaatttggggaaattataaaagtcaaagaaataccaataaaatttagaaagacatacaaagaagtaacatttttcgattatactttacctgaaccagtaccaaaattagacagacaaagagcagctaatttatgtttcctcttatatccaatgaataatggaaaatcaacagttcttgaattatttttacatttcgaaaatgaatttaaatatacacctattaaaatggttaccttcttcataaagaaaatcgtaaaaaacatgtatgaaaatattataaaatcctgtagaaattatgatcttttatattcagaattcttaatgaataatgccgaattctacatttggctagatgatcaaattaaaagatatatgaaaggaaaaaatgattctaaacttttatactcaataagtctcgaaagttatgatgaaccagaacataatgaggaactggatagtaagactTAG*GCACAAAAAAAAAAAAAAAAAATAAATAATAATAAATATATATATATATATATATGTATATATTATAACTATTAGATATAAACATAAATTTCAAAACAAATATATAATTAAATATCTTTAAAAAATATAAGACAAAATGTTATCCATTTTATTAAAATATGTTATATCGATATATCTATATTCAATTTGTCATCTTTTTACTTAATTAATTATATACAGTGTTTTTATATTTTATTTTGTTTATTTTCTTGTTAATATTTTTATTATATTCATATAATGTTCATTCTTTTAATTAAATTTTGTGCTTATTTATACATTGTTATTTTTTTTATTTTTTGTTTTGTTTTGTTTTGTTTTCTATATATAATATTCTTAGAATTAATTTTATATACATTTATATATAATATTATAGATACACATAGCATTATAACACAAAAATAATGTGTATATATTGCATGTTTTTATGTATATATAATATATATATATATATATATATTATAATGGCATACAAAAAAGAAAAAGAAAAATTGATATAAAGTAACCTTTTTTTATGTATAAATTATTTAAAGAAAGTTTATTTTATTTTAATTTATTTTATATTTATGTATTTATTTCATTTCATTTTTGAGTTTTCTAAACATAGGCCCTTTATAAAATTTACAATACATCATTGTGATATTAAATTTTTTCTCAAAAAATTGTTTGAAATTATATATATATATTATATATATGTATGTATTTTTTTATATATTTCCTTTTATTTATTTTATCTCTATTATTTCTTTATATTTAAATATAAATATTAAACCATTTCTATTAATACAGGTTTTATTAAAATTACGATTTGAATGTTCTGTAAATACATATTTTTTAATAAGAAAAAAGTAAAATGAAATAAGAATTGAATTGAATTGAATTTAATGAAGAATATAAATATAAAAATTTCCTTTTTTTGGAGTCGCACATATGAATATAATACACACTTGTATTATTAATATATATATATATATATGTAAATTATTTCGACGAATTTTATTTTTATTACGCATATATTTTTTTTCTTTAAAAATTAATAATATTTTCTTCTTTTTTTGTAAATGAAGGGAAAAAAAAAAAAAAAAAAAAAAAAAAGGAAAGGATAGGAAAATTACTTAAATTTTTATTTATTTATTTATTTATTATTATTATTTTATTTCTTTTTTTTTTTATACATATTTCAAATTTTCTTATAAATAATTTTTTCTCATTTTGTTTGCTAATATAATCGATTAGCTATATTTAAAAATATTATGTAATATTGAGGCTTTTTTTTTTTTCTTTTTTTTTTTTTTTTAATTTATTTTAAAGAAATTAATATATTTTAATTTTATGTAACGTTATTCCATATATTGTTTCCCTTTTTATTTATGAATTATTATAAATTTCTTTTATTTTAATTTTAATATATATATATATATATATATGTATGTGTATAAATTAAGGACAAGAAAAAAAA*

**5’ amplicon size: 1216bps**

**3’ amplicon size: 1545bps**

### > PF3D7_1463500 (PfSTART3)

ctttttaagaatatattttattttgaataagaattattattaatttatatttaaaaaattttcttcttattttttattaatacttattttatgttatattattttattttattttttattttttgggttatatttagaacgtttacttaacaaattttagattttattataatagaaatgtcttaacttttccttatgcaaaataagtgttataaatatatttatgtatatatatatttattcatatatatatttttatttaatatatatatatatatatatatatatatatatatatatatttaatatttattttgtgtccttctttgcatgcgcaatataatacttgttatttttttatattttaataaaatcctttacatgaatatttttcatattttaaaacaaataaatttatattataataattatgtattatttagtaaataaataagtaacataaaaatatataaaggggaatttatctatggaatatatatattatatataaaaacaagaatattataaagaaatactcaaacaaaggaattaaaagaatgatactatataaaattgaatattaaaaaaaatataatcataaatataaatacaaatataatatttatatataatgatttatatatattgttatataatatatatatgtaataaacatataggagctatataaacattattttagcataaagatatcttcaatttaggaatatgcaattttcagattaattctttgatcattttttattttattattttaaaagtttctttttcctttatcagaaatacagctatctataatatatttttatatattccaatatatatatatatatatatatatatatatatatatacaaatacatatacatattttattatatttatattacatatgtatatatttttcttctttctatataattattttttttggtatacatcaaatgcaagtacatatttcaaaATGAGTAATTCAAATGAAAATAACGAATTGGACATAGACGATCGTTTAAAGTCAATGGAACATTTGGTTTGTAAAGATGAGAAAGAAATAATGAAAGTTAATGAAATTATAGAAGAAGCTTCAAATGTGTTATATAACTTTTCAATAAAGCAAGATGATTATTACAAATATAGTACGATTGATGAAGACTCTCATTTGTATTTTAAGAAAGTTAACAACACTGATGTGGGAAAAATTGATTTACTCTTTCAGGATCCATCAAAA*gtagacttataaatagaaaaattatattctattaaaaaaaaaaaaaaaaaaaagatgtaatatgtaataatttggtacatatatatatatatatattatattatattattatgagaaaaaaaaaaaaaatagacagaaatttattattaacatacatgatccatattaataacttttttatttttatccttcttacaataaaaaaaatataaatgtttgtatcatttttttaatttttgttag*TTT**GAAAAAATTATACAAACCATTTGGGACGAAAATGGCACAAGAAAATTTGATCCATATTTTATAGAAG*gttctaaaagaataaaaaaaaaaataaaataaaatatatatatatatatatatatgtgatataatttgatattttaatgttatattattattattatatacttatatatttgatttttttgtaattttttctcaag*GCAAAATATTGCGCATATATGACAAGGATACAATTTTAGTGAGACAAAGTTACAAAGGAACTTTAGGAAAAGAAGGAAGATATTTTTATATTGTAGCTCAAAAAAAGAAG*gtagaataaataattcatattattattttattttattttattttttgattctcatataaaatttataataattttatttattatacaatttatataatatatattatttcattataatatatataaatatatttcttaattttgtag*CTAAATAATAATACGTATCTTATAACTTGTGTGTCCTTAAATATCAATGATAATAATGAAAAGTGCACAAGTAATTTTATTAATCCATTTATACATAGCGCTAATTCCTTTACCCTAAACATTGAATGCGATGAAAATATTAAGAATTCGTCATTAAAAAGGATGTATATTAATTTATCTGGTTATCATATTAAACGTGAAGATAACTGTATAAAATTTACTTATGTATCTTCT*gtaagtttataattatattaaattatatatatatatatatttctaaaattattccttaataataaaattttttttttgtttttttttttctctctttttatag*ATTGAACTAGATACGTCTCCTTTGATTCCCCAATTTATTATAAGAAAAGTCAAAGCGAAGAAAATGTTGCAGTTAAATACCTTAAGACAAAGTATT**TGAtaaatcgataaaatatatgctacatgaataaataaataaataaatataaatatatatatatatatatatatatatatatataagatatatttattcatatgtatataaaaactttatatatttttttttctcattatttacttttaatataaatgggaacatatatatatatatatatatatatatttataacattcctgaaaccaaacgtatctgctaaatatatcattatgatatttatgataatatttggtacatatcttcacttatatgtacttacattttttttttttttaccttttcgagttatataaaacttgaatttatatatttatattattatatacattttgtatatatatatatatatatatatatatatatatatatatatatatatatatatatatttgttatatgacgttttttattgaaaaatgtcttttttcaaaaatgatacaaaaacagtttttttttttttttttaatgaacaattataatatatacatattaatatatatatatatatatatatattgttataattttaataaaacagaaaaaaaagaaaaaattagaaacatttattgcaaatatgaatacaatacttacaattattatgtcatta

**Original Locus (OL) amplicon size: 1396bps**

### > PF3D7_1463500-GFP-Sandwich GFP Neo-R

ctttttaagaatatattttattttgaataagaattattattaatttatatttaaaaaattttcttcttattttttattaatacttattttatgttatattattttattttattttttattttttgggttatatttagaacgtttacttaacaaattttagattttattataatagaaatgtcttaacttttccttatgcaaaataagtgttataaatatatttatgtatatatatatttattcatatatatatttttatttaatatatatatatatatatatatatatatatatatatatatttaatatttattttgtgtccttctttgcatgcgcaatataatacttgttatttttttatattttaataaaatcctttacatgaatatttttcatattttaaaacaaataaatttatattataataattatgtattatttagtaaataaataagtaacataaaaatatataaaggggaatttatctatggaatatatatattatatataaaaacaagaatattataaagaaatactcaaacaaaggaattaaaagaatgatactatataaaattgaatattaaaaaaaatataatcataaatataaatacaaatataatatttatatataatgatttatatatattgttatataatatatatatgtaataaacatataggagctatataaacattattttagcataaagatatcttcaatttaggaatatgcaattttcagattaattctttgatcattttttattttattattttaaaagtttctttttcctttatcagaaatacagctatctataatatatttttatatattccaatatatatatatatatatatatatatatatatatatacaaatacatatacatattttattatatttatattacatatgtatatatttttcttctttctatataattattttttttggtatacatcaaatgcaagtacatatttcaaaATGAGTAATTCAAATGAAAATAACGAATTGGACATAGACGATCGTTTAAAGTCAATGGAACATTTGGTTTGTAAAGATGAGAAAGAAATAATGAAAGTTAATGAAATTATAGAAGAAGCTTCAAATGTGTTATATAACTTTTCAATAAAGCAAGATGATTATTACAAATATAGTACGATTGATGAAGACTCTCATTTGTATTTTAAGAAAGTTAACAACACTGATGTGGGAAAAATTGATTTACTCTTTCAGGATCCATCAAAA*gtagacttataaatagaaaaattatattctattaaaaaaaaaaaaaaaaaaaagatgtaatatgtaataatttggtacatatatatatatatatattatattatattattatgagaaaaaaaaaaaaaatagacagaaatttattattaacatacatgatccatattaataacttttttatttttatccttcttacaataaaaaaaatataaatgtttgtatcatttttttaatttttgttag*TTT**GAAAAAATTATACAAACCATTTGGGACGAAAATGGCACAAGAAAATTTGATCCATATTTTATAGAAG*gttctaaaagaataaaaaaaaaaataaaataaaatatatatatatatatatatatgtgatataatttgatattttaatgttatattattattattatatacttatatatttgatttttttgtaattttttctcaag*GCAAAATATTGCGCATATATGACAAGGATACAATTTTAGTGAGACAAAGTTACAAAGGAACTTTAGGAAAAGAAGGAAGATATTTTTATATTGTAGCTCAAAAAAAGAAG*gtagaataaataattcatattattattttattttattttattttttgattctcatataaaatttataataattttatttattatacaatttatataatatatattatttcattataatatatataaatatatttcttaattttgtag*CTAAATAATAATACGTATCTTATAACTTGTGTGTCCTTAAATATCAATGATAATAATGAAAAGTGCACAAGTAATTTTATTAATCCATTTATACATAGCGCTAATTCCTTTACCCTAAACATTGAATGCGATGAAAATATTAAGAATTCGTCATTAAAAAGGATGTATATTAATTTATCTGGTTATCATATTAAACGTGAAGATAACTGTATAAAATTTACTTATGTATCTTCT*gtaagtttataattatattaaattatatatatatatatatttctaaaattattccttaataataaaattttttttttgtttttttttttctctctttttatag*ATTGAACTAGATACGTCTCCTTTGATTCCCCAATTTATTATAAGAAAAGTCAAAGCGAAGAAAATGTTGCAGTTAAATACCTTAAGACAAAGTATT**CCTAGGTCAGGATTGAGATCAAGATCTGCTGCTGCTGGTGCTGGTGGTGCTGCTAGAGCTGCTCTGCAGAGAGGAGTACAAGTTGAAACAATATCACCAGGAGATGGTCGTACATTTCCAAAAAGAGGTCAAACTTGTGTTGTACATTATACTGGAATGCTTGAAGATGGAAAGAAATTTGATTCATCTCGTGATAGAAATAAACCATTTAAATTTATGCTAGGTAAACAAGAAGTAATACGAGGTTGGGAAGAAGGAGTTGCTCAAATGAGTGTAGGTCAAAGAGCAAAACTTACTATATCTCCAGATTATGCTTATGGTGCAACTGGACATCCAGGTATAATTCCACCTCATGCAACTCTTGTATTTGATGTGGAGCTTCTAAAACTAGAAACTAGAGGTGTTCAGGTTGAAACAATTTCACCTGGAGATGGCAGAACCTTTCCTAAAAGAGGACAGACTTGCGTAGTTCATTATACAGGCATGCTAGAGGATGGTAAGAAATTTGATTCTAGTCGAGATAGAAATAAGCCATTCAAGTTTATGCTAGGTAAACAGGAAGTAATAAGAGGTTGGGAAGAGGGTGTAGCACAGATGTCAGTTGGACAAAGAGCAAAGTTAACAATATCACCAGATTATGCATACGGTGCAACAGGCCATCCTGGCATCATCCCTCCACATGCAACTTTAGTATTCGACGTTGAATTGTTAAAGTTAGAGACAACGCGTGCTAGAGGTGCTGCTGCTGGTGCTGGAGGTGCAGGTAGACGTACGATGAGTAAAGGAGAAGAACTTTTCACTGGAGTTGTCCCAATTCTTGTTGAATTAGATGGTGATGTTAATGGGCACAAATTTTCTGTCAGTGGAGAGGGTGAAGGTGATGCAACATACGGAAAACTTACCCTTAAATTTATTTGCACTACTGGAAAACTACCTGTTCCATGGCCAACACTTGTCACTACTTTCGCGTATGGTCTTCAATGCTTTGCGAGATACCCAGATCATATGAAACAGCATGACTTTTTCAAGAGTGCCATGCCCGAAGGTTATGTACAGGAAAGAACTATATTTTTCAAAGATGACGGGAACTACAAGACACGTGCTGAAGTCAAGTTTGAAGGTGATACCCTTGTTAATAGAATCGAGTTAAAAGGTATTGATTTTAAAGAAGATGGAAACATTCTTGGACACAAATTGGAATACAACTATAACTCACACAATGTATACATCATGGCAGACAAACAAAAGAATGGAATCAAAGTTAACTTCAAAATTAGACACAACATTGAAGATGGAAGCGTTCAACTAGCAGACCATTATCAACAAAATACTCCAATTGGCGATGGCCCTGTCCTTTTACCAGACAACCATTACCTGTCCACACAATCTGCCCTTTCGAAAGATCCCAACGAAAAGAGAGACCACATGGTCCTTCTTGAGTTTGTAACAGCTGCTGGGATTACACATGGCATGGATGAGCTCTACAAAGTCGACGCCAGGGGAGCAGCCGCAGGAGCAGGGGGGGCAGGAAGGCGTGGTGTTCAGGTCGAGACTATTAGCCCTGGAGATGGACGCACGTTTCCTAAGCGTGGACAGACATGCGTAGTTCACTACACAGGTATGTTGGAGGACGGTAAAAAGTTCGACAGCTCACGCGACCGCAATAAACCTTTCAAGTTTATGCTTGGCAAGCAGGAGGTTATTCGTGGATGGGAGGAGGGTGTAGCACAGATGTCTGTTGGACAGCGTGCTAAGTTGACAATTTCACCTGACTATGCTTATGGCGCTACGGGCCATCCCGGGATCATTCCGCCACATGCGACTCTGGTATTCGACGTTGAATTATTAAAGTTAGAGACAGCTAGAGGGGCCGCTGCAGGTGCTGGTGGAGCTGGAAGACGTGGAGTACAAGTAGAGACTATCTCTCCAGGTGACGGTCGCACTTTCCCAAAGCGTGGCCAAACCTGTGTTGTACATTACACTGGTATGCTGGAGGATGGGAAAAAGTTCGATTCCAGTCGCGACCGTAACAAACCGTTCAAATTCATGTTGGGAAAGCAGGAAGTGATCCGCGGGTGGGAGGAAGGCGTGGCGCAAATGAGCGTCGGTCAGCGGGCTAAATTGACCATTTCCCCTGACTACGCGTATGGGGCTACTGGGCACCCAGGGATTATTCCGCCTCACGCTACACTTGTGTTTGATGTCGAACTTTTGAAACTGGAAACTGTCGACGGAGAAGGAAGAGGAAGTTTATTAACATGTGGAGATGTAGAAGAAAATCCAGGACCAATGATTGAACAAGATGGATTGCACGCAGGTTCTCCGGCCGCTTGGGTGGAGAGGCTATTCGGCTATGACTGGGCACAACAGACAATCGGCTGCTCTGATGCCGCCGTGTTCCGGCTGTCAGCGCAGGGGCGCCCGGTTCTTTTTGTCAAGACCGACCTGTCCGGTGCCCTGAATGAACTGCAGGACGAGGCAGCGCGGCTATCGTGGCTGGCCACGACGGGCGTTCCTTGCGCAGCTGTGCTCGACGTTGTCACTGAAGCGGGAAGGGACTGGCTGCTATTGGGCGAAGTGCCGGGGCAGGATCTCCTGTCATCTCACCTTGCTCCTGCCGAGAAAGTATCCATCATGGCTGATGCAATGCGGCGGCTGCATACGCTTGATCCGGCTACCTGCCCATTCGACCACCAAGCGAAACATCGCATCGAGCGAGCACGTACTCGGATGGAAGCCGGTCTTGTCGATCAGGATGATCTGGACGAAGAGCATCAGGGGCTCGCGCCAGCCGAACTGTTCGCCAGGCTCAAGGCGCGCATGCCCGACGGCGAGGATCTCGTCGTGACCCATGGCGATGCCTGCTTGCCGAATATCATGGTGGAAAATGGCCGCTTTTCTGGATTCATCGACTGTGGCCGGCTGGGTGTGGCGGACCGCTATCAGGACATAGCGTTGGCTACCCGTGATATTGCTGAAGAGCTTGGCGGCGAATGGGCTGACCGCTTCCTCGTGCTTTACGGTATCGCCGCTCCCGATTCGCAGCGCATCGCCTTCTATCGCCTTCTTGACGAGTTCTTCTAActcgagggatatggcagcttaatgttcgtttttcttatttatatatttataccaattgattgtatttataactgtaaaaatgtgtatgttgtgtgcatatttttttttgtgcatgcacatgcatgtaaatagctaaaattatgaacattttattttttgttcagaaaaaaaaaactttacacacataaaatggctagtatgaatagccatattttatataaattaaatcctatgaatttatgaccatattaaaaatttagatatttatggaacataatatgtttgaaacaataagacaaaattattattattattattatttttactgttataattatgtgtctccttcaatgattcataaatagttggacttgatttttaaaatgtttataatatgattagcatagttaaataaaaaaagttgaaaaattaaaaaaaaacatataaacacaaatgatggtttttccttcaatttcgatatcaatttatagaaacaaaatatatacttgtataattttatttttttatataaatcattacatatataattatacaatattttttctaagagataattatatattaatatatataaaaaaaggtgttttttttttttttttttatttttatttttattttatggtaatattttattttccttattttataaattatattagtttatatgtgattaattttatatattatcaatttatatatttttaaatgcttacttaattatctttttttttttttttttttttttttcccctctttttatattaatttatttttgaaaaaattgatatatatatatatatataatatatatatatacatgtagtagtattaaacaatgtataatatatataaataatatatttatatatttcatttcaattttaattttttttggttttttttttttttctttttgtcatatttaaaaaaaattatattcatataagttatgcattttttataaacattattcaatatatgtataatataatatatatatatatattaatgtattattccaatgtgcatgataaaagaaaaaaataatatttataaaaaaaaagaaaaataaaacaaaaaaagaaaaaaaaaaaaaaaaaaaaaaaaatacaaaaataaataatataatttataattatatattcttgtcacaataaaaatatatatatatatatatatatttataatatgtatattttaaactagaaaaggaataactaatattttatttattatcattcaagatttatattttataataataaatacctaatagaaatatatcaggatccatgcatggttcgctaaactgcatcgtcgctgtgtcccagaacatgggcatcggcaagaacggggactacccctggccaccgctcaggaacgaatttagatatttccagagaatgaccacaacctcttcagtagaaggtaaacagaatctggtgattatgggtaagaagacctggttctccattcctgagaagaatcgacctttaaagggtagaattaatttagttctcagcagagaactcaaggaacctccacaaggagctcattttctttccagaagtctagatgatgccttaaaacttactgaacaaccagaattagcaaataaagtagacatggtctggatagttggtggcagttctgtttataaggaagccatgaatcacccaggccatcttaaactatttgtgacaaggatcatgcaagactttgaaagtgacacgttttttccagaaattgatttggagaaatataaacttctgccagaatacccaggtgttctctctgatgtccaggaggagaaaggcattaagtacaaatttgaagtatatgagaagaatgattaagcttatttaataatagattaaaaatattataaaaataaaaacataaacacagaaattacaaaaaaaatacatatgaattttttttttgtaatcttccttataaatatagaataatgaatcatataaaacatatcattattcatttatttacatttaaaattattgtttcagtatctttaatttattatgtatatataaaaataacttacaattttattaataaacaatatatgtttattaattcatgttttgtaatttatgggatagcgattttttttactgtctgtatttttcttttttaattatgttttaattgtattttatttttattattgttctttttatagtattattttaaaacaaaatgtattttctaagaacttataataataataaatataaattttaataaaaattatatttatcttttacaatatgaacataaagtacaacattaatatatagcttttaatatttttattcctaatcatgtaaatcttaaatttttctttttaaacatatgttaaatatttatttctcattatatataagaacatatttattaaatctagaattctatagtgagtcgtattacaattcactggccgtcgttttacaacgtcgtgactgggaaaaccctggcgttacccaacttaatcgccttgcagcacatccccctttcgccagctggcgtaatagcgaagaggcccgcaccgatcgcccttcccaacagttgcgcagcctgaatggcgaatggcgcctgatgcggtattttctccttacgcatctgtgcggtatttcacaccgcatatggtgcactctcagtacaatctgctctgatgccgcatagttaagccagccccgacacccgccaacacccgctgacgcgccctgacgggcttgtctgctcccggcatccgcttacagacaagctgtgaccgtctccgggagctgcatgtgtcagaggttttcaccgtcatcaccgaaacgcgcgagacgaaagggcctcgtgatacgcctatttttataggttaatgtcatgataataatggtttcttagacgtcaggtggcacttttcggggaaatgtgcgcggaacccctatttgtttatttttctaaatacattcaaatatgtatccgctcatgagacaataaccctgataaatgcttcaataatattgaaaaaggaagagtatgagtattcaacatttccgtgtcgcccttattcccttttttgcggcattttgccttcctgtttttgctcacccagaaacgctggtgaaagtaaaagatgctgaagatcagttgggtgcacgagtgggttacatcgaactggatctcaacagcggtaagatccttgagagttttcgccccgaagaacgttttccaatgatgagcacttttaaagttctgctatgtggcgcggtattatcccgtattgacgccgggcaagagcaactcggtcgccgcatacactattctcagaatgacttggttgagtactcaccagtcacagaaaagcatcttacggatggcatgacagtaagagaattatgcagtgctgccataaccatgagtgataacactgcggccaacttacttctgacaacgatcggaggaccgaaggagctaaccgcttttttgcacaacatgggggatcatgtaactcgccttgatcgttgggaaccggagctgaatgaagccataccaaacgacgagcgtgacaccacgatgcctgtagcaatgccaacaacgttgcgcaaactattaactggcgaactacttactctagcttcccggcaacaattaatagactggatggaggcggataaagttgcaggaccacttctgcgctcggcccttccggctggctggtttattgctgataaatctggagccggtgagcgtgggtctcgcggtatcattgcagcactggggccagatggtaagccctcccgtatcgtagttatctacacgacggggagtcaggcaactatggatgaacgaaatagacagatcgctgagataggtgcctcactgattaagcattggtaactgtcagaccaagtttactcatatatactttagattgatttaaaacttcatttttaatttaaaaggatctaggtgaagatcctttttgataatctcatgaccaaaatcccttaacgtgagttttcgttccactgagcgtcagaccccgtagaaaagatcaaaggatcttcttgagatcctttttttctgcgcgtaatctgctgcttgcaaacaaaaaaaccaccgctaccagcggtggtttgtttgccggatcaagagctaccaactctttttccgaaggtaactggcttcagcagagcgcagataccaaatactgtccttctagtgtagccgtagttaggccaccacttcaagaactctgtagcaccgcctacatacctcgctctgctaatcctgttaccagtggctgctgccagtggcgataagtcgtgtcttaccgggttggactcaagacgatagttaccggataaggcgcagcggtcgggctgaacggggggttcgtgcacacagcccagcttggagcgaacgacctacaccgaactgagatacctacagcgtgagctatgagaaagcgccacgcttcccgaagggagaaaggcggacaggtatccggtaagcggcagggtcggaacaggagagcgcacgagggagcttccagggggaaacgcctggtatctttatagtcctgtcgggtttcgccacctctgacttgagcgtcgatttttgtgatgctcgtcaggggggcggagcctatcgaaaaacgccagcaacgcggcctttttacggttcctggccttttgctggccttttgctcacatgttctttcctgcgttatcccctgattctgtggataaccgtattaccgcctttgagtgagctgataccgctcgccgcagccgaacgaccgagcgcagcgagtcagtgagcgaggaagcggaagagcgcccaatacgcaaaccgcctctccccgcgcgttggccgattcattaatgcagctggcacgacaggtttcccgactggaaagcgggcagtgagcgcaacgcaattaatgtgagttagctcactcattaggcaccccaggctttacactttatgcttccggctcgtatgttgtgtggaattgtgagcggataacaatttcacacaggaaacagctatgaccatgattacgccaagctatttaggtgacactatagaatactcgcggccgcgaaaaaattatacaaaccatttgggacgaaaatggcacaagaaaatttgatccatattttatagaaggttctaaaagaataaaaaaaaaaataaaataaaatatatatatatatatatatatgtgatataatttgatattttaatgttatattattattattatatacttatatatttgatttttttgtaattttttctcaaggcaaaatattgcgcatatatgacaaggatacaattttagtgagacaaagttacaaaggaactttaggaaaagaaggaagatatttttatattgtagctcaaaaaaagaaggtagaataaataattcatattattattttattttattttattttttgattctcatataaaatttataataattttatttattatacaatttatataatatatattatttcattataatatatataaatatatttcttaattttgtagctaaataataatacgtatcttataacttgtgtgtccttaaatatcaatgataataatgaaaagtgcacaagtaattttattaatccatttatacatagcgctaattcctttaccctaaacattgaatgcgatgaaaatattaagaattcgtcattaaaaaggatgtatattaatttatctggttatcatattaaacgtgaagataactgtataaaatttacttatgtatcttctgtaagtttataattatattaaattatatatatatatatatttctaaaattattccttaataataaaattttttttttgtttttttttttctctctttttatagattgaactagatacgtctcctttgattccccaatttattataagaaaagtcaaagcgaagaaaatgttgcagttaaataccttaagacaaagtattTGAtaaatcgataaaatatatgctacatgaataaataaataaataaatataaatatatatatatatatatatatatatatatataagatatatttattcatatgtatataaaaactttatatatttttttttctcattatttacttttaatataaatgggaacatatatatatatatatatatatatatttataacattcctgaaaccaaacgtatctgctaaatatatcattatgatatttatgataatatttggtacatatcttcacttatatgtacttacattttttttttttttaccttttcgagttatataaaacttgaatttatatatttatattattatatacattttgtatatatatatatatatatatatatatatatatatatatatatatatatatatatatttgttatatgacgttttttattgaaaaatgtcttttttcaaaaatgatacaaaaacagtttttttttttttttttaatgaacaattataatatatacatattaatatatatatatatatatatatattgttataattttaataaaacagaaaaaaaagaaaaaattagaaacatttattgcaaatatgaatacaatacttacaattattatgtcatta

**5’ amplicon size: 1298bps**

**3’ amplicon size: 1218bps**

### > PF3D7_0214800 (PfTUPA)

cattttattccgtaaaattaaaatgaattccaccaaaggtttaacatatacataatatatatttatatatatatatatatgtatatatatatatataaaataaaaataataatatataatatatttttatattacattattatattatattatattatattatattttattttatttttttaatatagaacttcatgtagctaaaataaaaatgagttcacatttttcgttaattaaatatattatatatatatatatatgcaaaattggtattatatataaatataatatataatttgtttccttttttttagtttttaaaatatagatttataaataatttttttcttttttttaagaattattttgatttaccatatatattatttaaaatcccaagttaaaattaaaaaaaaaaaggagtttttattataatattatatatatttatatatatattttaactactttaattttaaaagtgtgagagtaagaaaatgaaaaaaaaaaaaaaaaaaaaaaaaaggattattttttatttatatattataatatatataatatattaataataaataaaataaacatgtaaataaatattatatatatatatcataatatagtatcatgaaataatatatatttataaaaatattattataatatatagttaattattatattatatattacataatatttataatatattatatattggattatagttttacctattgggggatcttggttctataaaaaaatttcaatttttttttaccctttatttttttttgttcactatatatatatattgttatatataacataatatattaatagtttcataatattatatgtatatagaaagaaaaacatatatatatatatatatatattgttcattatttctaaatatatatattattatatatattatataaaaaatatattattttagatttactcctttatatataaattaatattttttatagatttcgattttttcctatttacataatttttaattttttataattttaatttttctacattatatatgtattatatatatatataaataatctatatatttttaaaataattcaatgtcattttattaattcaattaaatacaaattaatttcttattttctttaattatatatatatatattataatatttatagaacctatattatagatttaatttttaatctgttaaaagaaaaaaagtacaaaaaattatacatttatttacatatatatatatatatatatatatatatgtataatttttaatcttttatttttagaggtattttatgttattttttattttacatatgttgttgtataattttttttgtatatgcccttatattgattttcttttttttttttaaataatatagtgtttgctttaataaaaaaaaggaaatatcaggattttattgaatacatacatatatatatatatatatatatatatatatatatatatatatatatataactattagaattcccatagatgaatatataaacatataataatattatatctataatatttatatttttattatacatacatactgaatatattttaaagttacgaataatgaatgctttttttttttttttttttttgttgctttaaataaatggtaatcataaatgaaattagtctagagctaaaatgaaaattaaagaaaatataaaaagtatataataataaatgtgttataatatttaaaataaaacaaaaaaaaaaaaaaaaaaaaaaaaaaaagtaaattacaataatttattctataaagttcagaATG**TTGGAATTATTACGTAATGCTTATAATAAATGTAAAGAGTATATTTTGGGCTTATCGAAAAAATCACAAATATATAAAATAAATATATTAAATTATTGTTTATTATTTTCGTTTTTTTTATTTTATGCATCATATTTTTTTTTTATAAATGGAATAATAAATCCAGCAGTAAAAATTGTTTTCCATGGAA*gtaagtaccttatatgaaaaaatgaaagtgtatccataaaagaagatatacacaaaggtgtatatatatatatatatatatatatatatatatatatgtatgtatgtatgtatttatttatttatttattatttatatatatattttctttatattttctttaag*GTATTATTGAGGACAAAACCAAAGTTTACGAGAACTCGATAATTGAATCAATATGGCTATTGTTTACCAAGGGAAGTTATTTTAATATGGCCATGCTTCTATTCTTTTCTATTATAATACCGTTTCTCAAGTTACTTATGGTTAGTGATAACTTTTATAGTTTTATCGTATTATATAAAATGAATAAAAAACGAGAAGAGGAAAAGAGAAGGAGACGAAGAAGAATACGTGCACGAGAATATAACAATGCTAAATATAACAAGAATAGATATAAAATGAATAAATATAATAGAAATAAGAATATTAATGATATATACAAGGATGATATATATTATAGTGAAAATATTTTTAAGAATGATGAGATAAATTATATTAATACAATAAATGAGAATGAAGAATTTATTTTAAAGAAATTTAAAATTTTAAATTTCATATCGCGTTTTCAATTTGTTGATGTATTTATATCTTTATTTATAGTATCATCATTAAATTTATATTTATTAGAGGCAAGGATGTTAAATGGTGC**ATATTATTTTTTGAATTATTGTATGTTATCAACAATATCATCTTTTTTGTTATTTAGTTTTACTTCTTTAAAAATACATATATTTAAAAATGGAAATATTAAGATATCTGCATGTCTCAATGAATCAAATCTAGAGGTTACTACAAGCGGTCCCCTTTCAACAAAAGATCTTGTCGAAGAAGAAGGACATGCACAAATAAATAATCTAATTATTAATGATAAGAATATGACTAGTGGTGTTGTTAATGATTTTTCAGGTAATGGAAACAATGTCGAATTAACAGACGACCTTAAGAGTGGCGAACCCCAAAACAGAGATGATATACAAACTGAAGAAACAAAAAAAGAAAAAATGAATACTAGAACTCATAATGATGAAGATAATACAAAAAAAAAAAATATAAAGGATAAAAAGAAAGCTAACGGTGATGATAAAGTAATACAAAAATGTATTGATAATGAAAGGAAAAAAAAACAAAATGGTATGATTCAAAGTGTAAACGATGGTGATAAAAATAGTAATTTTAATAATAACAATAATAATAATATTAATGGTGATAGTAATAATAATAATATTAATGGTGATAGTAATAATAATAATATTAATGGTGATAGTAATAATAATAATATTAATGGTGATAGTAATAATAATAATATTAATGGTGATAGTAATAATAATAATATTAATGGTGATAGTAATAATAATAATTATCATAATAATTATCATAATAATTATCGTAATAATTATCATAATAATTATCGTAATAATAATTGTAGGAATAATATTTTAGAACAAAATAAATGTGATAAAAATGTTTTGTGTTATAATAATATATATAATACAATGAAAGATAATGATACTTATATATATTTAAAAAAGAATAAATTTAATTCGTTATTAAAAAGTAATTGTATCAAAACTAATTTTAATATGATAAAGATAGGTTATGTAATATTTTTATTTGTGTTATTATGTTTATGTATATATTTAATAACAGGAGTAGAATGCAGTTTATTTGGTATATATATATATTTGAGTTATTTTAATTTTAATATTGAAGGAATATTAATTGATTATATGGATATGTTAAATATTTTAAAATTAAAAATAAAGAAAGGATATATCTATCCTTTTTTTGTTATGTTACCATTTATTTTCCCTGTAATAATATCTATGTGTTTCTTTTTAAGTGTATTTTTTTTAAATATGTATTATGAAAGCTTTTCAAAGTTATATAAAAAGATCAGTGAACTAAAGAATGAATTCATAAACTCCTCAGAAAATGACAATGTGAACGAAAGAATATTGGTATCCGAAACTTCTAATCATTTATGTTTAAATGAAAGTAACGATAAAGTATCTAATACATCAGATGATTTTTTATCAAGAAATAATAGCAACATATCTAGTTCAAAAAGTGAAATGATAAATTCAAATTTTGTATTTAATAAACTTTTGAATTTTTATTTTTCATTTGCAGTTTTCTTTTCTTATTTAGGTTCTGCATTTTTACATATATCCTTAGGTGAAATTATATGTATCGCATTATTAACATTTTATCAAATTGTAAAGCATACCAATAATTTGAATATCACAATTTTACTTAAGAGTGAGAAAATTAAATTTTGTAAATTCCTTTTGTTTATATTATATGGGTTGTTATGCTTTTCCATAAATTTGTATGTAAACCAATGGGAGGAATATATAACGAAATTAAAAAGGTTAAAAAGACGTATTTTATTGTTTGAAAAAAATAAATTTAGTGAGATTGTGGATCTGAATACTCAAAAGGGTGATGGTGACCATTTTGATGAAACACAAATATTTTCCATATTTTTTTCTTTTTTAATAAAAAAAAATGAGGGTTCAAAAATGAGGGATAATGATATGAATAGTGATAGCGAAGACAGTATATATGATGCTTATGAGCAGCAGATACAATTACACCATGGTGATAATATGGTCAATGGTATGTTGATGATGAGAAGGATATCGATGCAAAATTTAGAGGACGACGAAACCCAAGTTGAATATATAAATAGGGAGATCCATACGCAAGGTGATTTACATGTACGAAGAACGAATCAAGGTATTTTAAGATTTAATATGAGAAGGGGCAAAAAAGGGTCCAATGAAAATATGGGTGTCCACCATGAGAGTGGTAATGTGGATGATGCGAATGGTATGAATAATGTGGATGATACGAATAATATGAATAATGTGGATGGTACGAATAATATGAATAATGTGGATGGTACGAATAATATGAATAATATGGATGGTAGGAATAATATGAATAATATAAATAGTGTGGATAATATGAACAATTTAAATAATAATGATGGTGAAGAAGAAGAAGAATGTGTGAACGATGTTTTAAATTATGATAATAATAATTATGCTATTAATGAAGATGCTGAGGAATATATAAAAAATACGTCAGGAGAACGAGCTGTTATTATTTGTTCTGAAAAAAGAATATATGAAAAGAACGGAAATGGTGATATAATAACTAGAAATTATAAAAATGAAGAACGATATATATATTTAAAAAAATGGATTCCTTTTAAATCTATGATATTGTCAAAATTAGAAAAGAGGAAAAGGAATAGAAAAGAGGCATATAATACTCCTAGAGTATTAATATTAATACATTCCTTTTTATTTATACTTATTGTTTTCATATTTTTAATGGTATTCTTTAAAAAAGAGCCTATTTTTCGTTTTAATATGCCATCAGTTAATAAGCGTTTAAATAATTTTTTTAAATCAACAAGTTTTCACGAAATTATACCAAATAGTGTAGGGAAATGTAAAACAAAAAAATATATAGCAAAAGAACCATGTTTTAATGTGGGTCATATATACCATGAAGAGAAGACATTTTATCATGCAACTCTTTTATTTTTACAAGGTTTAAGATCTGTAAAAATTATGAATATGAATTTTTATTATGAAAAAGGCATATATTATTTATCTTTGGATGGATATTTTAAACATATTATAGGTCCTCTCTTTCTTAAATTATGTTTAGGTACAAACTTTTGTCCCATAAGTACATATGCATTTTTAGTAGGAAGTAAACCAACTTTCTCAGTAAATGTTGCTGTTCAATGTAATAATAAAAAACCACCATATTATATGACGGATATTATTGTAAAAGATTTAAAAATTACAAAAATAGAAATAGTTAAACATTCAGATGTTATCGATAATGTGGATATTAAATTAGATGATGTACAAGATAGAGTACAAGAAAAGGTTAATGCCATGTTAGAAGCAAAAAAAAAAATTATCGTCTGGAAAAATCAAAAGTATCATTTGGAGGGTTTTCTCAATTATTTAATATCTAAAAATGCATTATCTGGGTTTTCCTGTGAACCCATAAATTATTGAacttgacaaattaaaacacgttaaatgtatatatgtatatatattatatatgtatatatttatttatatttttcttacatttccatttgggtgaacatatattcttacccttttcataaattgttttttttttttttgatatttatcaataatatattcagtaaaaatatattatttcacataatttaatattttttggaataacatataaatgtcaatgtatatgtgtatatatatatatatatatatatatgtatatatttttttttctttttcatatatttatatgcgcccatttgtttttatctatatatatatatatttatatatccgaatttctttatattgttatattccgatttgattatatataaaattttttttaattcataatttttttttttttttttttcttttttttcatttccatgtgatttatttcaatccgatattatttatatgttatatatatattattctataatatgataaaaggatatttttttttcttatttgtatttttagattataaatataaagatattcttatatatagattatatctgcttttttttttttttttttttttttttctattattaatttttaaattatctacttacgaattttttaattgttgatatggccttcatgagtatattattatgcatgaaagaaaaaaaaaaaaaaaaatatatatatgtatataatatatatatatatatatataatataccaatttttattatcttatattttacatgcactataaatgcatagaatttttcttttttttttttttgatttcttctccccaaatttttggttatttatataatatacgtattataaatatgtacatataatatgtatatattataatgtgtaattttccacatatatatataaaatatatatatatatatatatatattttaataatacttgaacttttaaaattaaatatataaaaaggaatccaattaatttatggttatattatttgttattcactagaaaattctctagaatttaataaaacatatat

**Original Locus (OL) amplicon size: 1055bp**

### > GFP-2xFKBP-PF3D7_0214800 GFP yDHODH recodonizedsequence

Similar integration and PCR products for the S850N mutant.

cattttattccgtaaaattaaaatgaattccaccaaaggtttaacatatacataatatatatttatatatatatatatatgtatatatatatatataaaataaaaataataatatataatatatttttatattacattattatattatattatattatattatattttattttatttttttaatatagaacttcatgtagctaaaataaaaatgagttcacatttttcgttaattaaatatattatatatatatatatatgcaaaattggtattatatataaatataatatataatttgtttccttttttttagtttttaaaatatagatttataaataatttttttcttttttttaagaattattttgatttaccatatatattatttaaaatcccaagttaaaattaaaaaaaaaaaggagtttttattataatattatatatatttatatatatattttaactactttaattttaaaagtgtgagagtaagaaaatgaaaaaaaaaaaaaaaaaaaaaaaaaggattattttttatttatatattataatatatataatatattaataataaataaaataaacatgtaaataaatattatatatatatatcataatatagtatcatgaaataatatatatttataaaaatattattataatatatagttaattattatattatatattacataatatttataatatattatatattggattatagttttacctattgggggatcttggttctataaaaaaatttcaatttttttttaccctttatttttttttgttcactatatatatatattgttatatataacataatatattaatagtttcataatattatatgtatatagaaagaaaaacatatatatatatatatatatattgttcattatttctaaatatatatattattatatatattatataaaaaatatattattttagatttactcctttatatataaattaatattttttatagatttcgattttttcctatttacataatttttaattttttataattttaatttttctacattatatatgtattatatatatatataaataatctatatatttttaaaataattcaatgtcattttattaattcaattaaatacaaattaatttcttattttctttaattatatatatatatattataatatttatagaacctatattatagatttaatttttaatctgttaaaagaaaaaaagtacaaaaaattatacatttatttacatatatatatatatatatatatatatatgtataatttttaatcttttatttttagaggtattttatgttattttttattttacatatgttgttgtataattttttttgtatatgcccttatattgattttcttttttttttttaaataatatagtgtttgctttaataaaaaaaaggaaatatcaggattttattgaatacatacatatatatatatatatatatatatatatatatatatatatatatatataactattagaattcccatagatgaatatataaacatataataatattatatctataatatttatatttttattatacatacatactgaatatattttaaagttacgaataatgaatgctttttttttttttttttttttgttgctttaaataaatggtaatcataaatgaaattagtctagagctaaaatgaaaattaaagaaaatataaaaagtatataataataaatgtgttataatatttaaaataaaacaaaaaaaaaaaaaaaaaaaaaaaaaaaagtaaattacaataatttattctataaagttcagaATG**TTGGAATTATTACGTAATGCTTATAATAAATGTAAAGAGTATATTTTGGGCTTATCGAAAAAATCACAAATATATAAAATAAATATATTAAATTATTGTTTATTATTTTCGTTTTTTTTATTTTATGCATCATATTTTTTTTTTATAAATGGAATAATAAATCCAGCAGTAAAAATTGTTTTCCATGGAA*gtaagtaccttatatgaaaaaatgaaagtgtatccataaaagaagatatacacaaaggtgtatatatatatatatatatatatatatatatatatatgtatgtatgtatgtatttatttatttatttattatttatatatatattttctttatattttctttaag*GTATTATTGAGGACAAAACCAAAGTTTACGAGAACTCGATAATTGAATCAATATGGCTATTGTTTACCAAGGGAAGTTATTTTAATATGGCCATGCTTCTATTCTTTTCTATTATAATACCGTTTCTCAAGTTACTTATGGTTAGTGATAACTTTTATAGTTTTATCGTATTATATAAAATGAATAAAAAACGAGAAGAGGAAAAGAGAAGGAGACGAAGAAGAATACGTGCACGAGAATATAACAATGCTAAATATAACAAGAATAGATATAAAATGAATAAATATAATAGAAATAAGAATATTAATGATATATACAAGGATGATATATATTATAGTGAAAATATTTTTAAGAATGATGAGATAAATTATATTAATACAATAAATGAGAATGAAGAATTTATTTTAAAGAAATTTAAAATTTTAAATTTCATATCGCGTTTTCAATTTGTTGATGTATTTATATCTTTATTTATAGTATCATCATTAAATTTATATTTATTAGAGGCAAGGATGTTAAATGGTGC**AGGTTTAAACGAGCAGAAGTTAATATCAGAAGAGGATTTGGGTGAACAAAAACTCATAAGCGAAGAAGATTTAATAACTTCGTATAGCATACATTATACGAAGTTATCCGGAGAAGGAAGAGGAAGTTTATTAACATGTGGAGATGTAGAAGAAAATCCAGGACCAATGACAGCCAGTTTAACTACCAAGTTCTTGAACAATACCTATGAAAACCCATTTATGAATGCATCCGGTGTTCATTGCATGACTACACAAGAATTAGATGAATTAGCAAACTCTAAAGCTGGCGCATTCATTACAAAGAGTGCTACAACCTTAGAAAGAGAAGGTAACCCTGAACCACGTTACATTTCTGTCCCTCTAGGCAGTATCAACTCCATGGGTTTACCAAACGAAGGTATCGACTACTATTTGTCCTATGTATTAAACCGTCAAAAGAATTATCCTGATGCACCTGCTATTTTCTTCTCAGTTGCTGGTATGAGCATTGATGAAAATTTAAATTTGTTGAGGAAAATCCAAGATAGCGAATTCAACGGTATTACCGAGTTAAACTTGTCTTGTCCTAATGTGCCTGGGAAACCACAAGTTGCTTATGACTTTGACTTGACAAAGGAAACCTTGGAAAAGGTTTTTGCCTTTTTCAAAAAACCTCTTGGTGTCAAGTTGCCTCCTTATTTTGATTTTGCCCATTTTGATATCATGGCAAAAATATTGAACGAGTTCCCATTAGCTTATGTCAACTCTATCAATAGTATAGGAAATGGTCTTTTCATTGATGTGGAGAAGGAGAGTGTAGTAGTGAAGCCAAAGAATGGTTTCGGGGGTATTGGAGGTGAATATGTTAAGCCAACCGCGCTCGCCAATGTTCGTGCATTTTACACTCGTTTGAGACCTGAAATCAAAGTTATCGGTACAGGTGGAATTAAGTCCGGTAAGGATGCATTTGAACATCTTCTATGTGGTGCCTCTATGCTACAGATTGGTACAGAATTACAAAAAGAGGGCGTCAAGATTTTTGAACGTATCGAAAAAGAATTAAAAGACATAATGGAAGCTAAGGGTTATACATCCATAGATCAGTTCCGTGGGAAGTTGAACAGCATTGGTGAAGGTAGAGGTTCTTTGTTGACTTGTGGTGATGTTGAAGAAAATCCAGGTCCAGCTAGCATGAGTAAAGGAGAAGAACTTTTCACTGGAGTTGTCCCAATTCTTGTTGAATTAGATGGTGATGTTAATGGGCACAAATTTTCTGTCAGTGGAGAGGGTGAAGGTGATGCAACATACGGAAAACTTACCCTTAAATTTATTTGCACTACTGGAAAACTACCAGTTCCATGGCCAACACTTGTCACTACTTTCGCGTATGGTCTTCAATGCTTTGCGAGATACCCAGATCATATGAAACAGCATGACTTTTTCAAGAGTGCCATGCCCGAAGGTTATGTACAGGAAAGAACTATATTTTTCAAAGATGACGGGAACTACAAGACACGTGCTGAAGTCAAGTTTGAAGGTGATACCCTTGTTAATAGAATCGAGTTAAAAGGTATTGATTTTAAAGAAGATGGAAACATTCTTGGACACAAATTGGAATACAACTATAACTCACACAATGTATACATCATGGCAGACAAACAAAAGAATGGAATCAAAGTTAACTTCAAAATTAGACACAACATTGAAGATGGAAGCGTTCAACTAGCAGACCATTATCAACAAAATACTCCAATTGGCGATGGCCCTGTCCTTTTACCAGACAACCATTACCTGTCCACACAATCTGCCCTTTCGAAAGATCCCAACGAAAAGAGAGACCACATGGTCCTTCTTGAGTTTGTAACAGCTGCTGGGATTACACATGGCATGGATGAGCTCTACAAACCTAGCTCAGGATTGAGATCAAGATCTGCTGCTGCTGGTGCTGGTGGTGCTGCTAGAGCTGCTCTGCAGAGAGGAGTACAAGTTGAAACAATATCACCAGGAGATGGTCGTACATTTCCAAAAAGAGGTCAAACTTGTGTTGTACATTATACTGGAATGCTTGAAGATGGAAAGAAATTTGATTCATCTCGTGATAGAAATAAACCATTTAAATTTATGCTAGGTAAACAAGAAGTAATACGAGGTTGGGAAGAAGGAGTTGCTCAAATGAGTGTAGGTCAAAGAGCAAAACTTACTATATCTCCAGATTATGCTTATGGTGCAACTGGACATCCAGGTATAATTCCACCTCATGCAACTCTTGTATTTGATGTGGAGCTTCTAAAACTAGAAACTAGAGGTGTTCAGGTTGAAACAATTTCACCTGGAGATGGCAGAACCTTTCCTAAAAGAGGACAGACTTGCGTAGTTCATTATACAGGCATGCTAGAGGATGGTAAGAAATTTGATTCTAGTCGAGATAGAAATAAGCCATTCAAGTTTATGCTAGGTAAACAGGAAGTAATAAGAGGTTGGGAAGAGGGTGTAGCACAGATGTCAGTTGGACAAAGAGCAAAGTTAACAATATCACCAGATTATGCATACGGTGCAACAGGCCATCCTGGCATCATCCCTCCACATGCAACTTTAGTATTCGACGTTGAATTGTTAAAGTTAGAGACAACGCGTGCTAGAGGTGCTGCTGCTGGTGCTGGAGGTGCAGGTAGACCTAGGATGCTTGAGCTTCTTAGAAACGCATACAACAAGTGCAAGGAATACATACTTGGACTTAGTAAGAAGAGTCAGATTTACAAGATTAACATTCTTAACTACTGCCTTCTTTTCAGTTTCTTCCTTTTCTACGCTAGTTACTTCTTCTTCATTAACGGTATTATTAACCCTGCTGTTAAGATAGTATTTCACGGTTCAATAATAGAAGATAAGACAAAGGTATATGAAAATAGTATTATAGAGAGTATTTGGTTGCTTTTCACAAAAGGTTCATACTTCAACATGGCAATGTTATTGTTTTTCAGCATAATTATTCCATTCTTAAAACTTTTAATGGTATCAGACAATTTCTACTCATTCATAGTTCTTTACAAGATGAACAAGAAGAGGGAGGAAGAGAAACGTCGTCGTAGGCGTCGTATTAGAGCTAGGGAGTACAATAACGCAAAGTACAATAAAAACCGTTACAAGATGAACAAGTACAACCGTAACAAAAACATAAACGACATTTATAAAGACGACATTTACTACTCAGAGAACATATTCAAAAACGACGAAATTAACTACATAAACACTATTAACGAAAACGAGGAGTTCATACTTAAAAAGTTCAAGATACTTAACTTTATTAGTAGATTCCAGTTCGTAGACGTTTTCATTAGCCTTTTCATTGTTAGTAGTCTTAACCTTTACCTTCTTGAAGCTCGTATGCTTAACGGAGCTTACTACTTCCTTAACTACTGCATGCTTAGTACTATTAGTAGCTTCCTTCTTTTCTCATTCACAAGCCTTAAGATTCACATTTTCAAGAACGGTAACATAAAAATTAGCGCTTGCTTAAACGAGAGTAACTTGGAAGTAACAACTTCAGGACCATTAAGTACTAAGGACTTAGTTGAGGAGGAGGGTCACGCTCAGATTAACAACTTGATAATAAACGACAAAAACATGACATCAGGAGTAGTAAACGACTTCAGTGGAAACGGTAATAACGTTGAGCTTACTGATGATTTAAAATCAGGAGAGCCACAGAATCGTGACGACATTCAGACAGAGGAGACTAAGAAGGAGAAGATGAACACACGTACACACAACGACGAGGACAACACTAAGAAGAAGAACATTAAAGACAAGAAAAAGGCAAATGGAGACGACAAGGTTATTCAGAAGTGCATAGACAACGAGCGTAAGAAGAAGCAGAACGGAATGATACAGTCAGTTAATGACGGAGACAAGAACTCAAACTTCAACAACAATAACAACAACAACATAAACGGAGACTCAAACAACAACAACATAAACGGAGACTCAAACAACAACAACATAAACGGAGACTCAAACAACAACAACATAAACGGAGACTCAAACAACAACAACATAAACGGAGACTCAAACAACAACAACATAAACGGAGACTCAAACAACAACAACTACCACAACAACTACCACAACAACTACAGAAACAACTACCACAACAACTACAGAAACAACAACTGCCGTAACAACATACTTGAGCAGAACAAGTGCGACAAGAACGTACTTTGCTACAACAACATTTACAACACTATGAAGGACAACGACACATACATTTACCTTAAGAAAAACAAGTTCAACAGTCTTCTTAAGTCAAACTGCATAAAGACAAACTTCAACATGATTAAAATTGGATACGTTATTTTCCTTTTCGTTCTTCTTTGCCTTTGCATTTACCTTATTACTGGTGTTGAGTGTTCACTTTTCGGAATTTACATTTACCTTTCATACTTCAACTTCAACATAGAGGGTATTCTTATAGACTACATGGACATGCTTAACATACTTAAGCTTAAGATTAAAAAGGGTTACATATACCCATTCTTCGTAATGCTTCCTTTCATATTTCCAGTTATTATTAGCATGTGCTTTTTCCTTTCAGTTTTCTTCCTTAACATGTACTACGAGTCATTCAGTAAACTTTACAAGAAAATATCAGAGTTGAAAAACGAGTTTATTAATAGTAGTGAGAACGATAACGTTAATGAGCGTATTCTTGTTAGTGAGACAAGCAACCACCTTTGCCTTAACGAGTCAAATGACAAGGTTAGCAACACTAGTGACGACTTCCTTAGTCGTAACAACTCAAATATTAGCTCAAGTAAGTCAGAGATGATTAACAGTAACTTCGTTTTCAACAAGTTACTTAACTTCTACTTCAGTTTCGCTGTATTTTTCAGCTACCTTGGAAGCGCTTTCCTTCACATTAGTCTTGGAGAGATAATTTGCATAGCTCTTCTTACTTTCTACCAGATAGTTAAACACACAAACAACCTTAACATAACTATACTTTTAAAATCAGAAAAGATAAAGTTCTGCAAGTTTTTACTTTTCATTCTTTACGGACTTCTTTGTTTCAGTATTAACCTTTACGTTAATCAGTGGGAAGAGTACATTACAAAGCTTAAGCGTCTTAAGCGTAGAATACTTCTTTTCGAGAAGAACAAGTTCTCAGAAATAGTTGACTTAAACACACAGAAAGGAGACGGAGATCACTTCGACGAGACTCAGATTTTCAGTATTTTCTTCAGCTTCCTTATTAAGAAGAACGAAGGAAGTAAGATGCGTGACAACGACATGAACTCAGACTCAGAGGATTCAATTTACGACGCATACGAACAACAAATTCAGCTTCATCACGGAGACAACATGGTTAACGGAATGCTTATGATGCGTCGTATTAGTATGCAGAACCTTGAAGATGATGAGACACAGGTAGAGTACATTAACCGTGAAATACACACACAGGGAGACCTTCACGTTAGGCGTACAAACCAGGGAATACTTCGTTTCAACATGCGTCGTGGAAAGAAGGGAAGTAACGAGAACATGGGAGTTCATCACGAATCAGGAAACGTTGACGACGCAAACGGAATGAACAACGTTGACGACACAAACAACATGAACAACGTTGACGGAACAAACAACATGAACAACGTTGACGGAACAAACAACATGAACAACATGGACGGACGTAACAACATGAACAACATTAACTCAGTTGACAACATGAATAACCTTAACAACAACGACGGAGAGGAGGAGGAGGAGTGCGTTAATGACGTACTTAACTACGACAACAACAACTACGCAATAAACGAGGACGCAGAAGAGTACATTAAGAACACAAGTGGTGAGAGGGCAGTAATAATATGCAGCGAGAAGCGTATTTACGAGAAAAATGGTAACGGAGACATTATTACACGTAACTACAAGAACGAGGAGAGGTACATTTACCTTAAGAAGTGGATACCATTCAAGAGCATGATTCTTAGTAAGCTTGAGAAACGTAAGCGTAACCGTAAGGAAGCTTACAACACACCACGTGTTCTTATTCTTATTCACAGTTTCCTTTTCATTTTAATAGTATTTATTTTCCTTATGGTTTTTTTCAAGAAGGAACCAATATTCAGATTCAACATGCCTAGTGTAAACAAAAGACTTAACAACTTCTTCAAGAGTACTTCATTCCATGAGATAATTCCTAACTCAGTTGGAAAGTGCAAGACTAAGAAGTACATTGCTAAGGAGCCTTGCTTCAACGTTGGACACATTTATCACGAGGAAAAAACTTTCTACCACGCTACATTACTTTTCCTTCAGGGACTTCGTAGCGTTAAGATAATGAACATGAACTTCTACTACGAGAAGGGAATTTACTACCTTAGCCTTGACGGTTACTTCAAGCACATAATTGGACCATTATTCTTAAAGCTTTGCCTTGGAACTAATTTCTGCCCAATTTCAACTTACGCTTTCCTTGTTGGTTCAAAGCCTACATTTAGTGTTAACGTAGCAGTACAGTGCAACAACAAGAAGCCTCCTTACTACATGACAGACATAATAGTTAAGGACCTTAAGATAACTAAGATTGAGATTGTAAAGCACAGTGACGTAATAGACAACGTTGACATAAAGCTTGACGACGTTCAGGACCGTGTTCAGGAGAAAGTAAACGCAATGCTTGAGGCTAAGAAGAAGATAATAGTTTGGAAGAACCAGAAATACCACCTTGAAGGATTCTTAAACTACCTTATTAGCAAGAACGCTCTTAGCGGATTCAGTTGCGAGCCAATTAACTACtagaggcctataacttcgtatagcatacattatacgaagttattatgactcgagggatatggcagcttaatgttcgtttttcttatttatatatttataccaattgattgtatttataactgtaaaaatgtgtatgttgtgtgcatatttttttttgtgcatgcacatgcatgtaaatagctaaaattatgaacattttattttttgttcagaaaaaaaaaactttacacacataaaatggctagtatgaatagccatattttatataaattaaatcctatgaatttatgaccatattaaaaatttagatatttatggaacataatatgtttgaaacaataagacaaaattattattattattattatttttactgttataattatgtgtctccttcaatgattcataaatagttggacttgatttttaaaatgtttataatatgattagcatagttaaataaaaaaagttgaaaaattaaaaaaaaacatataaacacaaatgatggtttttccttcaatttcgatatcaatttatagaaacaaaatatatacttgtataattttatttttttatataaatcattacatatataattatacaatattttttctaagagataattatatattaatatatataaaaaaaggtgttttttttttttttttttatttttatttttattttatggtaatattttattttccttattttataaattatattagtttatatgtgattaattttatatattatcaatttatatatttttaaatgcttacttaattatctttttttttttttttttttttttttcccctctttttatattaatttatttttgaaaaaattgatatatatatatatatataatatatatatatacatgtagtagtattaaacaatgtataatatatataaataatatatttatatatttcatttcaattttaattttttttggttttttttttttttctttttgtcatatttaaaaaaaattatattcatataagttatgcattttttataaacattattcaatatatgtataatataatatatatatatatattaatgtattattccaatgtgcatgataaaagaaaaaaataatatttataaaaaaaaagaaaaataaaacaaaaaaagaaaaaaaaaaaaaaaaaaaaaaaaatacaaaaataaataatataatttataattatatattcttgtcacaataaaaatatatatatatatatatatatttataatatgtatattttaaactagaaaaggaataactaatattttatttattatcattcaagatttatattttataataataaatacctaatagaaatatatcaggatccatgcatggttcgctaaactgcatcgtcgctgtgtcccagaacatgggcatcggcaagaacggggactacccctggccaccgctcaggaacgaatttagatatttccagagaatgaccacaacctcttcagtagaaggtaaacagaatctggtgattatgggtaagaagacctggttctccattcctgagaagaatcgacctttaaagggtagaattaatttagttctcagcagagaactcaaggaacctccacaaggagctcattttctttccagaagtctagatgatgccttaaaacttactgaacaaccagaattagcaaataaagtagacatggtctggatagttggtggcagttctgtttataaggaagccatgaatcacccaggccatcttaaactatttgtgacaaggatcatgcaagactttgaaagtgacacgttttttccagaaattgatttggagaaatataaacttctgccagaatacccaggtgttctctctgatgtccaggaggagaaaggcattaagtacaaatttgaagtatatgagaagaatgattaagcttatttaataatagattaaaaatattataaaaataaaaacataaacacagaaattacaaaaaaaatacatatgaattttttttttgtaatcttccttataaatatagaataatgaatcatataaaacatatcattattcatttatttacatttaaaattattgtttcagtatctttaatttattatgtatatataaaaataacttacaattttattaataaacaatatatgtttattaattcatgttttgtaatttatgggatagcgattttttttactgtctgtatttttcttttttaattatgttttaattgtattttatttttattattgttctttttatagtattattttaaaacaaaatgtattttctaagaacttataataataataaatataaattttaataaaaattatatttatcttttacaatatgaacataaagtacaacattaatatatagcttttaatatttttattcctaatcatgtaaatcttaaatttttctttttaaacatatgttaaatatttatttctcattatatataagaacatatttattaaatctagaattctatagtgagtcgtattacaattcactggccgtcgttttacaacgtcgtgactgggaaaaccctggcgttacccaacttaatcgccttgcagcacatccccctttcgccagctggcgtaatagcgaagaggcccgcaccgatcgcccttcccaacagttgcgcagcctgaatggcgaatggcgcctgatgcggtattttctccttacgcatctgtgcggtatttcacaccgcatatggtgcactctcagtacaatctgctctgatgccgcatagttaagccagccccgacacccgccaacacccgctgacgcgccctgacgggcttgtctgctcccggcatccgcttacagacaagctgtgaccgtctccgggagctgcatgtgtcagaggttttcaccgtcatcaccgaaacgcgcgagacgaaagggcctcgtgatacgcctatttttataggttaatgtcatgataataatggtttcttagacgtcaggtggcacttttcggggaaatgtgcgcggaacccctatttgtttatttttctaaatacattcaaatatgtatccgctcatgagacaataaccctgataaatgcttcaataatattgaaaaaggaagagtatgagtattcaacatttccgtgtcgcccttattcccttttttgcggcattttgccttcctgtttttgctcacccagaaacgctggtgaaagtaaaagatgctgaagatcagttgggtgcacgagtgggttacatcgaactggatctcaacagcggtaagatccttgagagttttcgccccgaagaacgttttccaatgatgagcacttttaaagttctgctatgtggcgcggtattatcccgtattgacgccgggcaagagcaactcggtcgccgcatacactattctcagaatgacttggttgagtactcaccagtcacagaaaagcatcttacggatggcatgacagtaagagaattatgcagtgctgccataaccatgagtgataacactgcggccaacttacttctgacaacgatcggaggaccgaaggagctaaccgcttttttgcacaacatgggggatcatgtaactcgccttgatcgttgggaaccggagctgaatgaagccataccaaacgacgagcgtgacaccacgatgcctgtagcaatgccaacaacgttgcgcaaactattaactggcgaactacttactctagcttcccggcaacaattaatagactggatggaggcggataaagttgcaggaccacttctgcgctcggcccttccggctggctggtttattgctgataaatctggagccggtgagcgtgggtctcgcggtatcattgcagcactggggccagatggtaagccctcccgtatcgtagttatctacacgacggggagtcaggcaactatggatgaacgaaatagacagatcgctgagataggtgcctcactgattaagcattggtaactgtcagaccaagtttactcatatatactttagattgatttaaaacttcatttttaatttaaaaggatctaggtgaagatcctttttgataatctcatgaccaaaatcccttaacgtgagttttcgttccactgagcgtcagaccccgtagaaaagatcaaaggatcttcttgagatcctttttttctgcgcgtaatctgctgcttgcaaacaaaaaaaccaccgctaccagcggtggtttgtttgccggatcaagagctaccaactctttttccgaaggtaactggcttcagcagagcgcagataccaaatactgtccttctagtgtagccgtagttaggccaccacttcaagaactctgtagcaccgcctacatacctcgctctgctaatcctgttaccagtggctgctgccagtggcgataagtcgtgtcttaccgggttggactcaagacgatagttaccggataaggcgcagcggtcgggctgaacggggggttcgtgcacacagcccagcttggagcgaacgacctacaccgaactgagatacctacagcgtgagctatgagaaagcgccacgcttcccgaagggagaaaggcggacaggtatccggtaagcggcagggtcggaacaggagagcgcacgagggagcttccagggggaaacgcctggtatctttatagtcctgtcgggtttcgccacctctgacttgagcgtcgatttttgtgatgctcgtcaggggggcggagcctatcgaaaaacgccagcaacgcggcctttttacggttcctggccttttgctggccttttgctcacatgttctttcctgcgttatcccctgattctgtggataaccgtattaccgcctttgagtgagctgataccgctcgccgcagccgaacgaccgagcgcagcgagtcagtgagcgaggaagcggaagagcgcccaatacgcaaaccgcctctccccgcgcgttggccgattcattaatgcagctggcacgacaggtttcccgactggaaagcgggcagtgagcgcaacgcaattaatgtgagttagctcactcattaggcaccccaggctttacactttatgcttccggctcgtatgttgtgtggaattgtgagcggataacaatttcacacaggaaacagctatgaccatgattacgccaagctatttaggtgacactatagaatactcgcggccgcTAGTTGGAATTATTACGTAATGCTTATAATAAATGTAAAGAGTATATTTTGGGCTTATCGAAAAAATCACAAATATATAAAATAAATATATTAAATTATTGTTTATTATTTTCGTTTTTTTTATTTTATGCATCATATTTTTTTTTTATAAATGGAATAATAAATCCAGCAGTAAAAATTGTTTTCCATGGAAGTAAGTACCTTATATGAAAAAATGAAAGTGTATCCATAAAAGAAGATATACACAAAGGTGTATATATATATATATATATATATATATATATATATATGTATGTATGTATGTATTTATTTATTTATTTATTATTTATATATATATTTTCTTTATATTTTCTTTAAGGTATTATTGAGGACAAAACCAAAGTTTACGAGAACTCGATAATTGAATCAATATGGCTATTGTTTACCAAGGGAAGTTATTTTAATATGGCCATGCTTCTATTCTTTTCTATTATAATACCGTTTCTCAAGTTACTTATGGTTAGTGATAACTTTTATAGTTTTATCGTATTATATAAAATGAATAAAAAACGAGAAGAGGAAAAGAGAAGGAGACGAAGAAGAATACGTGCACGAGAATATAACAATGCTAAATATAACAAGAATAGATATAAAATGAATAAATATAATAGAAATAAGAATATTAATGATATATACAAGGATGATATATATTATAGTGAAAATATTTTTAAGAATGATGAGATAAATTATATTAATACAATAAATGAGAATGAAGAATTTATTTTAAAGAAATTTAAAATTTTAAATTTCATATCGCGTTTTCAATTTGTTGATGTATTTATATCTTTATTTATAGTATCATCATTAAATTTATATTTATTAGAGGCAAGGATGTTAAATGGTGCATATTATTTTTTGAATTATTGTATGTTATCAACAATATCATCTTTTTTGTTATTTAGTTTTACTTCTTTAAAAATACATATATTTAAAAATGGAAATATTAAGATATCTGCATGTCTCAATGAATCAAATCTAGAGGTTACTACAAGCGGTCCCCTTTCAACAAAAGATCTTGTCGAAGAAGAAGGACATGCACAAATAAATAATCTAATTATTAATGATAAGAATATGACTAGTGGTGTTGTTAATGATTTTTCAGGTAATGGAAACAATGTCGAATTAACAGACGACCTTAAGAGTGGCGAACCCCAAAACAGAGATGATATACAAACTGAAGAAACAAAAAAAGAAAAAATGAATACTAGAACTCATAATGATGAAGATAATACAAAAAAAAAAAATATAAAGGATAAAAAGAAAGCTAACGGTGATGATAAAGTAATACAAAAATGTATTGATAATGAAAGGAAAAAAAAACAAAATGGTATGATTCAAAGTGTAAACGATGGTGATAAAAATAGTAATTTTAATAATAACAATAATAATAATATTAATGGTGATAGTAATAATAATAATATTAATGGTGATAGTAATAATAATAATATTAATGGTGATAGTAATAATAATAATATTAATGGTGATAGTAATAATAATAATATTAATGGTGATAGTAATAATAATAATATTAATGGTGATAGTAATAATAATAATTATCATAATAATTATCATAATAATTATCGTAATAATTATCATAATAATTATCGTAATAATAATTGTAGGAATAATATTTTAGAACAAAATAAATGTGATAAAAATGTTTTGTGTTATAATAATATATATAATACAATGAAAGATAATGATACTTATATATATTTAAAAAAGAATAAATTTAATTCGTTATTAAAAAGTAATTGTATCAAAACTAATTTTAATATGATAAAGATAGGTTATGTAATATTTTTATTTGTGTTATTATGTTTATGTATATATTTAATAACAGGAGTAGAATGCAGTTTATTTGGTATATATATATATTTGAGTTATTTTAATTTTAATATTGAAGGAATATTAATTGATTATATGGATATGTTAAATATTTTAAAATTAAAAATAAAGAAAGGATATATCTATCCTTTTTTTGTTATGTTACCATTTATTTTCCCTGTAATAATATCTATGTGTTTCTTTTTAAGTGTATTTTTTTTAAATATGTATTATGAAAGCTTTTCAAAGTTATATAAAAAGATCAGTGAACTAAAGAATGAATTCATAAACTCCTCAGAAAATGACAATGTGAACGAAAGAATATTGGTATCCGAAACTTCTAATCATTTATGTTTAAATGAAAGTAACGATAAAGTATCTAATACATCAGATGATTTTTTATCAAGAAATAATAGCAACATATCTAGTTCAAAAAGTGAAATGATAAATTCAAATTTTGTATTTAATAAACTTTTGAATTTTTATTTTTCATTTGCAGTTTTCTTTTCTTATTTAGGTTCTGCATTTTTACATATATCCTTAGGTGAAATTATATGTATCGCATTATTAACATTTTATCAAATTGTAAAGCATACCAATAATTTGAATATCACAATTTTACTTAAGAGTGAGAAAATTAAATTTTGTAAATTCCTTTTGTTTATATTATATGGGTTGTTATGCTTTTCCATAAATTTGTATGTAAACCAATGGGAGGAATATATAACGAAATTAAAAAGGTTAAAAAGACGTATTTTATTGTTTGAAAAAAATAAATTTAGTGAGATTGTGGATCTGAATACTCAAAAGGGTGATGGTGACCATTTTGATGAAACACAAATATTTTCCATATTTTTTTCTTTTTTAATAAAAAAAAATGAGGGTTCAAAAATGAGGGATAATGATATGAATAGTGATAGCGAAGACAGTATATATGATGCTTATGAGCAGCAGATACAATTACACCATGGTGATAATATGGTCAATGGTATGTTGATGATGAGAAGGATATCGATGCAAAATTTAGAGGACGACGAAACCCAAGTTGAATATATAAATAGGGAGATCCATACGCAAGGTGATTTACATGTACGAAGAACGAATCAAGGTATTTTAAGATTTAATATGAGAAGGGGCAAAAAAGGGTCCAATGAAAATATGGGTGTCCACCATGAGAGTGGTAATGTGGATGATGCGAATGGTATGAATAATGTGGATGATACGAATAATATGAATAATGTGGATGGTACGAATAATATGAATAATGTGGATGGTACGAATAATATGAATAATATGGATGGTAGGAATAATATGAATAATATAAATAGTGTGGATAATATGAACAATTTAAATAATAATGATGGTGAAGAAGAAGAAGAATGTGTGAACGATGTTTTAAATTATGATAATAATAATTATGCTATTAATGAAGATGCTGAGGAATATATAAAAAATACGTCAGGAGAACGAGCTGTTATTATTTGTTCTGAAAAAAGAATATATGAAAAGAACGGAAATGGTGATATAATAACTAGAAATTATAAAAATGAAGAACGATATATATATTTAAAAAAATGGATTCCTTTTAAATCTATGATATTGTCAAAATTAGAAAAGAGGAAAAGGAATAGAAAAGAGGCATATAATACTCCTAGAGTATTAATATTAATACATTCCTTTTTATTTATACTTATTGTTTTCATATTTTTAATGGTATTCTTTAAAAAAGAGCCTATTTTTCGTTTTAATATGCCATCAGTTAATAAGCGTTTAAATAATTTTTTTAAATCAACAAGTTTTCACGAAATTATACCAAATAGTGTAGGGAAATGTAAAACAAAAAAATATATAGCAAAAGAACCATGTTTTAATGTGGGTCATATATACCATGAAGAGAAGACATTTTATCATGCAACTCTTTTATTTTTACAAGGTTTAAGATCTGTAAAAATTATGAATATGAATTTTTATTATGAAAAAGGCATATATTATTTATCTTTGGATGGATATTTTAAACATATTATAGGTCCTCTCTTTCTTAAATTATGTTTAGGTACAAACTTTTGTCCCATAAGTACATATGCATTTTTAGTAGGAAGTAAACCAACTTTCTCAGTAAATGTTGCTGTTCAATGTAATAATAAAAAACCACCATATTATATGACGGATATTATTGTAAAAGATTTAAAAATTACAAAAATAGAAATAGTTAAACATTCAGATGTTATCGATAATGTGGATATTAAATTAGATGATGTACAAGATAGAGTACAAGAAAAGGTTAATGCCATGTTAGAAGCAAAAAAAAAAATTATCGTCTGGAAAAATCAAAAGTATCATTTGGAGGGTTTTCTCAATTATTTAATATCTAAAAATGCATTATCTGGGTTTTCCTGTGAACCCATAAATTATTGAacttgacaaattaaaacacgttaaatgtatatatgtatatatattatatatgtatatatttatttatatttttcttacatttccatttgggtgaacatatattcttacccttttcataaattgttttttttttttttgatatttatcaataatatattcagtaaaaatatattatttcacataatttaatattttttggaataacatataaatgtcaatgtatatgtgtatatatatatatatatatatatatgtatatatttttttttctttttcatatatttatatgcgcccatttgtttttatctatatatatatatatttatatatccgaatttctttatattgttatattccgatttgattatatataaaattttttttaattcataatttttttttttttttttttcttttttttcatttccatgtgatttatttcaatccgatattatttatatgttatatatatattattctataatatgataaaaggatatttttttttcttatttgtatttttagattataaatataaagatattcttatatatagattatatctgcttttttttttttttttttttttttttctattattaatttttaaattatctacttacgaattttttaattgttgatatggccttcatgagtatattattatgcatgaaagaaaaaaaaaaaaaaaaatatatatatgtatataatatatatatatatatatataatataccaatttttattatcttatattttacatgcactataaatgcatagaatttttcttttttttttttttgatttcttctccccaaatttttggttatttatataatatacgtattataaatatgtacatataatatgtatatattataatgtgtaattttccacatatatatataaaatatatatatatatatatatatattttaataatacttgaacttttaaaattaaatatataaaaaggaatccaattaatttatggttatattatttgttattcactagaaaattctctagaatttaataaaacatatat

**5’ amplicon size: 2935bps**

**3’ amplicon size: 1116bps**

### > Halo-PF3D7_0214800 Halo yDHODH recodonizedsequence

cattttattccgtaaaattaaaatgaattccaccaaaggtttaacatatacataatatatatttatatatatatatatatgtatatatatatatataaaataaaaataataatatataatatatttttatattacattattatattatattatattatattatattttattttatttttttaatatagaacttcatgtagctaaaataaaaatgagttcacatttttcgttaattaaatatattatatatatatatatatgcaaaattggtattatatataaatataatatataatttgtttccttttttttagtttttaaaatatagatttataaataatttttttcttttttttaagaattattttgatttaccatatatattatttaaaatcccaagttaaaattaaaaaaaaaaaggagtttttattataatattatatatatttatatatatattttaactactttaattttaaaagtgtgagagtaagaaaatgaaaaaaaaaaaaaaaaaaaaaaaaaggattattttttatttatatattataatatatataatatattaataataaataaaataaacatgtaaataaatattatatatatatatcataatatagtatcatgaaataatatatatttataaaaatattattataatatatagttaattattatattatatattacataatatttataatatattatatattggattatagttttacctattgggggatcttggttctataaaaaaatttcaatttttttttaccctttatttttttttgttcactatatatatatattgttatatataacataatatattaatagtttcataatattatatgtatatagaaagaaaaacatatatatatatatatatatattgttcattatttctaaatatatatattattatatatattatataaaaaatatattattttagatttactcctttatatataaattaatattttttatagatttcgattttttcctatttacataatttttaattttttataattttaatttttctacattatatatgtattatatatatatataaataatctatatatttttaaaataattcaatgtcattttattaattcaattaaatacaaattaatttcttattttctttaattatatatatatatattataatatttatagaacctatattatagatttaatttttaatctgttaaaagaaaaaaagtacaaaaaattatacatttatttacatatatatatatatatatatatatatatgtataatttttaatcttttatttttagaggtattttatgttattttttattttacatatgttgttgtataattttttttgtatatgcccttatattgattttcttttttttttttaaataatatagtgtttgctttaataaaaaaaaggaaatatcaggattttattgaatacatacatatatatatatatatatatatatatatatatatatatatatatatataactattagaattcccatagatgaatatataaacatataataatattatatctataatatttatatttttattatacatacatactgaatatattttaaagttacgaataatgaatgctttttttttttttttttttttgttgctttaaataaatggtaatcataaatgaaattagtctagagctaaaatgaaaattaaagaaaatataaaaagtatataataataaatgtgttataatatttaaaataaaacaaaaaaaaaaaaaaaaaaaaaaaaaaaagtaaattacaataatttattctataaagttcagaATG**TTGGAATTATTACGTAATGCTTATAATAAATGTAAAGAGTATATTTTGGGCTTATCGAAAAAATCACAAATATATAAAATAAATATATTAAATTATTGTTTATTATTTTCGTTTTTTTTATTTTATGCATCATATTTTTTTTTTATAAATGGAATAATAAATCCAGCAGTAAAAATTGTTTTCCATGGAA*gtaagtaccttatatgaaaaaatgaaagtgtatccataaaagaagatatacacaaaggtgtatatatatatatatatatatatatatatatatatatgtatgtatgtatgtatttatttatttatttattatttatatatatattttctttatattttctttaag*GTATTATTGAGGACAAAACCAAAGTTTACGAGAACTCGATAATTGAATCAATATGGCTATTGTTTACCAAGGGAAGTTATTTTAATATGGCCATGCTTCTATTCTTTTCTATTATAATACCGTTTCTCAAGTTACTTATGGTTAGTGATAACTTTTATAGTTTTATCGTATTATATAAAATGAATAAAAAACGAGAAGAGGAAAAGAGAAGGAGACGAAGAAGAATACGTGCACGAGAATATAACAATGCTAAATATAACAAGAATAGATATAAAATGAATAAATATAATAGAAATAAGAATATTAATGATATATACAAGGATGATATATATTATAGTGAAAATATTTTTAAGAATGATGAGATAAATTATATTAATACAATAAATGAGAATGAAGAATTTATTTTAAAGAAATTTAAAATTTTAAATTTCATATCGCGTTTTCAATTTGTTGATGTATTTATATCTTTATTTATAGTATCATCATTAAATTTATATTTATTAGAGGCAAGGATGTTAAATGGTGC**AGGTTTAAACGAGCAGAAGTTAATATCAGAAGAGGATTTGGGTGAACAAAAACTCATAAGCGAAGAAGATTTAATAACTTCGTATAGCATACATTATACGAAGTTATCCGGAGAAGGAAGAGGAAGTTTATTAACATGTGGAGATGTAGAAGAAAATCCAGGACCAATGACAGCCAGTTTAACTACCAAGTTCTTGAACAATACCTATGAAAACCCATTTATGAATGCATCCGGTGTTCATTGCATGACTACACAAGAATTAGATGAATTAGCAAACTCTAAAGCTGGCGCATTCATTACAAAGAGTGCTACAACCTTAGAAAGAGAAGGTAACCCTGAACCACGTTACATTTCTGTCCCTCTAGGCAGTATCAACTCCATGGGTTTACCAAACGAAGGTATCGACTACTATTTGTCCTATGTATTAAACCGTCAAAAGAATTATCCTGATGCACCTGCTATTTTCTTCTCAGTTGCTGGTATGAGCATTGATGAAAATTTAAATTTGTTGAGGAAAATCCAAGATAGCGAATTCAACGGTATTACCGAGTTAAACTTGTCTTGTCCTAATGTGCCTGGGAAACCACAAGTTGCTTATGACTTTGACTTGACAAAGGAAACCTTGGAAAAGGTTTTTGCCTTTTTCAAAAAACCTCTTGGTGTCAAGTTGCCTCCTTATTTTGATTTTGCCCATTTTGATATCATGGCAAAAATATTGAACGAGTTCCCATTAGCTTATGTCAACTCTATCAATAGTATAGGAAATGGTCTTTTCATTGATGTGGAGAAGGAGAGTGTAGTAGTGAAGCCAAAGAATGGTTTCGGGGGTATTGGAGGTGAATATGTTAAGCCAACCGCGCTCGCCAATGTTCGTGCATTTTACACTCGTTTGAGACCTGAAATCAAAGTTATCGGTACAGGTGGAATTAAGTCCGGTAAGGATGCATTTGAACATCTTCTATGTGGTGCCTCTATGCTACAGATTGGTACAGAATTACAAAAAGAGGGCGTCAAGATTTTTGAACGTATCGAAAAAGAATTAAAAGACATAATGGAAGCTAAGGGTTATACATCCATAGATCAGTTCCGTGGGAAGTTGAACAGCATTGGTGAAGGTAGAGGTTCTTTGTTGACTTGTGGTGATGTTGAAGAAAATCCAGGTCCAGCTAGCATGGCAGAAATTGGTACGGGTTTTCCATTTGATCCTCATTATGTGGAGGTGCTGGGGGAAAGGATGCATTATGTTGACGTAGGACCAAGAGATGGTACTCCAGTGTTATTTTTGCATGGAAACCCAACCTCGAGTTATGTATGGAGAAATATAATTCCACATGTAGCACCAACACATAGATGTATAGCTCCTGATTTAATTGGTATGGGAAAAAGTGATAAACCTGACTTAGGATATTTTTTTGATGATCATGTCCGTTTTATGGATGCTTTCATTGAAGCTCTTGGCCTTGAAGAAGTAGTATTAGTTATACATGATTGGGGATCCGCCTTAGGATTTCATTGGGCCAAGAGGAATCCTGAAAGAGTAAAAGGAATAGCATTCATGGAATTCATACGACCAATCCCCACATGGGATGAATGGCCAGAATTTGCACGCGAAACATTTCAAGCTTTTAGAACTACAGATGTTGGTAGAAAATTAATAATAGATCAAAATGTATTTATAGAAGGAACTTTACCTATGGGTGTTGTAAGGCCGTTAACAGAAGTTGAAATGGACCACTACCGTGAACCTTTTTTAAATCCAGTAGATAGAGAGCCCTTATGGAGATTTCCTAATGAATTACCTATTGCAGGTGAACCCGCGAATATTGTTGCTTTAGTAGAAGAATATATGGATTGGTTACATCAGTCTCCTGTTCCTAAACTTCTATTTTGGGGTACACCTGGAGTTCTAATACCACCAGCTGAAGCAGCAAGATTAGCAAAATCATTACCAAATTGTAAAGCTGTTGATATAGGTCCTGGGTTGAATTTATTACAAGAAGATAATCCAGATTTGATTGGATCTGAGATAGCTAGATGGCTAAGTACATTAGAAATTTCAGGTACGCGTGCTAGAGGTGCTGCTGCTGGTGCTGGAGGTGCAGGTAGACCTAGGATGCTTGAGCTTCTTAGAAACGCATACAACAAGTGCAAGGAATACATACTTGGACTTAGTAAGAAGAGTCAGATTTACAAGATTAACATTCTTAACTACTGCCTTCTTTTCAGTTTCTTCCTTTTCTACGCTAGTTACTTCTTCTTCATTAACGGTATTATTAACCCTGCTGTTAAGATAGTATTTCACGGTTCAATAATAGAAGATAAGACAAAGGTATATGAAAATAGTATTATAGAGAGTATTTGGTTGCTTTTCACAAAAGGTTCATACTTCAACATGGCAATGTTATTGTTTTTCAGCATAATTATTCCATTCTTAAAACTTTTAATGGTATCAGACAATTTCTACTCATTCATAGTTCTTTACAAGATGAACAAGAAGAGGGAGGAAGAGAAACGTCGTCGTAGGCGTCGTATTAGAGCTAGGGAGTACAATAACGCAAAGTACAATAAAAACCGTTACAAGATGAACAAGTACAACCGTAACAAAAACATAAACGACATTTATAAAGACGACATTTACTACTCAGAGAACATATTCAAAAACGACGAAATTAACTACATAAACACTATTAACGAAAACGAGGAGTTCATACTTAAAAAGTTCAAGATACTTAACTTTATTAGTAGATTCCAGTTCGTAGACGTTTTCATTAGCCTTTTCATTGTTAGTAGTCTTAACCTTTACCTTCTTGAAGCTCGTATGCTTAACGGAGCTTACTACTTCCTTAACTACTGCATGCTTAGTACTATTAGTAGCTTCCTTCTTTTCTCATTCACAAGCCTTAAGATTCACATTTTCAAGAACGGTAACATAAAAATTAGCGCTTGCTTAAACGAGAGTAACTTGGAAGTAACAACTTCAGGACCATTAAGTACTAAGGACTTAGTTGAGGAGGAGGGTCACGCTCAGATTAACAACTTGATAATAAACGACAAAAACATGACATCAGGAGTAGTAAACGACTTCAGTGGAAACGGTAATAACGTTGAGCTTACTGATGATTTAAAATCAGGAGAGCCACAGAATCGTGACGACATTCAGACAGAGGAGACTAAGAAGGAGAAGATGAACACACGTACACACAACGACGAGGACAACACTAAGAAGAAGAACATTAAAGACAAGAAAAAGGCAAATGGAGACGACAAGGTTATTCAGAAGTGCATAGACAACGAGCGTAAGAAGAAGCAGAACGGAATGATACAGTCAGTTAATGACGGAGACAAGAACTCAAACTTCAACAACAATAACAACAACAACATAAACGGAGACTCAAACAACAACAACATAAACGGAGACTCAAACAACAACAACATAAACGGAGACTCAAACAACAACAACATAAACGGAGACTCAAACAACAACAACATAAACGGAGACTCAAACAACAACAACATAAACGGAGACTCAAACAACAACAACTACCACAACAACTACCACAACAACTACAGAAACAACTACCACAACAACTACAGAAACAACAACTGCCGTAACAACATACTTGAGCAGAACAAGTGCGACAAGAACGTACTTTGCTACAACAACATTTACAACACTATGAAGGACAACGACACATACATTTACCTTAAGAAAAACAAGTTCAACAGTCTTCTTAAGTCAAACTGCATAAAGACAAACTTCAACATGATTAAAATTGGATACGTTATTTTCCTTTTCGTTCTTCTTTGCCTTTGCATTTACCTTATTACTGGTGTTGAGTGTTCACTTTTCGGAATTTACATTTACCTTTCATACTTCAACTTCAACATAGAGGGTATTCTTATAGACTACATGGACATGCTTAACATACTTAAGCTTAAGATTAAAAAGGGTTACATATACCCATTCTTCGTAATGCTTCCTTTCATATTTCCAGTTATTATTAGCATGTGCTTTTTCCTTTCAGTTTTCTTCCTTAACATGTACTACGAGTCATTCAGTAAACTTTACAAGAAAATATCAGAGTTGAAAAACGAGTTTATTAATAGTAGTGAGAACGATAACGTTAATGAGCGTATTCTTGTTAGTGAGACAAGCAACCACCTTTGCCTTAACGAGTCAAATGACAAGGTTAGCAACACTAGTGACGACTTCCTTAGTCGTAACAACTCAAATATTAGCTCAAGTAAGTCAGAGATGATTAACAGTAACTTCGTTTTCAACAAGTTACTTAACTTCTACTTCAGTTTCGCTGTATTTTTCAGCTACCTTGGAAGCGCTTTCCTTCACATTAGTCTTGGAGAGATAATTTGCATAGCTCTTCTTACTTTCTACCAGATAGTTAAACACACAAACAACCTTAACATAACTATACTTTTAAAATCAGAAAAGATAAAGTTCTGCAAGTTTTTACTTTTCATTCTTTACGGACTTCTTTGTTTCAGTATTAACCTTTACGTTAATCAGTGGGAAGAGTACATTACAAAGCTTAAGCGTCTTAAGCGTAGAATACTTCTTTTCGAGAAGAACAAGTTCTCAGAAATAGTTGACTTAAACACACAGAAAGGAGACGGAGATCACTTCGACGAGACTCAGATTTTCAGTATTTTCTTCAGCTTCCTTATTAAGAAGAACGAAGGAAGTAAGATGCGTGACAACGACATGAACTCAGACTCAGAGGATTCAATTTACGACGCATACGAACAACAAATTCAGCTTCATCACGGAGACAACATGGTTAACGGAATGCTTATGATGCGTCGTATTAGTATGCAGAACCTTGAAGATGATGAGACACAGGTAGAGTACATTAACCGTGAAATACACACACAGGGAGACCTTCACGTTAGGCGTACAAACCAGGGAATACTTCGTTTCAACATGCGTCGTGGAAAGAAGGGAAGTAACGAGAACATGGGAGTTCATCACGAATCAGGAAACGTTGACGACGCAAACGGAATGAACAACGTTGACGACACAAACAACATGAACAACGTTGACGGAACAAACAACATGAACAACGTTGACGGAACAAACAACATGAACAACATGGACGGACGTAACAACATGAACAACATTAACTCAGTTGACAACATGAATAACCTTAACAACAACGACGGAGAGGAGGAGGAGGAGTGCGTTAATGACGTACTTAACTACGACAACAACAACTACGCAATAAACGAGGACGCAGAAGAGTACATTAAGAACACAAGTGGTGAGAGGGCAGTAATAATATGCAGCGAGAAGCGTATTTACGAGAAAAATGGTAACGGAGACATTATTACACGTAACTACAAGAACGAGGAGAGGTACATTTACCTTAAGAAGTGGATACCATTCAAGAGCATGATTCTTAGTAAGCTTGAGAAACGTAAGCGTAACCGTAAGGAAGCTTACAACACACCACGTGTTCTTATTCTTATTCACAGTTTCCTTTTCATTTTAATAGTATTTATTTTCCTTATGGTTTTTTTCAAGAAGGAACCAATATTCAGATTCAACATGCCTAGTGTAAACAAAAGACTTAACAACTTCTTCAAGAGTACTTCATTCCATGAGATAATTCCTAACTCAGTTGGAAAGTGCAAGACTAAGAAGTACATTGCTAAGGAGCCTTGCTTCAACGTTGGACACATTTATCACGAGGAAAAAACTTTCTACCACGCTACATTACTTTTCCTTCAGGGACTTCGTAGCGTTAAGATAATGAACATGAACTTCTACTACGAGAAGGGAATTTACTACCTTAGCCTTGACGGTTACTTCAAGCACATAATTGGACCATTATTCTTAAAGCTTTGCCTTGGAACTAATTTCTGCCCAATTTCAACTTACGCTTTCCTTGTTGGTTCAAAGCCTACATTTAGTGTTAACGTAGCAGTACAGTGCAACAACAAGAAGCCTCCTTACTACATGACAGACATAATAGTTAAGGACCTTAAGATAACTAAGATTGAGATTGTAAAGCACAGTGACGTAATAGACAACGTTGACATAAAGCTTGACGACGTTCAGGACCGTGTTCAGGAGAAAGTAAACGCAATGCTTGAGGCTAAGAAGAAGATAATAGTTTGGAAGAACCAGAAATACCACCTTGAAGGATTCTTAAACTACCTTATTAGCAAGAACGCTCTTAGCGGATTCAGTTGCGAGCCAATTAACTACtagaggcctataacttcgtatagcatacattatacgaagttattatgactcgagggatatggcagcttaatgttcgtttttcttatttatatatttataccaattgattgtatttataactgtaaaaatgtgtatgttgtgtgcatatttttttttgtgcatgcacatgcatgtaaatagctaaaattatgaacattttattttttgttcagaaaaaaaaaactttacacacataaaatggctagtatgaatagccatattttatataaattaaatcctatgaatttatgaccatattaaaaatttagatatttatggaacataatatgtttgaaacaataagacaaaattattattattattattatttttactgttataattatgtgtctccttcaatgattcataaatagttggacttgatttttaaaatgtttataatatgattagcatagttaaataaaaaaagttgaaaaattaaaaaaaaacatataaacacaaatgatggtttttccttcaatttcgatatcaatttatagaaacaaaatatatacttgtataattttatttttttatataaatcattacatatataattatacaatattttttctaagagataattatatattaatatatataaaaaaaggtgttttttttttttttttttatttttatttttattttatggtaatattttattttccttattttataaattatattagtttatatgtgattaattttatatattatcaatttatatatttttaaatgcttacttaattatctttttttttttttttttttttttttcccctctttttatattaatttatttttgaaaaaattgatatatatatatatatataatatatatatatacatgtagtagtattaaacaatgtataatatatataaataatatatttatatatttcatttcaattttaattttttttggttttttttttttttctttttgtcatatttaaaaaaaattatattcatataagttatgcattttttataaacattattcaatatatgtataatataatatatatatatatattaatgtattattccaatgtgcatgataaaagaaaaaaataatatttataaaaaaaaagaaaaataaaacaaaaaaagaaaaaaaaaaaaaaaaaaaaaaaaatacaaaaataaataatataatttataattatatattcttgtcacaataaaaatatatatatatatatatatatttataatatgtatattttaaactagaaaaggaataactaatattttatttattatcattcaagatttatattttataataataaatacctaatagaaatatatcaggatccatgcatggttcgctaaactgcatcgtcgctgtgtcccagaacatgggcatcggcaagaacggggactacccctggccaccgctcaggaacgaatttagatatttccagagaatgaccacaacctcttcagtagaaggtaaacagaatctggtgattatgggtaagaagacctggttctccattcctgagaagaatcgacctttaaagggtagaattaatttagttctcagcagagaactcaaggaacctccacaaggagctcattttctttccagaagtctagatgatgccttaaaacttactgaacaaccagaattagcaaataaagtagacatggtctggatagttggtggcagttctgtttataaggaagccatgaatcacccaggccatcttaaactatttgtgacaaggatcatgcaagactttgaaagtgacacgttttttccagaaattgatttggagaaatataaacttctgccagaatacccaggtgttctctctgatgtccaggaggagaaaggcattaagtacaaatttgaagtatatgagaagaatgattaagcttatttaataatagattaaaaatattataaaaataaaaacataaacacagaaattacaaaaaaaatacatatgaattttttttttgtaatcttccttataaatatagaataatgaatcatataaaacatatcattattcatttatttacatttaaaattattgtttcagtatctttaatttattatgtatatataaaaataacttacaattttattaataaacaatatatgtttattaattcatgttttgtaatttatgggatagcgattttttttactgtctgtatttttcttttttaattatgttttaattgtattttatttttattattgttctttttatagtattattttaaaacaaaatgtattttctaagaacttataataataataaatataaattttaataaaaattatatttatcttttacaatatgaacataaagtacaacattaatatatagcttttaatatttttattcctaatcatgtaaatcttaaatttttctttttaaacatatgttaaatatttatttctcattatatataagaacatatttattaaatctagaattctatagtgagtcgtattacaattcactggccgtcgttttacaacgtcgtgactgggaaaaccctggcgttacccaacttaatcgccttgcagcacatccccctttcgccagctggcgtaatagcgaagaggcccgcaccgatcgcccttcccaacagttgcgcagcctgaatggcgaatggcgcctgatgcggtattttctccttacgcatctgtgcggtatttcacaccgcatatggtgcactctcagtacaatctgctctgatgccgcatagttaagccagccccgacacccgccaacacccgctgacgcgccctgacgggcttgtctgctcccggcatccgcttacagacaagctgtgaccgtctccgggagctgcatgtgtcagaggttttcaccgtcatcaccgaaacgcgcgagacgaaagggcctcgtgatacgcctatttttataggttaatgtcatgataataatggtttcttagacgtcaggtggcacttttcggggaaatgtgcgcggaacccctatttgtttatttttctaaatacattcaaatatgtatccgctcatgagacaataaccctgataaatgcttcaataatattgaaaaaggaagagtatgagtattcaacatttccgtgtcgcccttattcccttttttgcggcattttgccttcctgtttttgctcacccagaaacgctggtgaaagtaaaagatgctgaagatcagttgggtgcacgagtgggttacatcgaactggatctcaacagcggtaagatccttgagagttttcgccccgaagaacgttttccaatgatgagcacttttaaagttctgctatgtggcgcggtattatcccgtattgacgccgggcaagagcaactcggtcgccgcatacactattctcagaatgacttggttgagtactcaccagtcacagaaaagcatcttacggatggcatgacagtaagagaattatgcagtgctgccataaccatgagtgataacactgcggccaacttacttctgacaacgatcggaggaccgaaggagctaaccgcttttttgcacaacatgggggatcatgtaactcgccttgatcgttgggaaccggagctgaatgaagccataccaaacgacgagcgtgacaccacgatgcctgtagcaatgccaacaacgttgcgcaaactattaactggcgaactacttactctagcttcccggcaacaattaatagactggatggaggcggataaagttgcaggaccacttctgcgctcggcccttccggctggctggtttattgctgataaatctggagccggtgagcgtgggtctcgcggtatcattgcagcactggggccagatggtaagccctcccgtatcgtagttatctacacgacggggagtcaggcaactatggatgaacgaaatagacagatcgctgagataggtgcctcactgattaagcattggtaactgtcagaccaagtttactcatatatactttagattgatttaaaacttcatttttaatttaaaaggatctaggtgaagatcctttttgataatctcatgaccaaaatcccttaacgtgagttttcgttccactgagcgtcagaccccgtagaaaagatcaaaggatcttcttgagatcctttttttctgcgcgtaatctgctgcttgcaaacaaaaaaaccaccgctaccagcggtggtttgtttgccggatcaagagctaccaactctttttccgaaggtaactggcttcagcagagcgcagataccaaatactgtccttctagtgtagccgtagttaggccaccacttcaagaactctgtagcaccgcctacatacctcgctctgctaatcctgttaccagtggctgctgccagtggcgataagtcgtgtcttaccgggttggactcaagacgatagttaccggataaggcgcagcggtcgggctgaacggggggttcgtgcacacagcccagcttggagcgaacgacctacaccgaactgagatacctacagcgtgagctatgagaaagcgccacgcttcccgaagggagaaaggcggacaggtatccggtaagcggcagggtcggaacaggagagcgcacgagggagcttccagggggaaacgcctggtatctttatagtcctgtcgggtttcgccacctctgacttgagcgtcgatttttgtgatgctcgtcaggggggcggagcctatcgaaaaacgccagcaacgcggcctttttacggttcctggccttttgctggccttttgctcacatgttctttcctgcgttatcccctgattctgtggataaccgtattaccgcctttgagtgagctgataccgctcgccgcagccgaacgaccgagcgcagcgagtcagtgagcgaggaagcggaagagcgcccaatacgcaaaccgcctctccccgcgcgttggccgattcattaatgcagctggcacgacaggtttcccgactggaaagcgggcagtgagcgcaacgcaattaatgtgagttagctcactcattaggcaccccaggctttacactttatgcttccggctcgtatgttgtgtggaattgtgagcggataacaatttcacacaggaaacagctatgaccatgattacgccaagctatttaggtgacactatagaatactcgcggccgcTAGTTGGAATTATTACGTAATGCTTATAATAAATGTAAAGAGTATATTTTGGGCTTATCGAAAAAATCACAAATATATAAAATAAATATATTAAATTATTGTTTATTATTTTCGTTTTTTTTATTTTATGCATCATATTTTTTTTTTATAAATGGAATAATAAATCCAGCAGTAAAAATTGTTTTCCATGGAAGTAAGTACCTTATATGAAAAAATGAAAGTGTATCCATAAAAGAAGATATACACAAAGGTGTATATATATATATATATATATATATATATATATATATGTATGTATGTATGTATTTATTTATTTATTTATTATTTATATATATATTTTCTTTATATTTTCTTTAAGGTATTATTGAGGACAAAACCAAAGTTTACGAGAACTCGATAATTGAATCAATATGGCTATTGTTTACCAAGGGAAGTTATTTTAATATGGCCATGCTTCTATTCTTTTCTATTATAATACCGTTTCTCAAGTTACTTATGGTTAGTGATAACTTTTATAGTTTTATCGTATTATATAAAATGAATAAAAAACGAGAAGAGGAAAAGAGAAGGAGACGAAGAAGAATACGTGCACGAGAATATAACAATGCTAAATATAACAAGAATAGATATAAAATGAATAAATATAATAGAAATAAGAATATTAATGATATATACAAGGATGATATATATTATAGTGAAAATATTTTTAAGAATGATGAGATAAATTATATTAATACAATAAATGAGAATGAAGAATTTATTTTAAAGAAATTTAAAATTTTAAATTTCATATCGCGTTTTCAATTTGTTGATGTATTTATATCTTTATTTATAGTATCATCATTAAATTTATATTTATTAGAGGCAAGGATGTTAAATGGTGCATATTATTTTTTGAATTATTGTATGTTATCAACAATATCATCTTTTTTGTTATTTAGTTTTACTTCTTTAAAAATACATATATTTAAAAATGGAAATATTAAGATATCTGCATGTCTCAATGAATCAAATCTAGAGGTTACTACAAGCGGTCCCCTTTCAACAAAAGATCTTGTCGAAGAAGAAGGACATGCACAAATAAATAATCTAATTATTAATGATAAGAATATGACTAGTGGTGTTGTTAATGATTTTTCAGGTAATGGAAACAATGTCGAATTAACAGACGACCTTAAGAGTGGCGAACCCCAAAACAGAGATGATATACAAACTGAAGAAACAAAAAAAGAAAAAATGAATACTAGAACTCATAATGATGAAGATAATACAAAAAAAAAAAATATAAAGGATAAAAAGAAAGCTAACGGTGATGATAAAGTAATACAAAAATGTATTGATAATGAAAGGAAAAAAAAACAAAATGGTATGATTCAAAGTGTAAACGATGGTGATAAAAATAGTAATTTTAATAATAACAATAATAATAATATTAATGGTGATAGTAATAATAATAATATTAATGGTGATAGTAATAATAATAATATTAATGGTGATAGTAATAATAATAATATTAATGGTGATAGTAATAATAATAATATTAATGGTGATAGTAATAATAATAATATTAATGGTGATAGTAATAATAATAATTATCATAATAATTATCATAATAATTATCGTAATAATTATCATAATAATTATCGTAATAATAATTGTAGGAATAATATTTTAGAACAAAATAAATGTGATAAAAATGTTTTGTGTTATAATAATATATATAATACAATGAAAGATAATGATACTTATATATATTTAAAAAAGAATAAATTTAATTCGTTATTAAAAAGTAATTGTATCAAAACTAATTTTAATATGATAAAGATAGGTTATGTAATATTTTTATTTGTGTTATTATGTTTATGTATATATTTAATAACAGGAGTAGAATGCAGTTTATTTGGTATATATATATATTTGAGTTATTTTAATTTTAATATTGAAGGAATATTAATTGATTATATGGATATGTTAAATATTTTAAAATTAAAAATAAAGAAAGGATATATCTATCCTTTTTTTGTTATGTTACCATTTATTTTCCCTGTAATAATATCTATGTGTTTCTTTTTAAGTGTATTTTTTTTAAATATGTATTATGAAAGCTTTTCAAAGTTATATAAAAAGATCAGTGAACTAAAGAATGAATTCATAAACTCCTCAGAAAATGACAATGTGAACGAAAGAATATTGGTATCCGAAACTTCTAATCATTTATGTTTAAATGAAAGTAACGATAAAGTATCTAATACATCAGATGATTTTTTATCAAGAAATAATAGCAACATATCTAGTTCAAAAAGTGAAATGATAAATTCAAATTTTGTATTTAATAAACTTTTGAATTTTTATTTTTCATTTGCAGTTTTCTTTTCTTATTTAGGTTCTGCATTTTTACATATATCCTTAGGTGAAATTATATGTATCGCATTATTAACATTTTATCAAATTGTAAAGCATACCAATAATTTGAATATCACAATTTTACTTAAGAGTGAGAAAATTAAATTTTGTAAATTCCTTTTGTTTATATTATATGGGTTGTTATGCTTTTCCATAAATTTGTATGTAAACCAATGGGAGGAATATATAACGAAATTAAAAAGGTTAAAAAGACGTATTTTATTGTTTGAAAAAAATAAATTTAGTGAGATTGTGGATCTGAATACTCAAAAGGGTGATGGTGACCATTTTGATGAAACACAAATATTTTCCATATTTTTTTCTTTTTTAATAAAAAAAAATGAGGGTTCAAAAATGAGGGATAATGATATGAATAGTGATAGCGAAGACAGTATATATGATGCTTATGAGCAGCAGATACAATTACACCATGGTGATAATATGGTCAATGGTATGTTGATGATGAGAAGGATATCGATGCAAAATTTAGAGGACGACGAAACCCAAGTTGAATATATAAATAGGGAGATCCATACGCAAGGTGATTTACATGTACGAAGAACGAATCAAGGTATTTTAAGATTTAATATGAGAAGGGGCAAAAAAGGGTCCAATGAAAATATGGGTGTCCACCATGAGAGTGGTAATGTGGATGATGCGAATGGTATGAATAATGTGGATGATACGAATAATATGAATAATGTGGATGGTACGAATAATATGAATAATGTGGATGGTACGAATAATATGAATAATATGGATGGTAGGAATAATATGAATAATATAAATAGTGTGGATAATATGAACAATTTAAATAATAATGATGGTGAAGAAGAAGAAGAATGTGTGAACGATGTTTTAAATTATGATAATAATAATTATGCTATTAATGAAGATGCTGAGGAATATATAAAAAATACGTCAGGAGAACGAGCTGTTATTATTTGTTCTGAAAAAAGAATATATGAAAAGAACGGAAATGGTGATATAATAACTAGAAATTATAAAAATGAAGAACGATATATATATTTAAAAAAATGGATTCCTTTTAAATCTATGATATTGTCAAAATTAGAAAAGAGGAAAAGGAATAGAAAAGAGGCATATAATACTCCTAGAGTATTAATATTAATACATTCCTTTTTATTTATACTTATTGTTTTCATATTTTTAATGGTATTCTTTAAAAAAGAGCCTATTTTTCGTTTTAATATGCCATCAGTTAATAAGCGTTTAAATAATTTTTTTAAATCAACAAGTTTTCACGAAATTATACCAAATAGTGTAGGGAAATGTAAAACAAAAAAATATATAGCAAAAGAACCATGTTTTAATGTGGGTCATATATACCATGAAGAGAAGACATTTTATCATGCAACTCTTTTATTTTTACAAGGTTTAAGATCTGTAAAAATTATGAATATGAATTTTTATTATGAAAAAGGCATATATTATTTATCTTTGGATGGATATTTTAAACATATTATAGGTCCTCTCTTTCTTAAATTATGTTTAGGTACAAACTTTTGTCCCATAAGTACATATGCATTTTTAGTAGGAAGTAAACCAACTTTCTCAGTAAATGTTGCTGTTCAATGTAATAATAAAAAACCACCATATTATATGACGGATATTATTGTAAAAGATTTAAAAATTACAAAAATAGAAATAGTTAAACATTCAGATGTTATCGATAATGTGGATATTAAATTAGATGATGTACAAGATAGAGTACAAGAAAAGGTTAATGCCATGTTAGAAGCAAAAAAAAAAATTATCGTCTGGAAAAATCAAAAGTATCATTTGGAGGGTTTTCTCAATTATTTAATATCTAAAAATGCATTATCTGGGTTTTCCTGTGAACCCATAAATTATTGAacttgacaaattaaaacacgttaaatgtatatatgtatatatattatatatgtatatatttatttatatttttcttacatttccatttgggtgaacatatattcttacccttttcataaattgttttttttttttttgatatttatcaataatatattcagtaaaaatatattatttcacataatttaatattttttggaataacatataaatgtcaatgtatatgtgtatatatatatatatatatatatatgtatatatttttttttctttttcatatatttatatgcgcccatttgtttttatctatatatatatatatttatatatccgaatttctttatattgttatattccgatttgattatatataaaattttttttaattcataatttttttttttttttttttcttttttttcatttccatgtgatttatttcaatccgatattatttatatgttatatatatattattctataatatgataaaaggatatttttttttcttatttgtatttttagattataaatataaagatattcttatatatagattatatctgcttttttttttttttttttttttttttctattattaatttttaaattatctacttacgaattttttaattgttgatatggccttcatgagtatattattatgcatgaaagaaaaaaaaaaaaaaaaatatatatatgtatataatatatatatatatatatataatataccaatttttattatcttatattttacatgcactataaatgcatagaatttttcttttttttttttttgatttcttctccccaaatttttggttatttatataatatacgtattataaatatgtacatataatatgtatatattataatgtgtaattttccacatatatatataaaatatatatatatatatatatatattttaataatacttgaacttttaaaattaaatatataaaaaggaatccaattaatttatggttatattatttgttattcactagaaaattctctagaatttaataaaacatatat

**5’ amplicon size: 1149bps**

**3’ amplicon size: 1116bps**

### > mChSEP-PF3D7_0214800 mCh SEP yDHODH recodonizedsequence

cattttattccgtaaaattaaaatgaattccaccaaaggtttaacatatacataatatatatttatatatatatatatatgtatatatatatatataaaataaaaataataatatataatatatttttatattacattattatattatattatattatattatattttattttatttttttaatatagaacttcatgtagctaaaataaaaatgagttcacatttttcgttaattaaatatattatatatatatatatatgcaaaattggtattatatataaatataatatataatttgtttccttttttttagtttttaaaatatagatttataaataatttttttcttttttttaagaattattttgatttaccatatatattatttaaaatcccaagttaaaattaaaaaaaaaaaggagtttttattataatattatatatatttatatatatattttaactactttaattttaaaagtgtgagagtaagaaaatgaaaaaaaaaaaaaaaaaaaaaaaaaggattattttttatttatatattataatatatataatatattaataataaataaaataaacatgtaaataaatattatatatatatatcataatatagtatcatgaaataatatatatttataaaaatattattataatatatagttaattattatattatatattacataatatttataatatattatatattggattatagttttacctattgggggatcttggttctataaaaaaatttcaatttttttttaccctttatttttttttgttcactatatatatatattgttatatataacataatatattaatagtttcataatattatatgtatatagaaagaaaaacatatatatatatatatatatattgttcattatttctaaatatatatattattatatatattatataaaaaatatattattttagatttactcctttatatataaattaatattttttatagatttcgattttttcctatttacataatttttaattttttataattttaatttttctacattatatatgtattatatatatatataaataatctatatatttttaaaataattcaatgtcattttattaattcaattaaatacaaattaatttcttattttctttaattatatatatatatattataatatttatagaacctatattatagatttaatttttaatctgttaaaagaaaaaaagtacaaaaaattatacatttatttacatatatatatatatatatatatatatatgtataatttttaatcttttatttttagaggtattttatgttattttttattttacatatgttgttgtataattttttttgtatatgcccttatattgattttcttttttttttttaaataatatagtgtttgctttaataaaaaaaaggaaatatcaggattttattgaatacatacatatatatatatatatatatatatatatatatatatatatatatatataactattagaattcccatagatgaatatataaacatataataatattatatctataatatttatatttttattatacatacatactgaatatattttaaagttacgaataatgaatgctttttttttttttttttttttgttgctttaaataaatggtaatcataaatgaaattagtctagagctaaaatgaaaattaaagaaaatataaaaagtatataataataaatgtgttataatatttaaaataaaacaaaaaaaaaaaaaaaaaaaaaaaaaaaagtaaattacaataatttattctataaagttcagaATG**TTGGAATTATTACGTAATGCTTATAATAAATGTAAAGAGTATATTTTGGGCTTATCGAAAAAATCACAAATATATAAAATAAATATATTAAATTATTGTTTATTATTTTCGTTTTTTTTATTTTATGCATCATATTTTTTTTTTATAAATGGAATAATAAATCCAGCAGTAAAAATTGTTTTCCATGGAA*gtaagtaccttatatgaaaaaatgaaagtgtatccataaaagaagatatacacaaaggtgtatatatatatatatatatatatatatatatatatatgtatgtatgtatgtatttatttatttatttattatttatatatatattttctttatattttctttaag*GTATTATTGAGGACAAAACCAAAGTTTACGAGAACTCGATAATTGAATCAATATGGCTATTGTTTACCAAGGGAAGTTATTTTAATATGGCCATGCTTCTATTCTTTTCTATTATAATACCGTTTCTCAAGTTACTTATGGTTAGTGATAACTTTTATAGTTTTATCGTATTATATAAAATGAATAAAAAACGAGAAGAGGAAAAGAGAAGGAGACGAAGAAGAATACGTGCACGAGAATATAACAATGCTAAATATAACAAGAATAGATATAAAATGAATAAATATAATAGAAATAAGAATATTAATGATATATACAAGGATGATATATATTATAGTGAAAATATTTTTAAGAATGATGAGATAAATTATATTAATACAATAAATGAGAATGAAGAATTTATTTTAAAGAAATTTAAAATTTTAAATTTCATATCGCGTTTTCAATTTGTTGATGTATTTATATCTTTATTTATAGTATCATCATTAAATTTATATTTATTAGAGGCAAGGATGTTAAATGGTGC**AGGTTTAAACGAGCAGAAGTTAATATCAGAAGAGGATTTGGGTGAACAAAAACTCATAAGCGAAGAAGATTTAATAACTTCGTATAGCATACATTATACGAAGTTATCCGGAGAAGGAAGAGGAAGTTTATTAACATGTGGAGATGTAGAAGAAAATCCAGGACCAATGACAGCCAGTTTAACTACCAAGTTCTTGAACAATACCTATGAAAACCCATTTATGAATGCATCCGGTGTTCATTGCATGACTACACAAGAATTAGATGAATTAGCAAACTCTAAAGCTGGCGCATTCATTACAAAGAGTGCTACAACCTTAGAAAGAGAAGGTAACCCTGAACCACGTTACATTTCTGTCCCTCTAGGCAGTATCAACTCCATGGGTTTACCAAACGAAGGTATCGACTACTATTTGTCCTATGTATTAAACCGTCAAAAGAATTATCCTGATGCACCTGCTATTTTCTTCTCAGTTGCTGGTATGAGCATTGATGAAAATTTAAATTTGTTGAGGAAAATCCAAGATAGCGAATTCAACGGTATTACCGAGTTAAACTTGTCTTGTCCTAATGTGCCTGGGAAACCACAAGTTGCTTATGACTTTGACTTGACAAAGGAAACCTTGGAAAAGGTTTTTGCCTTTTTCAAAAAACCTCTTGGTGTCAAGTTGCCTCCTTATTTTGATTTTGCCCATTTTGATATCATGGCAAAAATATTGAACGAGTTCCCATTAGCTTATGTCAACTCTATCAATAGTATAGGAAATGGTCTTTTCATTGATGTGGAGAAGGAGAGTGTAGTAGTGAAGCCAAAGAATGGTTTCGGGGGTATTGGAGGTGAATATGTTAAGCCAACCGCGCTCGCCAATGTTCGTGCATTTTACACTCGTTTGAGACCTGAAATCAAAGTTATCGGTACAGGTGGAATTAAGTCCGGTAAGGATGCATTTGAACATCTTCTATGTGGTGCCTCTATGCTACAGATTGGTACAGAATTACAAAAAGAGGGCGTCAAGATTTTTGAACGTATCGAAAAAGAATTAAAAGACATAATGGAAGCTAAGGGTTATACATCCATAGATCAGTTCCGTGGGAAGTTGAACAGCATTGGTGAAGGTAGAGGTTCTTTGTTGACTTGTGGTGATGTTGAAGAAAATCCAGGTCCAGCTAGCATGGTGAGCAAGGGCGAGGAGGATAACATGGCCATCATCAAGGAGTTCATGCGCTTCAAGGTGCACATGGAGGGCTCCGTGAACGGCCACGAGTTCGAGATCGAGGGCGAGGGCGAGGGCCGCCCCTACGAGGGCACCCAGACCGCCAAGCTGAAGGTGACCAAGGGTGGCCCCCTGCCCTTCGCCTGGGACATCCTGTCCCCTCAGTTCATGTACGGCTCCAAGGCCTACGTGAAGCACCCCGCCGACATCCCCGACTACTTGAAGCTGTCCTTCCCCGAGGGCTTCAAGTGGGAGCGCGTGATGAACTTCGAGGACGGCGGCGTGGTGACCGTGACCCAGGACTCCTCCCTGCAGGACGGCGAGTTCATCTACAAGGTGAAGCTGCGCGGCACCAACTTCCCCTCCGACGGCCCCGTAATGCAGAAGAAGACCATGGGCTGGGAGGCCTCCTCCGAGCGGATGTACCCCGAGGACGGCGCCCTGAAGGGCGAGATCAAGCAGAGGCTGAAGCTGAAGGACGGCGGCCACTACGACGCTGAGGTCAAGACCACCTACAAGGCCAAGAAGCCCGTGCAGCTGCCCGGCGCCTACAACGTCAACATCAAGTTGGACATCACCTCCCACAACGAGGACTACACCATCGTGGAACAGTACGAACGCGCCGAGGGCCGCCACTCCACCGGCGGCATGGACGAGCTGTACAAGACGCGTGTTATCGGAGCAGGTGCAGGTATGGGAAGTAAAGGAGAAGAACTTTTCACTGGAGTTGTCCCAATTCTTGTTGAATTAGATGGTGATGTTAATGGGCACAAATTTTCTGTCAGTGGAGAGGGTGAAGGTGATGCAACATACGGAAAACTTACCCTTAAATTTATTTGCACTACTGGAAAACTACCTGTTCCTTGGCCAACACTTGTCACTACTTTAACTTATGGTGTTCAATGCTTTTCAAGATACCCAGATCATATGAAACGGCATGACTTTTTCAAGAGTGCCATGCCCGAAGGTTATGTACAGGAAAGAACTATATTTTTCAAAGATGACGGGAACTACAAGACACGTGCTGAAGTCAAGTTTGAAGGTGATACCCTTGTTAATAGAATCGAGTTAAAAGGTATTGATTTTAAAGAAGATGGAAACATTCTTGGACACAAATTGGAATACAACTATAACGATCACCAGGTGTACATCATGGCAGACAAACAAAAGAATGGAATCAAAGCTAACTTCAAAATTAGACACAACATTGAAGATGGAGGCGTTCAACTAGCAGACCATTATCAACAAAATACTCCAATTGGCGATGGGCCCGTCCTTTTACCAGACAACCATTACCTGTTTACAACTTCTACTCTTTCGAAAGATCCCAACGAAAAGAGAGACCACATGGTCCTTCTTGAGTTTGTAACAGCTGCTGGGATTACACATGGCATGGATGAACTATACAAAGATAACACGCGTGCTAGAGGTGCTGCTGCTGGTGCTGGAGGTGCAGGTAGACCTAGGATGCTTGAGCTTCTTAGAAACGCATACAACAAGTGCAAGGAATACATACTTGGACTTAGTAAGAAGAGTCAGATTTACAAGATTAACATTCTTAACTACTGCCTTCTTTTCAGTTTCTTCCTTTTCTACGCTAGTTACTTCTTCTTCATTAACGGTATTATTAACCCTGCTGTTAAGATAGTATTTCACGGTTCAATAATAGAAGATAAGACAAAGGTATATGAAAATAGTATTATAGAGAGTATTTGGTTGCTTTTCACAAAAGGTTCATACTTCAACATGGCAATGTTATTGTTTTTCAGCATAATTATTCCATTCTTAAAACTTTTAATGGTATCAGACAATTTCTACTCATTCATAGTTCTTTACAAGATGAACAAGAAGAGGGAGGAAGAGAAACGTCGTCGTAGGCGTCGTATTAGAGCTAGGGAGTACAATAACGCAAAGTACAATAAAAACCGTTACAAGATGAACAAGTACAACCGTAACAAAAACATAAACGACATTTATAAAGACGACATTTACTACTCAGAGAACATATTCAAAAACGACGAAATTAACTACATAAACACTATTAACGAAAACGAGGAGTTCATACTTAAAAAGTTCAAGATACTTAACTTTATTAGTAGATTCCAGTTCGTAGACGTTTTCATTAGCCTTTTCATTGTTAGTAGTCTTAACCTTTACCTTCTTGAAGCTCGTATGCTTAACGGAGCTTACTACTTCCTTAACTACTGCATGCTTAGTACTATTAGTAGCTTCCTTCTTTTCTCATTCACAAGCCTTAAGATTCACATTTTCAAGAACGGTAACATAAAAATTAGCGCTTGCTTAAACGAGAGTAACTTGGAAGTAACAACTTCAGGACCATTAAGTACTAAGGACTTAGTTGAGGAGGAGGGTCACGCTCAGATTAACAACTTGATAATAAACGACAAAAACATGACATCAGGAGTAGTAAACGACTTCAGTGGAAACGGTAATAACGTTGAGCTTACTGATGATTTAAAATCAGGAGAGCCACAGAATCGTGACGACATTCAGACAGAGGAGACTAAGAAGGAGAAGATGAACACACGTACACACAACGACGAGGACAACACTAAGAAGAAGAACATTAAAGACAAGAAAAAGGCAAATGGAGACGACAAGGTTATTCAGAAGTGCATAGACAACGAGCGTAAGAAGAAGCAGAACGGAATGATACAGTCAGTTAATGACGGAGACAAGAACTCAAACTTCAACAACAATAACAACAACAACATAAACGGAGACTCAAACAACAACAACATAAACGGAGACTCAAACAACAACAACATAAACGGAGACTCAAACAACAACAACATAAACGGAGACTCAAACAACAACAACATAAACGGAGACTCAAACAACAACAACATAAACGGAGACTCAAACAACAACAACTACCACAACAACTACCACAACAACTACAGAAACAACTACCACAACAACTACAGAAACAACAACTGCCGTAACAACATACTTGAGCAGAACAAGTGCGACAAGAACGTACTTTGCTACAACAACATTTACAACACTATGAAGGACAACGACACATACATTTACCTTAAGAAAAACAAGTTCAACAGTCTTCTTAAGTCAAACTGCATAAAGACAAACTTCAACATGATTAAAATTGGATACGTTATTTTCCTTTTCGTTCTTCTTTGCCTTTGCATTTACCTTATTACTGGTGTTGAGTGTTCACTTTTCGGAATTTACATTTACCTTTCATACTTCAACTTCAACATAGAGGGTATTCTTATAGACTACATGGACATGCTTAACATACTTAAGCTTAAGATTAAAAAGGGTTACATATACCCATTCTTCGTAATGCTTCCTTTCATATTTCCAGTTATTATTAGCATGTGCTTTTTCCTTTCAGTTTTCTTCCTTAACATGTACTACGAGTCATTCAGTAAACTTTACAAGAAAATATCAGAGTTGAAAAACGAGTTTATTAATAGTAGTGAGAACGATAACGTTAATGAGCGTATTCTTGTTAGTGAGACAAGCAACCACCTTTGCCTTAACGAGTCAAATGACAAGGTTAGCAACACTAGTGACGACTTCCTTAGTCGTAACAACTCAAATATTAGCTCAAGTAAGTCAGAGATGATTAACAGTAACTTCGTTTTCAACAAGTTACTTAACTTCTACTTCAGTTTCGCTGTATTTTTCAGCTACCTTGGAAGCGCTTTCCTTCACATTAGTCTTGGAGAGATAATTTGCATAGCTCTTCTTACTTTCTACCAGATAGTTAAACACACAAACAACCTTAACATAACTATACTTTTAAAATCAGAAAAGATAAAGTTCTGCAAGTTTTTACTTTTCATTCTTTACGGACTTCTTTGTTTCAGTATTAACCTTTACGTTAATCAGTGGGAAGAGTACATTACAAAGCTTAAGCGTCTTAAGCGTAGAATACTTCTTTTCGAGAAGAACAAGTTCTCAGAAATAGTTGACTTAAACACACAGAAAGGAGACGGAGATCACTTCGACGAGACTCAGATTTTCAGTATTTTCTTCAGCTTCCTTATTAAGAAGAACGAAGGAAGTAAGATGCGTGACAACGACATGAACTCAGACTCAGAGGATTCAATTTACGACGCATACGAACAACAAATTCAGCTTCATCACGGAGACAACATGGTTAACGGAATGCTTATGATGCGTCGTATTAGTATGCAGAACCTTGAAGATGATGAGACACAGGTAGAGTACATTAACCGTGAAATACACACACAGGGAGACCTTCACGTTAGGCGTACAAACCAGGGAATACTTCGTTTCAACATGCGTCGTGGAAAGAAGGGAAGTAACGAGAACATGGGAGTTCATCACGAATCAGGAAACGTTGACGACGCAAACGGAATGAACAACGTTGACGACACAAACAACATGAACAACGTTGACGGAACAAACAACATGAACAACGTTGACGGAACAAACAACATGAACAACATGGACGGACGTAACAACATGAACAACATTAACTCAGTTGACAACATGAATAACCTTAACAACAACGACGGAGAGGAGGAGGAGGAGTGCGTTAATGACGTACTTAACTACGACAACAACAACTACGCAATAAACGAGGACGCAGAAGAGTACATTAAGAACACAAGTGGTGAGAGGGCAGTAATAATATGCAGCGAGAAGCGTATTTACGAGAAAAATGGTAACGGAGACATTATTACACGTAACTACAAGAACGAGGAGAGGTACATTTACCTTAAGAAGTGGATACCATTCAAGAGCATGATTCTTAGTAAGCTTGAGAAACGTAAGCGTAACCGTAAGGAAGCTTACAACACACCACGTGTTCTTATTCTTATTCACAGTTTCCTTTTCATTTTAATAGTATTTATTTTCCTTATGGTTTTTTTCAAGAAGGAACCAATATTCAGATTCAACATGCCTAGTGTAAACAAAAGACTTAACAACTTCTTCAAGAGTACTTCATTCCATGAGATAATTCCTAACTCAGTTGGAAAGTGCAAGACTAAGAAGTACATTGCTAAGGAGCCTTGCTTCAACGTTGGACACATTTATCACGAGGAAAAAACTTTCTACCACGCTACATTACTTTTCCTTCAGGGACTTCGTAGCGTTAAGATAATGAACATGAACTTCTACTACGAGAAGGGAATTTACTACCTTAGCCTTGACGGTTACTTCAAGCACATAATTGGACCATTATTCTTAAAGCTTTGCCTTGGAACTAATTTCTGCCCAATTTCAACTTACGCTTTCCTTGTTGGTTCAAAGCCTACATTTAGTGTTAACGTAGCAGTACAGTGCAACAACAAGAAGCCTCCTTACTACATGACAGACATAATAGTTAAGGACCTTAAGATAACTAAGATTGAGATTGTAAAGCACAGTGACGTAATAGACAACGTTGACATAAAGCTTGACGACGTTCAGGACCGTGTTCAGGAGAAAGTAAACGCAATGCTTGAGGCTAAGAAGAAGATAATAGTTTGGAAGAACCAGAAATACCACCTTGAAGGATTCTTAAACTACCTTATTAGCAAGAACGCTCTTAGCGGATTCAGTTGCGAGCCAATTAACTACTAGAGGCCTATAACTTCGTATAGCATACATTATACGAAGTTATTATGACTCGAGGGATATGGCAGCTTAATGTTCGTTTTTCTTATTTATATATTTATACCAATTGATTGTATTTATAACTGTAAAAATGTGTATGTTGTGTGCATATTTTTTTTTGTGCATGCACATGCATGTAAATAGCTAAAATTATGAACATTTTATTTTTTGTTCAGAAAAAAAAAACTTTACACACATAAAATGGCTAGTATGAATAGCCATATTTTATATAAATTAAATCCTATGAATTTATGACCATATTAAAAATTTAGATATTTATGGAACATAATATGTTTGAAACAATAAGACAAAATTATTATTATTATTATTATTTTTACTGTTATAATTATGTGTCTCCTTCAATGATTCATAAATAGTTGGACTTGATTTTTAAAATGTTTATAATATGATTAGCATAGTTAAATAAAAAAAGTTGAAAAATTAAAAAAAAACATATAAACACAAATGATGGTTTTTCCTTCAATTTCGATATCAATTTATAGAAACAAAATATATACTTGTATAATTTTATTTTTTTATATAAATCATTACATATATAATTATACAATATTTTTTCTAAGAGATAATTATATATTAATATATATAAAAAAAGGTGTTTTTTTTTTTTTTTTTTATTTTTATTTTTATTTTATGGTAATATTTTATTTTCCTTATTTTATAAATTATATTAGTTTATATGTGATTAATTTTATATATTATCAATTTATATATTTTTAAATGCTTACTTAATTATCTTTTTTTTTTTTTTTTTTTTTTTTTCCCCTCTTTTTATATTAATTTATTTTTGAAAAAATTGATATATATATATATATATAATATATATATATACATGTAGTAGTATTAAACAATGTATAATATATATAAATAATATATTTATATATTTCATTTCAATTTTAATTTTTTTTGGTTTTTTTTTTTTTTCTTTTTGTCATATTTAAAAAAAATTATATTCATATAAGTTATGCATTTTTTATAAACATTATTCAATATATGTATAATATAATATATATATATATATTAATGTATTATTCCAATGTGCATGATAAAAGAAAAAAATAATATTTATAAAAAAAAAGAAAAATAAAACAAAAAAAGAAAAAAAAAAAAAAAAAAAAAAAAATACAAAAATAAATAATATAATTTATAATTATATATTCTTGTCACAATAAAAATATATATATATATATATATATTTATAATATGTATATTTTAAACTAGAAAAGGAATAACTAATATTTTATTTATTATCATTCAAGATTTATATTTTATAATAATAAATACCTAATAGAAATATATCAGGATCCATGCATGGTTCGCTAAACTGCATCGTCGCTGTGTCCCAGAACATGGGCATCGGCAAGAACGGGGACTACCCCTGGCCACCGCTCAGGAACGAATTTAGATATTTCCAGAGAATGACCACAACCTCTTCAGTAGAAGGTAAACAGAATCTGGTGATTATGGGTAAGAAGACCTGGTTCTCCATTCCTGAGAAGAATCGACCTTTAAAGGGTAGAATTAATTTAGTTCTCAGCAGAGAACTCAAGGAACCTCCACAAGGAGCTCATTTTCTTTCCAGAAGTCTAGATGATGCCTTAAAACTTACTGAACAACCAGAATTAGCAAATAAAGTAGACATGGTCTGGATAGTTGGTGGCAGTTCTGTTTATAAGGAAGCCATGAATCACCCAGGCCATCTTAAACTATTTGTGACAAGGATCATGCAAGACTTTGAAAGTGACACGTTTTTTCCAGAAATTGATTTGGAGAAATATAAACTTCTGCCAGAATACCCAGGTGTTCTCTCTGATGTCCAGGAGGAGAAAGGCATTAAGTACAAATTTGAAGTATATGAGAAGAATGATTAAGCTTATTTAATAATAGATTAAAAATATTATAAAAATAAAAACATAAACACAGAAATTACAAAAAAAATACATATGAATTTTTTTTTTGTAATCTTCCTTATAAATATAGAATAATGAATCATATAAAACATATCATTATTCATTTATTTACATTTAAAATTATTGTTTCAGTATCTTTAATTTATTATGTATATATAAAAATAACTTACAATTTTATTAATAAACAATATATGTTTATTAATTCATGTTTTGTAATTTATGGGATAGCGATTTTTTTTACTGTCTGTATTTTTCTTTTTTAATTATGTTTTAATTGTATTTTATTTTTATTATTGTTCTTTTTATAGTATTATTTTAAAACAAAATGTATTTTCTAAGAACTTATAATAATAATAAATATAAATTTTAATAAAAATTATATTTATCTTTTACAATATGAACATAAAGTACAACATTAATATATAGCTTTTAATATTTTTATTCCTAATCATGTAAATCTTAAATTTTTCTTTTTAAACATATGTTAAATATTTATTTCTCATTATATATAAGAACATATTTATTAAATCTAGAATTCTATAGTGAGTCGTATTACAATTCACTGGCCGTCGTTTTACAACGTCGTGACTGGGAAAACCCTGGCGTTACCCAACTTAATCGCCTTGCAGCACATCCCCCTTTCGCCAGCTGGCGTAATAGCGAAGAGGCCCGCACCGATCGCCCTTCCCAACAGTTGCGCAGCCTGAATGGCGAATGGCGCCTGATGCGGTATTTTCTCCTTACGCATCTGTGCGGTATTTCACACCGCATATGGTGCACTCTCAGTACAATCTGCTCTGATGCCGCATAGTTAAGCCAGCCCCGACACCCGCCAACACCCGCTGACGCGCCCTGACGGGCTTGTCTGCTCCCGGCATCCGCTTACAGACAAGCTGTGACCGTCTCCGGGAGCTGCATGTGTCAGAGGTTTTCACCGTCATCACCGAAACGCGCGAGACGAAAGGGCCTCGTGATACGCCTATTTTTATAGGTTAATGTCATGATAATAATGGTTTCTTAGACGTCAGGTGGCACTTTTCGGGGAAATGTGCGCGGAACCCCTATTTGTTTATTTTTCTAAATACATTCAAATATGTATCCGCTCATGAGACAATAACCCTGATAAATGCTTCAATAATATTGAAAAAGGAAGAGTATGAGTATTCAACATTTCCGTGTCGCCCTTATTCCCTTTTTTGCGGCATTTTGCCTTCCTGTTTTTGCTCACCCAGAAACGCTGGTGAAAGTAAAAGATGCTGAAGATCAGTTGGGTGCACGAGTGGGTTACATCGAACTGGATCTCAACAGCGGTAAGATCCTTGAGAGTTTTCGCCCCGAAGAACGTTTTCCAATGATGAGCACTTTTAAAGTTCTGCTATGTGGCGCGGTATTATCCCGTATTGACGCCGGGCAAGAGCAACTCGGTCGCCGCATACACTATTCTCAGAATGACTTGGTTGAGTACTCACCAGTCACAGAAAAGCATCTTACGGATGGCATGACAGTAAGAGAATTATGCAGTGCTGCCATAACCATGAGTGATAACACTGCGGCCAACTTACTTCTGACAACGATCGGAGGACCGAAGGAGCTAACCGCTTTTTTGCACAACATGGGGGATCATGTAACTCGCCTTGATCGTTGGGAACCGGAGCTGAATGAAGCCATACCAAACGACGAGCGTGACACCACGATGCCTGTAGCAATGCCAACAACGTTGCGCAAACTATTAACTGGCGAACTACTTACTCTAGCTTCCCGGCAACAATTAATAGACTGGATGGAGGCGGATAAAGTTGCAGGACCACTTCTGCGCTCGGCCCTTCCGGCTGGCTGGTTTATTGCTGATAAATCTGGAGCCGGTGAGCGTGGGTCTCGCGGTATCATTGCAGCACTGGGGCCAGATGGTAAGCCCTCCCGTATCGTAGTTATCTACACGACGGGGAGTCAGGCAACTATGGATGAACGAAATAGACAGATCGCTGAGATAGGTGCCTCACTGATTAAGCATTGGTAACTGTCAGACCAAGTTTACTCATATATACTTTAGATTGATTTAAAACTTCATTTTTAATTTAAAAGGATCTAGGTGAAGATCCTTTTTGATAATCTCATGACCAAAATCCCTTAACGTGAGTTTTCGTTCCACTGAGCGTCAGACCCCGTAGAAAAGATCAAAGGATCTTCTTGAGATCCTTTTTTTCTGCGCGTAATCTGCTGCTTGCAAACAAAAAAACCACCGCTACCAGCGGTGGTTTGTTTGCCGGATCAAGAGCTACCAACTCTTTTTCCGAAGGTAACTGGCTTCAGCAGAGCGCAGATACCAAATACTGTCCTTCTAGTGTAGCCGTAGTTAGGCCACCACTTCAAGAACTCTGTAGCACCGCCTACATACCTCGCTCTGCTAATCCTGTTACCAGTGGCTGCTGCCAGTGGCGATAAGTCGTGTCTTACCGGGTTGGACTCAAGACGATAGTTACCGGATAAGGCGCAGCGGTCGGGCTGAACGGGGGGTTCGTGCACACAGCCCAGCTTGGAGCGAACGACCTACACCGAACTGAGATACCTACAGCGTGAGCTATGAGAAAGCGCCACGCTTCCCGAAGGGAGAAAGGCGGACAGGTATCCGGTAAGCGGCAGGGTCGGAACAGGAGAGCGCACGAGGGAGCTTCCAGGGGGAAACGCCTGGTATCTTTATAGTCCTGTCGGGTTTCGCCACCTCTGACTTGAGCGTCGATTTTTGTGATGCTCGTCAGGGGGGCGGAGCCTATCGAAAAACGCCAGCAACGCGGCCTTTTTACGGTTCCTGGCCTTTTGCTGGCCTTTTGCTCACATGTTCTTTCCTGCGTTATCCCCTGATTCTGTGGATAACCGTATTACCGCCTTTGAGTGAGCTGATACCGCTCGCCGCAGCCGAACGACCGAGCGCAGCGAGTCAGTGAGCGAGGAAGCGGAAGAGCGCCCAATACGCAAACCGCCTCTCCCCGCGCGTTGGCCGATTCATTAATGCAGCTGGCACGACAGGTTTCCCGACTGGAAAGCGGGCAGTGAGCGCAACGCAATTAATGTGAGTTAGCTCACTCATTAGGCACCCCAGGCTTTACACTTTATGCTTCCGGCTCGTATGTTGTGTGGAATTGTGAGCGGATAACAATTTCACACAGGAAACAGCTATGACCATGATTACGCCAAGCTATTTAGGTGACACTATAGAATACTCGCGGCCGCTAGTTGGAATTATTACGTAATGCTTATAATAAATGTAAAGAGTATATTTTGGGCTTATCGAAAAAATCACAAATATATAAAATAAATATATTAAATTATTGTTTATTATTTTCGTTTTTTTTATTTTATGCATCATATTTTTTTTTTATAAATGGAATAATAAATCCAGCAGTAAAAATTGTTTTCCATGGAAGTAAGTACCTTATATGAAAAAATGAAAGTGTATCCATAAAAGAAGATATACACAAAGGTGTATATATATATATATATATATATATATATATATATATGTATGTATGTATGTATTTATTTATTTATTTATTATTTATATATATATTTTCTTTATATTTTCTTTAAGGTATTATTGAGGACAAAACCAAAGTTTACGAGAACTCGATAATTGAATCAATATGGCTATTGTTTACCAAGGGAAGTTATTTTAATATGGCCATGCTTCTATTCTTTTCTATTATAATACCGTTTCTCAAGTTACTTATGGTTAGTGATAACTTTTATAGTTTTATCGTATTATATAAAATGAATAAAAAACGAGAAGAGGAAAAGAGAAGGAGACGAAGAAGAATACGTGCACGAGAATATAACAATGCTAAATATAACAAGAATAGATATAAAATGAATAAATATAATAGAAATAAGAATATTAATGATATATACAAGGATGATATATATTATAGTGAAAATATTTTTAAGAATGATGAGATAAATTATATTAATACAATAAATGAGAATGAAGAATTTATTTTAAAGAAATTTAAAATTTTAAATTTCATATCGCGTTTTCAATTTGTTGATGTATTTATATCTTTATTTATAGTATCATCATTAAATTTATATTTATTAGAGGCAAGGATGTTAAATGGTGCATATTATTTTTTGAATTATTGTATGTTATCAACAATATCATCTTTTTTGTTATTTAGTTTTACTTCTTTAAAAATACATATATTTAAAAATGGAAATATTAAGATATCTGCATGTCTCAATGAATCAAATCTAGAGGTTACTACAAGCGGTCCCCTTTCAACAAAAGATCTTGTCGAAGAAGAAGGACATGCACAAATAAATAATCTAATTATTAATGATAAGAATATGACTAGTGGTGTTGTTAATGATTTTTCAGGTAATGGAAACAATGTCGAATTAACAGACGACCTTAAGAGTGGCGAACCCCAAAACAGAGATGATATACAAACTGAAGAAACAAAAAAAGAAAAAATGAATACTAGAACTCATAATGATGAAGATAATACAAAAAAAAAAAATATAAAGGATAAAAAGAAAGCTAACGGTGATGATAAAGTAATACAAAAATGTATTGATAATGAAAGGAAAAAAAAACAAAATGGTATGATTCAAAGTGTAAACGATGGTGATAAAAATAGTAATTTTAATAATAACAATAATAATAATATTAATGGTGATAGTAATAATAATAATATTAATGGTGATAGTAATAATAATAATATTAATGGTGATAGTAATAATAATAATATTAATGGTGATAGTAATAATAATAATATTAATGGTGATAGTAATAATAATAATATTAATGGTGATAGTAATAATAATAATTATCATAATAATTATCATAATAATTATCGTAATAATTATCATAATAATTATCGTAATAATAATTGTAGGAATAATATTTTAGAACAAAATAAATGTGATAAAAATGTTTTGTGTTATAATAATATATATAATACAATGAAAGATAATGATACTTATATATATTTAAAAAAGAATAAATTTAATTCGTTATTAAAAAGTAATTGTATCAAAACTAATTTTAATATGATAAAGATAGGTTATGTAATATTTTTATTTGTGTTATTATGTTTATGTATATATTTAATAACAGGAGTAGAATGCAGTTTATTTGGTATATATATATATTTGAGTTATTTTAATTTTAATATTGAAGGAATATTAATTGATTATATGGATATGTTAAATATTTTAAAATTAAAAATAAAGAAAGGATATATCTATCCTTTTTTTGTTATGTTACCATTTATTTTCCCTGTAATAATATCTATGTGTTTCTTTTTAAGTGTATTTTTTTTAAATATGTATTATGAAAGCTTTTCAAAGTTATATAAAAAGATCAGTGAACTAAAGAATGAATTCATAAACTCCTCAGAAAATGACAATGTGAACGAAAGAATATTGGTATCCGAAACTTCTAATCATTTATGTTTAAATGAAAGTAACGATAAAGTATCTAATACATCAGATGATTTTTTATCAAGAAATAATAGCAACATATCTAGTTCAAAAAGTGAAATGATAAATTCAAATTTTGTATTTAATAAACTTTTGAATTTTTATTTTTCATTTGCAGTTTTCTTTTCTTATTTAGGTTCTGCATTTTTACATATATCCTTAGGTGAAATTATATGTATCGCATTATTAACATTTTATCAAATTGTAAAGCATACCAATAATTTGAATATCACAATTTTACTTAAGAGTGAGAAAATTAAATTTTGTAAATTCCTTTTGTTTATATTATATGGGTTGTTATGCTTTTCCATAAATTTGTATGTAAACCAATGGGAGGAATATATAACGAAATTAAAAAGGTTAAAAAGACGTATTTTATTGTTTGAAAAAAATAAATTTAGTGAGATTGTGGATCTGAATACTCAAAAGGGTGATGGTGACCATTTTGATGAAACACAAATATTTTCCATATTTTTTTCTTTTTTAATAAAAAAAAATGAGGGTTCAAAAATGAGGGATAATGATATGAATAGTGATAGCGAAGACAGTATATATGATGCTTATGAGCAGCAGATACAATTACACCATGGTGATAATATGGTCAATGGTATGTTGATGATGAGAAGGATATCGATGCAAAATTTAGAGGACGACGAAACCCAAGTTGAATATATAAATAGGGAGATCCATACGCAAGGTGATTTACATGTACGAAGAACGAATCAAGGTATTTTAAGATTTAATATGAGAAGGGGCAAAAAAGGGTCCAATGAAAATATGGGTGTCCACCATGAGAGTGGTAATGTGGATGATGCGAATGGTATGAATAATGTGGATGATACGAATAATATGAATAATGTGGATGGTACGAATAATATGAATAATGTGGATGGTACGAATAATATGAATAATATGGATGGTAGGAATAATATGAATAATATAAATAGTGTGGATAATATGAACAATTTAAATAATAATGATGGTGAAGAAGAAGAAGAATGTGTGAACGATGTTTTAAATTATGATAATAATAATTATGCTATTAATGAAGATGCTGAGGAATATATAAAAAATACGTCAGGAGAACGAGCTGTTATTATTTGTTCTGAAAAAAGAATATATGAAAAGAACGGAAATGGTGATATAATAACTAGAAATTATAAAAATGAAGAACGATATATATATTTAAAAAAATGGATTCCTTTTAAATCTATGATATTGTCAAAATTAGAAAAGAGGAAAAGGAATAGAAAAGAGGCATATAATACTCCTAGAGTATTAATATTAATACATTCCTTTTTATTTATACTTATTGTTTTCATATTTTTAATGGTATTCTTTAAAAAAGAGCCTATTTTTCGTTTTAATATGCCATCAGTTAATAAGCGTTTAAATAATTTTTTTAAATCAACAAGTTTTCACGAAATTATACCAAATAGTGTAGGGAAATGTAAAACAAAAAAATATATAGCAAAAGAACCATGTTTTAATGTGGGTCATATATACCATGAAGAGAAGACATTTTATCATGCAACTCTTTTATTTTTACAAGGTTTAAGATCTGTAAAAATTATGAATATGAATTTTTATTATGAAAAAGGCATATATTATTTATCTTTGGATGGATATTTTAAACATATTATAGGTCCTCTCTTTCTTAAATTATGTTTAGGTACAAACTTTTGTCCCATAAGTACATATGCATTTTTAGTAGGAAGTAAACCAACTTTCTCAGTAAATGTTGCTGTTCAATGTAATAATAAAAAACCACCATATTATATGACGGATATTATTGTAAAAGATTTAAAAATTACAAAAATAGAAATAGTTAAACATTCAGATGTTATCGATAATGTGGATATTAAATTAGATGATGTACAAGATAGAGTACAAGAAAAGGTTAATGCCATGTTAGAAGCAAAAAAAAAAATTATCGTCTGGAAAAATCAAAAGTATCATTTGGAGGGTTTTCTCAATTATTTAATATCTAAAAATGCATTATCTGGGTTTTCCTGTGAACCCATAAATTATTGAACTTGACAAATTAAAACACGTTAAATGTATATATGTATATATATTATATATGTATATATTTATTTATATTTTTCTTACATTTCCATTTGGGTGAACATATATTCTTACCCTTTTCATAAATTGTTTTTTTTTTTTTTGATATTTATCAATAATATATTCAGTAAAAATATATTATTTCACATAATTTAATATTTTTTGGAATAACATATAAATGTCAATGTATATGTGTATATATATATATATATATATATATGTATATATTTTTTTTTCTTTTTCATATATTTATATGCGCCCATTTGTTTTTATCTATATATATATATATTTATATATCCGAATTTCTTTATATTGTTATATTCCGATTTGATTATATATAAAATTTTTTTTAATTCATAATTTTTTTTTTTTTTTTTTTCTTTTTTTTCATTTCCATGTGATTTATTTCAATCCGATATTATTTATATGTTATATATATATTATTCTATAATATGATAAAAGGATATTTTTTTTTCTTATTTGTATTTTTAGATTATAAATATAAAGATATTCTTATATATAGATTATATCTGCTTTTTTTTTTTTTTTTTTTTTTTTTTCTATTATTAATTTTTAAATTATCTACTTACGAATTTTTTAATTGTTGATATGGCCTTCATGAGTATATTATTATGCATGAAAGAAAAAAAAAAAAAAAAATATATATATGTATATAATATATATATATATATATATAATATACCAATTTTTATTATCTTATATTTTACATGCACTATAAATGCATAGAATTTTTCTTTTTTTTTTTTTTGATTTCTTCTCCCCAAATTTTTGGTTATTTATATAATATACGTATTATAAATATGTACATATAATATGTATATATTATAATGTGTAATTTTCCACATATATATATAAAATATATATATATATATATATATATTTTAATAATACTTGAACTTTTAAAATTAAATATATAAAAAGGAATCCAATTAATTTATGGTTATATTATTTGTTATTCACTAGAAAATTCTCTAGAATTTAATAAAACATATAT

**5’ amplicon size: 1149bps**

**3’ amplicon size: 1116bps**

### PF3D7_0214800- mChSEP mCh SEP yDHODH recodonizedsequence

cattttattccgtaaaattaaaatgaattccaccaaaggtttaacatatacataatatatatttatatatatatatatatgtatatatatatatataaaataaaaataataatatataatatatttttatattacattattatattatattatattatattatattttattttatttttttaatatagaacttcatgtagctaaaataaaaatgagttcacatttttcgttaattaaatatattatatatatatatatatgcaaaattggtattatatataaatataatatataatttgtttccttttttttagtttttaaaatatagatttataaataatttttttcttttttttaagaattattttgatttaccatatatattatttaaaatcccaagttaaaattaaaaaaaaaaaggagtttttattataatattatatatatttatatatatattttaactactttaattttaaaagtgtgagagtaagaaaatgaaaaaaaaaaaaaaaaaaaaaaaaaggattattttttatttatatattataatatatataatatattaataataaataaaataaacatgtaaataaatattatatatatatatcataatatagtatcatgaaataatatatatttataaaaatattattataatatatagttaattattatattatatattacataatatttataatatattatatattggattatagttttacctattgggggatcttggttctataaaaaaatttcaatttttttttaccctttatttttttttgttcactatatatatatattgttatatataacataatatattaatagtttcataatattatatgtatatagaaagaaaaacatatatatatatatatatatattgttcattatttctaaatatatatattattatatatattatataaaaaatatattattttagatttactcctttatatataaattaatattttttatagatttcgattttttcctatttacataatttttaattttttataattttaatttttctacattatatatgtattatatatatatataaataatctatatatttttaaaataattcaatgtcattttattaattcaattaaatacaaattaatttcttattttctttaattatatatatatatattataatatttatagaacctatattatagatttaatttttaatctgttaaaagaaaaaaagtacaaaaaattatacatttatttacatatatatatatatatatatatatatatgtataatttttaatcttttatttttagaggtattttatgttattttttattttacatatgttgttgtataattttttttgtatatgcccttatattgattttcttttttttttttaaataatatagtgtttgctttaataaaaaaaaggaaatatcaggattttattgaatacatacatatatatatatatatatatatatatatatatatatatatatatatataactattagaattcccatagatgaatatataaacatataataatattatatctataatatttatatttttattatacatacatactgaatatattttaaagttacgaataatgaatgctttttttttttttttttttttgttgctttaaataaatggtaatcataaatgaaattagtctagagctaaaatgaaaattaaagaaaatataaaaagtatataataataaatgtgttataatatttaaaataaaacaaaaaaaaaaaaaaaaaaaaaaaaaaaagtaaattacaataatttattctataaagttcagaATG**TTGGAATTATTACGTAATGCTTATAATAAATGTAAAGAGTATATTTTGGGCTTATCGAAAAAATCACAAATATATAAAATAAATATATTAAATTATTGTTTATTATTTTCGTTTTTTTTATTTTATGCATCATATTTTTTTTTTATAAATGGAATAATAAATCCAGCAGTAAAAATTGTTTTCCATGGAA*gtaagtaccttatatgaaaaaatgaaagtgtatccataaaagaagatatacacaaaggtgtatatatatatatatatatatatatatatatatatatgtatgtatgtatgtatttatttatttatttattatttatatatatattttctttatattttctttaag*GTATTATTGAGGACAAAACCAAAGTTTACGAGAACTCGATAATTGAATCAATATGGCTATTGTTTACCAAGGGAAGTTATTTTAATATGGCCATGCTTCTATTCTTTTCTATTATAATACCGTTTCTCAAGTTACTTATGGTTAGTGATAACTTTTATAGTTTTATCGTATTATATAAAATGAATAAAAAACGAGAAGAGGAAAAGAGAAGGAGACGAAGAAGAATACGTGCACGAGAATATAACAATGCTAAATATAACAAGAATAGATATAAAATGAATAAATATAATAGAAATAAGAATATTAATGATATATACAAGGATGATATATATTATAGTGAAAATATTTTTAAGAATGATGAGATAAATTATATTAATACAATAAATGAGAATGAAGAATTTATTTTAAAGAAATTTAAAATTTTAAATTTCATATCGCGTTTTCAATTTGTTGATGTATTTATATCTTTATTTATAGTATCATCATTAAATTTATATTTATTAGAGGCAAGGATGTTAAATGGTGC**AGGTTTAAACGAGCAGAAGTTAATATCAGAAGAGGATTTGGGTGAACAAAAACTCATAAGCGAAGAAGATTTAATAACTTCGTATAGCATACATTATACGAAGTTATCCGGAGAAGGAAGAGGAAGTTTATTAACATGTGGAGATGTAGAAGAAAATCCAGGACCAATGACAGCCAGTTTAACTACCAAGTTCTTGAACAATACCTATGAAAACCCATTTATGAATGCATCCGGTGTTCATTGCATGACTACACAAGAATTAGATGAATTAGCAAACTCTAAAGCTGGCGCATTCATTACAAAGAGTGCTACAACCTTAGAAAGAGAAGGTAACCCTGAACCACGTTACATTTCTGTCCCTCTAGGCAGTATCAACTCCATGGGTTTACCAAACGAAGGTATCGACTACTATTTGTCCTATGTATTAAACCGTCAAAAGAATTATCCTGATGCACCTGCTATTTTCTTCTCAGTTGCTGGTATGAGCATTGATGAAAATTTAAATTTGTTGAGGAAAATCCAAGATAGCGAATTCAACGGTATTACCGAGTTAAACTTGTCTTGTCCTAATGTGCCTGGGAAACCACAAGTTGCTTATGACTTTGACTTGACAAAGGAAACCTTGGAAAAGGTTTTTGCCTTTTTCAAAAAACCTCTTGGTGTCAAGTTGCCTCCTTATTTTGATTTTGCCCATTTTGATATCATGGCAAAAATATTGAACGAGTTCCCATTAGCTTATGTCAACTCTATCAATAGTATAGGAAATGGTCTTTTCATTGATGTGGAGAAGGAGAGTGTAGTAGTGAAGCCAAAGAATGGTTTCGGGGGTATTGGAGGTGAATATGTTAAGCCAACCGCGCTCGCCAATGTTCGTGCATTTTACACTCGTTTGAGACCTGAAATCAAAGTTATCGGTACAGGTGGAATTAAGTCCGGTAAGGATGCATTTGAACATCTTCTATGTGGTGCCTCTATGCTACAGATTGGTACAGAATTACAAAAAGAGGGCGTCAAGATTTTTGAACGTATCGAAAAAGAATTAAAAGACATAATGGAAGCTAAGGGTTATACATCCATAGATCAGTTCCGTGGGAAGTTGAACAGCATTGGTGAAGGTAGAGGTTCTTTGTTGACTTGTGGTGATGTTGAAGAAAATCCAGGTCCAGCTAGCATGACGCGTGCTAGAGGTGCTGCTGCTGGTGCTGGAGGTGCAGGTAGACCTAGGATGCTTGAGCTTCTTAGAAACGCATACAACAAGTGCAAGGAATACATACTTGGACTTAGTAAGAAGAGTCAGATTTACAAGATTAACATTCTTAACTACTGCCTTCTTTTCAGTTTCTTCCTTTTCTACGCTAGTTACTTCTTCTTCATTAACGGTATTATTAACCCTGCTGTTAAGATAGTATTTCACGGTTCAATAATAGAAGATAAGACAAAGGTATATGAAAATAGTATTATAGAGAGTATTTGGTTGCTTTTCACAAAAGGTTCATACTTCAACATGGCAATGTTATTGTTTTTCAGCATAATTATTCCATTCTTAAAACTTTTAATGGTATCAGACAATTTCTACTCATTCATAGTTCTTTACAAGATGAACAAGAAGAGGGAGGAAGAGAAACGTCGTCGTAGGCGTCGTATTAGAGCTAGGGAGTACAATAACGCAAAGTACAATAAAAACCGTTACAAGATGAACAAGTACAACCGTAACAAAAACATAAACGACATTTATAAAGACGACATTTACTACTCAGAGAACATATTCAAAAACGACGAAATTAACTACATAAACACTATTAACGAAAACGAGGAGTTCATACTTAAAAAGTTCAAGATACTTAACTTTATTAGTAGATTCCAGTTCGTAGACGTTTTCATTAGCCTTTTCATTGTTAGTAGTCTTAACCTTTACCTTCTTGAAGCTCGTATGCTTAACGGAGCTTACTACTTCCTTAACTACTGCATGCTTAGTACTATTAGTAGCTTCCTTCTTTTCTCATTCACAAGCCTTAAGATTCACATTTTCAAGAACGGTAACATAAAAATTAGCGCTTGCTTAAACGAGAGTAACTTGGAAGTAACAACTTCAGGACCATTAAGTACTAAGGACTTAGTTGAGGAGGAGGGTCACGCTCAGATTAACAACTTGATAATAAACGACAAAAACATGACATCAGGAGTAGTAAACGACTTCAGTGGAAACGGTAATAACGTTGAGCTTACTGATGATTTAAAATCAGGAGAGCCACAGAATCGTGACGACATTCAGACAGAGGAGACTAAGAAGGAGAAGATGAACACACGTACACACAACGACGAGGACAACACTAAGAAGAAGAACATTAAAGACAAGAAAAAGGCAAATGGAGACGACAAGGTTATTCAGAAGTGCATAGACAACGAGCGTAAGAAGAAGCAGAACGGAATGATACAGTCAGTTAATGACGGAGACAAGAACTCAAACTTCAACAACAATAACAACAACAACATAAACGGAGACTCAAACAACAACAACATAAACGGAGACTCAAACAACAACAACATAAACGGAGACTCAAACAACAACAACATAAACGGAGACTCAAACAACAACAACATAAACGGAGACTCAAACAACAACAACATAAACGGAGACTCAAACAACAACAACTACCACAACAACTACCACAACAACTACAGAAACAACTACCACAACAACTACAGAAACAACAACTGCCGTAACAACATACTTGAGCAGAACAAGTGCGACAAGAACGTACTTTGCTACAACAACATTTACAACACTATGAAGGACAACGACACATACATTTACCTTAAGAAAAACAAGTTCAACAGTCTTCTTAAGTCAAACTGCATAAAGACAAACTTCAACATGATTAAAATTGGATACGTTATTTTCCTTTTCGTTCTTCTTTGCCTTTGCATTTACCTTATTACTGGTGTTGAGTGTTCACTTTTCGGAATTTACATTTACCTTTCATACTTCAACTTCAACATAGAGGGTATTCTTATAGACTACATGGACATGCTTAACATACTTAAGCTTAAGATTAAAAAGGGTTACATATACCCATTCTTCGTAATGCTTCCTTTCATATTTCCAGTTATTATTAGCATGTGCTTTTTCCTTTCAGTTTTCTTCCTTAACATGTACTACGAGTCATTCAGTAAACTTTACAAGAAAATATCAGAGTTGAAAAACGAGTTTATTAATAGTAGTGAGAACGATAACGTTAATGAGCGTATTCTTGTTAGTGAGACAAGCAACCACCTTTGCCTTAACGAGTCAAATGACAAGGTTAGCAACACTAGTGACGACTTCCTTAGTCGTAACAACTCAAATATTAGCTCAAGTAAGTCAGAGATGATTAACAGTAACTTCGTTTTCAACAAGTTACTTAACTTCTACTTCAGTTTCGCTGTATTTTTCAGCTACCTTGGAAGCGCTTTCCTTCACATTAGTCTTGGAGAGATAATTTGCATAGCTCTTCTTACTTTCTACCAGATAGTTAAACACACAAACAACCTTAACATAACTATACTTTTAAAATCAGAAAAGATAAAGTTCTGCAAGTTTTTACTTTTCATTCTTTACGGACTTCTTTGTTTCAGTATTAACCTTTACGTTAATCAGTGGGAAGAGTACATTACAAAGCTTAAGCGTCTTAAGCGTAGAATACTTCTTTTCGAGAAGAACAAGTTCTCAGAAATAGTTGACTTAAACACACAGAAAGGAGACGGAGATCACTTCGACGAGACTCAGATTTTCAGTATTTTCTTCAGCTTCCTTATTAAGAAGAACGAAGGAAGTAAGATGCGTGACAACGACATGAACTCAGACTCAGAGGATTCAATTTACGACGCATACGAACAACAAATTCAGCTTCATCACGGAGACAACATGGTTAACGGAATGCTTATGATGCGTCGTATTAGTATGCAGAACCTTGAAGATGATGAGACACAGGTAGAGTACATTAACCGTGAAATACACACACAGGGAGACCTTCACGTTAGGCGTACAAACCAGGGAATACTTCGTTTCAACATGCGTCGTGGAAAGAAGGGAAGTAACGAGAACATGGGAGTTCATCACGAATCAGGAAACGTTGACGACGCAAACGGAATGAACAACGTTGACGACACAAACAACATGAACAACGTTGACGGAACAAACAACATGAACAACGTTGACGGAACAAACAACATGAACAACATGGACGGACGTAACAACATGAACAACATTAACTCAGTTGACAACATGAATAACCTTAACAACAACGACGGAGAGGAGGAGGAGGAGTGCGTTAATGACGTACTTAACTACGACAACAACAACTACGCAATAAACGAGGACGCAGAAGAGTACATTAAGAACACAAGTGGTGAGAGGGCAGTAATAATATGCAGCGAGAAGCGTATTTACGAGAAAAATGGTAACGGAGACATTATTACACGTAACTACAAGAACGAGGAGAGGTACATTTACCTTAAGAAGTGGATACCATTCAAGAGCATGATTCTTAGTAAGCTTGAGAAACGTAAGCGTAACCGTAAGGAAGCTTACAACACACCACGTGTTCTTATTCTTATTCACAGTTTCCTTTTCATTTTAATAGTATTTATTTTCCTTATGGTTTTTTTCAAGAAGGAACCAATATTCAGATTCAACATGCCTAGTGTAAACAAAAGACTTAACAACTTCTTCAAGAGTACTTCATTCCATGAGATAATTCCTAACTCAGTTGGAAAGTGCAAGACTAAGAAGTACATTGCTAAGGAGCCTTGCTTCAACGTTGGACACATTTATCACGAGGAAAAAACTTTCTACCACGCTACATTACTTTTCCTTCAGGGACTTCGTAGCGTTAAGATAATGAACATGAACTTCTACTACGAGAAGGGAATTTACTACCTTAGCCTTGACGGTTACTTCAAGCACATAATTGGACCATTATTCTTAAAGCTTTGCCTTGGAACTAATTTCTGCCCAATTTCAACTTACGCTTTCCTTGTTGGTTCAAAGCCTACATTTAGTGTTAACGTAGCAGTACAGTGCAACAACAAGAAGCCTCCTTACTACATGACAGACATAATAGTTAAGGACCTTAAGATAACTAAGATTGAGATTGTAAAGCACAGTGACGTAATAGACAACGTTGACATAAAGCTTGACGACGTTCAGGACCGTGTTCAGGAGAAAGTAAACGCAATGCTTGAGGCTAAGAAGAAGATAATAGTTTGGAAGAACCAGAAATACCACCTTGAAGGATTCTTAAACTACCTTATTAGCAAGAACGCTCTTAGCGGATTCAGTTGCGAGCCAATTAACTACACAAGTTATCCATATGATAATCCAGATTATGCACCAGTTGCAACATTAGGTAGTTTAAACATGGTGAGCAAGGGCGAGGAGGATAACATGGCCATCATCAAGGAGTTCATGCGCTTCAAGGTGCACATGGAGGGCTCCGTGAACGGCCACGAGTTCGAGATCGAGGGCGAGGGCGAGGGCCGCCCCTACGAGGGCACCCAGACCGCCAAGCTGAAGGTGACCAAGGGTGGCCCCCTGCCCTTCGCCTGGGACATCCTGTCCCCTCAGTTCATGTACGGCTCCAAGGCCTACGTGAAGCACCCCGCCGACATCCCCGACTACTTGAAGCTGTCCTTCCCCGAGGGCTTCAAGTGGGAGCGCGTGATGAACTTCGAGGACGGCGGCGTGGTGACCGTGACCCAGGACTCCTCCCTGCAGGACGGCGAGTTCATCTACAAGGTGAAGCTGCGCGGCACCAACTTCCCCTCCGACGGCCCCGTAATGCAGAAGAAGACCATGGGCTGGGAGGCCTCCTCCGAGCGGATGTACCCCGAGGACGGCGCCCTGAAGGGCGAGATCAAGCAGAGGCTGAAGCTGAAGGACGGCGGCCACTACGACGCTGAGGTCAAGACCACCTACAAGGCCAAGAAGCCCGTGCAGCTGCCCGGCGCCTACAACGTCAACATCAAGTTGGACATCACCTCCCACAACGAGGACTACACCATCGTGGAACAGTACGAACGCGCCGAGGGCCGCCACTCCACCGGCGGCATGGACGAGCTGTACAAGACGCGTGTTATCGGAGCAGGTGCAGGTATGGGAAGTAAAGGAGAAGAACTTTTCACTGGAGTTGTCCCAATTCTTGTTGAATTAGATGGTGATGTTAATGGGCACAAATTTTCTGTCAGTGGAGAGGGTGAAGGTGATGCAACATACGGAAAACTTACCCTTAAATTTATTTGCACTACTGGAAAACTACCTGTTCCTTGGCCAACACTTGTCACTACTTTAACTTATGGTGTTCAATGCTTTTCAAGATACCCAGATCATATGAAACGGCATGACTTTTTCAAGAGTGCCATGCCCGAAGGTTATGTACAGGAAAGAACTATATTTTTCAAAGATGACGGGAACTACAAGACACGTGCTGAAGTCAAGTTTGAAGGTGATACCCTTGTTAATAGAATCGAGTTAAAAGGTATTGATTTTAAAGAAGATGGAAACATTCTTGGACACAAATTGGAATACAACTATAACGATCACCAGGTGTACATCATGGCAGACAAACAAAAGAATGGAATCAAAGCTAACTTCAAAATTAGACACAACATTGAAGATGGAGGCGTTCAACTAGCAGACCATTATCAACAAAATACTCCAATTGGCGATGGGCCCGTCCTTTTACCAGACAACCATTACCTGTTTACAACTTCTACTCTTTCGAAAGATCCCAACGAAAAGAGAGACCACATGGTCCTTCTTGAGTTTGTAACAGCTGCTGGGATTACACATGGCATGGATGAACTATACAAAGATAACtagaggcctataacttcgtatagcatacattatacgaagttattatgactcgagggatatggcagcttaatgttcgtttttcttatttatatatttataccaattgattgtatttataactgtaaaaatgtgtatgttgtgtgcatatttttttttgtgcatgcacatgcatgtaaatagctaaaattatgaacattttattttttgttcagaaaaaaaaaactttacacacataaaatggctagtatgaatagccatattttatataaattaaatcctatgaatttatgaccatattaaaaatttagatatttatggaacataatatgtttgaaacaataagacaaaattattattattattattatttttactgttataattatgtgtctccttcaatgattcataaatagttggacttgatttttaaaatgtttataatatgattagcatagttaaataaaaaaagttgaaaaattaaaaaaaaacatataaacacaaatgatggtttttccttcaatttcgatatcaatttatagaaacaaaatatatacttgtataattttatttttttatataaatcattacatatataattatacaatattttttctaagagataattatatattaatatatataaaaaaaggtgttttttttttttttttttatttttatttttattttatggtaatattttattttccttattttataaattatattagtttatatgtgattaattttatatattatcaatttatatatttttaaatgcttacttaattatctttttttttttttttttttttttttcccctctttttatattaatttatttttgaaaaaattgatatatatatatatatataatatatatatatacatgtagtagtattaaacaatgtataatatatataaataatatatttatatatttcatttcaattttaattttttttggttttttttttttttctttttgtcatatttaaaaaaaattatattcatataagttatgcattttttataaacattattcaatatatgtataatataatatatatatatatattaatgtattattccaatgtgcatgataaaagaaaaaaataatatttataaaaaaaaagaaaaataaaacaaaaaaagaaaaaaaaaaaaaaaaaaaaaaaaatacaaaaataaataatataatttataattatatattcttgtcacaataaaaatatatatatatatatatatatttataatatgtatattttaaactagaaaaggaataactaatattttatttattatcattcaagatttatattttataataataaatacctaatagaaatatatcaggatccatgcatggttcgctaaactgcatcgtcgctgtgtcccagaacatgggcatcggcaagaacggggactacccctggccaccgctcaggaacgaatttagatatttccagagaatgaccacaacctcttcagtagaaggtaaacagaatctggtgattatgggtaagaagacctggttctccattcctgagaagaatcgacctttaaagggtagaattaatttagttctcagcagagaactcaaggaacctccacaaggagctcattttctttccagaagtctagatgatgccttaaaacttactgaacaaccagaattagcaaataaagtagacatggtctggatagttggtggcagttctgtttataaggaagccatgaatcacccaggccatcttaaactatttgtgacaaggatcatgcaagactttgaaagtgacacgttttttccagaaattgatttggagaaatataaacttctgccagaatacccaggtgttctctctgatgtccaggaggagaaaggcattaagtacaaatttgaagtatatgagaagaatgattaagcttatttaataatagattaaaaatattataaaaataaaaacataaacacagaaattacaaaaaaaatacatatgaattttttttttgtaatcttccttataaatatagaataatgaatcatataaaacatatcattattcatttatttacatttaaaattattgtttcagtatctttaatttattatgtatatataaaaataacttacaattttattaataaacaatatatgtttattaattcatgttttgtaatttatgggatagcgattttttttactgtctgtatttttcttttttaattatgttttaattgtattttatttttattattgttctttttatagtattattttaaaacaaaatgtattttctaagaacttataataataataaatataaattttaataaaaattatatttatcttttacaatatgaacataaagtacaacattaatatatagcttttaatatttttattcctaatcatgtaaatcttaaatttttctttttaaacatatgttaaatatttatttctcattatatataagaacatatttattaaatctagaattctatagtgagtcgtattacaattcactggccgtcgttttacaacgtcgtgactgggaaaaccctggcgttacccaacttaatcgccttgcagcacatccccctttcgccagctggcgtaatagcgaagaggcccgcaccgatcgcccttcccaacagttgcgcagcctgaatggcgaatggcgcctgatgcggtattttctccttacgcatctgtgcggtatttcacaccgcatatggtgcactctcagtacaatctgctctgatgccgcatagttaagccagccccgacacccgccaacacccgctgacgcgccctgacgggcttgtctgctcccggcatccgcttacagacaagctgtgaccgtctccgggagctgcatgtgtcagaggttttcaccgtcatcaccgaaacgcgcgagacgaaagggcctcgtgatacgcctatttttataggttaatgtcatgataataatggtttcttagacgtcaggtggcacttttcggggaaatgtgcgcggaacccctatttgtttatttttctaaatacattcaaatatgtatccgctcatgagacaataaccctgataaatgcttcaataatattgaaaaaggaagagtatgagtattcaacatttccgtgtcgcccttattcccttttttgcggcattttgccttcctgtttttgctcacccagaaacgctggtgaaagtaaaagatgctgaagatcagttgggtgcacgagtgggttacatcgaactggatctcaacagcggtaagatccttgagagttttcgccccgaagaacgttttccaatgatgagcacttttaaagttctgctatgtggcgcggtattatcccgtattgacgccgggcaagagcaactcggtcgccgcatacactattctcagaatgacttggttgagtactcaccagtcacagaaaagcatcttacggatggcatgacagtaagagaattatgcagtgctgccataaccatgagtgataacactgcggccaacttacttctgacaacgatcggaggaccgaaggagctaaccgcttttttgcacaacatgggggatcatgtaactcgccttgatcgttgggaaccggagctgaatgaagccataccaaacgacgagcgtgacaccacgatgcctgtagcaatgccaacaacgttgcgcaaactattaactggcgaactacttactctagcttcccggcaacaattaatagactggatggaggcggataaagttgcaggaccacttctgcgctcggcccttccggctggctggtttattgctgataaatctggagccggtgagcgtgggtctcgcggtatcattgcagcactggggccagatggtaagccctcccgtatcgtagttatctacacgacggggagtcaggcaactatggatgaacgaaatagacagatcgctgagataggtgcctcactgattaagcattggtaactgtcagaccaagtttactcatatatactttagattgatttaaaacttcatttttaatttaaaaggatctaggtgaagatcctttttgataatctcatgaccaaaatcccttaacgtgagttttcgttccactgagcgtcagaccccgtagaaaagatcaaaggatcttcttgagatcctttttttctgcgcgtaatctgctgcttgcaaacaaaaaaaccaccgctaccagcggtggtttgtttgccggatcaagagctaccaactctttttccgaaggtaactggcttcagcagagcgcagataccaaatactgtccttctagtgtagccgtagttaggccaccacttcaagaactctgtagcaccgcctacatacctcgctctgctaatcctgttaccagtggctgctgccagtggcgataagtcgtgtcttaccgggttggactcaagacgatagttaccggataaggcgcagcggtcgggctgaacggggggttcgtgcacacagcccagcttggagcgaacgacctacaccgaactgagatacctacagcgtgagctatgagaaagcgccacgcttcccgaagggagaaaggcggacaggtatccggtaagcggcagggtcggaacaggagagcgcacgagggagcttccagggggaaacgcctggtatctttatagtcctgtcgggtttcgccacctctgacttgagcgtcgatttttgtgatgctcgtcaggggggcggagcctatcgaaaaacgccagcaacgcggcctttttacggttcctggccttttgctggccttttgctcacatgttctttcctgcgttatcccctgattctgtggataaccgtattaccgcctttgagtgagctgataccgctcgccgcagccgaacgaccgagcgcagcgagtcagtgagcgaggaagcggaagagcgcccaatacgcaaaccgcctctccccgcgcgttggccgattcattaatgcagctggcacgacaggtttcccgactggaaagcgggcagtgagcgcaacgcaattaatgtgagttagctcactcattaggcaccccaggctttacactttatgcttccggctcgtatgttgtgtggaattgtgagcggataacaatttcacacaggaaacagctatgaccatgattacgccaagctatttaggtgacactatagaatactcgcggccgcTAGTTGGAATTATTACGTAATGCTTATAATAAATGTAAAGAGTATATTTTGGGCTTATCGAAAAAATCACAAATATATAAAATAAATATATTAAATTATTGTTTATTATTTTCGTTTTTTTTATTTTATGCATCATATTTTTTTTTTATAAATGGAATAATAAATCCAGCAGTAAAAATTGTTTTCCATGGAAGTAAGTACCTTATATGAAAAAATGAAAGTGTATCCATAAAAGAAGATATACACAAAGGTGTATATATATATATATATATATATATATATATATATATGTATGTATGTATGTATTTATTTATTTATTTATTATTTATATATATATTTTCTTTATATTTTCTTTAAGGTATTATTGAGGACAAAACCAAAGTTTACGAGAACTCGATAATTGAATCAATATGGCTATTGTTTACCAAGGGAAGTTATTTTAATATGGCCATGCTTCTATTCTTTTCTATTATAATACCGTTTCTCAAGTTACTTATGGTTAGTGATAACTTTTATAGTTTTATCGTATTATATAAAATGAATAAAAAACGAGAAGAGGAAAAGAGAAGGAGACGAAGAAGAATACGTGCACGAGAATATAACAATGCTAAATATAACAAGAATAGATATAAAATGAATAAATATAATAGAAATAAGAATATTAATGATATATACAAGGATGATATATATTATAGTGAAAATATTTTTAAGAATGATGAGATAAATTATATTAATACAATAAATGAGAATGAAGAATTTATTTTAAAGAAATTTAAAATTTTAAATTTCATATCGCGTTTTCAATTTGTTGATGTATTTATATCTTTATTTATAGTATCATCATTAAATTTATATTTATTAGAGGCAAGGATGTTAAATGGTGCATATTATTTTTTGAATTATTGTATGTTATCAACAATATCATCTTTTTTGTTATTTAGTTTTACTTCTTTAAAAATACATATATTTAAAAATGGAAATATTAAGATATCTGCATGTCTCAATGAATCAAATCTAGAGGTTACTACAAGCGGTCCCCTTTCAACAAAAGATCTTGTCGAAGAAGAAGGACATGCACAAATAAATAATCTAATTATTAATGATAAGAATATGACTAGTGGTGTTGTTAATGATTTTTCAGGTAATGGAAACAATGTCGAATTAACAGACGACCTTAAGAGTGGCGAACCCCAAAACAGAGATGATATACAAACTGAAGAAACAAAAAAAGAAAAAATGAATACTAGAACTCATAATGATGAAGATAATACAAAAAAAAAAAATATAAAGGATAAAAAGAAAGCTAACGGTGATGATAAAGTAATACAAAAATGTATTGATAATGAAAGGAAAAAAAAACAAAATGGTATGATTCAAAGTGTAAACGATGGTGATAAAAATAGTAATTTTAATAATAACAATAATAATAATATTAATGGTGATAGTAATAATAATAATATTAATGGTGATAGTAATAATAATAATATTAATGGTGATAGTAATAATAATAATATTAATGGTGATAGTAATAATAATAATATTAATGGTGATAGTAATAATAATAATATTAATGGTGATAGTAATAATAATAATTATCATAATAATTATCATAATAATTATCGTAATAATTATCATAATAATTATCGTAATAATAATTGTAGGAATAATATTTTAGAACAAAATAAATGTGATAAAAATGTTTTGTGTTATAATAATATATATAATACAATGAAAGATAATGATACTTATATATATTTAAAAAAGAATAAATTTAATTCGTTATTAAAAAGTAATTGTATCAAAACTAATTTTAATATGATAAAGATAGGTTATGTAATATTTTTATTTGTGTTATTATGTTTATGTATATATTTAATAACAGGAGTAGAATGCAGTTTATTTGGTATATATATATATTTGAGTTATTTTAATTTTAATATTGAAGGAATATTAATTGATTATATGGATATGTTAAATATTTTAAAATTAAAAATAAAGAAAGGATATATCTATCCTTTTTTTGTTATGTTACCATTTATTTTCCCTGTAATAATATCTATGTGTTTCTTTTTAAGTGTATTTTTTTTAAATATGTATTATGAAAGCTTTTCAAAGTTATATAAAAAGATCAGTGAACTAAAGAATGAATTCATAAACTCCTCAGAAAATGACAATGTGAACGAAAGAATATTGGTATCCGAAACTTCTAATCATTTATGTTTAAATGAAAGTAACGATAAAGTATCTAATACATCAGATGATTTTTTATCAAGAAATAATAGCAACATATCTAGTTCAAAAAGTGAAATGATAAATTCAAATTTTGTATTTAATAAACTTTTGAATTTTTATTTTTCATTTGCAGTTTTCTTTTCTTATTTAGGTTCTGCATTTTTACATATATCCTTAGGTGAAATTATATGTATCGCATTATTAACATTTTATCAAATTGTAAAGCATACCAATAATTTGAATATCACAATTTTACTTAAGAGTGAGAAAATTAAATTTTGTAAATTCCTTTTGTTTATATTATATGGGTTGTTATGCTTTTCCATAAATTTGTATGTAAACCAATGGGAGGAATATATAACGAAATTAAAAAGGTTAAAAAGACGTATTTTATTGTTTGAAAAAAATAAATTTAGTGAGATTGTGGATCTGAATACTCAAAAGGGTGATGGTGACCATTTTGATGAAACACAAATATTTTCCATATTTTTTTCTTTTTTAATAAAAAAAAATGAGGGTTCAAAAATGAGGGATAATGATATGAATAGTGATAGCGAAGACAGTATATATGATGCTTATGAGCAGCAGATACAATTACACCATGGTGATAATATGGTCAATGGTATGTTGATGATGAGAAGGATATCGATGCAAAATTTAGAGGACGACGAAACCCAAGTTGAATATATAAATAGGGAGATCCATACGCAAGGTGATTTACATGTACGAAGAACGAATCAAGGTATTTTAAGATTTAATATGAGAAGGGGCAAAAAAGGGTCCAATGAAAATATGGGTGTCCACCATGAGAGTGGTAATGTGGATGATGCGAATGGTATGAATAATGTGGATGATACGAATAATATGAATAATGTGGATGGTACGAATAATATGAATAATGTGGATGGTACGAATAATATGAATAATATGGATGGTAGGAATAATATGAATAATATAAATAGTGTGGATAATATGAACAATTTAAATAATAATGATGGTGAAGAAGAAGAAGAATGTGTGAACGATGTTTTAAATTATGATAATAATAATTATGCTATTAATGAAGATGCTGAGGAATATATAAAAAATACGTCAGGAGAACGAGCTGTTATTATTTGTTCTGAAAAAAGAATATATGAAAAGAACGGAAATGGTGATATAATAACTAGAAATTATAAAAATGAAGAACGATATATATATTTAAAAAAATGGATTCCTTTTAAATCTATGATATTGTCAAAATTAGAAAAGAGGAAAAGGAATAGAAAAGAGGCATATAATACTCCTAGAGTATTAATATTAATACATTCCTTTTTATTTATACTTATTGTTTTCATATTTTTAATGGTATTCTTTAAAAAAGAGCCTATTTTTCGTTTTAATATGCCATCAGTTAATAAGCGTTTAAATAATTTTTTTAAATCAACAAGTTTTCACGAAATTATACCAAATAGTGTAGGGAAATGTAAAACAAAAAAATATATAGCAAAAGAACCATGTTTTAATGTGGGTCATATATACCATGAAGAGAAGACATTTTATCATGCAACTCTTTTATTTTTACAAGGTTTAAGATCTGTAAAAATTATGAATATGAATTTTTATTATGAAAAAGGCATATATTATTTATCTTTGGATGGATATTTTAAACATATTATAGGTCCTCTCTTTCTTAAATTATGTTTAGGTACAAACTTTTGTCCCATAAGTACATATGCATTTTTAGTAGGAAGTAAACCAACTTTCTCAGTAAATGTTGCTGTTCAATGTAATAATAAAAAACCACCATATTATATGACGGATATTATTGTAAAAGATTTAAAAATTACAAAAATAGAAATAGTTAAACATTCAGATGTTATCGATAATGTGGATATTAAATTAGATGATGTACAAGATAGAGTACAAGAAAAGGTTAATGCCATGTTAGAAGCAAAAAAAAAAATTATCGTCTGGAAAAATCAAAAGTATCATTTGGAGGGTTTTCTCAATTATTTAATATCTAAAAATGCATTATCTGGGTTTTCCTGTGAACCCATAAATTATTGAacttgacaaattaaaacacgttaaatgtatatatgtatatatattatatatgtatatatttatttatatttttcttacatttccatttgggtgaacatatattcttacccttttcataaattgttttttttttttttgatatttatcaataatatattcagtaaaaatatattatttcacataatttaatattttttggaataacatataaatgtcaatgtatatgtgtatatatatatatatatatatatatgtatatatttttttttctttttcatatatttatatgcgcccatttgtttttatctatatatatatatatttatatatccgaatttctttatattgttatattccgatttgattatatataaaattttttttaattcataatttttttttttttttttttcttttttttcatttccatgtgatttatttcaatccgatattatttatatgttatatatatattattctataatatgataaaaggatatttttttttcttatttgtatttttagattataaatataaagatattcttatatatagattatatctgcttttttttttttttttttttttttttctattattaatttttaaattatctacttacgaattttttaattgttgatatggccttcatgagtatattattatgcatgaaagaaaaaaaaaaaaaaaaatatatatatgtatataatatatatatatatatatataatataccaatttttattatcttatattttacatgcactataaatgcatagaatttttcttttttttttttttgatttcttctccccaaatttttggttatttatataatatacgtattataaatatgtacatataatatgtatatattataatgtgtaattttccacatatatatataaaatatatatatatatatatatatattttaataatacttgaacttttaaaattaaatatataaaaaggaatccaattaatttatggttatattatttgttattcactagaaaattctctagaatttaataaaacatatat

**5’ amplicon size: 1149bps**

**3’ amplicon size: 1116bps**

### > GFP-2xFKBP-PF3D7_0214800-3UTR-mCh- PF3D7_0214800 mCh GFP yDHODH recodonizedsequence1 recodonizedsequence2

Similar integration and PCR products for the LW/VW mutant.

cattttattccgtaaaattaaaatgaattccaccaaaggtttaacatatacataatatatatttatatatatatatatatgtatatatatatatataaaataaaaataataatatataatatatttttatattacattattatattatattatattatattatattttattttatttttttaatatagaacttcatgtagctaaaataaaaatgagttcacatttttcgttaattaaatatattatatatatatatatatgcaaaattggtattatatataaatataatatataatttgtttccttttttttagtttttaaaatatagatttataaataatttttttcttttttttaagaattattttgatttaccatatatattatttaaaatcccaagttaaaattaaaaaaaaaaaggagtttttattataatattatatatatttatatatatattttaactactttaattttaaaagtgtgagagtaagaaaatgaaaaaaaaaaaaaaaaaaaaaaaaaggattattttttatttatatattataatatatataatatattaataataaataaaataaacatgtaaataaatattatatatatatatcataatatagtatcatgaaataatatatatttataaaaatattattataatatatagttaattattatattatatattacataatatttataatatattatatattggattatagttttacctattgggggatcttggttctataaaaaaatttcaatttttttttaccctttatttttttttgttcactatatatatatattgttatatataacataatatattaatagtttcataatattatatgtatatagaaagaaaaacatatatatatatatatatatattgttcattatttctaaatatatatattattatatatattatataaaaaatatattattttagatttactcctttatatataaattaatattttttatagatttcgattttttcctatttacataatttttaattttttataattttaatttttctacattatatatgtattatatatatatataaataatctatatatttttaaaataattcaatgtcattttattaattcaattaaatacaaattaatttcttattttctttaattatatatatatatattataatatttatagaacctatattatagatttaatttttaatctgttaaaagaaaaaaagtacaaaaaattatacatttatttacatatatatatatatatatatatatatatgtataatttttaatcttttatttttagaggtattttatgttattttttattttacatatgttgttgtataattttttttgtatatgcccttatattgattttcttttttttttttaaataatatagtgtttgctttaataaaaaaaaggaaatatcaggattttattgaatacatacatatatatatatatatatatatatatatatatatatatatatatatataactattagaattcccatagatgaatatataaacatataataatattatatctataatatttatatttttattatacatacatactgaatatattttaaagttacgaataatgaatgctttttttttttttttttttttgttgctttaaataaatggtaatcataaatgaaattagtctagagctaaaatgaaaattaaagaaaatataaaaagtatataataataaatgtgttataatatttaaaataaaacaaaaaaaaaaaaaaaaaaaaaaaaaaaagtaaattacaataatttattctataaagttcagaATG**TTGGAATTATTACGTAATGCTTATAATAAATGTAAAGAGTATATTTTGGGCTTATCGAAAAAATCACAAATATATAAAATAAATATATTAAATTATTGTTTATTATTTTCGTTTTTTTTATTTTATGCATCATATTTTTTTTTTATAAATGGAATAATAAATCCAGCAGTAAAAATTGTTTTCCATGGAA*gtaagtaccttatatgaaaaaatgaaagtgtatccataaaagaagatatacacaaaggtgtatatatatatatatatatatatatatatatatatatgtatgtatgtatgtatttatttatttatttattatttatatatatattttctttatattttctttaag*GTATTATTGAGGACAAAACCAAAGTTTACGAGAACTCGATAATTGAATCAATATGGCTATTGTTTACCAAGGGAAGTTATTTTAATATGGCCATGCTTCTATTCTTTTCTATTATAATACCGTTTCTCAAGTTACTTATGGTTAGTGATAACTTTTATAGTTTTATCGTATTATATAAAATGAATAAAAAACGAGAAGAGGAAAAGAGAAGGAGACGAAGAAGAATACGTGCACGAGAATATAACAATGCTAAATATAACAAGAATAGATATAAAATGAATAAATATAATAGAAATAAGAATATTAATGATATATACAAGGATGATATATATTATAGTGAAAATATTTTTAAGAATGATGAGATAAATTATATTAATACAATAAATGAGAATGAAGAATTTATTTTAAAGAAATTTAAAATTTTAAATTTCATATCGCGTTTTCAATTTGTTGATGTATTTATATCTTTATTTATAGTATCATCATTAAATTTATATTTATTAGAGGCAAGGATGTTAAATGGTG**AGgtttaaacgagcagaagttaatatcagaagaggatttgggtgaacaaaaactcataagcgaagaagatttaataacttcgtatagcatacattatacgaagttatccggagaaggaagaggaagtttattaacatgtggagatgtagaagaaaatccaggaccaatgacagccagtttaactaccaagttcttgaacaatacctatgaaaacccatttatgaatgcatccggtgttcattgcatgactacacaagaattagatgaattagcaaactctaaagctggcgcattcattacaaagagtgctacaaccttagaaagagaaggtaaccctgaaccacgttacatttctgtccctctaggcagtatcaactccatgggtttaccaaacgaaggtatcgactactatttgtcctatgtattaaaccgtcaaaagaattatcctgatgcacctgctattttcttctcagttgctggtatgagcattgatgaaaatttaaatttgttgaggaaaatccaagatagcgaattcaacggtattaccgagttaaacttgtcttgtcctaatgtgcctgggaaaccacaagttgcttatgactttgacttgacaaaggaaaccttggaaaaggtttttgcctttttcaaaaaacctcttggtgtcaagttgcctccttattttgattttgcccattttgatatcatggcaaaaatattgaacgagttcccattagcttatgtcaactctatcaatagtataggaaatggtcttttcattgatgtggagaaggagagtgtagtagtgaagccaaagaatggtttcgggggtattggaggtgaatatgttaagccaaccgcgctcgccaatgttcgtgcattttacactcgtttgagacctgaaatcaaagttatcggtacaggtggaattaagtccggtaaggatgcatttgaacatcttctatgtggtgcctctatgctacagattggtacagaattacaaaaagagggcgtcaagatttttgaacgtatcgaaaaagaattaaaagacataatggaagctaagggttatacatccatagatcagttccgtgggaagttgaacagcattggtgaaggtagaggttctttgttgacttgtggtgatgttgaagaaaatccaggtccagctagcatgagtaaaggagaagaacttttcactggagttgtcccaattcttgttgaattagatggtgatgttaatgggcacaaattttctgtcagtggagagggtgaaggtgatgcaacatacggaaaacttacccttaaatttatttgcactactggaaaactaccagttccatggccaacacttgtcactactttcgcgtatggtcttcaatgctttgcgagatacccagatcatatgaaacagcatgactttttcaagagtgccatgcccgaaggttatgtacaggaaagaactatatttttcaaagatgacgggaactacaagacacgtgctgaagtcaagtttgaaggtgatacccttgttaatagaatcgagttaaaaggtattgattttaaagaagatggaaacattcttggacacaaattggaatacaactataactcacacaatgtatacatcatggcagacaaacaaaagaatggaatcaaagttaacttcaaaattagacacaacattgaagatggaagcgttcaactagcagaccattatcaacaaaatactccaattggcgatggccctgtccttttaccagacaaccattacctgtccacacaatctgccctttcgaaagatcccaacgaaaagagagaccacatggtccttcttgagtttgtaacagctgctgggattacacatggcatggatgagctctacaaacctagctcaggattgagatcaagatctgctgctgctggtgctggtggtgctgctagagctgctctgcagagaggagtacaagttgaaacaatatcaccaggagatggtcgtacatttccaaaaagaggtcaaacttgtgttgtacattatactggaatgcttgaagatggaaagaaatttgattcatctcgtgatagaaataaaccatttaaatttatgctaggtaaacaagaagtaatacgaggttgggaagaaggagttgctcaaatgagtgtaggtcaaagagcaaaacttactatatctccagattatgcttatggtgcaactggacatccaggtataattccacctcatgcaactcttgtatttgatgtggagcttctaaaactagaaactagaggtgttcaggttgaaacaatttcacctggagatggcagaacctttcctaaaagaggacagacttgcgtagttcattatacaggcatgctagaggatggtaagaaatttgattctagtcgagatagaaataagccattcaagtttatgctaggtaaacaggaagtaataagaggttgggaagagggtgtagcacagatgtcagttggacaaagagcaaagttaacaatatcaccagattatgcatacggtgcaacaggccatcctggcatcatccctccacatgcaactttagtattcgacgttgaattgttaaagttagagacaacgcgtgctagaggtgctgctgctggtgctggaggtgcaggtagacctaggATGCTTGAATTACTAAGAAATGCATATAATAAGTGTAAAGAATACATTTTGGGTTTATCAAAGAAATCTCAAATTTACAAGATTAATATTTTAAACTACTGTTTATTGTTTTCTTTTTTTTTGTTTTACGCTAGTTATTTCTTTTTTATTAACGGTATAATTAATCCTGCGGTTAAAATTGTTTTCCATGGTTCAATAATTGAAGATAAGACAAAAGTTTATGAAAACTCTATAATTGAATCTATTTGGTTGTTATTTACTAAAGGTTCTTACTTTAACATGGCTATGCTGTTATTCTTTTCAATTATTATTCCTTTTTTGAAATTGTTGATGGTTTCAGATAACTTCTATTCTTTTATCGTATTATATAAGATGAATAAAAAGCGTGAGGAAGAAAAAAGAAGAAGAAGAAGAAGAATTAGAGCAAGAGAATATAATAATGCCAAATATAATAAGAATAGATATAAAATGAATAAATACAATAGAAATAAAAATATTAACGATATCTATAAGGATGATATTTATTATTCAGAAAACATTTTTAAAAATGATGAGATCAATTACATTAATACTATTAATGAAAATGAAGAATTTATATTAAAGAAATTTAAGATTTTAAATTTTATTTCTAGATTTCAATTCGTTGATGTTTTTATTTCATTGTTCATTGTTTCTTCACTTAACTTATACTTGTTAGAAGCTAGAATGTTAAATGGTGCTTATTACTTTTTGAACTATTGTATGTTGAGTACTATTTCTTCGTTTTTGTTGTTTTCCTTTACTAGTCTTAAAATCCATATCTTCAAAAATGGTAACATTAAGATTAGCGCTTGTTTAAACGAATCTAATCTAGAGGTCACCACCTCTGGTCCATTATCTACTAAGGATCTGGTCGAAGAGGAAGGTCATGCTCAAATAAACAATCTGATAATTAACGATAAAAATATGACCTCAGGTGTTGTTAACGATTTTAGCGGAAATGGCAATAATGTTGAATTAACAGATGATTTAAAGTCCGGTGAACCACAAAATAGAGACGATATTCAAACTGAAGAAACTAAAAAGGAAAAAATGAATACTAGAACCCATAACGATGAAGATAACACAAAAAAGAAGAACATCAAAGATAAAAAAAAAGCTAACGGTGATGATAAAGTTATTCAAAAGTGTATTGATAACGAAAGAAAAAAAAAACAAAATGGTATGATTCAATCTGTCAATGATGGCGACAAGAACTCTAATTTCAACAATAATAATAATAATAATATTAATGGAGACTCTAATAACAACAATATTAATGGTGACTCAAATAACAATAACATTAATGGTGACTCTAACAACAATAATATTAATGGTGATTCTAATAACAATAATATAAACGGTGATTCTAATAATAATAACATTAATGGGGACTCAAATAATAATAATTATCATAACAATTACCATAATAATTATAGAAATAATTACCATAATAATTATAGAAATAATAATTGTAGGAATAATATTCTAGAACAGAATAAGTGTGATAAAAATGTTCTTTGTTACAATAATATTTACAATACCATGAAAGATAACGATACTTATATATATCTAAAAAAGAACAAATTTAATTCACTGTTAAAGAGTAACTGTATTAAGACTAATTTTAACATGATTAAGATTGGTTATGTCATTTTTTTGTTCGTTTTGTTATGTTTGTGTATTTATTTGATTACTGGTGTTGAATGTTCTTTGTTCGGTATTTATATTTACTTGAGTTATTTTAATTTCAACATTGAAGGTATTTTAATTGATTATATGGATATGTTAAACATCTTAAAATTGAAAATCAAAAAAGGTTACATCTATCCTTTTTTCGTTATGTTACCTTTTATTTTTCCAGTCATTATCTCTATGTGCTTCTTTTTGTCTGTATTCTTTTTGAATATGTATTACGAATCTTTCTCTAAACTTTATAAAAAAATTAGTGAATTAAAAAACGAATTCATCAATTCTTCAGAGAATGATAATGTTAATGAAAGAATCTTGGTCTCTGAAACATCAAATCATTTATGTTTGAATGAATCTAATGATAAAGTATCTAATACTTCTGATGATTTTTTATCTAGAAATAACTCGAACATTTCTTCTTCTAAATCCGAAATGATTAATTCTAATTTTGTTTTTAACAAGTTGTTAAACTTTTATTTCTCTTTCGCAGTTTTCTTTTCATATTTAGGTTCTGCTTTCTTACATATTTCTTTGGGTGAAATTATTTGCATTGCTTTGTTAACTTTTTACCAAATCGTTAAACATACTAATAATTTGAACATTACAATCTTGTTAAAATCTGAAAAAATTAAATTCTGCAAGTTTCTACTGTTTATTTTGTATGGTTTATTGTGTTTCTCTATTAACTTATATGTTAATCAATGGGAAGAATATATTACTAAATTGAAAAGATTAAAGAGAAGAATTTTATTGTTTGAAAAGAATAAATTTTCCGAAATTGTTGATTTGAACACCCAAAAAGGTGATGGTGATCATTTTGATGAAACCCAAATTTTTTCCATTTTTTTTTCCTTCCTGATTAAAAAAAATGAAGGTTCTAAAATGAGAGATAATGACATGAACTCTGATTCTGAAGATTCTATTTATGACGCTTATGAACAACAAATTCAATTGCATCATGGTGACAACATGGTTAATGGTATGTTGATGATGAGAAGAATTTCCATGCAAAATTTAGAAGATGATGAAACACAAGTTGAATATATTAATAGAGAAATCCATACGCAAGGTGATCTACATGTTAGAAGAACAAACCAAGGCATTTTAAGATTCAATATGAGAAGAGGTAAAAAAGGTTCTAATGAAAACATGGGTGTTCACCATGAAAGTGGAAACGTTGATGATGCAAACGGAATGAATAATGTTGACGATACAAATAATATGAATAATGTAGATGGTACTAACAATATGAATAATGTTGATGGTACTAATAATATGAATAATATGGATGGAAGAAACAATATGAATAACATTAATTCAGTCGATAATATGAATAATCTAAACAATAATGATGGGGAAGAGGAAGAAGAATGTGTTAACGACGTGTTGAACTACGATAATAATAATTATGCTATAAATGAAGATGCTGAAGAATATATTAAAAATACTAGCGGAGAAAGAGCTGTTATCATTTGTTCAGAAAAGAGAATTTACGAAAAGAATGGTAATGGTGATATTATTACTAGAAATTATAAGAACGAAGAAAGGTACATATATTTGAAAAAATGGATTCCATTTAAGTCAATGATATTATCAAAGTTGGAAAAAAGAAAAAGAAACAGAAAAGAAGCATATAATACTCCAAGAGTTTTGATATTAATTCATTCTTTTTTGTTCATTTTGATTGTTTTTATTTTCTTAATGGTTTTTTTCAAAAAAGAACCAATATTCAGATTTAATATGCCATCTGTGAATAAAAGATTGAACAACTTCTTTAAATCAACAAGTTTCCATGAAATCATTCCAAACTCAGTTGGCAAATGTAAAACAAAGAAATATATCGCTAAGGAACCATGCTTTAATGTCGGTCATATTTACCATGAAGAAAAGACTTTCTATCATGCTACATTATTGTTTTTACAAGGTCTCAGATCGGTGAAGATAATGAACATGAACTTTTACTATGAGAAAGGTATTTATTACTTATCCTTAGATGGTTATTTTAAACACATTATCGGGCCATTATTTTTGAAACTATGTTTGGGTACTAATTTCTGCCCAATTTCAACTTATGCCTTTCTAGTTGGTAGCAAGCCAACATTTTCAGTCAATGTTGCCGTTCAATGTAACAATAAAAAACCTCCGTACTATATGACTGATATTATTGTAAAAGATTTAAAAATTACTAAAATTGAGATTGTAAAACATTCAGATGTAATTGATAATGTCGATATTAAACTAGATGATGTTCAAGACAGAGTCCAAGAAAAAGTTAATGCAATGTTGGAAGCCAAAAAGAAAATTATTGTCTGGAAGAATCAAAAATACCACTTGGAGGGTTTTTTGAATTATTTGATTTCTAAAAATGCTCTTTCTGGTTTTTCTTGTGAACCAATCAATTACtagggtaccaaggttcggacattcagtattgattgctaacatttgaGCATTTTTAACTTTTCATGTGCATAAGGATTATATGTATATATATATATATATATATATATATTTATATATTTATATATATTTGTATAATTTTATGATATAGAAATAAATAAATAAATAAAGTAAAATAAAAAATAAATATAAAAAAATAATAATAAAATATTCTTAATTATATTTTTTTCATATTCCTAAAATATGTGAATACATTTTGTATATGTCTGTGTACATTTAATAACCACCTTATATATATATATATGTATATATTTATAATATATTTTTTTTTTGAATTTACATGTGTTTTTTTATATTCTAATTTTTTTTTTAATATGTTATTATTAAGACACACACACAAAAAAAAAAAAAAAAAAAAAAAAAAAATTAAATAAAAATTAAATAAAAAATTAATAAACAAAACAATAATGTATATATATATATATATATATATATATATATATATATATGTTTTAGGTTTAATATATCAATTTGGTATTACAAATAAGGGAAGACATATATACATATTATATGTATTGATGAATTAAAAAAAAAAAAAAAATGTTTATATTTAAAAATTATtaaataacttcgtatagcatacattatacgaagttattactcgagggggaagggagaggatcacttcttacctgtggggatgtcgaggaaaatcctggacctatggtgagcaagggcgaggaggataacatggccatcatCaaggagttcatgcgcttcaaggtgcacatggagggctccgtgaacggccacgagttcgaGatcgagggcgagggcgagggccgcccctacgagggcacccagaccgccaagctgaaggtGaccaagggtggccccctgcccttcgcctgggacatcctgtcccctcagttcatgtacggCtccaaggcctacgtgaagcaccccgccgacatccccgactacttgaagctgtccttcccCgagggcttcaagtgggagcgcgtgatgaacttcgaggacggcggcgtggtgaccgtgacCcaggactcctccctgcaggacggcgagttcatctacaaggtgaagctgcgcggcaccaaCttcccctccgacggccccgtaatgcagaagaagaccatgggctgggaggcctcctccgaGcggatgtaccccgaggacggcgccctgaagggcgagatcaagcagaggctgaagctgaaGgacggcggccactacgacgctgaggtcaagaccacctacaaggccaagaagcccgtgcaGctgcccggcgcctacaacgtcaacatcaagttggacatcacctcccacaacgaggactaCaccatcgtggaacagtacgaacgcgccgagggccgccactccaccggcggcatggacgagctgtacaagacgcgtgatccaacaagaagtgcaaatagtggagcaggagcaggagcaggagcaatattaagtagagccggcATGCTTGAGCTTCTTAGAAACGCATACAACAAGTGCAAGGAATACATACTTGGACTTAGTAAGAAGAGTCAGATTTACAAGATTAACATTCTTAACTACTGCCTTCTTTTCAGTTTCTTCCTTTTCTACGCTAGTTACTTCTTCTTCATTAACGGTATTATTAACCCTGCTGTTAAGATAGTATTTCACGGTTCAATAATAGAAGATAAGACAAAGGTATATGAAAATAGTATTATAGAGAGTATTTGGTTGCTTTTCACAAAAGGTTCATACTTCAACATGGCAATGTTATTGTTTTTCAGCATAATTATTCCATTCTTAAAACTTTTAATGGTATCAGACAATTTCTACTCATTCATAGTTCTTTACAAGATGAACAAGAAGAGGGAGGAAGAGAAACGTCGTCGTAGGCGTCGTATTAGAGCTAGGGAGTACAATAACGCAAAGTACAATAAAAACCGTTACAAGATGAACAAGTACAACCGTAACAAAAACATAAACGACATTTATAAAGACGACATTTACTACTCAGAGAACATATTCAAAAACGACGAAATTAACTACATAAACACTATTAACGAAAACGAGGAGTTCATACTTAAAAAGTTCAAGATACTTAACTTTATTAGTAGATTCCAGTTCGTAGACGTTTTCATTAGCCTTTTCATTGTTAGTAGTCTTAACCTTTACCTTCTTGAAGCTCGTATGCTTAACGGAGCTTACTACTTCCTTAACTACTGCATGCTTAGTACTATTAGTAGCTTCCTTCTTTTCTCATTCACAAGCCTTAAGATTCACATTTTCAAGAACGGTAACATAAAAATTAGCGCTTGCTTAAACGAGAGTAACTTGGAAGTAACAACTTCAGGACCATTAAGTACTAAGGACTTAGTTGAGGAGGAGGGTCACGCTCAGATTAACAACTTGATAATAAACGACAAAAACATGACATCAGGAGTAGTAAACGACTTCAGTGGAAACGGcAATAACGTTGAGCTTACTGATGATTTAAAATCAGGAGAGCCACAGAATCGTGACGACATTCAGACAGAGGAGACTAAGAAGGAGAAGATGAACACACGTACACACAACGACGAGGACAACACTAAGAAGAAGAACATTAAAGACAAGAAAAAGGCAAATGGAGACGACAAGGTTATTCAGAAGTGCATAGACAACGAGCGTAAGAAGAAGCAGAACGGAATGATACAGTCAGTTAATGACGGAGACAAGAACTCAAACTTCAACAACAATAACAACAACAACATAAACGGAGACTCAAACAACAACAACATAAACGGAGACTCAAACAACAACAACATAAACGGAGACTCAAACAACAACAACATAAACGGAGACTCAAACAACAACAACATAAACGGAGACTCAAACAACAACAACATAAACGGAGACTCAAACAACAACAACTACCACAACAACTACCACAACAACTACAGAAACAACTACCACAACAACTACAGAAACAACAACTGCCGTAACAACATACTTGAGCAGAACAAGTGCGACAAGAACGTACTTTGCTACAACAACATTTACAACACTATGAAGGACAACGACACATACATTTACCTTAAGAAAAACAAGTTCAACAGTCTTCTTAAGTCAAACTGCATAAAGACAAACTTCAACATGATTAAAATTGGATACGTTATTTTCCTTTTCGTTCTTCTTTGCCTTTGCATTTACCTTATTACTGGTGTTGAGTGTTCACTTTTCGGAATTTACATTTACCTTTCATACTTCAACTTCAACATAGAGGGTATTCTTATAGACTACATGGACATGCTTAACATACTTAAGCTTAAGATTAAAAAGGGTTACATATACCCATTCTTCGTAATGCTTCCTTTCATATTTCCAGTTATTATTAGCATGTGCTTTTTCCTTTCAGTTTTCTTCCTTAACATGTACTACGAGTCATTCAGTAAACTTTACAAGAAAATATCAGAGTTGAAAAACGAGTTTATTAATAGTAGTGAGAACGATAACGTTAATGAGCGTATTCTTGTTAGTGAGACAAGCAACCACCTTTGCCTTAACGAGTCAAATGACAAGGTTAGCAACACTAGTGACGACTTCCTTAGTCGTAACAACTCAAATATTAGCTCAAGTAAGTCAGAGATGATTAACAGTAACTTCGTTTTCAACAAGTTACTTAACTTCTACTTCAGTTTCGCTGTATTTTTCAGCTACCTTGGAAGCGCTTTCCTTCACATTAGTCTTGGAGAGATAATTTGCATAGCTCTTCTTACTTTCTACCAGATAGTTAAACACACAAACAACCTTAACATAACTATACTTTTAAAATCAGAAAAGATAAAGTTCTGCAAGTTTTTACTTTTCATTCTTTACGGACTTCTTTGTTTCAGTATTAACCTTTACGTTAATCAGTGGGAAGAGTACATTACAAAGCTTAAGCGTCTTAAGCGTAGAATACTTCTTTTCGAGAAGAACAAGTTCTCAGAAATAGTTGACTTAAACACACAGAAAGGAGACGGAGATCACTTCGACGAGACTCAGATTTTCAGTATTTTCTTCAGCTTCCTTATTAAGAAGAACGAAGGAAGTAAGATGCGTGACAACGACATGAACTCAGACTCAGAGGATTCAATTTACGACGCATACGAACAACAAATTCAGCTTCATCACGGAGACAACATGGTTAACGGAATGCTTATGATGCGTCGTATTAGTATGCAGAACCTTGAAGATGATGAGACACAGGTAGAGTACATTAACCGTGAAATACACACACAGGGAGACCTTCACGTTAGGCGTACAAACCAGGGAATACTTCGTTTCAACATGCGTCGTGGAAAGAAGGGAAGTAACGAGAACATGGGAGTTCATCACGAATCAGGAAACGTTGACGACGCAAACGGAATGAACAACGTTGACGACACAAACAACATGAACAACGTTGACGGAACAAACAACATGAACAACGTTGACGGAACAAACAACATGAACAACATGGACGGACGTAACAACATGAACAACATTAACTCAGTTGACAACATGAATAACCTTAACAACAACGACGGAGAGGAGGAGGAGGAGTGCGTTAATGACGTACTTAACTACGACAACAACAACTACGCAATAAACGAGGACGCAGAAGAGTACATTAAGAACACAAGTGGTGAGAGGGCAGTAATAATATGCAGCGAGAAGCGTATTTACGAGAAAAATGGTAACGGAGACATTATTACACGTAACTACAAGAACGAGGAGAGGTACATTTACCTTAAGAAGTGGATACCATTCAAGAGCATGATTCTTAGTAAGCTTGAGAAACGTAAGCGTAACCGTAAGGAAGCTTACAACACACCACGTGTTCTTATTCTTATTCACAGTTTCCTTTTCATTTTAATAGTATTTATTTTCCTTATGGTTTTTTTCAAGAAGGAACCAATATTCAGATTCAACATGCCTAGTGTAAACAAAAGACTTAACAACTTCTTCAAGAGTACTTCATTCCATGAGATAATTCCTAACTCAGTTGGAAAGTGCAAGACTAAGAAGTACATTGCTAAGGAGCCTTGCTTCAACGTTGGACACATTTATCACGAGGAAAAAACTTTCTACCACGCTACATTACTTTTCCTTCAGGGACTTCGTAGCGTTAAGATAATGAACATGAACTTCTACTACGAGAAGGGAATTTACTACCTTAGCCTTGACGGTTACTTCAAGCACATAATTGGACCATTATTCTTAAAGCTTTGCCTTGGAACTAATTTCTGCCCAATTTCAACTTACGCTTTCCTTGTTGGTTCAAAGCCTACATTTAGTGTTAACGTAGCAGTACAGTGCAACAACAAGAAGCCTCCTTACTACATGACAGACATAATAGTTAAGGACCTTAAGATAACTAAGATTGAGATTGTAAAGCACAGTGACGTAATAGACAACGTTGACATAAAGCTTGACGACGTTCAGGACCGTGTTCAGGAGAAAGTAAACGCAATGCTTGAGGCTAAGAAGAAGATAATAGTTTGGAAGAACCAGAAATACCACCTTGAAGGATTCTTAAACTACCTTATTAGCAAGAACGCTCTTAGCGGATTCAGTTGCGAGCCAATTAACTACtaggtcgacggatatggcagcttaatgttcgtttttcttatttatatatttataccaattgattgtatttataactgtaaaaatgtgtatgttgtgtgcatatttttttttgtgcatgcacatgcatgtaaatagctaaaattatgaacattttattttttgttcagaaaaaaaaaactttacacacataaaatggctagtatgaatagccatattttatataaattaaatcctatgaatttatgaccatattaaaaatttagatatttatggaacataatatgtttgaaacaataagacaaaattattattattattattatttttactgttataattatgtgtctccttcaatgattcataaatagttggacttgatttttaaaatgtttataatatgattagcatagttaaataaaaaaagttgaaaaattaaaaaaaaacatataaacacaaatgatggtttttccttcaatttcgatatcaatttatagaaacaaaatatatacttgtataattttatttttttatataaatcattacatatataattatacaatattttttctaagagataattatatattaatatatataaaaaaaggtgttttttttttttttttttatttttatttttattttatggtaatattttattttccttattttataaattatattagtttatatgtgattaattttatatattatcaatttatatatttttaaatgcttacttaattatctttttttttttttttttttttttttcccctctttttatattaatttatttttgaaaaaattgatatatatatatatatataatatatatatatacatgtagtagtattaaacaatgtataatatatataaataatatatttatatatttcatttcaattttaattttttttggttttttttttttttctttttgtcatatttaaaaaaaattatattcatataagttatgcattttttataaacattattcaatatatgtataatataatatatatatatatattaatgtattattccaatgtgcatgataaaagaaaaaaataatatttataaaaaaaaagaaaaataaaacaaaaaaagaaaaaaaaaaaaaaaaaaaaaaaaatacaaaaataaataatataatttataattatatattcttgtcacaataaaaatatatatatatatatatatatttataatatgtatattttaaactagaaaaggaataactaatattttatttattatcattcaagatttatattttataataataaatacctaatagaaatatatcaggatccatgcatggttcgctaaactgcatcgtcgctgtgtcccagaacatgggcatcggcaagaacggggactacccctggccaccgctcaggaacgaatttagatatttccagagaatgaccacaacctcttcagtagaaggtaaacagaatctggtgattatgggtaagaagacctggttctccattcctgagaagaatcgacctttaaagggtagaattaatttagttctcagcagagaactcaaggaacctccacaaggagctcattttctttccagaagtctagatgatgccttaaaacttactgaacaaccagaattagcaaataaagtagacatggtctggatagttggtggcagttctgtttataaggaagccatgaatcacccaggccatcttaaactatttgtgacaaggatcatgcaagactttgaaagtgacacgttttttccagaaattgatttggagaaatataaacttctgccagaatacccaggtgttctctctgatgtccaggaggagaaaggcattaagtacaaatttgaagtatatgagaagaatgattaagcttatttaataatagattaaaaatattataaaaataaaaacataaacacagaaattacaaaaaaaatacatatgaattttttttttgtaatcttccttataaatatagaataatgaatcatataaaacatatcattattcatttatttacatttaaaattattgtttcagtatctttaatttattatgtatatataaaaataacttacaattttattaataaacaatatatgtttattaattcatgttttgtaatttatgggatagcgattttttttactgtctgtatttttcttttttaattatgttttaattgtattttatttttattattgttctttttatagtattattttaaaacaaaatgtattttctaagaacttataataataataaatataaattttaataaaaattatatttatcttttacaatatgaacataaagtacaacattaatatatagcttttaatatttttattcctaatcatgtaaatcttaaatttttctttttaaacatatgttaaatatttatttctcattatatataagaacatatttattaaatctagaattctatagtgagtcgtattacaattcactggccgtcgttttacaacgtcgtgactgggaaaaccctggcgttacccaacttaatcgccttgcagcacatccccctttcgccagctggcgtaatagcgaagaggcccgcaccgatcgcccttcccaacagttgcgcagcctgaatggcgaatggcgcctgatgcggtattttctccttacgcatctgtgcggtatttcacaccgcatatggtgcactctcagtacaatctgctctgatgccgcatagttaagccagccccgacacccgccaacacccgctgacgcgccctgacgggcttgtctgctcccggcatccgcttacagacaagctgtgaccgtctccgggagctgcatgtgtcagaggttttcaccgtcatcaccgaaacgcgcgagacgaaagggcctcgtgatacgcctatttttataggttaatgtcatgataataatggtttcttagacgtcaggtggcacttttcggggaaatgtgcgcggaacccctatttgtttatttttctaaatacattcaaatatgtatccgctcatgagacaataaccctgataaatgcttcaataatattgaaaaaggaagagtatgagtattcaacatttccgtgtcgcccttattcccttttttgcggcattttgccttcctgtttttgctcacccagaaacgctggtgaaagtaaaagatgctgaagatcagttgggtgcacgagtgggttacatcgaactggatctcaacagcggtaagatccttgagagttttcgccccgaagaacgttttccaatgatgagcacttttaaagttctgctatgtggcgcggtattatcccgtattgacgccgggcaagagcaactcggtcgccgcatacactattctcagaatgacttggttgagtactcaccagtcacagaaaagcatcttacggatggcatgacagtaagagaattatgcagtgctgccataaccatgagtgataacactgcggccaacttacttctgacaacgatcggaggaccgaaggagctaaccgcttttttgcacaacatgggggatcatgtaactcgccttgatcgttgggaaccggagctgaatgaagccataccaaacgacgagcgtgacaccacgatgcctgtagcaatgccaacaacgttgcgcaaactattaactggcgaactacttactctagcttcccggcaacaattaatagactggatggaggcggataaagttgcaggaccacttctgcgctcggcccttccggctggctggtttattgctgataaatctggagccggtgagcgtgggtctcgcggtatcattgcagcactggggccagatggtaagccctcccgtatcgtagttatctacacgacggggagtcaggcaactatggatgaacgaaatagacagatcgctgagataggtgcctcactgattaagcattggtaactgtcagaccaagtttactcatatatactttagattgatttaaaacttcatttttaatttaaaaggatctaggtgaagatcctttttgataatctcatgaccaaaatcccttaacgtgagttttcgttccactgagcgtcagaccccgtagaaaagatcaaaggatcttcttgagatcctttttttctgcgcgtaatctgctgcttgcaaacaaaaaaaccaccgctaccagcggtggtttgtttgccggatcaagagctaccaactctttttccgaaggtaactggcttcagcagagcgcagataccaaatactgtccttctagtgtagccgtagttaggccaccacttcaagaactctgtagcaccgcctacatacctcgctctgctaatcctgttaccagtggctgctgccagtggcgataagtcgtgtcttaccgggttggactcaagacgatagttaccggataaggcgcagcggtcgggctgaacggggggttcgtgcacacagcccagcttggagcgaacgacctacaccgaactgagatacctacagcgtgagctatgagaaagcgccacgcttcccgaagggagaaaggcggacaggtatccggtaagcggcagggtcggaacaggagagcgcacgagggagcttccagggggaaacgcctggtatctttatagtcctgtcgggtttcgccacctctgacttgagcgtcgatttttgtgatgctcgtcaggggggcggagcctatcgaaaaacgccagcaacgcggcctttttacggttcctggccttttgctggccttttgctcacatgttctttcctgcgttatcccctgattctgtggataaccgtattaccgcctttgagtgagctgataccgctcgccgcagccgaacgaccgagcgcagcgagtcagtgagcgaggaagcggaagagcgcccaatacgcaaaccgcctctccccgcgcgttggccgattcattaatgcagctggcacgacaggtttcccgactggaaagcgggcagtgagcgcaacgcaattaatgtgagttagctcactcattaggcaccccaggctttacactttatgcttccggctcgtatgttgtgtggaattgtgagcggataacaatttcacacaggaaacagctatgaccatgattacgccaagctatttaggtgacactatagaatactcaagctgcggccgcTAGTTGGAATTATTACGTAATGCTTATAATAAATGTAAAGAGTATATTTTGGGCTTATCGAAAAAATCACAAATATATAAAATAAATATATTAAATTATTGTTTATTATTTTCGTTTTTTTTATTTTATGCATCATATTTTTTTTTTATAAATGGAATAATAAATCCAGCAGTAAAAATTGTTTTCCATGGAAGTAAGTACCTTATATGAAAAAATGAAAGTGTATCCATAAAAGAAGATATACACAAAGGTGTATATATATATATATATATATATATATATATATATATGTATGTATGTATGTATTTATTTATTTATTTATTATTTATATATATATTTTCTTTATATTTTCTTTAAGGTATTATTGAGGACAAAACCAAAGTTTACGAGAACTCGATAATTGAATCAATATGGCTATTGTTTACCAAGGGAAGTTATTTTAATATGGCCATGCTTCTATTCTTTTCTATTATAATACCGTTTCTCAAGTTACTTATGGTTAGTGATAACTTTTATAGTTTTATCGTATTATATAAAATGAATAAAAAACGAGAAGAGGAAAAGAGAAGGAGACGAAGAAGAATACGTGCACGAGAATATAACAATGCTAAATATAACAAGAATAGATATAAAATGAATAAATATAATAGAAATAAGAATATTAATGATATATACAAGGATGATATATATTATAGTGAAAATATTTTTAAGAATGATGAGATAAATTATATTAATACAATAAATGAGAATGAAGAATTTATTTTAAAGAAATTTAAAATTTTAAATTTCATATCGCGTTTTCAATTTGTTGATGTATTTATATCTTTATTTATAGTATCATCATTAAATTTATATTTATTAGAGGCAAGGATGTTAAATGGTGCATATTATTTTTTGAATTATTGTATGTTATCAACAATATCATCTTTTTTGTTATTTAGTTTTACTTCTTTAAAAATACATATATTTAAAAATGGAAATATTAAGATATCTGCATGTCTCAATGAATCAAATCTAGAGGTTACTACAAGCGGTCCCCTTTCAACAAAAGATCTTGTCGAAGAAGAAGGACATGCACAAATAAATAATCTAATTATTAATGATAAGAATATGACTAGTGGTGTTGTTAATGATTTTTCAGGTAATGGAAACAATGTCGAATTAACAGACGACCTTAAGAGTGGCGAACCCCAAAACAGAGATGATATACAAACTGAAGAAACAAAAAAAGAAAAAATGAATACTAGAACTCATAATGATGAAGATAATACAAAAAAAAAAAATATAAAGGATAAAAAGAAAGCTAACGGTGATGATAAAGTAATACAAAAATGTATTGATAATGAAAGGAAAAAAAAACAAAATGGTATGATTCAAAGTGTAAACGATGGTGATAAAAATAGTAATTTTAATAATAACAATAATAATAATATTAATGGTGATAGTAATAATAATAATATTAATGGTGATAGTAATAATAATAATATTAATGGTGATAGTAATAATAATAATATTAATGGTGATAGTAATAATAATAATATTAATGGTGATAGTAATAATAATAATATTAATGGTGATAGTAATAATAATAATTATCATAATAATTATCATAATAATTATCGTAATAATTATCATAATAATTATCGTAATAATAATTGTAGGAATAATATTTTAGAACAAAATAAATGTGATAAAAATGTTTTGTGTTATAATAATATATATAATACAATGAAAGATAATGATACTTATATATATTTAAAAAAGAATAAATTTAATTCGTTATTAAAAAGTAATTGTATCAAAACTAATTTTAATATGATAAAGATAGGTTATGTAATATTTTTATTTGTGTTATTATGTTTATGTATATATTTAATAACAGGAGTAGAATGCAGTTTATTTGGTATATATATATATTTGAGTTATTTTAATTTTAATATTGAAGGAATATTAATTGATTATATGGATATGTTAAATATTTTAAAATTAAAAATAAAGAAAGGATATATCTATCCTTTTTTTGTTATGTTACCATTTATTTTCCCTGTAATAATATCTATGTGTTTCTTTTTAAGTGTATTTTTTTTAAATATGTATTATGAAAGCTTTTCAAAGTTATATAAAAAGATCAGTGAACTAAAGAATGAATTCATAAACTCCTCAGAAAATGACAATGTGAACGAAAGAATATTGGTATCCGAAACTTCTAATCATTTATGTTTAAATGAAAGTAACGATAAAGTATCTAATACATCAGATGATTTTTTATCAAGAAATAATAGCAACATATCTAGTTCAAAAAGTGAAATGATAAATTCAAATTTTGTATTTAATAAACTTTTGAATTTTTATTTTTCATTTGCAGTTTTCTTTTCTTATTTAGGTTCTGCATTTTTACATATATCCTTAGGTGAAATTATATGTATCGCATTATTAACATTTTATCAAATTGTAAAGCATACCAATAATTTGAATATCACAATTTTACTTAAGAGTGAGAAAATTAAATTTTGTAAATTCCTTTTGTTTATATTATATGGGTTGTTATGCTTTTCCATAAATTTGTATGTAAACCAATGGGAGGAATATATAACGAAATTAAAAAGGTTAAAAAGACGTATTTTATTGTTTGAAAAAAATAAATTTAGTGAGATTGTGGATCTGAATACTCAAAAGGGTGATGGTGACCATTTTGATGAAACACAAATATTTTCCATATTTTTTTCTTTTTTAATAAAAAAAAATGAGGGTTCAAAAATGAGGGATAATGATATGAATAGTGATAGCGAAGACAGTATATATGATGCTTATGAGCAGCAGATACAATTACACCATGGTGATAATATGGTCAATGGTATGTTGATGATGAGAAGGATATCGATGCAAAATTTAGAGGACGACGAAACCCAAGTTGAATATATAAATAGGGAGATCCATACGCAAGGTGATTTACATGTACGAAGAACGAATCAAGGTATTTTAAGATTTAATATGAGAAGGGGCAAAAAAGGGTCCAATGAAAATATGGGTGTCCACCATGAGAGTGGTAATGTGGATGATGCGAATGGTATGAATAATGTGGATGATACGAATAATATGAATAATGTGGATGGTACGAATAATATGAATAATGTGGATGGTACGAATAATATGAATAATATGGATGGTAGGAATAATATGAATAATATAAATAGTGTGGATAATATGAACAATTTAAATAATAATGATGGTGAAGAAGAAGAAGAATGTGTGAACGATGTTTTAAATTATGATAATAATAATTATGCTATTAATGAAGATGCTGAGGAATATATAAAAAATACGTCAGGAGAACGAGCTGTTATTATTTGTTCTGAAAAAAGAATATATGAAAAGAACGGAAATGGTGATATAATAACTAGAAATTATAAAAATGAAGAACGATATATATATTTAAAAAAATGGATTCCTTTTAAATCTATGATATTGTCAAAATTAGAAAAGAGGAAAAGGAATAGAAAAGAGGCATATAATACTCCTAGAGTATTAATATTAATACATTCCTTTTTATTTATACTTATTGTTTTCATATTTTTAATGGTATTCTTTAAAAAAGAGCCTATTTTTCGTTTTAATATGCCATCAGTTAATAAGCGTTTAAATAATTTTTTTAAATCAACAAGTTTTCACGAAATTATACCAAATAGTGTAGGGAAATGTAAAACAAAAAAATATATAGCAAAAGAACCATGTTTTAATGTGGGTCATATATACCATGAAGAGAAGACATTTTATCATGCAACTCTTTTATTTTTACAAGGTTTAAGATCTGTAAAAATTATGAATATGAATTTTTATTATGAAAAAGGCATATATTATTTATCTTTGGATGGATATTTTAAACATATTATAGGTCCTCTCTTTCTTAAATTATGTTTAGGTACAAACTTTTGTCCCATAAGTACATATGCATTTTTAGTAGGAAGTAAACCAACTTTCTCAGTAAATGTTGCTGTTCAATGTAATAATAAAAAACCACCATATTATATGACGGATATTATTGTAAAAGATTTAAAAATTACAAAAATAGAAATAGTTAAACATTCAGATGTTATCGATAATGTGGATATTAAATTAGATGATGTACAAGATAGAGTACAAGAAAAGGTTAATGCCATGTTAGAAGCAAAAAAAAAAATTATCGTCTGGAAAAATCAAAAGTATCATTTGGAGGGTTTTCTCAATTATTTAATATCTAAAAATGCATTATCTGGGTTTTCCTGTGAACCCATAAATTATTGAacttgacaaattaaaacacgttaaatgtatatatgtatatatattatatatgtatatatttatttatatttttcttacatttccatttgggtgaacatatattcttacccttttcataaattgttttttttttttttgatatttatcaataatatattcagtaaaaatatattatttcacataatttaatattttttggaataacatataaatgtcaatgtatatgtgtatatatatatatatatatatatatgtatatatttttttttctttttcatatatttatatgcgcccatttgtttttatctatatatatatatatttatatatccgaatttctttatattgttatattccgatttgattatatataaaattttttttaattcataatttttttttttttttttttcttttttttcatttccatgtgatttatttcaatccgatattatttatatgttatatatatattattctataatatgataaaaggatatttttttttcttatttgtatttttagattataaatataaagatattcttatatatagattatatctgcttttttttttttttttttttttttttctattattaatttttaaattatctacttacgaattttttaattgttgatatggccttcatgagtatattattatgcatgaaagaaaaaaaaaaaaaaaaatatatatatgtatataatatatatatatatatatataatataccaatttttattatcttatattttacatgcactataaatgcatagaatttttcttttttttttttttgatttcttctccccaaatttttggttatttatataatatacgtattataaatatgtacatataatatgtatatattataatgtgtaattttccacatatatatataaaatatatatatatatatatatatattttaataatacttgaacttttaaaattaaatatataaaaaggaatccaattaatttatggttatattatttgttattcactagaaaattctctagaatttaataaaacatatat

**5’ amplicon size: 1149bps**

**3’ amplicon size: 1121bps**

## > PF3D7_1146200

TatttttttgttagttcgtttgttttttctttcttttttttttttttttttaatttcttataaacttttctttttaattataaacgttctgacaataattcttaaaacttgtgatttctaaattttatacctacataatatatatatgtatatatatatatataaatatatagatatagaataaataataaatatattaatatatccgatttacttttaatatatataatattaattatttctttgtgatgtacaaataaacatgttggtacgacaaaaataaaaaacactagttctttataaagtaccccaaaaatatgtacaaaatattacctatatatattttttcatatttaaatataatatgaagtgataaagcaaattcattattttttccaagaaaatttatataaatatatatatatgtacacactccaatacatatacagcatctttataaataaataacatattcatatttttgtaattttttttttttttttttttttccgacgaattgttttaacttttctgaatcatataaaaATG**AGTGATGATATTATGAATGGTGATACTCTCTCAGAGTATAGAAAGATTCATAAAAAAAATTATGAGGAGAGAATAATAAAAGAAAATGAATTAATAGAAAAACAAAAAACGGAAGAATTATTAAATGAAAAAAAAAAAAACGAAGAAATTTATAGACAACTACAAATAGATAAAGATAGAGAATACTCTAAACAATGTTATTATGAAATTAATAATTTTCAAAACATACATGAAAGAACTTACAATGATTCAGATATTAGAGAAAATTATACACAAAGATTAATTAACCCTTCGGCATATTCTTATATTGATAATAACATGACACAAAATTTGTTAGATGGTGATATGGGATATACTTTTTCTCAAAGAATTAGACGTTTTTTCCTGAACAA*gtaatcaaaaaaaaaaaatatataaatataaaaatataaatatataaaattaatatatatatatatatatatattattttcatcttttatag*CAATATCAGGCAGTGCTTT**GTTTGTACAACGGTTCAGTTGATATGTAACGTTGTCCTTATTATTTTATTAATTGCATTTATAATAATATTTGGCTTGTCATGAagtttttttaatagtttatatttttctaccgttttaatttttttaattcacacattattccatccttttattatattattttttttagtc

**Original Locus (OL) amplicon size: 657bp**

### > Halo-PF3D7_1146200 Halo yDHODH recodonizedsequence

tatttttttgttagttcgtttgttttttctttcttttttttttttttttttaatttcttataaacttttctttttaattataaacgttctgacaataattcttaaaacttgtgatttctaaattttatacctacataatatatatatgtatatatatatatataaatatatagatatagaataaataataaatatattaatatatccgatttacttttaatatatataatattaattatttctttgtgatgtacaaataaacatgttggtacgacaaaaataaaaaacactagttctttataaagtaccccaaaaatatgtacaaaatattacctatatatattttttcatatttaaatataatatgaagtgataaagcaaattcattattttttccaagaaaatttatataaatatatatatatgtacacactccaatacatatacagcatctttataaataaataacatattcatatttttgtaattttttttttttttttttttttccgacgaattgttttaacttttctgaatcatataaaaATG**AGTGATGATATTATGAATGGTGATACTCTCTCAGAGTATAGAAAGATTCATAAAAAAAATTATGAGGAGAGAATAATAAAAGAAAATGAATTAATAGAAAAACAAAAAACGGAAGAATTATTAAATGAAAAAAAAAAAAACGAAGAAATTTATAGACAACTACAAATAGATAAAGATAGAGAATACTCTAAACAATGTTATTATGAAATTAATAATTTTCAAAACATACATGAAAGAACTTACAATGATTCAGATATTAGAGAAAATTATACACAAAGATTAATTAACCCTTCGGCATATTCTTATATTGATAATAACATGACACAAAATTTGTTAGATGGTGATATGGGATATACTTTTTCTCAAAGAATTAGACGTTTTTTCCTGAACAAgtaatcaaaaaaaaaaaatatataaatataaaaatataaatatataaaattaatatatatatatatatatatattattttcatcttttatagCAATATCAGGCAGTGCTTT**GgtttaaacgagcagaagttaatatcagaagaggatttgggtgaacaaaaactcataagcgaagaagatttaataacttcgtatagcatacattatacgaagttatccggagaaggaagaggaagtttattaacatgtggagatgtagaagaaaatccaggaccaATGACAGCCAGTTTAACTACCAAGTTCTTGAACAATACCTATGAAAACCCATTTATGAATGCATCCGGTGTTCATTGCATGACTACACAAGAATTAGATGAATTAGCAAACTCTAAAGCTGGCGCATTCATTACAAAGAGTGCTACAACCTTAGAAAGAGAAGGTAACCCTGAACCACGTTACATTTCTGTCCCTCTAGGCAGTATCAACTCCATGGGTTTACCAAACGAAGGTATCGACTACTATTTGTCCTATGTATTAAACCGTCAAAAGAATTATCCTGATGCACCTGCTATTTTCTTCTCAGTTGCTGGTATGAGCATTGATGAAAATTTAAATTTGTTGAGGAAAATCCAAGATAGCGAATTCAACGGTATTACCGAGTTAAACTTGTCTTGTCCTAATGTGCCTGGGAAACCACAAGTTGCTTATGACTTTGACTTGACAAAGGAAACCTTGGAAAAGGTTTTTGCCTTTTTCAAAAAACCTCTTGGTGTCAAGTTGCCTCCTTATTTTGATTTTGCCCATTTTGATATCATGGCAAAAATATTGAACGAGTTCCCATTAGCTTATGTCAACTCTATCAATAGTATAGGAAATGGTCTTTTCATTGATGTGGAGAAGGAGAGTGTAGTAGTGAAGCCAAAGAATGGTTTCGGGGGTATTGGAGGTGAATATGTTAAGCCAACCGCGCTCGCCAATGTTCGTGCATTTTACACTCGTTTGAGACCTGAAATCAAAGTTATCGGTACAGGTGGAATTAAGTCCGGTAAGGATGCATTTGAACATCTTCTATGTGGTGCCTCTATGCTACAGATTGGTACAGAATTACAAAAAGAGGGCGTCAAGATTTTTGAACGTATCGAAAAAGAATTAAAAGACATAATGGAAGCTAAGGGTTATACATCCATAGATCAGTTCCGTGGGAAGTTGAACAGCATTGGTGAAGGTAGAGGTTCTTTGTTGACTTGTGGTGATGTTGAAGAAAATCCAGGTCCAGCTAGCATGGCAGAAATTGGTACGGGTTTTCCATTTGATCCTCATTATGTGGAGGTGCTGGGGGAAAGGATGCATTATGTTGACGTAGGACCAAGAGATGGTACTCCAGTGTTATTTTTGCATGGAAACCCAACCTCGAGTTATGTATGGAGAAATATAATTCCACATGTAGCACCAACACATAGATGTATAGCTCCTGATTTAATTGGTATGGGAAAAAGTGATAAACCTGACTTAGGATATTTTTTTGATGATCATGTCCGTTTTATGGATGCTTTCATTGAAGCTCTTGGCCTTGAAGAAGTAGTATTAGTTATACATGATTGGGGATCCGCCTTAGGATTTCATTGGGCCAAGAGGAATCCTGAAAGAGTAAAAGGAATAGCATTCATGGAATTCATACGACCAATCCCCACATGGGATGAATGGCCAGAATTTGCACGCGAAACATTTCAAGCTTTTAGAACTACAGATGTTGGTAGAAAATTAATAATAGATCAAAATGTATTTATAGAAGGAACTTTACCTATGGGTGTTGTAAGGCCGTTAACAGAAGTTGAAATGGACCACTACCGTGAACCTTTTTTAAATCCAGTAGATAGAGAGCCCTTATGGAGATTTCCTAATGAATTACCTATTGCAGGTGAACCCGCGAATATTGTTGCTTTAGTAGAAGAATATATGGATTGGTTACATCAGTCTCCTGTTCCTAAACTTCTATTTTGGGGTACACCTGGAGTTCTAATACCACCAGCTGAAGCAGCAAGATTAGCAAAATCATTACCAAATTGTAAAGCTGTTGATATAGGTCCTGGGTTGAATTTATTACAAGAAGATAATCCAGATTTGATTGGATCTGAGATAGCTAGATGGCTAAGTACATTAGAAATTTCAGGTACGCGTGCTAGAGGTGCTGCTGCTGGTGCTGGAGGTGCAGGTAGACCTAGGATGTCTGATGATATAATGAATGGAGATACATTAAGTGAATATAGAAAAATACATAAAAAAAATTATGAAGAAAGAATAATAAAAGAAAATGAATTAATAGAAAAACAAAAAACAGAAGAATTATTAAATGAAAAGAAAAAAAATGAAGAAATTTATAGACAATTACAAATAGATAAAGATAGAGAATATAGTAAACAATGTTATTATGAAATTAATAATTTTCAAAATATACACGAAAGAACATATAATGATTCTGATATAAGAGAAAATTATACACAAAGACTTATAAATCCTTCTGCTTATAGTTATATTGATAATAATATGACACAAAATTTATTAGATGGTGATATGGGTTATACATTTTCACAAAGGATTAGAAGATTTTTTTTAAATAATAATATAAGACAATGTTTCGTATGTACGACTGTTCAATTAATTTGTAATGTAGTATTAATAATATTATTAATAGCATTTATAATAATATTTGGATTAAGTTAAaggcctataacttcgtatagcatacattatacgaagttattatgactcgagggatatggcagcttaatgttcgtttttcttatttatatatttataccaattgattgtatttataactgtaaaaatgtgtatgttgtgtgcatatttttttttgtgcatgcacatgcatgtaaatagctaaaattatgaacattttattttttgttcagaaaaaaaaaactttacacacataaaatggctagtatgaatagccatattttatataaattaaatcctatgaatttatgaccatattaaaaatttagatatttatggaacataatatgtttgaaacaataagacaaaattattattattattattatttttactgttataattatgtgtctccttcaatgattcataaatagttggacttgatttttaaaatgtttataatatgattagcatagttaaataaaaaaagttgaaaaattaaaaaaaaacatataaacacaaatgatggtttttccttcaatttcgatatcaatttatagaaacaaaatatatacttgtataattttatttttttatataaatcattacatatataattatacaatattttttctaagagataattatatattaatatatataaaaaaaggtgttttttttttttttttttatttttatttttattttatggtaatattttattttccttattttataaattatattagtttatatgtgattaattttatatattatcaatttatatatttttaaatgcttacttaattatctttttttttttttttttttttttttcccctctttttatattaatttatttttgaaaaaattgatatatatatatatatataatatatatatatacatgtagtagtattaaacaatgtataatatatataaataatatatttatatatttcatttcaattttaattttttttggttttttttttttttctttttgtcatatttaaaaaaaattatattcatataagttatgcattttttataaacattattcaatatatgtataatataatatatatatatatattaatgtattattccaatgtgcatgataaaagaaaaaaataatatttataaaaaaaaagaaaaataaaacaaaaaaagaaaaaaaaaaaaaaaaaaaaaaaaatacaaaaataaataatataatttataattatatattcttgtcacaataaaaatatatatatatatatatatatttataatatgtatattttaaactagaaaaggaataactaatattttatttattatcattcaagatttatattttataataataaatacctaatagaaatatatcaggatccatgcatggttcgctaaactgcatcgtcgctgtgtcccagaacatgggcatcggcaagaacggggactacccctggccaccgctcaggaacgaatttagatatttccagagaatgaccacaacctcttcagtagaaggtaaacagaatctggtgattatgggtaagaagacctggttctccattcctgagaagaatcgacctttaaagggtagaattaatttagttctcagcagagaactcaaggaacctccacaaggagctcattttctttccagaagtctagatgatgccttaaaacttactgaacaaccagaattagcaaataaagtagacatggtctggatagttggtggcagttctgtttataaggaagccatgaatcacccaggccatcttaaactatttgtgacaaggatcatgcaagactttgaaagtgacacgttttttccagaaattgatttggagaaatataaacttctgccagaatacccaggtgttctctctgatgtccaggaggagaaaggcattaagtacaaatttgaagtatatgagaagaatgattaagcttatttaataatagattaaaaatattataaaaataaaaacataaacacagaaattacaaaaaaaatacatatgaattttttttttgtaatcttccttataaatatagaataatgaatcatataaaacatatcattattcatttatttacatttaaaattattgtttcagtatctttaatttattatgtatatataaaaataacttacaattttattaataaacaatatatgtttattaattcatgttttgtaatttatgggatagcgattttttttactgtctgtatttttcttttttaattatgttttaattgtattttatttttattattgttctttttatagtattattttaaaacaaaatgtattttctaagaacttataataataataaatataaattttaataaaaattatatttatcttttacaatatgaacataaagtacaacattaatatatagcttttaatatttttattcctaatcatgtaaatcttaaatttttctttttaaacatatgttaaatatttatttctcattatatataagaacatatttattaaatctagaattctatagtgagtcgtattacaattcactggccgtcgttttacaacgtcgtgactgggaaaaccctggcgttacccaacttaatcgccttgcagcacatccccctttcgccagctggcgtaatagcgaagaggcccgcaccgatcgcccttcccaacagttgcgcagcctgaatggcgaatggcgcctgatgcggtattttctccttacgcatctgtgcggtatttcacaccgcatatggtgcactctcagtacaatctgctctgatgccgcatagttaagccagccccgacacccgccaacacccgctgacgcgccctgacgggcttgtctgctcccggcatccgcttacagacaagctgtgaccgtctccgggagctgcatgtgtcagaggttttcaccgtcatcaccgaaacgcgcgagacgaaagggcctcgtgatacgcctatttttataggttaatgtcatgataataatggtttcttagacgtcaggtggcacttttcggggaaatgtgcgcggaacccctatttgtttatttttctaaatacattcaaatatgtatccgctcatgagacaataaccctgataaatgcttcaataatattgaaaaaggaagagtatgagtattcaacatttccgtgtcgcccttattcccttttttgcggcattttgccttcctgtttttgctcacccagaaacgctggtgaaagtaaaagatgctgaagatcagttgggtgcacgagtgggttacatcgaactggatctcaacagcggtaagatccttgagagttttcgccccgaagaacgttttccaatgatgagcacttttaaagttctgctatgtggcgcggtattatcccgtattgacgccgggcaagagcaactcggtcgccgcatacactattctcagaatgacttggttgagtactcaccagtcacagaaaagcatcttacggatggcatgacagtaagagaattatgcagtgctgccataaccatgagtgataacactgcggccaacttacttctgacaacgatcggaggaccgaaggagctaaccgcttttttgcacaacatgggggatcatgtaactcgccttgatcgttgggaaccggagctgaatgaagccataccaaacgacgagcgtgacaccacgatgcctgtagcaatgccaacaacgttgcgcaaactattaactggcgaactacttactctagcttcccggcaacaattaatagactggatggaggcggataaagttgcaggaccacttctgcgctcggcccttccggctggctggtttattgctgataaatctggagccggtgagcgtgggtctcgcggtatcattgcagcactggggccagatggtaagccctcccgtatcgtagttatctacacgacggggagtcaggcaactatggatgaacgaaatagacagatcgctgagataggtgcctcactgattaagcattggtaactgtcagaccaagtttactcatatatactttagattgatttaaaacttcatttttaatttaaaaggatctaggtgaagatcctttttgataatctcatgaccaaaatcccttaacgtgagttttcgttccactgagcgtcagaccccgtagaaaagatcaaaggatcttcttgagatcctttttttctgcgcgtaatctgctgcttgcaaacaaaaaaaccaccgctaccagcggtggtttgtttgccggatcaagagctaccaactctttttccgaaggtaactggcttcagcagagcgcagataccaaatactgtccttctagtgtagccgtagttaggccaccacttcaagaactctgtagcaccgcctacatacctcgctctgctaatcctgttaccagtggctgctgccagtggcgataagtcgtgtcttaccgggttggactcaagacgatagttaccggataaggcgcagcggtcgggctgaacggggggttcgtgcacacagcccagcttggagcgaacgacctacaccgaactgagatacctacagcgtgagctatgagaaagcgccacgcttcccgaagggagaaaggcggacaggtatccggtaagcggcagggtcggaacaggagagcgcacgagggagcttccagggggaaacgcctggtatctttatagtcctgtcgggtttcgccacctctgacttgagcgtcgatttttgtgatgctcgtcaggggggcggagcctatcgaaaaacgccagcaacgcggcctttttacggttcctggccttttgctggccttttgctcacatgttctttcctgcgtAtatcccctgattctgtggataaccgtattaccgcctttgagtgagctgataccgctcgccgcagccgaacgaccgagcgcagcgagtcagtgagcgaggaagcggaagagcgcccaatacgcaaaccgcctctccccgcgcgttggccgattcattaatgcagctggcacgacaggtttcccgactggaaagcgggcagtgagcgcaacgcaattaatgtgagttagctcactcattaggcaccccaggctttacactttatgcttccggctcgtatgttgtgtggaattgtgagcggataacaatttcacacaggaaacagctatgaccatgattacgccaagctatttaggtgacactatagaatactcgcggccgcTAGAGTGATGATATTATGAATGGTGATACTCTCTCAGAGTATAGAAAGATTCATAAAAAAAATTATGAGGAGAGAATAATAAAAGAAAATGAATTAATAGAAAAACAAAAAACGGAAGAATTATTAAATGAAAAAAAAAAAAACGAAGAAATTTATAGACAACTACAAATAGATAAAGATAGAGAATACTCTAAACAATGTTATTATGAAATTAATAATTTTCAAAACATACATGAAAGAACTTACAATGATTCAGATATTAGAGAAAATTATACACAAAGATTAATTAACCCTTCGGCATATTCTTATATTGATAATAACATGACACAAAATTTGTTAGATGGTGATATGGGATATACTTTTTCTCAAAGAATTAGACGTTTTTTCCTGAACAAgtaatcaaaaaaaaaaaatatataaatataaaaatataaatatataaaattaatatatatatatatatatatattattttcatcttttatagCAATATCAGGCAGTGCTTTGTTTGTACAACGGTTCAGTTGATATGTAACGTTGTCCTTATTATTTTATTAATTGCATTTATAATAATATTTGGCTTGTCATGAagtttttttaatagtttatatttttctaccgttttaatttttttaattcacacattattccatccttttattatattattttttttagtc

**5’ amplicon size: 857bps**

**3’ amplicon size: 631bps**

## > PF3D7_1217500

tattttcttctcaataccaaaaatttttataaataaatactaataaataaaatattatatttactttattatataaaaaatatatatatatatatatatatatatatattaaaaacgtacaaaataagaattaattttttttattttatataaatttaaatattgatgatatcatattatttatattatattattataaatataatgatttaaaaaagttaaaaaaaaaaaagtatatatatatatatatatatatatacatataaaggttccaatttttatatgaagtatgagtttatatataaatataaagaaattttttttttgcatgtttcttgcatgtttctctatgcataaaaatttatttatatatatatatatatactatacatataaattgaaaaagaatatacataaaattaatatataatatatttataatattatataacttttataaaataatgaaatatgcaaaaagaaaaaaaaaaaaaaaaaaaaaaaaagaaaaatacaaatatattaaaatttttataataatatataaaatataactatattatatatatatataattgtattgaataatatatttattttttaacataaattatggaatctttttttttttttacatattttataactttttctatatttttctaaaaaaaataatatattttattttttgctttttattctatatatatatatatatatatatatatataatatataacaatttttagatttatttatatgttattatatattatatataaatttttatttaaagaaaaaaaaattgaaactttttttttctttttcaaaactgttattatatatatatatatatatatataagaagtttatatatatataataataattataataatacagaatacttaattttttttcataatatgaaaaaaggatttttttaaacaataaatatatatatatatattttgaaagtctggagtattttataagaggtaccattgaaaatattatatatatatatatatatatatatatatatatatatgtatttatttatttatttattaattttacacttcacattagttatatatatagtatatatttatttatatgaacttttttttttatgaaagaaaaaatatttgtttatattattttataaataaattttatatatgtaatttattgtataattgctttctttttttttcgtttcttttatttttttttaattaaatataaaagaaagctattataaaaagaaaaaatataggagtatattaatataatgtacaaacgtttattttgtatgtaattttttttatttacgttgcctatattattatccatattttctatactataaaaaagaaaaaatatataaagagtctatttattttgaatagaaagaaaaaataaaaaaaatagtctttttatagtaaaattagagtgtatatatatatatatatatatataatatttaattatatacacaaaaatatatatgtgaagttttaccttattattcttatatattatatcttattattatattttaatttttaatcctatcacattatatatatatatatatataatatatatatattttgtcttaataacgcaaaaaaaaaaaaaaaataaataatatattttatatataATG**ACTATTACACTGGAAAGTTTAAGTGAGGAAAAGTTACAGAATTATAACAATAAAAGAAAATATAATTGTCGTATGACATTTAAAAGATTTACAAAAAAGAGATTTAAAAATAAAGATAAAAATAAAGAAGGAAAAATTAATATTAGAACGTCCTTAATTAAAAATGAAATTATTGGTGATGACAATGAGGAATTTATAGATAATAGTTTTATGAGAACTTGTCATAAATTTAAATTTTACAAAAGTAATCGAACTGATGTATATAATCCATATGATCGAAAATTAGAAAAATATAATTATGAAAGAGACGAAACATTTTGTTCATTAACAGATCTTGAGTATATTGTACATGATGATCAGAAAAAAAAAAATAAATATTTTATTTATAAAAATAAAAGTTTAAATGATTACAACTCGTGTTTTGAAAAAAAAAGTAAAAGACTTAGATGGTTTCGTTTAAAAAGAAAAAAAAAAAAAGATTATGTTAATGAAGAAAGTATTAAATATCTTAATGAATCTGTCTGCTCAAATTATAAAGAGGCAAATAAAAGTAATCAAATTCAAAACTCATCTGAATATACTAGAGGTTGTGTTTTAAATACTGTTAATGGAATTAAAGGATTTTCAACATGGATTATT**TGAaaaaaaaaaaaaaaaaaaaaaaaaaaaaatatgtgcatatgtaattgtaccttatagctagccataagtatatatatatatatatatatatatatatatatatatttatatatttatatatgtatgcattttagctagctacatatatttttaccttatcatgtaaggtaaataaataaatttatatttaacatatttatttgataaagaatgataacaaaatatacaaggaaaatataaagaaaatgatgaggggaaaaaaaaaaaataacagaaaaataaaaactgagagagaaaaaaaaaaaaaacagaataagtattagataatattgaaacaaaaaatgagttattgaaatattattccaaattttatatattaaaaaaattgaagaatatattttttttctattaagttgctttaataaacaaataaaaaatcaaaaaaaaaaaaaaaaaaaagaagaaagagatgtagaatattatcacgatttattatggatattagaaaaattatatatacacacttatatttatatatatatatatatatataatatatttaccccgctcccctatatagacattgataatatttttgaatggaatttttttcatttgagaattatataagaaacgttaataattttttaatgtttatatatttataaattttttttttttattttctttctatacatatgtatatatatatatatatatatataattttcctttatttattattataataaatgtaaggatagattgaaaaatgaaaataataaaaagaaaaaaaaggtatacaaaaaaaaaaaaaaaaataagagtaaaaattaaaaaataatatatatttatgttgaaatcaagcaactaatatattatatatatatttttttttttaatattttatag

**Original Locus (OL) amplicon size: 1094bp**

### > PF3D7_1217500-Halo-SW Halo Neo-R

tattttcttctcaataccaaaaatttttataaataaatactaataaataaaatattatatttactttattatataaaaaatatatatatatatatatatatatatatattaaaaacgtacaaaataagaattaattttttttattttatataaatttaaatattgatgatatcatattatttatattatattattataaatataatgatttaaaaaagttaaaaaaaaaaaagtatatatatatatatatatatatatacatataaaggttccaatttttatatgaagtatgagtttatatataaatataaagaaattttttttttgcatgtttcttgcatgtttctctatgcataaaaatttatttatatatatatatatatactatacatataaattgaaaaagaatatacataaaattaatatataatatatttataatattatataacttttataaaataatgaaatatgcaaaaagaaaaaaaaaaaaaaaaaaaaaaaaagaaaaatacaaatatattaaaatttttataataatatataaaatataactatattatatatatatataattgtattgaataatatatttattttttaacataaattatggaatctttttttttttttacatattttataactttttctatatttttctaaaaaaaataatatattttattttttgctttttattctatatatatatatatatatatatatatataatatataacaatttttagatttatttatatgttattatatattatatataaatttttatttaaagaaaaaaaaattgaaactttttttttctttttcaaaactgttattatatatatatatatatatatataagaagtttatatatatataataataattataataatacagaatacttaattttttttcataatatgaaaaaaggatttttttaaacaataaatatatatatatatattttgaaagtctggagtattttataagaggtaccattgaaaatattatatatatatatatatatatatatatatatatatatgtatttatttatttatttattaattttacacttcacattagttatatatatagtatatatttatttatatgaacttttttttttatgaaagaaaaaatatttgtttatattattttataaataaattttatatatgtaatttattgtataattgctttctttttttttcgtttcttttatttttttttaattaaatataaaagaaagctattataaaaagaaaaaatataggagtatattaatataatgtacaaacgtttattttgtatgtaattttttttatttacgttgcctatattattatccatattttctatactataaaaaagaaaaaatatataaagagtctatttattttgaatagaaagaaaaaataaaaaaaatagtctttttatagtaaaattagagtgtatatatatatatatatatatataatatttaattatatacacaaaaatatatatgtgaagttttaccttattattcttatatattatatcttattattatattttaatttttaatcctatcacattatatatatatatatatataatatatatatattttgtcttaataacgcaaaaaaaaaaaaaaaataaataatatattttatatataATG**ACTATTACACTGGAAAGTTTAAGTGAGGAAAAGTTACAGAATTATAACAATAAAAGAAAATATAATTGTCGTATGACATTTAAAAGATTTACAAAAAAGAGATTTAAAAATAAAGATAAAAATAAAGAAGGAAAAATTAATATTAGAACGTCCTTAATTAAAAATGAAATTATTGGTGATGACAATGAGGAATTTATAGATAATAGTTTTATGAGAACTTGTCATAAATTTAAATTTTACAAAAGTAATCGAACTGATGTATATAATCCATATGATCGAAAATTAGAAAAATATAATTATGAAAGAGACGAAACATTTTGTTCATTAACAGATCTTGAGTATATTGTACATGATGATCAGAAAAAAAAAAATAAATATTTTATTTATAAAAATAAAAGTTTAAATGATTACAACTCGTGTTTTGAAAAAAAAAGTAAAAGACTTAGATGGTTTCGTTTAAAAAGAAAAAAAAAAAAAGATTATGTTAATGAAGAAAGTATTAAATATCTTAATGAATCTGTCTGCTCAAATTATAAAGAGGCAAATAAAAGTAATCAAATTCAAAACTCATCTGAATATACTAGAGGTTGTGTTTTAAATACTGTTAATGGAATTAAAGGATTTTCAACATGGATTATT**CCTAGGTCAGGATTGAGATCAAGATCTGCTGCTGCTGGTGCTGGTGGTGCTGCTAGAGCTGCTCTGCAGAGAGGAGTACAAGTTGAAACAATATCACCAGGAGATGGTCGTACATTTCCAAAAAGAGGTCAAACTTGTGTTGTACATTATACTGGAATGCTTGAAGATGGAAAGAAATTTGATTCATCTCGTGATAGAAATAAACCATTTAAATTTATGCTAGGTAAACAAGAAGTAATACGAGGTTGGGAAGAAGGAGTTGCTCAAATGAGTGTAGGTCAAAGAGCAAAACTTACTATATCTCCAGATTATGCTTATGGTGCAACTGGACATCCAGGTATAATTCCACCTCATGCAACTCTTGTATTTGATGTGGAGCTTCTAAAACTAGAAACTAGAGGTGTTCAGGTTGAAACAATTTCACCTGGAGATGGCAGAACCTTTCCTAAAAGAGGACAGACTTGCGTAGTTCATTATACAGGCATGCTAGAGGATGGTAAGAAATTTGATTCTAGTCGAGATAGAAATAAGCCATTCAAGTTTATGCTAGGTAAACAGGAAGTAATAAGAGGTTGGGAAGAGGGTGTAGCACAGATGTCAGTTGGACAAAGAGCAAAGTTAACAATATCACCAGATTATGCATACGGTGCAACAGGCCATCCTGGCATCATCCCTCCACATGCAACTTTAGTATTCGACGTTGAATTGTTAAAGTTAGAGACAACGCGTGCTAGAGGTGCTGCTGCTGGTGCTGGAGGTGCAGGTAGACGTACGATGGCAGAAATTGGTACGGGTTTTCCATTTGATCCTCATTATGTGGAGGTGCTGGGGGAAAGGATGCATTATGTTGACGTAGGACCAAGAGATGGTACTCCAGTGTTATTTTTGCATGGAAACCCAACCTCGAGTTATGTATGGAGAAATATAATTCCACATGTAGCACCAACACATAGATGTATAGCTCCTGATTTAATTGGTATGGGAAAAAGTGATAAACCTGACTTAGGATATTTTTTTGATGATCATGTCCGTTTTATGGATGCTTTCATTGAAGCTCTTGGCCTTGAAGAAGTAGTATTAGTTATACATGATTGGGGATCCGCCTTAGGATTTCATTGGGCCAAGAGGAATCCTGAAAGAGTAAAAGGAATAGCATTCATGGAATTCATACGACCAATCCCCACATGGGATGAATGGCCAGAATTTGCACGCGAAACATTTCAAGCTTTTAGAACTACAGATGTTGGTAGAAAATTAATAATAGATCAAAATGTATTTATAGAAGGAACTTTACCTATGGGTGTTGTAAGGCCGTTAACAGAAGTTGAAATGGACCACTACCGTGAACCTTTTTTAAATCCAGTAGATAGAGAGCCCTTATGGAGATTTCCTAATGAATTACCTATTGCAGGTGAACCCGCGAATATTGTTGCTTTAGTAGAAGAATATATGGATTGGTTACATCAGTCTCCTGTTCCTAAACTTCTATTTTGGGGTACACCTGGAGTTCTAATACCACCAGCTGAAGCAGCAAGATTAGCAAAATCATTACCAAATTGTAAAGCTGTTGATATAGGTCCTGGGTTGAATTTATTACAAGAAGATAATCCAGATTTGATTGGATCTGAGATAGCTAGATGGCTAAGTACATTAGAAATTTCAGGTACCGGTGCCAGGGGAGCAGCCGCAGGAGCAGGGGGGGCAGGAAGGCGTGGTGTTCAGGTCGAGACTATTAGCCCTGGAGATGGACGCACGTTTCCTAAGCGTGGACAGACATGCGTAGTTCACTACACAGGTATGTTGGAGGACGGTAAAAAGTTCGACAGCTCACGCGACCGCAATAAACCTTTCAAGTTTATGCTTGGCAAGCAGGAGGTTATTCGTGGATGGGAGGAGGGTGTAGCACAGATGTCTGTTGGACAGCGTGCTAAGTTGACAATTTCACCTGACTATGCTTATGGCGCTACGGGCCATCCCGGGATCATTCCGCCACATGCGACTCTGGTATTCGACGTTGAATTATTAAAGTTAGAGACAGCTAGAGGGGCCGCTGCAGGTGCTGGTGGAGCTGGAAGACGTGGAGTACAAGTAGAGACTATCTCTCCAGGTGACGGTCGCACTTTCCCAAAGCGTGGCCAAACCTGTGTTGTACATTACACTGGTATGCTGGAGGATGGGAAAAAGTTCGATTCCAGTCGCGACCGTAACAAACCGTTCAAATTCATGTTGGGAAAGCAGGAAGTGATCCGCGGGTGGGAGGAAGGCGTGGCGCAAATGAGCGTCGGTCAGCGGGCTAAATTGACCATTTCCCCTGACTACGCGTATGGGGCTACTGGGCACCCAGGGATTATTCCGCCTCACGCTACACTTGTGTTTGATGTCGAACTTTTGAAACTGGAAACTGTCGACGGAGAAGGAAGAGGAAGTTTATTAACATGTGGAGATGTAGAAGAAAATCCAGGACCAATGATTGAACAAGATGGATTGCACGCAGGTTCTCCGGCCGCTTGGGTGGAGAGGCTATTCGGCTATGACTGGGCACAACAGACAATCGGCTGCTCTGATGCCGCCGTGTTCCGGCTGTCAGCGCAGGGGCGCCCGGTTCTTTTTGTCAAGACCGACCTGTCCGGTGCCCTGAATGAACTGCAGGACGAGGCAGCGCGGCTATCGTGGCTGGCCACGACGGGCGTTCCTTGCGCAGCTGTGCTCGACGTTGTCACTGAAGCGGGAAGGGACTGGCTGCTATTGGGCGAAGTGCCGGGGCAGGATCTCCTGTCATCTCACCTTGCTCCTGCCGAGAAAGTATCCATCATGGCTGATGCAATGCGGCGGCTGCATACGCTTGATCCGGCTACCTGCCCATTCGACCACCAAGCGAAACATCGCATCGAGCGAGCACGTACTCGGATGGAAGCCGGTCTTGTCGATCAGGATGATCTGGACGAAGAGCATCAGGGGCTCGCGCCAGCCGAACTGTTCGCCAGGCTCAAGGCGCGCATGCCCGACGGCGAGGATCTCGTCGTGACCCATGGCGATGCCTGCTTGCCGAATATCATGGTGGAAAATGGCCGCTTTTCTGGATTCATCGACTGTGGCCGGCTGGGTGTGGCGGACCGCTATCAGGACATAGCGTTGGCTACCCGTGATATTGCTGAAGAGCTTGGCGGCGAATGGGCTGACCGCTTCCTCGTGCTTTACGGTATCGCCGCTCCCGATTCGCAGCGCATCGCCTTCTATCGCCTTCTTGACGAGTTCTTCTAActcgagggatatggcagcttaatgttcgtttttcttatttatatatttataccaattgattgtatttataactgtaaaaatgtgtatgttgtgtgcatatttttttttgtgcatgcacatgcatgtaaatagctaaaattatgaacattttattttttgttcagaaaaaaaaaactttacacacataaaatggctagtatgaatagccatattttatataaattaaatcctatgaatttatgaccatattaaaaatttagatatttatggaacataatatgtttgaaacaataagacaaaattattattattattattatttttactgttataattatgtgtctccttcaatgattcataaatagttggacttgatttttaaaatgtttataatatgattagcatagttaaataaaaaaagttgaaaaattaaaaaaaaacatataaacacaaatgatggtttttccttcaatttcgatatcaatttatagaaacaaaatatatacttgtataattttatttttttatataaatcattacatatataattatacaatattttttctaagagataattatatattaatatatataaaaaaaggtgttttttttttttttttttatttttatttttattttatggtaatattttattttccttattttataaattatattagtttatatgtgattaattttatatattatcaatttatatatttttaaatgcttacttaattatctttttttttttttttttttttttttcccctctttttatattaatttatttttgaaaaaattgatatatatatatatatataatatatatatatacatgtagtagtattaaacaatgtataatatatataaataatatatttatatatttcatttcaattttaattttttttggttttttttttttttctttttgtcatatttaaaaaaaattatattcatataagttatgcattttttataaacattattcaatatatgtataatataatatatatatatatattaatgtattattccaatgtgcatgataaaagaaaaaaataatatttataaaaaaaaagaaaaataaaacaaaaaaagaaaaaaaaaaaaaaaaaaaaaaaaatacaaaaataaataatataatttataattatatattcttgtcacaataaaaatatatatatatatatatatatttataatatgtatattttaaactagaaaaggaataactaatattttatttattatcattcaagatttatattttataataataaatacctaatagaaatatatcaggatccatgcatggttcgctaaactgcatcgtcgctgtgtcccagaacatgggcatcggcaagaacggggactacccctggccaccgctcaggaacgaatttagatatttccagagaatgaccacaacctcttcagtagaaggtaaacagaatctggtgattatgggtaagaagacctggttctccattcctgagaagaatcgacctttaaagggtagaattaatttagttctcagcagagaactcaaggaacctccacaaggagctcattttctttccagaagtctagatgatgccttaaaacttactgaacaaccagaattagcaaataaagtagacatggtctggatagttggtggcagttctgtttataaggaagccatgaatcacccaggccatcttaaactatttgtgacaaggatcatgcaagactttgaaagtgacacgttttttccagaaattgatttggagaaatataaacttctgccagaatacccaggtgttctctctgatgtccaggaggagaaaggcattaagtacaaatttgaagtatatgagaagaatgattaagcttatttaataatagattaaaaatattataaaaataaaaacataaacacagaaattacaaaaaaaatacatatgaattttttttttgtaatcttccttataaatatagaataatgaatcatataaaacatatcattattcatttatttacatttaaaattattgtttcagtatctttaatttattatgtatatataaaaataacttacaattttattaataaacaatatatgtttattaattcatgttttgtaatttatgggatagcgattttttttactgtctgtatttttcttttttaattatgttttaattgtattttatttttattattgttctttttatagtattattttaaaacaaaatgtattttctaagaacttataataataataaatataaattttaataaaaattatatttatcttttacaatatgaacataaagtacaacattaatatatagcttttaatatttttattcctaatcatgtaaatcttaaatttttctttttaaacatatgttaaatatttatttctcattatatataagaacatatttattaaatctagaattctatagtgagtcgtattacaattcactggccgtcgttttacaacgtcgtgactgggaaaaccctggcgttacccaacttaatcgccttgcagcacatccccctttcgccagctggcgtaatagcgaagaggcccgcaccgatcgcccttcccaacagttgcgcagcctgaatggcgaatggcgcctgatgcggtattttctccttacgcatctgtgcggtatttcacaccgcatatggtgcactctcagtacaatctgctctgatgccgcatagttaagccagccccgacacccgccaacacccgctgacgcgccctgacgggcttgtctgctcccggcatccgcttacagacaagctgtgaccgtctccgggagctgcatgtgtcagaggttttcaccgtcatcaccgaaacgcgcgagacgaaagggcctcgtgatacgcctatttttataggttaatgtcatgataataatggtttcttagacgtcaggtggcacttttcggggaaatgtgcgcggaacccctatttgtttatttttctaaatacattcaaatatgtatccgctcatgagacaataaccctgataaatgcttcaataatattgaaaaaggaagagtatgagtattcaacatttccgtgtcgcccttattcccttttttgcggcattttgccttcctgtttttgctcacccagaaacgctggtgaaagtaaaagatgctgaagatcagttgggtgcacgagtgggttacatcgaactggatctcaacagcggtaagatccttgagagttttcgccccgaagaacgttttccaatgatgagcacttttaaagttctgctatgtggcgcggtattatcccgtattgacgccgggcaagagcaactcggtcgccgcatacactattctcagaatgacttggttgagtactcaccagtcacagaaaagcatcttacggatggcatgacagtaagagaattatgcagtgctgccataaccatgagtgataacactgcggccaacttacttctgacaacgatcggaggaccgaaggagctaaccgcttttttgcacaacatgggggatcatgtaactcgccttgatcgttgggaaccggagctgaatgaagccataccaaacgacgagcgtgacaccacgatgcctgtagcaatgccaacaacgttgcgcaaactattaactggcgaactacttactctagcttcccggcaacaattaatagactggatggaggcggataaagttgcaggaccacttctgcgctcggcccttccggctggctggtttattgctgataaatctggagccggtgagcgtgggtctcgcggtatcattgcagcactggggccagatggtaagccctcccgtatcgtagttatctacacgacggggagtcaggcaactatggatgaacgaaatagacagatcgctgagataggtgcctcactgattaagcattggtaactgtcagaccaagtttactcatatatactttagattgatttaaaacttcatttttaatttaaaaggatctaggtgaagatcctttttgataatctcatgaccaaaatcccttaacgtgagttttcgttccactgagcgtcagaccccgtagaaaagatcaaaggatcttcttgagatcctttttttctgcgcgtaatctgctgcttgcaaacaaaaaaaccaccgctaccagcggtggtttgtttgccggatcaagagctaccaactctttttccgaaggtaactggcttcagcagagcgcagataccaaatactgtccttctagtgtagccgtagttaggccaccacttcaagaactctgtagcaccgcctacatacctcgctctgctaatcctgttaccagtggctgctgccagtggcgataagtcgtgtcttaccgggttggactcaagacgatagttaccggataaggcgcagcggtcgggctgaacggggggttcgtgcacacagcccagcttggagcgaacgacctacaccgaactgagatacctacagcgtgagctatgagaaagcgccacgcttcccgaagggagaaaggcggacaggtatccggtaagcggcagggtcggaacaggagagcgcacgagggagcttccagggggaaacgcctggtatctttatagtcctgtcgggtttcgccacctctgacttgagcgtcgatttttgtgatgctcgtcaggggggcggagcctatcgaaaaacgccagcaacgcggcctttttacggttcctggccttttgctggccttttgctcacatgttctttcctgcgttatcccctgattctgtggataaccgtattaccgcctttgagtgagctgataccgctcgccgcagccgaacgaccgagcgcagcgagtcagtgagcgaggaagcggaagagcgcccaatacgcaaaccgcctctccccgcgcgttggccgattcattaatgcagctggcacgacaggtttcccgactggaaagcgggcagtgagcgcaacgcaattaatgtgagttagctcactcattaggcaccccaggctttacactttatgcttccggctcgtatgttgtgtggaattgtgagcggataacaatttcacacaggaaacagctatgaccatgattacgccaagctatttaggtgacactatagaagcggccgctagACTATTACACTGGAAAGTTTAAGTGAGGAAAAGTTACAGAATTATAACAATAAAAGAAAATATAATTGTCGTATGACATTTAAAAGATTTACAAAAAAGAGATTTAAAAATAAAGATAAAAATAAAGAAGGAAAAATTAATATTAGAACGTCCTTAATTAAAAATGAAATTATTGGTGATGACAATGAGGAATTTATAGATAATAGTTTTATGAGAACTTGTCATAAATTTAAATTTTACAAAAGTAATCGAACTGATGTATATAATCCATATGATCGAAAATTAGAAAAATATAATTATGAAAGAGACGAAACATTTTGTTCATTAACAGATCTTGAGTATATTGTACATGATGATCAGAAAAAAAAAAATAAATATTTTATTTATAAAAATAAAAGTTTAAATGATTACAACTCGTGTTTTGAAAAAAAAAGTAAAAGACTTAGATGGTTTCGTTTAAAAAGAAAAAAAAAAAAAGATTATGTTAATGAAGAAAGTATTAAATATCTTAATGAATCTGTCTGCTCAAATTATAAAGAGGCAAATAAAAGTAATCAAATTCAAAACTCATCTGAATATACTAGAGGTTGTGTTTTAAATACTGTTAATGGAATTAAAGGATTTTCAACATGGATTATTtgaaaaaaaaaaaaaaaaaaaaaaaaaaaaaatatgtgcatatgtaattgtaccttatagctagccataagtatatatatatatatatatatatatatatatatatatttatatatttatatatgtatgcattttagctagctacatatatttttaccttatcatgtaaggtaaataaataaatttatatttaacatatttatttgataaagaatgataacaaaatatacaaggaaaatataaagaaaatgatgaggggaaaaaaaaaaaataacagaaaaataaaaactgagagagaaaaaaaaaaaaaacagaataagtattagataatattgaaacaaaaaatgagttattgaaatattattccaaattttatatattaaaaaaattgaagaatatattttttttctattaagttgctttaataaacaaataaaaaatcaaaaaaaaaaaaaaaaaaaagaagaaagagatgtagaatattatcacgatttattatggatattagaaaaattatatatacacacttatatttatatatatatatatatatataatatatttaccccgctcccctatatagacattgataatatttttgaatggaatttttttcatttgagaattatataagaaacgttaataattttttaatgtttatatatttataaattttttttttttattttctttctatacatatgtatatatatatatatatatatataattttcctttatttattattataataaatgtaaggatagattgaaaaatgaaaataataaaaagaaaaaaaaggtatacaaaaaaaaaaaaaaaaataagagtaaaaattaaaaaataatatatatttatgttgaaatcaagcaactaatatattatatatatatttttttttttaatattttatag

**5’ amplicon size: 1165bps**

**3’ amplicon size: 793bps**

## > PF3D7_0520100

tttccatcctaatattataatattatatatatatatttaatattaagtaccctttttaattttgttgtttttatgatatatctttttattgatatattattattctttttattatattaattcaatagttaaaaaaattgataatttttctagtaaatattatattctaatatattaagagtaataaaggcatttattaaaataacatacgtacataagtatataaaaaaaaaaaaaaaaatataaatatatatataaatatatatatagatataatacatatgatacattaattttattctctacagttcgtatttttttatatatatttttctaatttttagttatattttttacttttaaaaagtaaatattaaatgatagaaaataaaagaaaaaaaaggaaaaagaatatatagtaatattatacatatgttatatacatatatatatgtatatatgagtattataaaatatatatatttactgtttttttttttgttgcttgttcaaaaatgagtacatacaaatagttgtatatttaaaatatatttttctataacattcttttgtgagtaaaaataattatatgatataatatttcaggggaaacatataccattataatatatgtatatgtatatatatatatatatataatacttaaatatatatatatatatatatatattattacttttttttttttttttttctattATGAGTTTATTAACCAATTATAGAGGTAAGAATAATAATGAAACCATGATAACAGAAAGGAGTGTTAATAATGTGAATGAATCTGATAAAGTTAATAATATAAAAGAAATGAATAATGTGAATGATATGAATAATATTAATAATGCAGAGAAGATAATTGAATATAATAGTATAAATGATTTAAAATATTGTAAAAGTATTATAGACATTGTAAATTATATTATAACTATTGGTATATGGGAAATGTATACATACGAATGGGTATTTAATAGAAAGACGTTTTTATATTATCATACATTATTAGAGAAATATTATTTTTATGACAAAAATAATAATTTGACTTTATTTAATGAGAAGTCTTTTTTTTTATATGTATGCACATTTGAAAATAATGAATTTGTATCTGATTTTTTTTTTCAAAGGAATAAAAAAAAAAAAGAATTAAAATTAACTAGTTATACAAAAACTATGCAAGGTAGAAGAAAAAAGCAAGAAGATCGTTACCTTGTTATAACTGATCTTACTAAATATATTGATTCAAATGATTATAAAACTTTATATTTTTATAAAAAAAATCCATTATACTTTTATTCAATATTTGATGGCCATAGAGGAATAAAAGCTTGTGAATATTGTATGAGTCACATTATAAAAAATATTATTTATTATTTTTATAATCAAAATATGGAAGATGATCAATCTTCTACTACTATTAATAAATTAAATTCTAATGATATTAATATATCACCTGATAAAAGAAGAAAATTTGATAATGTAAAGAATGAGGACATACCATTAGGTGATTCAACAATTAATATAAATAATATAAATGATATGTGTAATATAAATAATAATATAAACAAATGTAATGAAAATTTTAATGATTTACATATAGAAGGAGATAATACAAATGATAATAATTATTGCATGAACAATAAAACAGAAGAAGAAAGAAAAGAAAATTTATTTATTAATACAACATCTAAAAATGATTATTTCTTAAAAAATATTTCTTCTGTTAATCATATATCAAATGACAATATTTTAAATAATATTCATACCAATAAAAAAGATATTGAAAATAACACACCAATAAAAAATAAAGATAAAAGATTAATTGATAGTTCGTATTTAAATACTATTAGTGATGATGTAGAATATAGGAAGAAAAGAAAAGTTGAGGAAATGCATATTATTAAGCAAAATATAAATAATAAGAATGAATATATTAATTTGAATATATCTGAAGAATGTTCCAATAATATAGACCAAACTCATAATAATAAGAAGAAGAAAATATCTGATAATTCCAATTCTTATGAAAAATTAAAATTTCTAGACGAATTAACAGATGAACAAATAATGGAATATATAAAACTAGCCTTTTTAAAAACTGATGAACAATTTTTAAAAGTATCAAAATTTCCTAACCATGGCTGTACTATTATTTCATTAATTATATTTAGAAATAAAATGTTTGTAGCTAACTTGGGAGATTGTAGAGCCATAGGGGTTGTTAATATAAGTAACACTTTG*gtatatataaaaagaagaaaaatataaatcaataaataattaaataaataaatacatatatatatatatatatatatattattcatatttattttattttatatataattttttcatattgctctttctttatgttttgtattattttcctgttaaaatatttcatag*AAA**ACGGAAGTGTTGTCAAATGATCATAAGCCCAATGATCCTAAAGAGAAGGAGAGGATCAAAAAAATGGGAGGAGATGTCATATGCTTACAAAATGTTTATCGTGTTAAGGCAAATGCTAGAAAG*gtaaagatataaaatagcaacaaaaaaaattattacaaagtaaatatttgtatatttatgtatatgtatgaaaattcattattattttcatgtag*AATAATAAAGACAAGCCTAGTTTACTTGAGAGGCTTTCCATGAAGGAGGAAGTATATCTTGCAG*gtaaatattaataataacatatatgtatatatatataaatatatgtatatatgtatatattaatttatattgtattattattttctccttttttttttttag*TTTCAAGAGCTATTGGTGACAAAGACTTTAAATTTAATAATGTCATTTCAGCTACTCCTGATGTTATATGTAAAGAAATATATAGTGAAAAATGTAATGAGAAGAAAGAAGAAAAGGAAATTACAAAACGTACTAATGTTATTGAAAAAAGCGATATTGATGAAAATTATTTTAAGGAGACTGATAATTTATATTATTCAGCTCACGAAGTGAATTATCATTATGTTGTTATGGCATGTGATGGAGTATGGGATACCATGTCTAATAAG*gtacatacgaatctgttaatgagtacaattatgtcacatatatatatatatatatatatttttatttatttatgtttatttttttttccatttcattttatattttttatag*GATATCGTTAAAATATTACAAACATATAATAACGATCCGGATAAAGCATGTAGCGAAATAATAAAAACGGCTTATGCCTATGGTAGTCAG*gtacattgaaatgtaaataaatcacatatatatatatatgtaatatatatgtatatttattcatttatatgtctataattaatatgaacttatccatataatttgatttgttttaatttatttctgtag*GATAATTTAACCGCTATGTTGCTCAAATTTTAT**TGAaatcatattcttattttattttttttattttttatattcatccttattttttaaaaatcactacgattaatatatatatattttacattatgttttaccctctgtttatatgtctaatttttttggtgatatttttctcacacatttaccagagatgtgattcagttatatatatatatatatataaatttatatatatgatgtttgtgattcatttattataaggaataaataaatatttcctaatcattgatatatttattcatatttttatttaaccaaagaaatattctaatgttattttattttatcctatttttttcattattaatttttatttatttatttatttatttattaccatttattgttctgattc

**Original Locus (OL) amplicon size: 1327bp**

### > PF3D7_0520100-Halo-SW Halo Neo-R

tttccatcctaatattataatattatatatatatatttaatattaagtaccctttttaattttgttgtttttatgatatatctttttattgatatattattattctttttattatattaattcaatagttaaaaaaattgataatttttctagtaaatattatattctaatatattaagagtaataaaggcatttattaaaataacatacgtacataagtatataaaaaaaaaaaaaaaaatataaatatatatataaatatatatatagatataatacatatgatacattaattttattctctacagttcgtatttttttatatatatttttctaatttttagttatattttttacttttaaaaagtaaatattaaatgatagaaaataaaagaaaaaaaaggaaaaagaatatatagtaatattatacatatgttatatacatatatatatgtatatatgagtattataaaatatatatatttactgtttttttttttgttgcttgttcaaaaatgagtacatacaaatagttgtatatttaaaatatatttttctataacattcttttgtgagtaaaaataattatatgatataatatttcaggggaaacatataccattataatatatgtatatgtatatatatatatatatataatacttaaatatatatatatatatatatatattattacttttttttttttttttttctattATGAGTTTATTAACCAATTATAGAGGTAAGAATAATAATGAAACCATGATAACAGAAAGGAGTGTTAATAATGTGAATGAATCTGATAAAGTTAATAATATAAAAGAAATGAATAATGTGAATGATATGAATAATATTAATAATGCAGAGAAGATAATTGAATATAATAGTATAAATGATTTAAAATATTGTAAAAGTATTATAGACATTGTAAATTATATTATAACTATTGGTATATGGGAAATGTATACATACGAATGGGTATTTAATAGAAAGACGTTTTTATATTATCATACATTATTAGAGAAATATTATTTTTATGACAAAAATAATAATTTGACTTTATTTAATGAGAAGTCTTTTTTTTTATATGTATGCACATTTGAAAATAATGAATTTGTATCTGATTTTTTTTTTCAAAGGAATAAAAAAAAAAAAGAATTAAAATTAACTAGTTATACAAAAACTATGCAAGGTAGAAGAAAAAAGCAAGAAGATCGTTACCTTGTTATAACTGATCTTACTAAATATATTGATTCAAATGATTATAAAACTTTATATTTTTATAAAAAAAATCCATTATACTTTTATTCAATATTTGATGGCCATAGAGGAATAAAAGCTTGTGAATATTGTATGAGTCACATTATAAAAAATATTATTTATTATTTTTATAATCAAAATATGGAAGATGATCAATCTTCTACTACTATTAATAAATTAAATTCTAATGATATTAATATATCACCTGATAAAAGAAGAAAATTTGATAATGTAAAGAATGAGGACATACCATTAGGTGATTCAACAATTAATATAAATAATATAAATGATATGTGTAATATAAATAATAATATAAACAAATGTAATGAAAATTTTAATGATTTACATATAGAAGGAGATAATACAAATGATAATAATTATTGCATGAACAATAAAACAGAAGAAGAAAGAAAAGAAAATTTATTTATTAATACAACATCTAAAAATGATTATTTCTTAAAAAATATTTCTTCTGTTAATCATATATCAAATGACAATATTTTAAATAATATTCATACCAATAAAAAAGATATTGAAAATAACACACCAATAAAAAATAAAGATAAAAGATTAATTGATAGTTCGTATTTAAATACTATTAGTGATGATGTAGAATATAGGAAGAAAAGAAAAGTTGAGGAAATGCATATTATTAAGCAAAATATAAATAATAAGAATGAATATATTAATTTGAATATATCTGAAGAATGTTCCAATAATATAGACCAAACTCATAATAATAAGAAGAAGAAAATATCTGATAATTCCAATTCTTATGAAAAATTAAAATTTCTAGACGAATTAACAGATGAACAAATAATGGAATATATAAAACTAGCCTTTTTAAAAACTGATGAACAATTTTTAAAAGTATCAAAATTTCCTAACCATGGCTGTACTATTATTTCATTAATTATATTTAGAAATAAAATGTTTGTAGCTAACTTGGGAGATTGTAGAGCCATAGGGGTTGTTAATATAAGTAACACTTTG*gtatatataaaaagaagaaaaatataaatcaataaataattaaataaataaatacatatatatatatatatatatatattattcatatttattttattttatatataattttttcatattgctctttctttatgttttgtattattttcctgttaaaatatttcatag*AAA**ACGGAAGTGTTGTCAAATGATCATAAGCCCAATGATCCTAAAGAGAAGGAGAGGATCAAAAAAATGGGAGGAGATGTCATATGCTTACAAAATGTTTATCGTGTTAAGGCAAATGCTAGAAAG*gtaaagatataaaatagcaacaaaaaaaattattacaaagtaaatatttgtatatttatgtatatgtatgaaaattcattattattttcatgtag*AATAATAAAGACAAGCCTAGTTTACTTGAGAGGCTTTCCATGAAGGAGGAAGTATATCTTGCAG*gtaaatattaataataacatatatgtatatatatataaatatatgtatatatgtatatattaatttatattgtattattattttctccttttttttttttag*TTTCAAGAGCTATTGGTGACAAAGACTTTAAATTTAATAATGTCATTTCAGCTACTCCTGATGTTATATGTAAAGAAATATATAGTGAAAAATGTAATGAGAAGAAAGAAGAAAAGGAAATTACAAAACGTACTAATGTTATTGAAAAAAGCGATATTGATGAAAATTATTTTAAGGAGACTGATAATTTATATTATTCAGCTCACGAAGTGAATTATCATTATGTTGTTATGGCATGTGATGGAGTATGGGATACCATGTCTAATAAG*gtacatacgaatctgttaatgagtacaattatgtcacatatatatatatatatatatatttttatttatttatgtttatttttttttccatttcattttatattttttatag*GATATCGTTAAAATATTACAAACATATAATAACGATCCGGATAAAGCATGTAGCGAAATAATAAAAACGGCTTATGCCTATGGTAGTCAG*gtacattgaaatgtaaataaatcacatatatatatatatgtaatatatatgtatatttattcatttatatgtctataattaatatgaacttatccatataatttgatttgttttaatttatttctgtag*GATAATTTAACCGCTATGTTGCTCAAATTTTAT**CCTAGGTCAGGATTGAGATCAAGATCTGCTGCTGCTGGTGCTGGTGGTGCTGCTAGAGCTGCTCTGCAGAGAGGAGTACAAGTTGAAACAATATCACCAGGAGATGGTCGTACATTTCCAAAAAGAGGTCAAACTTGTGTTGTACATTATACTGGAATGCTTGAAGATGGAAAGAAATTTGATTCATCTCGTGATAGAAATAAACCATTTAAATTTATGCTAGGTAAACAAGAAGTAATACGAGGTTGGGAAGAAGGAGTTGCTCAAATGAGTGTAGGTCAAAGAGCAAAACTTACTATATCTCCAGATTATGCTTATGGTGCAACTGGACATCCAGGTATAATTCCACCTCATGCAACTCTTGTATTTGATGTGGAGCTTCTAAAACTAGAAACTAGAGGTGTTCAGGTTGAAACAATTTCACCTGGAGATGGCAGAACCTTTCCTAAAAGAGGACAGACTTGCGTAGTTCATTATACAGGCATGCTAGAGGATGGTAAGAAATTTGATTCTAGTCGAGATAGAAATAAGCCATTCAAGTTTATGCTAGGTAAACAGGAAGTAATAAGAGGTTGGGAAGAGGGTGTAGCACAGATGTCAGTTGGACAAAGAGCAAAGTTAACAATATCACCAGATTATGCATACGGTGCAACAGGCCATCCTGGCATCATCCCTCCACATGCAACTTTAGTATTCGACGTTGAATTGTTAAAGTTAGAGACAACGCGTGCTAGAGGTGCTGCTGCTGGTGCTGGAGGTGCAGGTAGACGTACGATGGCAGAAATTGGTACGGGTTTTCCATTTGATCCTCATTATGTGGAGGTGCTGGGGGAAAGGATGCATTATGTTGACGTAGGACCAAGAGATGGTACTCCAGTGTTATTTTTGCATGGAAACCCAACCTCGAGTTATGTATGGAGAAATATAATTCCACATGTAGCACCAACACATAGATGTATAGCTCCTGATTTAATTGGTATGGGAAAAAGTGATAAACCTGACTTAGGATATTTTTTTGATGATCATGTCCGTTTTATGGATGCTTTCATTGAAGCTCTTGGCCTTGAAGAAGTAGTATTAGTTATACATGATTGGGGATCCGCCTTAGGATTTCATTGGGCCAAGAGGAATCCTGAAAGAGTAAAAGGAATAGCATTCATGGAATTCATACGACCAATCCCCACATGGGATGAATGGCCAGAATTTGCACGCGAAACATTTCAAGCTTTTAGAACTACAGATGTTGGTAGAAAATTAATAATAGATCAAAATGTATTTATAGAAGGAACTTTACCTATGGGTGTTGTAAGGCCGTTAACAGAAGTTGAAATGGACCACTACCGTGAACCTTTTTTAAATCCAGTAGATAGAGAGCCCTTATGGAGATTTCCTAATGAATTACCTATTGCAGGTGAACCCGCGAATATTGTTGCTTTAGTAGAAGAATATATGGATTGGTTACATCAGTCTCCTGTTCCTAAACTTCTATTTTGGGGTACACCTGGAGTTCTAATACCACCAGCTGAAGCAGCAAGATTAGCAAAATCATTACCAAATTGTAAAGCTGTTGATATAGGTCCTGGGTTGAATTTATTACAAGAAGATAATCCAGATTTGATTGGATCTGAGATAGCTAGATGGCTAAGTACATTAGAAATTTCAGGTACCGGTGCCAGGGGAGCAGCCGCAGGAGCAGGGGGGGCAGGAAGGCGTGGTGTTCAGGTCGAGACTATTAGCCCTGGAGATGGACGCACGTTTCCTAAGCGTGGACAGACATGCGTAGTTCACTACACAGGTATGTTGGAGGACGGTAAAAAGTTCGACAGCTCACGCGACCGCAATAAACCTTTCAAGTTTATGCTTGGCAAGCAGGAGGTTATTCGTGGATGGGAGGAGGGTGTAGCACAGATGTCTGTTGGACAGCGTGCTAAGTTGACAATTTCACCTGACTATGCTTATGGCGCTACGGGCCATCCCGGGATCATTCCGCCACATGCGACTCTGGTATTCGACGTTGAATTATTAAAGTTAGAGACAGCTAGAGGGGCCGCTGCAGGTGCTGGTGGAGCTGGAAGACGTGGAGTACAAGTAGAGACTATCTCTCCAGGTGACGGTCGCACTTTCCCAAAGCGTGGCCAAACCTGTGTTGTACATTACACTGGTATGCTGGAGGATGGGAAAAAGTTCGATTCCAGTCGCGACCGTAACAAACCGTTCAAATTCATGTTGGGAAAGCAGGAAGTGATCCGCGGGTGGGAGGAAGGCGTGGCGCAAATGAGCGTCGGTCAGCGGGCTAAATTGACCATTTCCCCTGACTACGCGTATGGGGCTACTGGGCACCCAGGGATTATTCCGCCTCACGCTACACTTGTGTTTGATGTCGAACTTTTGAAACTGGAAACTGTCGACGGAGAAGGAAGAGGAAGTTTATTAACATGTGGAGATGTAGAAGAAAATCCAGGACCAATGATTGAACAAGATGGATTGCACGCAGGTTCTCCGGCCGCTTGGGTGGAGAGGCTATTCGGCTATGACTGGGCACAACAGACAATCGGCTGCTCTGATGCCGCCGTGTTCCGGCTGTCAGCGCAGGGGCGCCCGGTTCTTTTTGTCAAGACCGACCTGTCCGGTGCCCTGAATGAACTGCAGGACGAGGCAGCGCGGCTATCGTGGCTGGCCACGACGGGCGTTCCTTGCGCAGCTGTGCTCGACGTTGTCACTGAAGCGGGAAGGGACTGGCTGCTATTGGGCGAAGTGCCGGGGCAGGATCTCCTGTCATCTCACCTTGCTCCTGCCGAGAAAGTATCCATCATGGCTGATGCAATGCGGCGGCTGCATACGCTTGATCCGGCTACCTGCCCATTCGACCACCAAGCGAAACATCGCATCGAGCGAGCACGTACTCGGATGGAAGCCGGTCTTGTCGATCAGGATGATCTGGACGAAGAGCATCAGGGGCTCGCGCCAGCCGAACTGTTCGCCAGGCTCAAGGCGCGCATGCCCGACGGCGAGGATCTCGTCGTGACCCATGGCGATGCCTGCTTGCCGAATATCATGGTGGAAAATGGCCGCTTTTCTGGATTCATCGACTGTGGCCGGCTGGGTGTGGCGGACCGCTATCAGGACATAGCGTTGGCTACCCGTGATATTGCTGAAGAGCTTGGCGGCGAATGGGCTGACCGCTTCCTCGTGCTTTACGGTATCGCCGCTCCCGATTCGCAGCGCATCGCCTTCTATCGCCTTCTTGACGAGTTCTTCTAACTCGAGGGATATGGCAGCTTAATGTTCGTTTTTCTTATTTATATATTTATACCAATTGATTGTATTTATAACTGTAAAAATGTGTATGTTGTGTGCATATTTTTTTTTGTGCATGCACATGCATGTAAATAGCTAAAATTATGAACATTTTATTTTTTGTTCAGAAAAAAAAAACTTTACACACATAAAATGGCTAGTATGAATAGCCATATTTTATATAAATTAAATCCTATGAATTTATGACCATATTAAAAATTTAGATATTTATGGAACATAATATGTTTGAAACAATAAGACAAAATTATTATTATTATTATTATTTTTACTGTTATAATTATGTGTCTCCTTCAATGATTCATAAATAGTTGGACTTGATTTTTAAAATGTTTATAATATGATTAGCATAGTTAAATAAAAAAAGTTGAAAAATTAAAAAAAAACATATAAACACAAATGATGGTTTTTCCTTCAATTTCGATATCAATTTATAGAAACAAAATATATACTTGTATAATTTTATTTTTTTATATAAATCATTACATATATAATTATACAATATTTTTTCTAAGAGATAATTATATATTAATATATATAAAAAAAGGTGTTTTTTTTTTTTTTTTTTATTTTTATTTTTATTTTATGGTAATATTTTATTTTCCTTATTTTATAAATTATATTAGTTTATATGTGATTAATTTTATATATTATCAATTTATATATTTTTAAATGCTTACTTAATTATCTTTTTTTTTTTTTTTTTTTTTTTTTCCCCTCTTTTTATATTAATTTATTTTTGAAAAAATTGATATATATATATATATATAATATATATATATACATGTAGTAGTATTAAACAATGTATAATATATATAAATAATATATTTATATATTTCATTTCAATTTTAATTTTTTTTGGTTTTTTTTTTTTTTCTTTTTGTCATATTTAAAAAAAATTATATTCATATAAGTTATGCATTTTTTATAAACATTATTCAATATATGTATAATATAATATATATATATATATTAATGTATTATTCCAATGTGCATGATAAAAGAAAAAAATAATATTTATAAAAAAAAAGAAAAATAAAACAAAAAAAGAAAAAAAAAAAAAAAAAAAAAAAAATACAAAAATAAATAATATAATTTATAATTATATATTCTTGTCACAATAAAAATATATATATATATATATATATTTATAATATGTATATTTTAAACTAGAAAAGGAATAACTAATATTTTATTTATTATCATTCAAGATTTATATTTTATAATAATAAATACCTAATAGAAATATATCAGGATCCATGCATGGTTCGCTAAACTGCATCGTCGCTGTGTCCCAGAACATGGGCATCGGCAAGAACGGGGACTACCCCTGGCCACCGCTCAGGAACGAATTTAGATATTTCCAGAGAATGACCACAACCTCTTCAGTAGAAGGTAAACAGAATCTGGTGATTATGGGTAAGAAGACCTGGTTCTCCATTCCTGAGAAGAATCGACCTTTAAAGGGTAGAATTAATTTAGTTCTCAGCAGAGAACTCAAGGAACCTCCACAAGGAGCTCATTTTCTTTCCAGAAGTCTAGATGATGCCTTAAAACTTACTGAACAACCAGAATTAGCAAATAAAGTAGACATGGTCTGGATAGTTGGTGGCAGTTCTGTTTATAAGGAAGCCATGAATCACCCAGGCCATCTTAAACTATTTGTGACAAGGATCATGCAAGACTTTGAAAGTGACACGTTTTTTCCAGAAATTGATTTGGAGAAATATAAACTTCTGCCAGAATACCCAGGTGTTCTCTCTGATGTCCAGGAGGAGAAAGGCATTAAGTACAAATTTGAAGTATATGAGAAGAATGATTAAGCTTATTTAATAATAGATTAAAAATATTATAAAAATAAAAACATAAACACAGAAATTACAAAAAAAATACATATGAATTTTTTTTTTGTAATCTTCCTTATAAATATAGAATAATGAATCATATAAAACATATCATTATTCATTTATTTACATTTAAAATTATTGTTTCAGTATCTTTAATTTATTATGTATATATAAAAATAACTTACAATTTTATTAATAAACAATATATGTTTATTAATTCATGTTTTGTAATTTATGGGATAGCGATTTTTTTTACTGTCTGTATTTTTCTTTTTTAATTATGTTTTAATTGTATTTTATTTTTATTATTGTTCTTTTTATAGTATTATTTTAAAACAAAATGTATTTTCTAAGAACTTATAATAATAATAAATATAAATTTTAATAAAAATTATATTTATCTTTTACAATATGAACATAAAGTACAACATTAATATATAGCTTTTAATATTTTTATTCCTAATCATGTAAATCTTAAATTTTTCTTTTTAAACATATGTTAAATATTTATTTCTCATTATATATAAGAACATATTTATTAAATCTAGAATTCTATAGTGAGTCGTATTACAATTCACTGGCCGTCGTTTTACAACGTCGTGACTGGGAAAACCCTGGCGTTACCCAACTTAATCGCCTTGCAGCACATCCCCCTTTCGCCAGCTGGCGTAATAGCGAAGAGGCCCGCACCGATCGCCCTTCCCAACAGTTGCGCAGCCTGAATGGCGAATGGCGCCTGATGCGGTATTTTCTCCTTACGCATCTGTGCGGTATTTCACACCGCATATGGTGCACTCTCAGTACAATCTGCTCTGATGCCGCATAGTTAAGCCAGCCCCGACACCCGCCAACACCCGCTGACGCGCCCTGACGGGCTTGTCTGCTCCCGGCATCCGCTTACAGACAAGCTGTGACCGTCTCCGGGAGCTGCATGTGTCAGAGGTTTTCACCGTCATCACCGAAACGCGCGAGACGAAAGGGCCTCGTGATACGCCTATTTTTATAGGTTAATGTCATGATAATAATGGTTTCTTAGACGTCAGGTGGCACTTTTCGGGGAAATGTGCGCGGAACCCCTATTTGTTTATTTTTCTAAATACATTCAAATATGTATCCGCTCATGAGACAATAACCCTGATAAATGCTTCAATAATATTGAAAAAGGAAGAGTATGAGTATTCAACATTTCCGTGTCGCCCTTATTCCCTTTTTTGCGGCATTTTGCCTTCCTGTTTTTGCTCACCCAGAAACGCTGGTGAAAGTAAAAGATGCTGAAGATCAGTTGGGTGCACGAGTGGGTTACATCGAACTGGATCTCAACAGCGGTAAGATCCTTGAGAGTTTTCGCCCCGAAGAACGTTTTCCAATGATGAGCACTTTTAAAGTTCTGCTATGTGGCGCGGTATTATCCCGTATTGACGCCGGGCAAGAGCAACTCGGTCGCCGCATACACTATTCTCAGAATGACTTGGTTGAGTACTCACCAGTCACAGAAAAGCATCTTACGGATGGCATGACAGTAAGAGAATTATGCAGTGCTGCCATAACCATGAGTGATAACACTGCGGCCAACTTACTTCTGACAACGATCGGAGGACCGAAGGAGCTAACCGCTTTTTTGCACAACATGGGGGATCATGTAACTCGCCTTGATCGTTGGGAACCGGAGCTGAATGAAGCCATACCAAACGACGAGCGTGACACCACGATGCCTGTAGCAATGCCAACAACGTTGCGCAAACTATTAACTGGCGAACTACTTACTCTAGCTTCCCGGCAACAATTAATAGACTGGATGGAGGCGGATAAAGTTGCAGGACCACTTCTGCGCTCGGCCCTTCCGGCTGGCTGGTTTATTGCTGATAAATCTGGAGCCGGTGAGCGTGGGTCTCGCGGTATCATTGCAGCACTGGGGCCAGATGGTAAGCCCTCCCGTATCGTAGTTATCTACACGACGGGGAGTCAGGCAACTATGGATGAACGAAATAGACAGATCGCTGAGATAGGTGCCTCACTGATTAAGCATTGGTAACTGTCAGACCAAGTTTACTCATATATACTTTAGATTGATTTAAAACTTCATTTTTAATTTAAAAGGATCTAGGTGAAGATCCTTTTTGATAATCTCATGACCAAAATCCCTTAACGTGAGTTTTCGTTCCACTGAGCGTCAGACCCCGTAGAAAAGATCAAAGGATCTTCTTGAGATCCTTTTTTTCTGCGCGTAATCTGCTGCTTGCAAACAAAAAAACCACCGCTACCAGCGGTGGTTTGTTTGCCGGATCAAGAGCTACCAACTCTTTTTCCGAAGGTAACTGGCTTCAGCAGAGCGCAGATACCAAATACTGTCCTTCTAGTGTAGCCGTAGTTAGGCCACCACTTCAAGAACTCTGTAGCACCGCCTACATACCTCGCTCTGCTAATCCTGTTACCAGTGGCTGCTGCCAGTGGCGATAAGTCGTGTCTTACCGGGTTGGACTCAAGACGATAGTTACCGGATAAGGCGCAGCGGTCGGGCTGAACGGGGGGTTCGTGCACACAGCCCAGCTTGGAGCGAACGACCTACACCGAACTGAGATACCTACAGCGTGAGCTATGAGAAAGCGCCACGCTTCCCGAAGGGAGAAAGGCGGACAGGTATCCGGTAAGCGGCAGGGTCGGAACAGGAGAGCGCACGAGGGAGCTTCCAGGGGGAAACGCCTGGTATCTTTATAGTCCTGTCGGGTTTCGCCACCTCTGACTTGAGCGTCGATTTTTGTGATGCTCGTCAGGGGGGCGGAGCCTATCGAAAAACGCCAGCAACGCGGCCTTTTTACGGTTCCTGGCCTTTTGCTGGCCTTTTGCTCACATGTTCTTTCCTGCGTTATCCCCTGATTCTGTGGATAACCGTATTACCGCCTTTGAGTGAGCTGATACCGCTCGCCGCAGCCGAACGACCGAGCGCAGCGAGTCAGTGAGCGAGGAAGCGGAAGAGCGCCCAATACGCAAACCGCCTCTCCCCGCGCGTTGGCCGATTCATTAATGCAGCTGGCACGACAGGTTTCCCGACTGGAAAGCGGGCAGTGAGCGCAACGCAATTAATGTGAGTTAGCTCACTCATTAGGCACCCCAGGCTTTACACTTTATGCTTCCGGCTCGTATGTTGTGTGGAATTGTGAGCGGATAACAATTTCACACAGGAAACAGCTATGACCATGATTACGCCAAGCTATTTAGGTGACACTATAGAAGCGGCCGCTAGACGGAAGTGTTGTCAAATGATCATAAGCCCAATGATCCTAAAGAGAAGGAGAGGATCAAAAAAATGGGAGGAGATGTCATATGCTTACAAAATGTTTATCGTGTTAAGGCAAATGCTAGAAAGGTAAAGATATAAAATAGCAACAAAAAAAATTATTACAAAGTAAATATTTGTATATTTATGTATATGTATGAAAATTCATTATTATTTTCATGTAGAATAATAAAGACAAGCCTAGTTTACTTGAGAGGCTTTCCATGAAGGAGGAAGTATATCTTGCAGGTAAATATTAATAATAACATATATGTATATATATATAAATATATGTATATATGTATATATTAATTTATATTGTATTATTATTTTCTCCTTTTTTTTTTTTAGTTTCAAGAGCTATTGGTGACAAAGACTTTAAATTTAATAATGTCATTTCAGCTACTCCTGATGTTATATGTAAAGAAATATATAGTGAAAAATGTAATGAGAAGAAAGAAGAAAAGGAAATTACAAAACGTACTAATGTTATTGAAAAAAGCGATATTGATGAAAATTATTTTAAGGAGACTGATAATTTATATTATTCAGCTCACGAAGTGAATTATCATTATGTTGTTATGGCATGTGATGGAGTATGGGATACCATGTCTAATAAGGTACATACGAATCTGTTAATGAGTACAATTATGTCACATATATATATATATATATATATTTTTATTTATTTATGTTTATTTTTTTTTCCATTTCATTTTATATTTTTTATAGGATATCGTTAAAATATTACAAACATATAATAACGATCCGGATAAAGCATGTAGCGAAATAATAAAAACGGCTTATGCCTATGGTAGTCAGGTACATTGAAATGTAAATAAATCACATATATATATATATGTAATATATATGTATATTTATTCATTTATATGTCTATAATTAATATGAACTTATCCATATAATTTGATTTGTTTTAATTTATTTCTGTAGGATAATTTAACCGCTATGTTGCTCAAATTTTATTGAAATCATATTCTTATTTTATTTTTTTTATTTTTTATATTCATCCTTATTTTTTAAAAATCACTACGATTAATATATATATATTTTACATTATGTTTTACCCTCTGTTTATATGTCTAATTTTTTTGGTGATATTTTTCTCACACATTTACCAGAGATGTGATTCAGTTATATATATATATATATATAAATTTATATATATGATGTTTGTGATTCATTTATTATAAGGAATAAATAAATATTTCCTAATCATTGATATATTTATTCATATTTTTATTTAACCAAAGAAATATTCTAATGTTATTTTATTTTATCCTATTTTTTTCATTATTAATTTTTATTTATTTATTTATTTATTTATTACCATTTATTGTTCTGATTC

**5’ amplicon size: 1344bps**

**3’ amplicon size: 1225bps**
